# Genomic Distortion of Jawed Vertebrate Phylogeny

**DOI:** 10.64898/2026.06.28.735080

**Authors:** Chase D. Brownstein, Liandong Yang, Alex Dornburg, Thomas J. Near

## Abstract

Reconstructing patterns of evolution requires understanding the interrelationships of species, yet evolutionary relationships that defy resolution and calibration in time are commonplace across the Tree of Life. Here, we investigate the dynamics of temporal and topological uncertainty by generating a phylogeny of jawed vertebrates using 1105 exonic loci sampled for 540 species spanning all major orders and most families of gnathostomes. Across loci and DNA sequence sites, we observe rapid reductions in statistical support for the monophyly of jawed vertebrate clades that originated around the Cretaceous-Paleogene mass extinction. Phylogenetic signal was scrambled to different degrees during rapid successive divergences in multiple unrelated jawed vertebrate lineages that radiated in this interval, including birds, snakes, placental mammals, and acanthomorph fishes. In addition to showing that particular events have modified phylogenetic signal across the same loci in distantly related vertebrate clades, we also demonstrate how rates of genomic evolution affect our ability to infer the timescale of vertebrate evolution. By testing how the inclusion of lineages of ray-finned fishes with very fast and slow rates of molecular evolution changes inferences of the vertebrate evolutionary timescale, we show that the deepest divergences in ray-finned fishes may be impossible to accurately infer using sequence data and calibrations from a limited fossil record. These results hint at the macroevolutionary realities underlying topological and divergence time uncertainty across evolutionary trees.

## Introduction

Phylogenetics has emerged as a foundational framework that unifies the biological sciences (Garland et al. 2005; Cavender-Bares et al. 2009; Dunn et al. 2014; Smith et al. 2020). The widespread application of genomic sequencing to non-model organisms has enabled unprecedented reconstruction of evolutionary relationships across the Tree of Life, including illuminating the origins of animals (Dunn et al. 2008, 2014; Simakov et al. 2022; Schultz et al. 2023) and vertebrates (Delsuc et al. 2006, 2018; Smith et al. 2013; Zhu et al. 2021; Marlétaz et al. 2024; Yu et al. 2024), and the phylogenetic relationships of the major clades of flowering plants (Zhao et al. 2021a; Zuntini et al. 2024), arthropods (Giribet and Edgecombe 2019; Su et al. 2024), mammals (Foley et al. 2023), birds (Stiller et al. 2024), and fishes (Thompson et al. 2021; Parey et al. 2023). Despite the increasing use of genome-scale data to tackle phylogenetic problems, considerable uncertainty persists surrounding the relationships of several species-rich lineages (Song et al. 2012; Jarvis et al. 2014; Prum et al. 2015; Esselstyn et al. 2017; Feng et al. 2017; Hughes et al. 2018; Friedman et al. 2019; Giribet and Edgecombe 2019; Kimball et al. 2019; Dornburg and Near 2021; Kuhl et al. 2021; Ghezelayagh et al. 2022; Mongiardino Koch et al. 2022, 2023; Álvarez-Carretero et al. 2022; Budd and Mann 2023; Foley et al. 2023; Glass et al. 2023; Su et al. 2024). In many lineages, such as neoavian birds or placental mammals, significant improvement in phylogenetic resolution is only accomplishable using tens or hundreds of thousands of loci, which necessitates the use of analyses targeting lineage-specific conserved regions (Foley et al. 2023; Stiller et al. 2024). Different types of genomic markers may result in the inference of alternative phylogenetic relationships within lineages, reflecting the varied nature of genomic ancestry and the different histories of different genes and syntenic blocks (Alda et al. 2021; Arcila et al. 2021; Foley et al. 2023; Mongiardino Koch et al. 2023; Stiller et al. 2024; Brownstein and Near 2026). For inferences at wider taxonomic scales, the assembly of such large genomic datasets is difficult owing to the ancient divergence of sampled species.

Regions of phylogenetic uncertainty frequently correspond to macroevolutionary events, especially mass extinctions, when trees are calibrated in time (Jarvis et al. 2014; Prum et al. 2015; Feng et al. 2017; Foley et al. 2023; Stiller et al. 2024; Somogyi et al. 2026). Although temporal patterns of phylogenetic discordance have been investigated within specific clades, including birds, mammals, and ray-finned fishes (Song et al. 2012; Jarvis et al. 2014; Prum et al. 2015; Esselstyn et al. 2017; Reddy et al. 2017; Ghezelayagh et al. 2022; Foley et al. 2023; Berv et al. 2024; Stiller et al. 2024), broader comparative analyses across the Tree of Life remain limited (Budd and Mann 2023). Consequently, the relative magnitude of phylogenetic uncertainty across distantly related clades and their correspondence to shared macroevolutionary phenomena remain poorly understood (O’Leary et al. 2013; Crouch et al. 2019). Moreover, lineage-specific biological attributes may influence both phylogenetic resolution and time-calibration of phylogeny (Sanderson 2002; Bromham and Woolfit 2004; Drummond et al. 2006; Lepage et al. 2007; Dornburg et al. 2012; Berv and Field 2018; Budd and Mann 2023; Berv et al. 2024; Duchêne et al. 2025). As phylogenetics enters an era of increased genome coverage for organismal diversity, data are now available to infer interactions among the structure of phylogenies and patterns of life history, ecology, and physiology in deep time (Berv and Field 2018; Berv et al. 2024).

Jawed vertebrates (*Gnathostomata*) comprise over 70,000 species present in nearly every ecosystem on the planet and are perhaps the best-studied of all animals. However, phylogenetic relationships among several jawed vertebrate clades remain contentious, a factor that for many lineages is often attributed to rapid diversification following major extinctions and biotic turnovers (Meredith et al. 2011; Song et al. 2012; Jarvis et al. 2014; Chen et al. 2015; Prum et al. 2015; Esselstyn et al. 2017; Irisarri et al. 2017; Hughes et al. 2018; Hime et al. 2020; Meyer et al. 2021; Álvarez-Carretero et al. 2022; Foley et al. 2023; Parey et al. 2023; Berv et al. 2024; Schartl et al. 2024; Stiller et al. 2024). No study has attempted to compare levels of phylogenetic uncertainty across a comprehensive sample of gnathostomes to assess global patterns of phylogenomic discordance through time in the three major gnathostome divisions, *Chondrichthyes* (cartilaginous fishes), *Actinopterygii* (ray-finned fishes), and *Sarcopterygii* (lobe-finned fishes), nor have previous analyses compared levels of phylogenetic discordance in classic vertebrate radiations such as birds, placental mammals, and acanthomorph fishes using the same genomic sequence data.

Here, we infer a phylogeny of 540 jawed vertebrates from 117 of 133 order-level (Foley et al. 2023; Near and Thacker 2024; Stiller et al. 2024) clades using a conserved set of 1,105 exonic loci (Hughes et al. 2018, 2021) to test whether directly comparable levels of phylogenetic uncertainty exist across these classic post-extinction adaptive radiations. Originally designed to circumvent the paralogy introduced by the teleost-specific whole genome duplication and interrogate ray-finned fish relationships (Hughes et al. 2018), these exons allow us to examine common patterns of phylogenetic discordance in a genomic marker set conserved across jawed vertebrates. Jawed vertebrate phylogeny, when calibrated in time, reveals tree-wide signatures of phylogenetic uncertainty and lineage origination in distantly related lineages of jawed vertebrates that correspond to major extinctions and biotic revolutions, especially the K-Pg event. We also show that biological variation in rates of protein-coding gene evolution make inferring the ages of some of the most species-rich clades of jawed vertebrates untenable. This suggests that the ages of some branches on the tree of life may not be accurately constrained past the minimum ages provided by unambiguous extinct representatives from the fossil record. These dual patterns of uncertainty in phylogenetic space and geological time have sculpted the shape of vertebrate phylogeny and prescribed the limits of what we can learn about jawed vertebrate evolutionary history from genomic data.

## Methods

### (a) Sequence Dataset Assembly

Constructing a comprehensive jawed vertebrate phylogeny requires accounting for hidden paralogs in ray-finned fishes (several lineages underwent independent whole-genome duplication episodes) when creating alignments. As such, we used the set of 1,105 exon loci published by Hughes et al. (Hughes et al. 2018, 2021). as a basis for producing a genome-wide marker sequence dataset to investigate jawed vertebrate phylogenetic relationships. Although we recognize that tens of thousands of markers may be necessary to resolve the relationships of clades in rapid radiations such as neoavian birds (Stiller et al. 2024), our interests were in checking for the presence of similar patterns of phylogenetic discordance across a broad sample of jawed vertebrates. We pooled data from Hughes et al. (Hughes et al. 2018, 2021) with sequences extracted using available whole genome assemblies on NCBI for non-actinopterygian vertebrates, including 13 chondrichthyans, the coelacanth *Latimeria chalumnae*, two lungfishes, 11 amphibians, 69 mammals, 39 lepidosaurs, 22 turtles, four crocodylians, and 60 birds, as well as 20 additional teleosts and holosteans for a total of 542 tips representing 540 species (see Brownstein et al. (Brownstein et al. 2023b, 2024a) for the inclusion of all seven living species of gars). The dataset represents a more than five-fold increase in sampling relative to previous attempts to reconstruct jawed vertebrate phylogeny, which sampled between 58 and 100 taxa and did not sample all of the deepest divergences in ray-finned fishes (Chen et al. 2015; Irisarri et al. 2017; Hime et al. 2020). Exon sequences were located in available genomes using HMMER 3.1 (Wheeler and Eddy 2013), extracted using the custom python scripts from Hughes et al. (Hughes et al. 2018; Brownstein et al. 2024a), and finally aligned using MAFFT 7.3 (Katoh and Standley 2013). Together, the exon sequence dataset included 729935 bp with a mean length of 661 bp per exon. We also used *phyluce* 1.7.3 (Faircloth 2016) to produce a 75% complete taxon-sequence matrix in addition to the original 1,105 exons; this subsampling step retained 662 (60% of the total) exon loci. We also used *CIAlign* (118) to identify and remove chimaeric alignments. Table S1 includes information on all sequences extracted from genomes available on the NCBI repository GenBank. For the 1,105 exon dataset, the total and average number of parsimony informative, singleton, and invariant sites were 53824 and 1922.29, 9364 and 302, and 56066 and 1808.6, respectively. For the 662 exon dataset, the total and average number of parsimony informative, singleton, and invariant sites were 26844 and 2237, 4099 and 341.58, and 23312 and 1943.66, respectively.

### (b) Maximum Likelihood Phylogenetic Analyses and Nodal Support

We inferred phylogenies using the original and 75% complete datasets using both concatenation and multispecies coalescent approaches. First, we used IQ-TREE 2 (Nguyen et al. 2015; Minh et al. 2020b) to infer maximum likelihood phylogenies from the concatenated exon sequence datasets. In each case, we used ModelFinder (Kalyaanamoorthy et al. 2017) to select best-fit models of sequence evolution and calculated ultrafast bootstraps over 1000 replicates. For multispecies coalescent inference, we inferred gene trees for exons in IQ-TREE2 using ModelFinder (Kalyaanamoorthy et al. 2017) to find best-fit models, and then used ASTRAL-III (Zhang et al. 2018) to infer a multispecies coalescent tree from the individual gene trees. In addition to ultrafast bootstrap supports, we further quantified statistical support for individual branches along the inferred phylogenies by inferring gene and site concordance factors for the phylogenies inferred from the original and 75% complete exon matrices. Gene and site concordance factors measure the number of decisive gene trees and sites, respectively, that are congruent with an input species tree topology (in this case, the concatenated trees were used as input trees) (Minh et al. 2020a). As such, concordance factors can provide information on patterns of support across two different levels of information in the genome: genes and sites. We conducted all maximum likelihood phylogenetic analyses and concordance factor calculations on the Yale High Performance Computing Cluster McCleary. We examined associations between different levels of support and compared gene and site concordance factors, multispecies coalescent support values, and ultrafast bootstrap values for clades in the inferred phylogenies using heatmaps and scatterplots built using custom scripts in the R package *ggplot2* (Valero-Mora 2010).

### (c) Anomaly Zone Detection

We further interrogated statistical support for branches along the inferred phylogenies by inferring the position of anomaly zones along the ASTRAL species trees. Anomaly zones (AZs) are regions of species trees where an alternative topology is supported more strongly in a handful of gene trees than the consensus topology is in the species tree (Degnan and Rosenberg 2006, 2009). The presence of anomaly zones in regions of a tree is usually taken to indicate rapid successive divergences or high effective population sizes (Degnan and Rosenberg 2006, 2009; Liu and Edwards 2009; Linkem et al. 2016; Chafin et al. 2021; Brownstein et al. 2024b). We used custom python scripts(Linkem et al. 2016; Chafin et al. 2021) to check how many anomalous bipartitions included each branch in the input ASTRAL-III species trees generated using the original and 75% complete exon matrices. Branches that were present in more than 10 anomalous bipartitions were considered to be in anomaly zones.

### (d) Time Calibration of Jawed Vertebrate Phylogeny

In order to produce a time-calibrated phylogeny of jawed vertebrates, we conducted Bayesian node-dating analysis in BEAST 2.6.7 (Bouckaert et al. 2014, 2019) on three sets containing the 150 largest exon sequences from the 75% complete dataset for 375 representative species. For time-calibration, we fixed the phylogeny by pruning the concatenated maximum likelihood tree generated from the 75% complete matrix. We placed minimum bounds on nodes in the tree using 52 fossil calibrations. Because fossil calibrations can extensively change our understanding of the timescale of evolution, even fossils that have been allied with crown groups in phylogenetic analyses should be rigorously evaluated (Parham et al. 2012; Ksepka et al. 2015; Whiteside et al. 2022; Brownstein et al. 2023a). As such, we vetted the phylogenetic placement of included fossil calibrations through a comprehensive literature review and phylogenetic analysis of morphological character matrices when character states uniting taxa used as fossil calibrations with the relevant crown clades in jawed vertebrates were not provided (Brownstein et al. 2022, 2023a; Brownstein 2023b; Brownstein and Near 2024). This approach allowed us to supply rigorous justification of fossil calibration placement based on the observation of phylogenetically optimized character states. For each calibration, we also searched the literature for penecontemporaneous occurrences of equivalent calibrations, such that in many cases our fossil calibrations are based on multiple coeval extinct taxa or would only need slight modification in cases where the phylogenetic position of the principal fossil calibration is revised. To this extent, we reran the time-calibration analyses with a modified set of fossil calibrations to test for whether the exclusion of a handful of somewhat controversial mammal fossils affected the ages that we inferred (Table S2). A full list of fossil calibration justifications, including lists of optimized character states that unite extinct species used as fossil calibrations with crown clades in jawed vertebrates, is included in the Supplementary Material, following best practice recommendations (Parham et al. 2012).

For the root node, we set the origin prior to 443.0 Ma, the earliest Silurian, which is the earliest age for potential jawed vertebrate material from the fossil record (The distribution of the vertebrates in the Late Ordovician and Early Silurian palaeobasins of the Siberian Platform 1995). We set the bounds on the root prior to 439.0 Ma, the age of the oldest known unambiguous crown-group jawed vertebrate †*Fanjingshania renovata* (Pan-*Chondrichthyes*)(Andreev et al. 2022) and 509.0 Ma, the lower bound of the Cambrian Series 2. This is the maximum age of the Burgess Shale, which contains several fossil chordates but no crown vertebrates (Morris and Caron 2012, 2014; Mussini et al. 2024). For each fossil calibration, we inputted bounds on the prior distribution of specified clades such that 97.5% of the distribution fell before the age of the fossil used. Although maximum age bounds have been proposed for some nodes calibrated in this phylogeny, we chose to sculpt prior distributions in this manner to account for uncertainty regarding early fossil occurrences following previous studies (Prum et al. 2015; Ghezelayagh et al. 2022; Brownstein et al. 2024b, 2025), which should in any case postdate the origination of crown clades. We placed the fossil calibrations on the tree using monophyletic MRCA priors (Table S3).

We used a General Time Reversible model of sequence evolution, a Relaxed Lognormal Clock Model, and the BEAST2 implementation of the Fossilized Birth-Death (FBD) Model. We set the diversification rate to 0.025, which was found by rounding from the approximation found using the equation in Magallón and Sanderson (Magallón and Sanderson 2001) using the input values of 439.0 Ma for the clade age and 75,681 (the current sum of jawed vertebrate species as of February 2025 taken from Eschemeyer’s Catalogue of Fishes, the Reptile Database, The Clements Checklist of Birds of the World, the ASM Mammal Diversity Database, and AmphibiaWeb) for the species count. Finally, we ran each exon set three times independently over 4.0 x 10^8^ generations with a 1.0 x 10^8^ pre-burnin, checked the log files for convergence of the posteriors and ESS values >200 using Tracer v 1.7 (Rambaut et al. 2018), combined the top 5% of trees from the combined set of nine runs (=3.6 billion generations) using LogCombiner v.2.6.6 (Bouckaert et al. 2019), and summarized the posterior trees in a single tree with common ancestor heights annotated onto the input concatenated maximum likelihood topology. For each exon set, we also produced trees with median node heights annotated onto the maximum likelihood topology to check the similarity of divergence times estimated from different exon sets.

### (e) Analyses of Support Through Time

We matched gene and site concordance values, ultrafast bootstrap support values, and branch lengths in substitutions per site from the 75% concatenated maximum likelihood phylogeny to the corresponding branches in the time-calibrated phylogeny generated from pooling posterior tree sets from the nine independent node-dating runs to assess for patterns of change in phylogenetic support over deep time. Similarly, we assessed how the length of branches in millions of years changed through time to see whether certain intervals of jawed vertebrate evolutionary history have featured a higher number of short branching events and replicated this analysis across trees generated from each of the three exon sets. We used custom scripts in R to plot all these values through time (Valero-Mora 2010). Next, we subsampled nodes to only those with median ages within 10 million years of the K-Pg boundary (76.02 to 56.02 Ma) and explored variance in node support across ecological categories, clades, and evolutionary radiations. We determined whether nodes in this subsample belonged to classic post-extinction radiations, such as birds (Hackett et al. 2008; Jarvis et al. 2014; Brusatte et al. 2015; Prum et al. 2015; Kuhl et al. 2021; Berv et al. 2024; Stiller et al. 2024), acanthomorphs (Friedman 2010; Near et al. 2013; Friedman et al. 2019; Ghezelayagh et al. 2022), some otophysan fish orders (*Cypriniformes*, *Siluriformes*)(Kappas et al. 2016; Bagley et al. 2018; Roxo et al. 2019; Day et al. 2023), placental mammal orders (Meredith et al. 2011; dos Reis et al. 2012, 2014; Phillips and Fruciano 2018; Upham et al. 2021; Álvarez-Carretero et al. 2022; Foley et al. 2023), and snakes (Burbrink et al. 2020; Klein et al. 2021), and compared concordance factors for these nodes to those that appeared in the same time period but are not associated with classic adaptive radiations. For ecological characterization, we based our scorings on presumed ancestral ecology. For example, the nodes representing the MRCA of penguins (*Spheniscidae*) with other waterbirds were coded as ‘flighted’ since penguins are secondarily flightless (Cole et al. 2022).

### (f) In silico Divergence Time Estimation Experiments

To explore how lineage-specific rates of molecular evolution might affect our inferences of the timescale of jawed vertebrate evolution, we conducted *in silico* taxon and fossil calibration inclusion and exclusion experiments on a subset of 45 species of non-tetrapod sarcopterygians and actinopterygians sampling all major lineages in lungfishes, coelacanths, and ray-finned fishes (*Polypteridae*, *Acipenseriformes*, *Holostei*, *Oseanacephala, Otocephala*, and *Euteleostei*). We selected this region of the tree for *in silico* divergence time testing because of the known heterogeneity in rates of substitution per site among the first four divergences in ray-finned fishes (Takezaki 2018; Brownstein et al. 2024a) and the long ghost lineages for these divergences implied by the ages inferred by previous molecular phylogenies (Near et al. 2012; Betancur-R et al. 2013; Giles et al. 2017; Hughes et al. 2018), which suggest that the earliest known fossil actinopterygians, actinopterians, neopterygians, and teleosts postdate the origins of these clades by up to 100 million years (Hurley et al. 2006; Giles et al. 2017; Friedman 2022). For all tests on the subset of 45 taxa, we ran Bayesian node-dating analysis in BEAST2 on two of the three 50 exon sets over 1.0 x 10^8^ generations with a 1.0 x 10^8^ pre-burnin, checked the log files for convergence of the posteriors and ESS values >200 using Tracer v 1.7 (Rambaut et al. 2018), combined the top 5% of trees from the combined set of nine runs (=3.6 billion generations) using LogCombiner v.2.6.6 (Bouckaert et al. 2019), and summarized the posterior trees in a single tree with median node heights annotated onto the input concatenated maximum likelihood topology. We analyzed the following permutations to the 45 taxon dataset:

(1) The full 45 taxon dataset, with all relevant fossil calibrations included, to provide a base of comparison.

(2) A reduced 40 taxon dataset, excluding holosteans (gars and bowfins) and acipenseriforms (sturgeons and paddlefishes), to test whether the exclusion of living fossil ray-finned fish lineages with slow rates of nucleotide substitution modifies the inferred ages of crown *Actinopterygii* and crown *Teleostei*.

(3) The full the 45 taxon dataset, but with fossil calibrations providing hard constraints on the ages of the lepisosteid and acipenseriform crown groups removed. Because these fossil calibrations, †*Atractosteus falipoui* (Grande 2010) and †*Protopsephurus liui* (Grande et al. 2002) have been reliably placed within the gar and acipenseriform crown groups in numerous phylogenetic analyses of morphological characters (Grande et al. 2002; Grande 2010; Hilton et al. 2011, 2023; Brownstein 2022; Brownstein and Lyson 2022; Brownstein et al. 2023b), and because many similarly old or slightly geologically younger crown gars and acipenseriforms are known from complete skeletons (Grande and Bemis 1991; Grande 2010; Hilton et al. 2011, 2023; Brito et al. 2017; Sato et al. 2018; Brownstein and Lyson 2022; Brownstein et al. 2023b; Murray et al. 2023), appreciably younger ages for the gar and acipenseriform crown groups found when these fossil calibrations are excluded would conflict directly with a well-established fossil record and provide support for the hypothesis that slow and fast rates of nucleotide substitution confound inference of the ages of the deepest divergences in ray-finned fishes without minimum age constraints from fossils.

We then recorded the median and 95% highest posterior density bounds for divergence times and branch-wise rates of substitution across different analyses and plotted them together (Table S4). We compared these rate and divergence time values to those of the time-calibrated phylogeny built using all three exon sets and the 375 species set. Finally, we used the application FigTree (http://tree.bio.ed.ac.uk/software/figtree/) to visualize median rates of nucleotide substitution across the 375 taxon phylogenies made using each and all of the three exon sets and the 40 and 45 taxon phylogenies built using two of the three exon datasets.

## Results

### (a) Concordance and discordance across jawed vertebrate phylogeny and in time

We assembled a dataset of 1,105 orthologous exon loci (Hughes et al. 2018, 2021; Brownstein et al. 2024a) for 542 jawed vertebrates representing all major divisions (e.g., *Chondrichthyes, Polypteridae, Acipenseriformes, Holostei, Oseanacephala, Clupeocephala, Lepidosauria, Amphibia, Mammalia, Squamata, Testudines, Crocodylia,* and *Aves*) and 88% of orders, inferred phylogenies using multispecies coalescent (Zhang et al. 2018) and concatenated maximum likelihood (Minh et al. 2020b) approaches, and inferred a time-calibrated evolutionary tree under a Uncorrelated Relaxed Log-Normal clock and General Time Reversible model in a Bayesian framework (Bouckaert et al. 2014, 2019) using 52 fossil calibrations rigorously vetted using optimized character states from phylogenetic analyses of morphological characters (Supplementary Information).

Our phylogeny (Figure 1; Figure 2) compares favorably to previous large-scale phylogenomic analyses of jawed vertebrate clades in resolving several relationships that conflict with traditional morphology-based phylogenies, including the resolution of a clade containing odd-toed ungulates, carnivoran mammals, and pangolins (Song et al. 2012; Esselstyn et al. 2017; Álvarez-Carretero et al. 2022; Foley et al. 2023), the placement of turtles as the living sister to archosaurs (birds and crocodylians)(Cao et al. 2000; Chiari et al. 2012; Crawford et al. 2012; Wang et al. 2013; Chen et al. 2015; Hime et al. 2020), the placement of geckos as the earliest diverging clade of squamates (Pyron et al. 2013; Zheng and Wiens 2016; Streicher and Wiens 2017; Burbrink et al. 2020; Simões and Pyron 2021; Singhal et al. 2021; Title et al. 2024), the resolution of a clade containing snakes, iguanian, and anguimorph lizards (Pyron et al. 2013; Zheng and Wiens 2016; Streicher and Wiens 2017; Burbrink et al. 2020; Simões and Pyron 2021; Singhal et al. 2021; Title et al. 2024), and the resolution of the first four diverging clades of ray-finned fishes (Grande 2010; Near et al. 2012; Betancur-R et al. 2013; Hughes et al. 2018, 2021; Dornburg and Near 2021; Thompson et al. 2021; Near and Thacker 2024): bichirs and Reedfish (*Polypteridae*), sturgeons and paddlefishes (*Acipenseriformes*), gars and bowfins (*Holostei*), and all other ray-finned fishes (*Teleostei*) (Figure 1, Figure 2).

**Figure 1.**
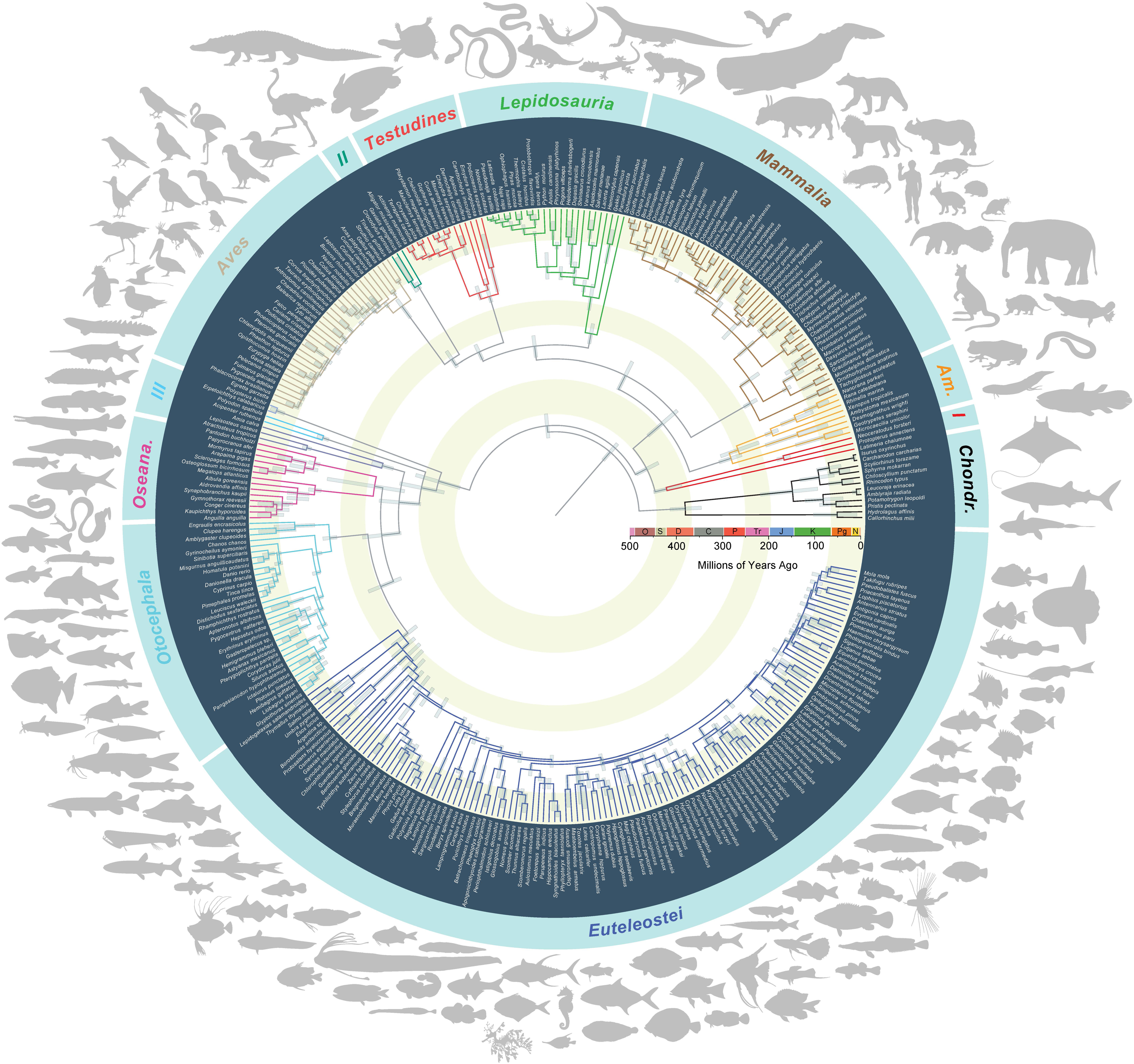
Time-Calibrated Phylogeny of Jawed Vertebrates. Radially projected Bayesian node-calibrated phylogeny of 375 representative species of jawed vertebrates based on the topology inferred for 1105 concatenated exon loci sampled for 542 species under a maximum likelihood criterion in IQ-TREE2. The phylogeny was time-calibrated in BEAST 2.6.7 using 52 vetted fossil calibrations justified in the Supplementary Information. Bars at nodes indicate 95% highest posterior density (HPD) intervals for divergence times. Major vertebrate clades are noted around the phylogeny, and abbreviations are as follows, counterclockwise: *Chond*., *Chondrichthyes*; *I*, *Latimeria-Dipnoi* clade; *Am*., *Amphibia*; *II*, *Crocodylia*; *III*, non-teleost actinopterygians, including *Polypteridae*, *Acipenseriformes*, and *Holostei*; *Oseana*., *Oseanacephala*.(Near and Thacker 2024) Borders between tan and white concentric regions denote the timing of the ‘big five’ mass extinctions: Ordovician-Silurian, End-Devonian, Permo-Triassic, End-Triassic, and Cretaceous-Paleogene. Silhouettes are public domain from phylopic.org or by CDB.

**Figure 2.**
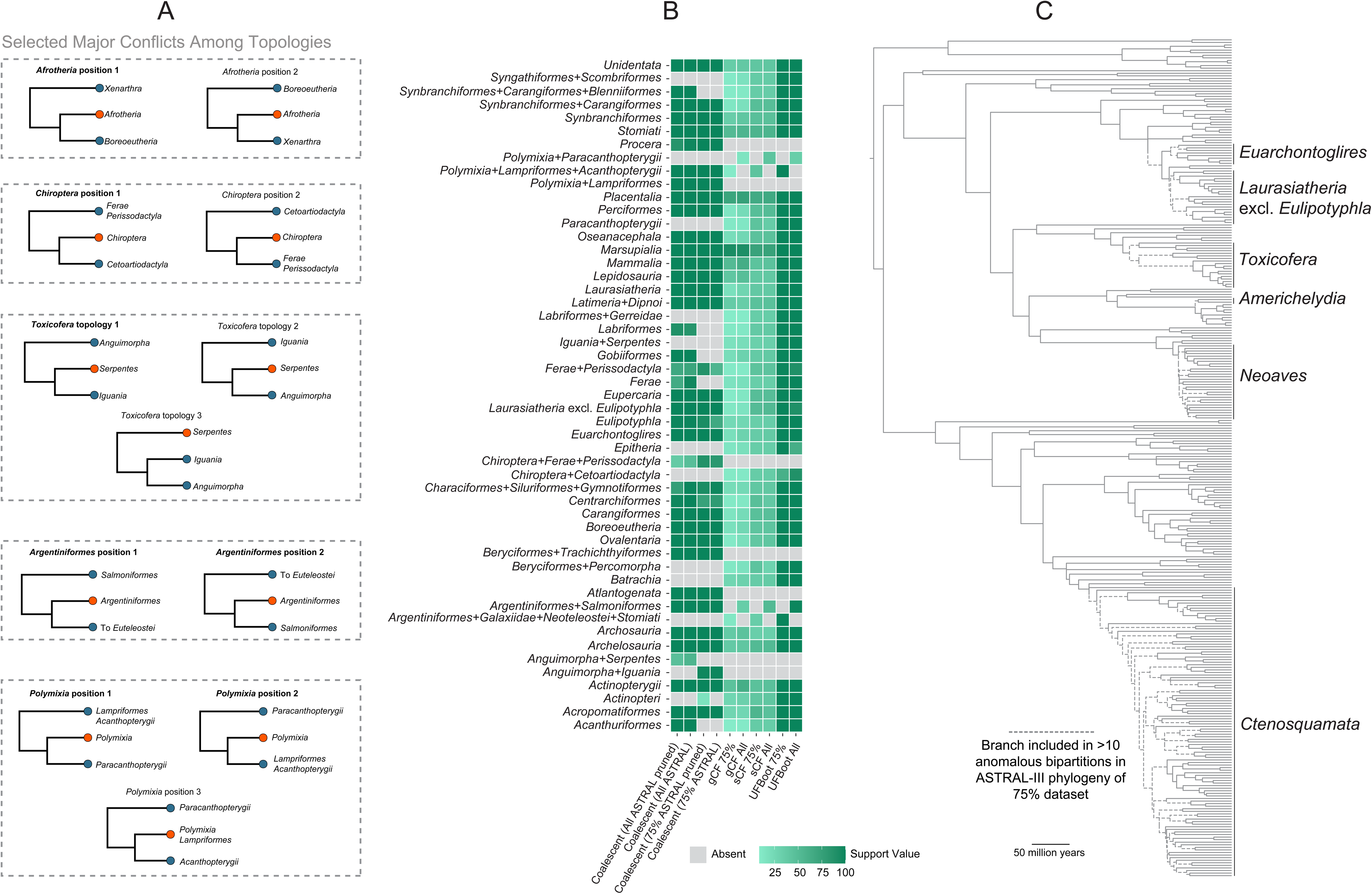
Discordance Across Jawed Vertebrate Phylogeny. (A) Selected major conflicts among trees built in this and previous(Song et al. 2012; dos Reis et al. 2014; Esselstyn et al. 2017; Hughes et al. 2018; Burbrink et al. 2020; Singhal et al. 2021; Ghezelayagh et al. 2022; Foley et al. 2023; Title et al. 2024) phylogenies built using genome-wide sequence data. (B) Heatmap of support values across different metrics for selected clades. gCF, gene concordance factor; sCF, site concordance factor; UFBoot, ultrafast bootstrap. (C) Time-calibrated phylogeny shown in Figure 1 annotated with the corresponding inferred locations of anomaly zones on the ASTRAL-III species tree, indicated by dashed lines.

In our analyses, osteoglossomorph (mooneyes, mormyrid elephantfishes, and arowanas) and elopomorph (eels and tarpons) fishes are found to comprise a lineage, *Oseanacephala* (Near and Thacker 2024), that is the sister to all other teleosts. This clade was only recently supported using data on genome structure evolution (Parey et al. 2023) and some (though not all) (Hughes et al. 2018, 2021) whole-genome-based phylogenies (Bian et al. 2016; Hao et al. 2020; Takezaki 2021). We also resolve a clade, *Stomiati* (Betancur-R et al. 2013; Hughes et al. 2018; Near and Thacker 2024), containing the deep-sea dragonfishes and relatives (*Stomiiformes*) and the smelts (*Osmeriformes*), reject the monophyly of the traditional *Characiformes* (containing citharinoid fishes)(Chakrabarty et al. 2017; Melo et al. 2022), and resolve the major lineages containing the over 18,900 species in percomorph fishes. In order of divergence, these are cusk eels and brotulas (*Ophidiiformes*), toadfishes (*Batrachoididae*), gobies and cardinalfishes (*Gobiiformes*), a clade containing seahorses and relatives (*Syngnathiformes*) and tunas and relatives (*Scombriformes*), and all other percomorphs, which form two major lineages (Ghezelayagh et al. 2022; Near and Thacker 2024). *Eupercaria* contains five major groups (Figure 1, Figure 2): darters, true perches, groupers, icefishes, sculpins, rockfishes, sea lions, snailfishes, sticklebacks, searobins, and eelpouts form the *Perciformes*, the sister lineage of all other *Eupercaria*. The other eupercarian clades are wrasses, parrotfishes, and stargazers (*Labriformes*), wreckfishes and deep-sea cardinalfishes (*Acropomatiformes*), black basses, sunfishes, flagtails, and relatives (*Centrarchiformes*), and a bizarre lineage containing anglerfishes, pufferfishes and *Mola mola*, butterflyfishes, seabasses, surgeonfishes, drums, and a host of other marine and especially reef-associated clades (*Acanthuriformes*). Sister to *Eupercaria* is an unnamed clade consisting of four orders: jacks, swordfishes, flatfishes, and relatives (*Carangiformes*), swamp eels and betta fishes (*Synbranchiformes*), the flyingfishes, livebearers, halfbeaks, ricefishes, New World silversides, and relatives (*Atheriniformes*), and the ‘*Blenniiformes*,’ a grade comprising species-rich families like cichlids, damselfishes, surfperches, and blennies (Figure 1; Figure 2) (Ghezelayagh et al. 2022; Near and Thacker 2024).

Oddly, we find strong support across different analyses for a clade containing coelacanths and lungfishes to the exclusion of tetrapods (Figure 1, Figure 2), which contrasts with many (Chen et al. 2015; Irisarri and Meyer 2016; Irisarri et al. 2017; Bi et al. 2021; Meyer et al. 2021; Wang et al. 2021; Schartl et al. 2024), but not all (Shan and Gras 2011), phylogenies built using large DNA sequence datasets. Many previous studies evaluating the relationships of coelacanths, lungfishes, and tetrapods did not sample the major early-diverging ray-finned fish clades (Irisarri and Meyer 2016; Meyer et al. 2021; Wang et al. 2021; Schartl et al. 2024). Our inclusion of representatives of the major chondrichthyan and actinopterygian clades means that we fully sample the initial divergences among living jawed vertebrates, which might partially account for this result. High-resolution stratigraphic columns and an appreciably complete fossil record from China show that the stem lineages of coelacanths, lungfishes, and tetrapods originated in rapid succession in the Early Devonian (Zhao et al. 2021b; Cui et al. 2022), which could lead to a lack of clarity about the relationships of these lineages (Degnan and Rosenberg 2006, 2009; Irisarri and Meyer 2016; Reddy et al. 2017; Berv et al. 2024). As such, this uncertainty might conceivably be due to the absence of phylogenetic signal retained in the initial rapid radiation of lobe-finned fishes, in addition to the inability for even over 1000 exon loci to provide clarity surrounding the relationships at the base of *Sarcopterygii*.

More importantly, our results demonstrate that the relationships of some of the most species-rich vertebrate clades, including neoavian birds (Hackett et al. 2008; Jarvis et al. 2014; Prum et al. 2015; Reddy et al. 2017; Kimball et al. 2019; Berv et al. 2024; Stiller et al. 2024), iguanians, anguimorphs, and snakes (Pyron et al. 2013; Zheng and Wiens 2016; Streicher and Wiens 2017; Burbrink et al. 2020; Singhal et al. 2021; Title et al. 2024), several interordinal relationships of placental mammals (Meredith et al. 2011; Song et al. 2012; Foley et al. 2016, 2023; Tarver et al. 2016; Esselstyn et al. 2017; Irisarri et al. 2017; Álvarez-Carretero et al. 2022), and euteleost and acanthomorph fishes (Near et al. 2012, 2013; Betancur-R et al. 2013; Hughes et al. 2018, 2021; Ghezelayagh et al. 2022) are all inferred with similar levels of uncertainty using the same genomic dataset. We investigated the factors hindering resolution of these clades’ relationships through multiple approaches (Figure 2A-C; Figures S1-S4), including comparing jawed vertebrate phylogenies using both multispecies coalescent and concatenation methods, evaluating gene and site concordance factors (Minh et al. 2020a; Kück et al. 2022), and identifying and inferring the location of anomaly zones (Degnan and Rosenberg 2006, 2009).

Concordance factors measure how many gene trees and sequence sites align with relationships in a species tree, while anomaly zones represent regions where a small number of gene trees strongly support alternative relationships compared to the consensus species tree. We find that phylogenetic discordance underlying the unresolved relationships of these species-rich clades (Figure 2A) is primarily due to a low number of gene histories that support a given topology (Figures S1-S4); many contentious relationships among acanthomorph and placental mammal clades, for example, are supported by only a handful of gene trees or none at all. Indeed, reducing the number of sampled exons to account for matrix incompleteness results in many branches in neoavian birds, placental mammals, and acanthomorph fishes collapsing (Figure 2B, Supplementary Information) despite receiving strong support in concatenated maximum likelihood phylogenies (Figure 1, Figure 2B) and in analyses using genome-wide data (Esselstyn et al. 2017; Hughes et al. 2018, 2021; Ghezelayagh et al. 2022; Foley et al. 2023). This supports the hypothesis that lineage-specific phylogenies that can incorporate a higher number of orthologs are necessary to resolve the relationships of clades such as neoavian birds (Stiller et al. 2024) that are entirely contained in anomaly zones in our jawed vertebrate phylogeny (Figure 2C).

When placed in a temporal context, these patterns of concordance and discordance across jawed vertebrate phylogeny are associated with major macroevolutionary events, especially the Cretaceous-Paleogene mass extinction 66.02 million years ago (Gradstein et al. 2021)(Figure 3; Figures S6-S9). Gene and site concordance factors, bootstrap and Bayesian posterior supports, and branch lengths all sharply decrease at the Cretaceous-Paleogene boundary before rebounding in the Cenozoic (Figure 3). These shifts correspond to an increase in the rapidity of successive divergences among many jawed vertebrate clades, including neoavian birds, several orders of placental mammals, and pelagic scombriform and carangiform fishes, around the Cretaceous-Paleogene boundary (Figure 1, Figure 3B; Figures S6-S9; Table S2).

**Figure 3.**
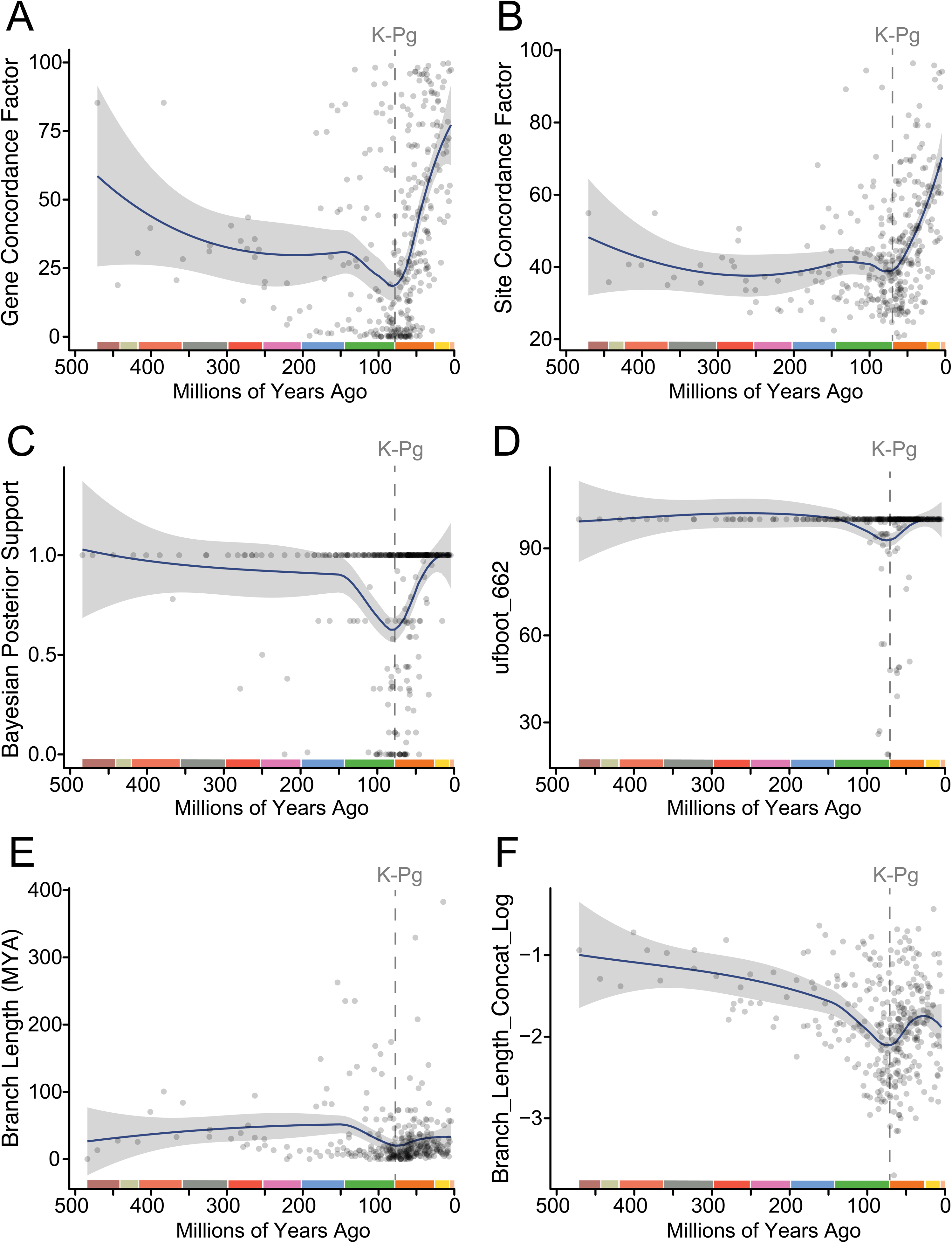
Signatures of Mass Extinction on Phylogenetic Resolution. Graphs show trendlines and 95% confidence intervals for several measures of nodal support (A-D) and branch length (E-F) through time. (A-C, F) are based on the concatenated maximum likelihood phylogeny built form the 75% complete matrix. (A) Gene concordance factors through time, (B) site concordance factors through time, (C) ultrafast bootstraps through time, (D) Bayesian posterior support through time, (E) branch length in millions of years through time, (F) branch length in substitutions per site through time. Dashed line indicates the Cretaceous-Paleogene Mass Extinction.

Although previous studies have found associations between phylogenetic discordance and rapid diversification following the Cretaceous-Paleogene boundary (Near et al. 2013; Jarvis et al. 2014; Prum et al. 2015; Alfaro et al. 2018; Friedman et al. 2019; Burbrink et al. 2020; Kuhl et al. 2021; Ghezelayagh et al. 2022; Foley et al. 2023; Glass et al. 2023; Berv et al. 2024; Stiller et al. 2024), these lineage-specific analyses do not allow for direct comparisons of phylogenomic discordance across jawed vertebrates as shown here. In addition to the decreases in branch lengths and phylogenetic support across genes and sites found across jawed vertebrate diversity at the K-Pg boundary (Figure 3), we also find that classic post-Cretaceous adaptive radiations recognized in the literature, including neoavian birds (Jarvis et al. 2014; Brusatte et al. 2015; Prum et al. 2015; Field et al. 2019; Kuhl et al. 2021; Berv et al. 2024; Stiller et al. 2024), various clades of acanthomorph fishes (Friedman 2010; Near et al. 2013; Alfaro et al. 2018; Friedman et al. 2019; Ghezelayagh et al. 2022; Brownstein et al. 2024b; Miller et al. 2024), and the interordinal diversifications of placental mammals (Meredith et al. 2011; Álvarez-Carretero et al. 2022; Foley et al. 2023), show moderately lower gene and site concordance factors than equivalent-age clades not associated with post-Cretaceous radiation (Figure 4). We also find a drop in phylogenetic support around the K-Pg extinction is concentrated in flighted, marine, freshwater-marine (euryhaline), and terrestrial clades; freshwater clades that diverge within 10 million years of the K-Pg are supported by comparatively high gene and site concordance factors (Figure 2, Figure 4). This result is consistent with the hypothesis that freshwater faunas are thought to have been the least affected by the K-Pg extinction (Sheehan and Fastovsky 1992; Robertson et al. 2013; Brownstein and Lyson 2022).

**Figure 4.**
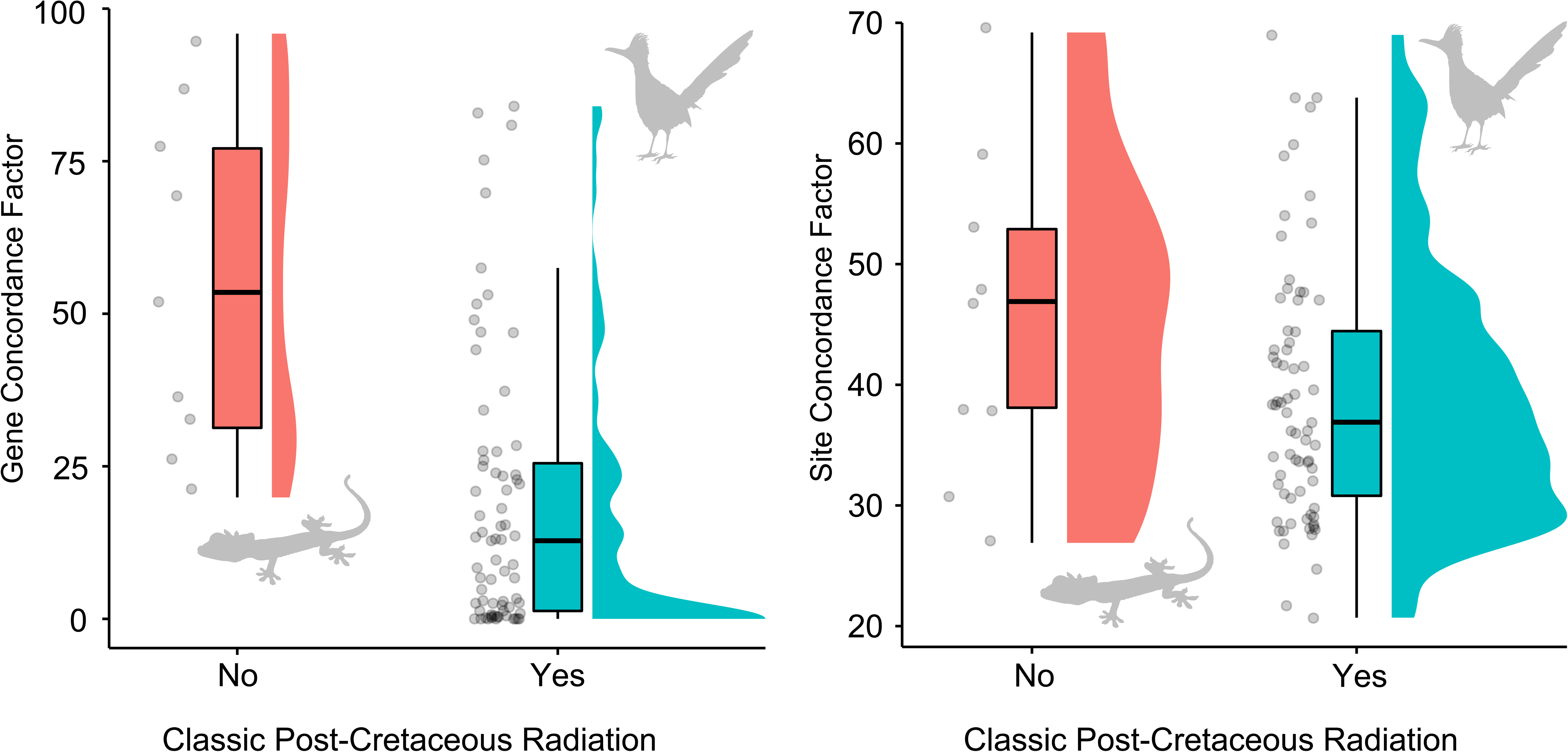
Classic Post-Cretaceous Evolutionary Radiations Show Increased Phylogenetic Discordance. Boxplot shows that, for nodes with ages falling within 10 million years of the K-Pg boundary (76.02 to 56.02 Ma), both (A) gene and (B) site concordance factors are on average lower in classic post-Cretaceous adaptive radiations, including neoavian birds, acanthomorph and placental mammal order-level clades, and snakes.

Our analysis reveals that mass extinctions are associated with sudden decreases in phylogenetic support across both genes and individual nucleotide sites throughout jawed vertebrate phylogeny (Figure 3). Because only a handful of lineages remain after a mass extinction and usually are represented by only small population sizes, rapid successive divergences into ecological opportunity created by mass extinctions in concert with the effects of bottleneck events might produce unclear evolutionary histories as genes and genomes experienced uneven recombination (Mirarab et al. 2024) fail to fix at the pace of rapid lineage divergence (Degnan and Rosenberg 2006; Suh et al. 2015; Reddy et al. 2017; Mirarab et al. 2024). Our analyses support the process of mass extinction and faunal recovery as a fundamental driver of discordance across the jawed vertebrate tree, which contains many of the classic examples of post-extinction evolutionary radiation (Jablonski 2001, 2005; Stroud and Losos 2016). By inducing sudden bottlenecks and opening up opportunities for lineage diversification as ecosystems reassemble, mass extinctions can scramble the genomic record of evolutionary history across diverse vertebrate lineages within a haze of rapid lineage origination, introgression, and repressed recombination.

### (b) Timing the jawed vertebrate radiation

Improved sampling of gnathostome clades allows us to chart jawed vertebrate diversification through time in detail. For example, our analyses strongly support a Mesozoic origin and diversification of placental mammal orders, followed by diversification within orders after the Cretaceous-Paleogene mass extinction (Springer et al. 2003; Meredith et al. 2011; dos Reis et al. 2012, 2014; Phillips and Fruciano 2018; Upham et al. 2021; Álvarez-Carretero et al. 2022; Carlisle et al. 2023; Foley et al. 2023), rather than diversification of both in the Cenozoic (Figure 1, Figure S3, 6)(Alroy 1999; Archibald and Deutschman 2001; Wible et al. 2007). The conservative placement of several of our node calibrations, including for *Carnivora* and *Arctoidea*, should skew posterior ages younger if crown clades originate in the Paleocene.

Furthermore, the estimated ages of placental mammal orders are robust to the use of different fossil calibration schemes that alternately exclude contentious and recently debated fossils for the setting of prior age distributions (Table S2).Thus, we remain confident in the inferred Cretaceous origin of placental order-level clades. Similarly, our results support the inference that three to five avian clades survived the Cretaceous-Paleogene extinction (Jarvis et al. 2014; Prum et al. 2015; Berv and Field 2018; Berv et al. 2024; Stiller et al. 2024) and soundly reject a Mesozoic diversification of crown birds (Figure 1, Figure S3, 6)(Jetz et al. 2012; Wu et al. 2024).

Our time-calibrated phylogenies using different sets of fossil calibrations and exonic loci also consistently estimate rapid Paleogene diversifications of several major ray-finned fish clades, including eels (*Anguilliformes*), minnows, carps, loaches, and suckers (*Cypriniformes*), catfishes (*Siluriformes*), opahs and oarfishes (*Lampriformes*)(Brownstein and Near 2024), tunas and mackerels (*Scombriformes*),(Friedman et al. 2019) jacks, swordfishes, and flatfishes (*Carangiformes*)(Ribeiro et al. 2018; Glass et al. 2023), anglerfishes (*Lophioidei*)(Brownstein et al. 2024b; Miller et al. 2024), and the orders of *Eupercaria* (Figure 1; Figures S4-6). Several deeply-divergent vertebrate clades also appear to have originated after mass extinctions. For example, the major reptile crown clades (*Lepidosauria* and *Archosauria*) and the amphibian crown group appear close to the Permo-Triassic mass extinction, which is consistent with the late Permian origins and rapid Triassic diversification of crown reptiles and amphibians implied by the fossil record (Marjanović and Laurin 2007; Anderson et al. 2008; Brusatte et al. 2010; Nesbitt 2011; Jones et al. 2013; Bernardi et al. 2015; Simões et al. 2018, 2022; Ezcurra et al. 2020; Sander et al. 2021; Simões and Pyron 2021; Kear et al. 2023; Kligman et al. 2023). Thus, the timescale of evolution for many jawed vertebrate clades matches expectations from the hypothesis that mass extinctions were key events in shaping living vertebrate diversity.

### (c) Some jawed vertebrate clade ages are highly unreliable

Our analyses reveal that molecular age estimates for certain clades remain unreliable even when hundreds of conserved loci are analyzed, particularly in groups with sparse fossil records. To investigate the patterns and underlying causes of age estimation inaccuracies across jawed vertebrate phylogeny, we conducted computational experiments examining how taxonomic sampling and fossil calibration selection influence the estimated ages of major sarcopterygian and actinopterygian crown clades (Figure 4). Divergence times for basal actinopterygian lineages estimated using molecular data consistently precede the oldest unambiguous crown-group fossils by 100 to 150 million years in time (Giles et al. 2017; Friedman 2022). This discrepancy has largely been attributed to a poor fossil record, and several putative members of the ray-finned fish crown have been described from the Paleozoic (Hurley et al. 2006; Argyriou et al. 2018, n.d.). However, this disagreement between the fossil record and molecular phylogenies could also plausibly stem from extreme heterogeneity in genomic evolutionary rates across the most deeply divergent vertebrate lineages. The earliest-diverging actinopterygian and sarcopterygian clades (lungfishes, coelacanths, gars, bowfins, sturgeons, and polypterids) are often considered to be ‘living fossils,’ due to their low species diversity, ancient (>250 million year) common ancestry with other vertebrates, and the observation that living species show few morphological differences from extinct relatives that lived tens of millions of years in the past (Darwin 1859; Stanley 1975; Schultze and Wiley 1984; Grande and Bemis 1991, 1998; Grande 2010; Hilton et al. 2011; Amemiya et al. 2013; Casane and Laurenti 2013; Braasch et al. 2016; Clarke et al. 2016; Brito et al. 2017; Giles et al. 2017; Lidgard and Love 2018; Cavin et al. 2021; Meyer et al. 2021; Brownstein et al. 2022, 2024a; Cui et al. 2022; Hilton and Grande 2023).

Several early diverging ray-finned fish clades, including gars and sturgeons, exhibit the slowest rates of nucleotide substitution among vertebrates (Braasch et al. 2016; Takezaki 2018; Du et al. 2020; Brownstein et al. 2024a), a phenomenon that may be linked to their capacity for successful hybridization between lineages separated by over 100 million years of common ancestry (Havelka et al. 2011; Bohn et al. 2017; Káldy et al. 2020; Taylor et al. 2020; Brownstein et al. 2024b). The juxtaposition of these exceptionally slow evolutionary rates with the accelerated rates characteristic of teleost fishes (Takezaki 2018; Brownstein et al. 2024a) can confound molecular clock models, leading to systematic errors in divergence time estimation (Dornburg et al. 2012). We investigated the impact of extreme molecular rate variation on divergence time estimation by inferring time-calibrated phylogenies that alternatively included and excluded species from the ‘living fossil’ clades *Lepisosteidae* (gars), *Amia* (bowfins), and *Acipenseriformes* (sturgeons and paddlefishes), and which included these species but without fossil calibrations placed on the nodes corresponding to their crown clades. As is expected if rate accommodation induces spurious inference of the ages of crown ray-finned fishes and the oldest actinopterygian divergences, we found that excluding living fossils resulted in crown ages for *Actinopterygii* and *Teleostei* that, on average, are 20 to 70 million years younger than when living fossil clades of ray-finned fishes are included (Figure 5). We did not observe similar shifts in estimated divergence times for deeply-divergent sarcopterygian clades (e.g., lungfishes; (Figure 5), suggesting that errors related to rate accommodation are largely restricted to ray-finned fishes. This observation is further supported by our analysis of nucleotide substitution rates across time-calibrated phylogenies from both subsampled and complete datasets (Figure 4B; Figures S10-11).

**Figure 5.**
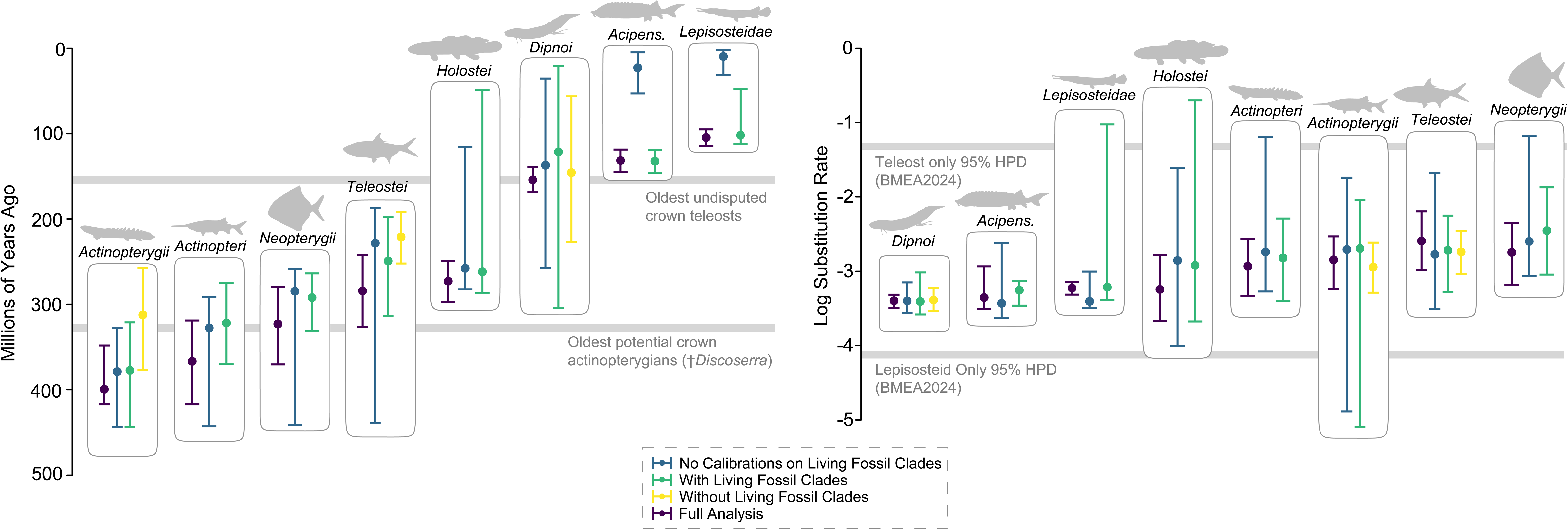
Variation in Genomic Evolution Renders Some Divergence Times Unknowable. Bowtie plot shows the median (dots) and 95% highest posterior density intervals (bars) for selected early-diverging vertebrate clades inferred across the taxon and fossil calibration inclusion experiments and the main tree. Silhouettes are public domain from phylopic.org or by CDB.

Previous research using both this exon dataset (Brownstein et al. 2024a) and other sequence data (Braasch et al. 2016; Takezaki 2018; Du et al. 2020) has demonstrated that gars and sturgeons exhibit slow molecular evolutionary rates relative to other vertebrates across their genomes. The mechanisms accounting for these slow rates of molecular evolution remain unclear, and may involve the suppression of transposable element activity (Braasch et al. 2016) or enhanced DNA repair capacity (Brownstein et al. 2024a). These slow rates represent biological characteristics rather than analytical artifacts (Braasch et al. 2016; Takezaki 2018; Du et al. 2020; Brownstein et al. 2024a), and so the inclusion of more sequence data from these and other ray-finned fishes would not necessarily remedy the problem.

When rates of substitution and divergence times are jointly estimated across ray-finned fishes, we found that the inferred rates for these typically slow-evolving clades converge more closely with the higher teleost rates, and vice versa (Figure 5). As rate variance is accommodated during inference of the time tree, the successive divergences among the four major actinopterygian clades (*Teleostei, Holostei, Acipenseriformes,* and *Polypteridae*) are pushed backward in time (Figure 5; Figures S10-S11).

Further evidence that the extreme disparity in molecular evolutionary rates among the major clades of ray-finned fishes contributes to erroneous divergence time estimation comes from analyses where holosteans and acipenseriforms are included without corresponding fossil calibrations. Crown gars are known from complete skeletons starting in the Late Cretaceous, ∼97.2 Ma, and many members of crown *Lepisosteidae* have been described from complete body fossils from the latest Mesozoic and early Cenozoic (Grande 2010; Brito et al. 2017; Brownstein 2022; Brownstein and Lyson 2022; Brownstein et al. 2023b). Similarly, crown group acipenseriforms and acipenserids are known from partial and complete skeletons from throughout the Cretaceous and Cenozoic (Grande and Bemis 1991; Hilton et al. 2011; Sato et al. 2018; Murray et al. 2020; Hilton et al. 2023; Murray et al. 2023; Brownstein 2023a; Hilton and Grande 2023). Yet, when fossil calibrations are not used for these clades, the median ages of crown gars and acipenseriforms shift far towards the Recent, between 9 and 23 million years ago (Figure 5). These age estimates directly contradict the fossil records of these clades and show that even relaxed clock models fail to accommodate the exceptionally slow rates of these living fossil lineages. The estimated ages of *Actinopterygii*, *Teleostei*, and the other major ray-finned fish divergences found in analyses in which living fossil clades are included without corresponding fossil calibrations are also similar to those estimated when the living fossil crown clades are calibrated using fossils, suggesting that the disparity between the exceptionally fast teleost rate and slow holostean and acipenseriform molecular evolutionary rates drives the overestimation of ray-finned fish divergences as the relaxed clock model attempts to accommodate these rate differences by pushing the estimated teleost crown age and successive ray-finned fish divergences farther back in time (Figure 5).

## Discussion

Here, we use a time-calibrated phylogeny of jawed vertebrates to investigate how a phylogeny of a species-rich lineage can be distorted by macroevolutionary events and intrinsic aspects of organismal molecular biology. Our analyses tie rapid diversifications of both terrestrial and marine gnathostome lineages to macroevolutionary events, such as the Cretaceous Terrestrial Revolution (Lloyd et al. 2008; Meredith et al. 2011; Weaver et al. 2024) and the Cretaceous-Paleogene Mass Extinction (Hackett et al. 2008; Jarvis et al. 2014; Prum et al. 2015; Feng et al. 2017; Alfaro et al. 2018; Burbrink et al. 2020; Kuhl et al. 2021; Ghezelayagh et al. 2022; Foley et al. 2023; Berv et al. 2024; Stiller et al. 2024). Phylogenetic signal within genes and sequence sites plunges at the K-Pg boundary (Figure 3), supporting the hypothesis that extinctions leave a lasting mark on evolutionary history by subjecting lineages to both major bottlenecks and sudden ecological opportunity as ecosystems reassemble (Gavrilets and Losos 2009; Prum et al. 2015; Stroud and Losos 2016; Reddy et al. 2017; Berv et al. 2024). The process of mass extinction and subsequent rapid radiation has sculpted the diversity of living jawed vertebrates across their phylogenetic and ecological diversity and resulted in some of the most famous evolutionary radiations, including birds, placental mammals, snakes, and pelagic predatory ray-finned fishes such as tunas, mackerels, jacks, and swordfishes (Figure 1). The intense diversification following the Cretaceous-Paleogene extinction manifests in our data as a distinct reduction in successive branching times among jawed vertebrates, centered at approximately 66 million years ago (Figure 3). Additionally, we observe anomaly zones – regions characterized by substantial conflict between gene trees and species trees – throughout the backbone phylogenies of these post-Cretaceous radiations (Figure 2C; Figure 3). Paradoxically, as much as genomes have provided consilience around jawed vertebrate relationships, these data also reveal the confounding effects of macroevolutionary events on inferring phylogenies.

Just as macroevolutionary events confound the resolution of jawed vertebrate phylogeny, the organismal biology of some jawed vertebrate clades hinders our ability to reconstruct their deep-time evolutionary history. Through experimental taxon and calibration exclusion experiments, we show that the inclusion of living representatives of *Acipenseriformes* and *Holostei*, which are two of the three major non-teleost lineages of ray-finned fishes and possess the slowest nucleotide substitution rates of all vertebrates (Braasch et al. 2016; Takezaki 2018; Du et al. 2020; Brownstein et al. 2024a), pushes the ages of *Teleostei* and *Actinopterygii* back in time by 29 to 65 million years (Figure 4). This temporal displacement helps explain a notable discrepancy in the fossil record. Although time-calibrated phylogenies built using molecular data suggest a Devonian-Carboniferous origin for the ray-finned fish crown group (Near et al. 2012; Betancur-R et al. 2013, 2017; Giles et al. 2017; Hughes et al. 2018), the earliest fossil evidence of major actinopterygian crown clades (including crown actinopterygians, crown neopterygians, and pan-acipenseriforms) appears 30 to 50 million years later (Argyriou et al. 2018, n.d.). This inconsistency suggests that molecular data alone cannot reliably determine the ages of *Actinopterygii* and its major internal divergences (*Actinopteri*, *Neopterygii*, and *Teleostei*). This result reemphasizes the importance of the fossil record for estimating the ages of this clades by providing minimum bounds.

The increasing availability of high-quality genomic data presents both opportunities and challenges in reconstructing the Tree of Life. Our phylogenetic analyses of 540 representative jawed vertebrates using genome-scale data corroborates several recently proposed relationships among ancient tetrapod, sarcopterygian, and ray-finned fishes (Figure 1)(Chakrabarty et al. 2017; Hughes et al. 2018; Thompson et al. 2021; Ghezelayagh et al. 2022; Melo et al. 2022; Foley et al. 2023; Parey et al. 2023), yet provide just as little clarity about others. Although comparisons of different types of genomic marker datasets show that deep-branching relationships across the vertebrate tree of life largely remain stable regardless of marker type (e.g., UCEs, exons, AHEs) used (Reddy et al. 2017; Alda et al. 2021; Arcila et al. 2021), analyses incorporating thousands loci still fail to unambiguously resolve the relationships of lineages in rapid radiations such as neoavian birds (Stiller et al. 2024) and placental mammals (Foley et al. 2023). Our phylogenetic analyses similarly fail to resolve neoavian, acanthomorph and laurasiatherian mammal relationships, which are largely unresolved in our multispecies coalescent analyses owing to intense gene tree discordance (Figure 2, Figure S2, Figure S3). To this end, the phylogeny that we infer cannot be considered a robust hypothesis of the relationships of these lineages and other rapidly diversifying clades.

However, our results are notable for facilitating direct comparisons of levels of support for particular relationships across unrelated vertebrate lineages. For example, our analyses demonstrate that the monophyly of recently proposed ray-finned fish lineages *Oseanacephala* and *Stomiati* receive as much support from the same genome-wide dataset as the monophyly of placental and marsupial mammals, squamates, and archosaurs (Figure 2). Conversely, our results are the first direct comparison of uncertainty surrounding the relationships of unrelated order-level clades of jawed vertebrates classically considered major post-Cretaceous radiations; although gene tree uncertainty for neoavian ordinal relationships far surpasses that in acanthomorph and otophysan fishes, placental mammals, and anguimorph lizards, for example, bootstrap support and site concordance factors are comparable across these radiations (Figure S7). Although we only use one marker set comprised of exons, which are expected to be under strong selection, the ages of rapid radiations that are poorly resolved in our phylogeny, especially birds, placental mammals, acanthomorph fishes, are closely comparable to those estimated in previous studies (Meredith et al. 2011; dos Reis et al. 2012, 2014; Jarvis et al. 2014; Prum et al. 2015; Hughes et al. 2018; Kuhl et al. 2021; Ghezelayagh et al. 2022; Álvarez-Carretero et al. 2022; Foley et al. 2023; Berv et al. 2024; Stiller et al. 2024). It is especially exciting to us that despite the obvious lack of resolution within *Neoaves* in our analyses, our time-calibrated phylogeny still places the rapid diversification of this lineage right at the K-Pg boundary (Chen et al. 2025), which has been found using genomic datasets containing different marker types, including exons (Jarvis et al. 2014), anchored hybrid enrichment loci (Prum et al. 2015; Berv et al. 2024), transcriptomes (Kuhl et al. 2021), and combinations of different marker sets (Stiller et al. 2024). Similarly, the middle Cretaceous age we infer for most divergences among placental mammal orders matches those found in phylogenomic analyses using different data types (Meredith et al. 2011; dos Reis et al. 2012, 2014; Álvarez-Carretero et al. 2022; Foley et al. 2023). Thus, our inference of the timing of these radiations cannot be attributed to data type, and the severity of phylogenetic discordance is indeed comparable across multiple distantly related jawed vertebrate lineages that diversified in response to the K-Pg mass extinction.

Our estimates of the ages of major jawed vertebrate clades are also affected by lineage-specific aspects of the genome, and most obviously by rates of sequence evolution (Figure 5). A striking example appears in ray-finned fishes, where substitution rate variation among the four earliest diverging lineages likely results in age overestimates of up to 70 million years for both the actinopterygian and teleost crown groups (Figure 5). These results challenge the current paradigm of phylogenetic reconstruction by demonstrating the limits of phylogenetic inference against the realities of confounding macroevolutionary events and organismal biology. Indeed, it is our inability to achieve complete phylogenetic resolution that illuminates macroevolutionary dynamics and lineage-specific patterns of genome evolution across more than 500 million years of jawed vertebrate evolution.

## Acknowledgements

We thank members of the Muñoz, Near, and Ogbunu labs at Yale University for discussions related to this manuscript.

## Author Contributions

CDB and TJN conceived the project. CDB and YL collected the data. CDB analyzed the data and wrote the manuscript. CDB, YL, and TJN reviewed and edited drafts of the manuscript.

## Data Availability

All data needed to replicate the results in this study are provided in the Supplementary Information or on the associated Yale Dataverse Repository (https://doi.org/10.60600/YU/YL2LSZ). All sequences come from genomic data available on the NCBI sequence read archive or Hughes et al. (Hughes et al. 2018), and are feature in Table S1.

## I. List of Fossil Calibrations

*†Phoebodus saidselachus*

**Justification of Placement:** †*Phoebodus saidselachus* calibrates the most recent common ancestor (MRCA) of crown *Chondrichthyes* (specifiers: *Callorhinchus milii*, *Carcharodon carcharias*) in our node-dating analyses. The placement of †*P. saidselachus* within crown *Chondrichthyes* on the stem of *Elasmobranchii* (sharks, skates and rays) is supported by parsimony and Bayesian phylogenetic analyses of morphological characters, including the 221 character, 60 taxon matrix of Frey et al. (Frey et al. 2019), the 230 character, 64 taxon matrix of Frey et al. (Frey et al. 2020; Klug et al. 2023), and the 236 character, 36 taxon matrix of Brownstein et al. (Brownstein, Near, et al. 2024). All of these matrices are modified from the matrix of Coates et al. (Coates et al. 2018). For the purposes of this justification, we rely on the phylogeny presented in figure 4 in Frey et al. (Frey et al. 2019) as the reference tree.

Phylogenetically optimized apomorphies uniting †*P. saidselachus* with Pan-*Elasmobranchii* are: smooth ceratohyal has a posterior, lateral fossa (character 51:1 in Frey et al. (Frey et al. 2019)), dorsal otic ridge forms a crest posteriorly (156:1 in Frey et al. (Frey et al. 2019)), slot-shaped endolymphatic fossa divides dorsal otic ridge along midline (158:1 in Frey et al. (Frey et al. 2019)), dorsal portion of occipital arch wedged between otic capsules (171:1 in Frey et al. (Frey et al. 2019)), posterior/pelvic-level dorsal fin possesses calcified base plate (200:1 in Frey et al. (Frey et al. 2019)).

**Stratigraphic Horizon**: Madene El Mrakib and Aguelmous, Ibâouane Formation (Lahfira Member, Thylacocephalan Layer)(Frey et al. 2020), southern Maïder region, eastern Anti-Atlas, Morocco; Middle Famennian Stage of the Devonian (Frey et al. 2019), 372.2 to 359.3-358.9 Ma (Becker et al. 2012; Gradstein et al. 2021).

**Equivalent Fossil Calibrations:** †*Ferromirum oukherbouchi* and †*Maghriboselache mohamezanei* are chondrichthyans known from whole-body fossils from the same stratigraphic horizon that are unambiguously placed as stem-holocephalans in crown *Chondrichthyes* in parsimony and Bayesian phylogenetic analyses of morphological characters (Frey et al. 2020; Klug et al. 2023; Brownstein, Near, et al. 2024).

**Fossil tip age**: 358.9 Ma. The node calibration was created such that 97.5% of the probability distribution fell before 358.9 Ma.

*†Guiyu oneiros*

**Justification of Placement:** †*Guiyu oneiros* calibrates the MRCA of crown *Osteichthyes* (specifiers: *Homo sapiens*, *Danio rerio*) in our node-dating analyses. The placement of †*Guiyu oneiros* within crown *Osteichthyes* is supported by multiple parsimony and Bayesian phylogenetic analyses of morphological characters, including the 153 character, 23 taxon matrix of Zhu et al. (Zhu et al. 2009), the 236 character, 78 taxon matrix of Giles et al.(Giles, Friedman, et al. 2015), the 182 character, 49 taxon matrix of Giles et al.(Giles, Darras, et al. 2015), the 269 character, 96 taxon matrix of Lu et al. (Lu, Giles, et al. 2016), the 335 character, 103 taxon matrix of Qiao et al. (Qiao et al. 2016), the 336 character, 104 taxon matrix of Choo et al. (Choo et al. 2017), the 347 character, 104 taxon matrix of Cui et al. (Cui et al. 2019), and the 694 character, 159 taxon matrix of Cui et al. (Cui et al. 2023). Notably, Lu et al. (Lu et al. 2017), King et al. (King et al. 2017), and Brazeau et al. (Brazeau et al. 2020) placed †*G. oneiros* within a clade of Devonian osteichthyans that is the sister lineage to crown *Osteichthyes*. However, analyses of an expanded version of the King et al. (King et al. 2017), and Brazeau et al.(Brazeau et al. 2020) matrices either place this clade in a polytomy with crown osteichthyans or on the sarcopterygian stem under a Bayesian tip-dating approach (Chase Doran Brownstein 2023). For the purposes of this justification, we rely on the phylogeny presented in supplementary figure 6 in Cui et al. (Cui et al. 2023) Because no character state optimizations were provided in Cui et al. (Cui et al. 2023), we reran the matrix presented in the paper using TNT v 1.5 (Goloboff and Catalano 2016) to optimize apomorphies; we ran an initial Wagner search over 1000 replicates and default parameters for ratchet, tfuse, drift, and sectorial search followed by traditional bisection reconnection branch swapping over 100,000 trees. We summarized the topologies in a strict consensus and then recorded apomorphies of two nodes: the clade containing †*G. oneiros* and crown *Osteichthyes* and †*Guiyu oneiros* and Pan-*Sarcopterygii*. Phylogenetically optimized apomorphies uniting †*G. oneiros* with crown *Osteichthyes* are: presence of differentiated lepidotrichia (28:1 in Cui et al. (Cui et al. 2023)), posterior end of supraorbital canal contained in postparietal (178:0 in Cui et al. (Cui et al. 2023)), and posterior expansion of the maxilla (270:0 in Cui et al. (Cui et al. 2023)). Phylogenetically optimized apomorphies uniting †*G. oneiros* with Pan-*Sarcopterygii* are: dermal cranial joint at the level of sphenoid-otic junction (111:1 in Cui et al. (Cui et al. 2023)), presence of the supraorbital (125:1 in Cui et al. (Cui et al. 2023)), presence of an endoskeletal intracranial joint (371:1 in Cui et al. (Cui et al. 2023)), internasal vacuities shallow, paired pits with strong midline ridge (390:1 in Cui et al. (Cui et al. 2023)), and presence of an unconstricted cranial notochord (411:1 in Cui et al. (Cui et al. 2023)).

**Stratigraphic Horizon**: Qujing, Kuanti Formation, Yunnan Province, China; late Ludfordian Stage of the Silurian (Zhu et al. 2013). The Ludfordian Stage ranges between 425.6 and 423.0 Ma (Gradstein et al. 2021).

**Equivalent Fossil Calibrations:** None. Slightly younger calibrations of the same node include the ‘psarolepids’ †*Achoania jarvikii* (Zhu et al. 2001) and †*Psarolepis romeri* (Qu et al. 2013), as well as the stem actinopterygian †*Meemania eos* (Lu, Giles, et al. 2016). However, these taxa are younger than the age of the oldest stem-lungfishes (see below)(Cui et al. 2022) and are therefore redundant for fossil calibration.

**Fossil tip age**: 423.0 Ma. The node calibration was created such that 97.5% of the probability distribution fell before 423.0 Ma.

*†Youngolepis praecursor*

**Justification of Placement:** †*Youngolepis praecursor* calibrates the MRCA of *Dipnoi* and *Latimeria* (specifiers: *Neoceratodus forsteri*, *Latimeria chalumnae*) in our node dating analyses.

The placement of †*Y. praecursor* on the stem of *Dipnoi* is supported by parsimony and Bayesian phylogenetic analyses of morphological characters, including the 335 character, 103 taxon matrix of Qiao et al. (Qiao et al. 2016), the 242 character, 37 taxon matrix of Lu et al. (Lu, Zhu, et al. 2016), the 70 character, 25 taxon matrix of Friedman (FRIEDMAN 2007), and the 277 character, 88 taxon matrix of Cui et al. (Cui et al. 2022). For the purposes of this justification, we rely on the phylogenies presented in supplementary figure 2 in Cui et al.(Cui et al. 2022) Because no character state optimizations were provided by Cui et al. (Cui et al. 2022) for the placement of †*Y. praecursor* sister to all other members of Pan-*Dipnoi*, we reran the matrix presented in the paper using TNT v 1.5 (Goloboff and Catalano 2016) to optimize apomorphies; we ran an initial Wagner search over 1000 replicates and default parameters for ratchet, tfuse, drift, and sectorial search followed by traditional bisection reconnection branch swapping over 100,000 trees. We summarized the topologies in a strict consensus and then recorded apomorphies of †*Y. praecursor*+Pan-*Dipnoi.* Phylogenetically optimized apomorphies uniting †*Y. praecursor* with Pan-*Dipnoi* are: closed pineal opening (1:1 in Cui et al. (Cui et al. 2022)), quadratojugal as large as jugal (34:0 in Cui et al. (Cui et al. 2022)), adductor fossa more than 1/3 length of jaw (133:0 in Cui et al. (Cui et al. 2022)), addition of large dentine elements at regular intervals to lateral margin of pterygoid and/or prearticular (166:0 in Cui et al. (Cui et al. 2022)), dentine sheet on resorption areas within tooth plate on palate (174:1 in Cui et al. (Cui et al. 2022)), marginal tooth ridge continuous (185:1 in Cui et al. (Cui et al. 2022)), intracranial joint present as a ventral cranial fissure (190:1 in Cui et al. (Cui et al. 2022)), and a triangular hyomandibular (271:1 in Cui et al. (Cui et al. 2022)).

**Stratigraphic Horizon**: Yunnan, China; Xitun Formation, earliest Lochkovian Stage of the Devonian (Zhao et al. 2021; Cui et al. 2022). We followed Cui et al. (Cui et al. 2022) as treating its age as 418.0 Ma.

**Equivalent Fossil Calibrations:** †*Diabolepis speratus* is known from the same formation and is unambiguously placed as a stem-dipnoan crownward of †*Youngolepis praecursor* in the phylogeny of Cui et al. (Cui et al. 2022), as well as a pan-dipnoan in other phylogenetic analyses (Qiao et al. 2016; Kemp et al. 2017; Brazeau et al. 2020; Cloutier et al. 2020; Zhu et al. 2021).

**Fossil tip age**: 418.0 Ma. The node calibration was created such that 97.5% of the probability distribution fell before 418.0 Ma.

*†Ferganoceratodus martini*

**Justification of Placement:** †*Ferganoceratodus martini* calibrates the MRCA of *Neoceratodus forsteri* and *Protopterus annectens* in our node-dating analyses. The placement of †*F. martini* in crown *Dipnoi* is supported by parsimony-based phylogenetic analysis of morphological characters, including the 14 character, 16 taxon matrix of Cavin et al. (CAVIN et al. 2007) We use the tree supplied in figure 18 of Cavin et al. (CAVIN et al. 2007) as the reference phylogeny for this placement. Because not all character state optimizations were provided by Cavin et al. (CAVIN et al. 2007) for the placement of †*F. martini* in *Dipnoi*, we reran the matrix presented in the paper using TNT v 1.5 (Goloboff and Catalano 2016) to optimize apomorphies; we ran an initial Wagner search over 1000 replicates and default parameters for ratchet, tfuse, drift, and sectorial search followed by traditional bisection reconnection branch swapping over 100,000 trees. †*Dipterus valenciennesi* was used as the outgroup. We summarized the topologies in a strict consensus and then recorded apomorphies of crown *Dipnoi* and Pan-*Lepidosirenidae* (*Lepidosiren* + *Protopterus* total clade). Phylogenetically optimized apomorphies that †*F. martini* shares with the dipnoan crown clade are: median series with 3 bones or bone pairs (1:1 in Cavin et al. (CAVIN et al. 2007)), and mediolateral series with four or more bones (3:0 in Cavin et al. (CAVIN et al. 2007)). Phylogenetically optimized apomorphies that †*F. martini* shares with Pan-*Lepidosirenidae* are: skull roof anteriorly straight, with a short rostrum without a supraorbital sensory canal (5:2 in Cavin et al. (CAVIN et al. 2007)).

**Stratigraphic Horizon**: Phu Nam Jun, Kalasin Province, Thailand; Upper Phu Kradung Formation, Tithonian Stage of the Late Jurassic to Berriasian Stage of the Early Cretaceous (CAVIN et al. 2007; Kemp et al. 2017).

**Equivalent Fossil Calibrations:** Numerous extinct lungfishes from as old as the Permian and Triassic have been placed within the lungfish crown clade in phylogenetic analyses of morphological characters (CAVIN et al. 2007; Kemp et al. 2017). Our use of †*Ferganoceratodus martini* provides a fairly conservative young age (earliest Cretaceous) for the prior on the crown lungfish divergence.

**Fossil tip age**: 140.2 Ma. The node calibration was created such that 97.5% of the probability distribution fell before 140.2 Ma.

*†Triadobatrachus massinoti*

**Justification of Placement:** †*Triadobatrachus massinoti* calibrates the MRCA of crown *Batrachia* (specifiers: *Ambystoma mexicanum*, *Xenopus tropicalis*) in our node-dating analyses. The placement of †*T. massinoti* in crown *Amphibia* on the anuran stem is supported by multiple parsimony and Bayesian phylogenetic analyses of morphological characters, including the 360 character, 62 taxon matrix of Schoch et al. (Schoch et al. 2020) the 308 character, 53 taxon matrix of Jones et al. (Jones et al. 2022), and the 355 character, 63 taxon matrix of Kligman et al. (Kligman et al. 2023). For the purposes of this justification, we rely on the phylogeny presented in Kligman et al. (Kligman et al. 2023) Because no character state optimizations were provided by Kligman et al. (Kligman et al. 2023) for the placement of †*T. massinoti* in crown *Batrachia* sister to *Anura*, we reran the matrix presented in the paper using TNT v 1.5(Goloboff and Catalano 2016) to optimize apomorphies; we ran an initial Wagner search over 1000 replicates and default parameters for ratchet, tfuse, drift, and sectorial search followed by traditional bisection reconnection branch swapping over 100,000 trees. We summarized the topologies in a strict consensus and then recorded apomorphies of three nodes: crown *Amphibia*, crown *Batrachia*, and the clade in Pan-*Anura* uniting †*T. massinoti* with crown frogs. Phylogenetically optimized apomorphies uniting †*T. massinoti* with crown *Amphibia* are: weak torsion of the humerus (191:1 in Kligman et al. (Kligman et al. 2023)), absence of the postorbital (226:1 in Kligman et al. (Kligman et al. 2023)), tabular absent (234:1 in Kligman et al. (Kligman et al. 2023)), lateral process of pterygoid contribution to posttemporal fossa absent (253:0 in Kligman et al. (Kligman et al. 2023)), trunk intercentra absent (272:1 in Kligman et al. (Kligman et al. 2023)), ectopterygoid absent (340:1 in Kligman et al. (Kligman et al. 2023)), interclavicle absent (341:1 in Kligman et al. (Kligman et al. 2023)), cleithrum absent (342:1 in Kligman et al. (Kligman et al. 2023)), and vomer irregularly shaped, with a posteromedially expanded choana (343:1 in Kligman et al. (Kligman et al. 2023)). Phylogenetically optimized apomorphies uniting †*T. massinoti* with crown *Batrachia* are: nasal as wide as long (15:1 in Kligman et al. (Kligman et al. 2023)), orbit and naris separated only by a narrow gap of bone (24:1 in Kligman et al.(Kligman et al. 2023)), vertebral transverse processes distally extended with laterally directed diapophyses (156:1* in Kligman et al. (Kligman et al. 2023); this character is coded as states 0 and 1 in †*T. massinoti*), ilium shaft slender and elongated (194:1 in Kligman et al. (Kligman et al. 2023)), internal trochanter of femur reduced to a short crest (201:1 in Kligman et al. (Kligman et al. 2023)), ossified hyoids present (270:0 in Kligman et al. (Kligman et al. 2023)), skull table T-shaped (307:1 in Kligman et al. (Kligman et al. 2023)), footplate of stapes much smaller than the fenestra vestibule (332:1 in Kligman et al. (Kligman et al. 2023)), and the presence of the batrachian operculum (350:1in Kligman et al. (Kligman et al. 2023)). Phylogenetically optimized apomorphies uniting †*T. massinoti* with Pan-*Anura* are: internarial distance wider than interorbital distance (14:1 in Kligman et al. (Kligman et al. 2023)), otic notch appears as a semicircular embayment between squamosal and posterior skull table (50:0 in Kligman et al. (Kligman et al. 2023)), main axis of ilium shaft inclined anteriorly (197:2 Kligman et al. (Kligman et al. 2023)), parahyoid present (271:1in Kligman et al. (Kligman et al. 2023)), single notochordal atlas centrum (283:1 in Kligman et al. (Kligman et al. 2023)), elongated tibiale and fibulare present (302:1 in Kligman et al. (Kligman et al. 2023)), and absence of a canal for palatal ramus of the facial nerve in antotic region (331:0 in Kligman et al. (Kligman et al. 2023)). **Stratigraphic Horizon**: There is considerable uncertainty surrounding the provenance of the only known fossil of †*Triadobatrachus massinoti,* which was recovered from the Sakamena Group of Madagascar (Ascarrunz et al. 2016). Many analyses have used an uppermost Early Triassic age for this taxon (Ascarrunz et al. 2016; Jones et al. 2022), primarily based on observations of vertebrate biostratigraphy at the type locality (Ascarrunz et al. 2016). A younger age (Olenekian, 249.9-246.7 Ma) was suggested on the basis of invertebrate faunas from other localities corresponding to the Triassic Sakamena Group in Madagascar (Ascarrunz et al. 2016). The youngest possible age of this fossil would be Middle Triassic, which is the younger bound on the Sakamena Group itself (Wescott and Diggens 1998). As such, we set the age of this fossil to this minimum bound, 237.0 Ma (Gradstein et al. 2021). In addition to representing a conservative interpretation of the age of this fossil, a Middle Triassic age of †*T. massinoti* allows for recognition of several equivalent fossil calibrations (see below), which provides a robust minimum age for the frog-salamander divergence.

**Equivalent Fossil Calibrations:** †*Triassurus sixtelae*, a member of Pan-*Caudata* known from partial articulated skeletons from the Ladinian-Carnian Madygen Formation of Kyrgyzstan (Schoch et al. 2020; Jones et al. 2022; Kligman et al. 2023), and †*Czatkobatrachus polonicus*, a possible member of Pan-*Anura* known from hypodigms of isolated bones from the fissure fills of the Early-Middle Triassic of Poland (Evans and Borsuk-Bialynicka 1998), are equivalent fossil calibrations. We did not use †*Czatkobatrachus polonicus* because of conceptual issues with the use of isolated bone hypodigms as fossil calibrations (Caldwell et al. 2015; Chase D. Brownstein et al. 2023).

**Fossil tip age**: 237.0 Ma. The node calibration was created such that 97.5% of the probability distribution fell before 237.0 Ma.

*†Rhadinosteus parvus*

**Justification of Placement:** †*Rhadinosteus parvus* calibrates the MRCA of crown *Pipanura* (specifiers: *Rana catesbeiana*, *Xenopus tropicalis*) in our node-dating analyses. The placement of †*R. parvus* along either the stem or within crown *Pipoidea* is supported by multiple parsimony phylogenetic analyses of morphological characters, including the 165 character, 36 taxon matrix of Gómez (Gómez 2016), the 18 character, 13 taxon matrix of Henrici (Henrici 1998), and the 72 character, 24 taxon matrix of Báez (Báez 2013) For the purposes of this justification, we rely on the phylogeny presented in Gómez (Gómez 2016). Because no character state optimizations were provided by Gómez (Gómez 2016) for the placement of †*R. parvus* in crown *Anura* sister to *Pipoidea*, we reran the matrix presented in the paper using TNT v 1.5 (Goloboff and Catalano 2016) to optimize apomorphies; we ran an initial Wagner search over 1000 replicates and default parameters for ratchet, tfuse, drift, and sectorial search followed by traditional bisection reconnection branch swapping over 100,000 trees. We summarized the topologies in a strict consensus and then recorded apomorphies of two nodes: crown *Pipanura* and the clade in Pan-*Pipoidea* including †*R. parvus.* Phylogenetically optimized apomorphies uniting †*R. parvus* with crown *Pipanura* are: scapula, ratio between maximum width of glenoid fossa and maximum width of shaft between 1.0 and 0.5 (109:1 in Gómez (Gómez 2016)), tibiale-fibulare less than 20% of the length of the hindlimb. Phylogenetically optimized apomorphies uniting †*R. parvus* with Pan-*Pipoidea* are: complete fusion of the frontoparietals (4:1 in Gómez (Gómez 2016)), absence of the subotic alae of the parasphenoid (17:1 in Gómez (Gómez 2016)), and anterior third of the cultriform process of the parasphenoid very narrow, such that the process is strongly tapering and pointed (19:2 in Gómez (Gómez 2016)).

**Stratigraphic Horizon**: Dinosaur National Monument (DNM) locality 96, Utah; Brushy Basin Member, Morrison Formation (Henrici 1998). The Brushy Basin Member of the Morrison Formation at DNM is considered to be form the Kimmeridgian to Tithonian Stages of the Late Jurassic (Henrici 1998); ^40^Ar/^39^Ar dating places this unit between 151 and 145 Ma (Kowallis et al. 1991; Trujillo and Kowallis 2015). The dinosaur quarry at DNM has been dated to between 150.91 and 150.04 Ma using magnetostratigraphy (Maidment et al. 2017). Following previous work on Morrison Formation herpetofauna (Brownstein et al. 2022; Meyer et al. 2023), we use an age of 145.0 Ma.

**Equivalent Fossil Calibrations: None.**

**Fossil tip age**: 145.0 Ma. The node calibration was created such that 97.5% of the probability distribution fell before 145.0 Ma.

*†Henkelotherium guimarotae*

**Justification of Placement:** †*Henkelotherium guimarotae* calibrates the MRCA of crown *Mammalia* (specifiers: *Homo sapiens*, *Ornithorhynchus anatinus*) in our node-dating analyses. The placement of †*H. guimarotae* as a crown mammal on the stem of *Theria* (Pan-*Marsupialia* + Pan-*Placentalia*) is supported by multiple parsimony and Bayesian phylogenetic analyses of morphological characters, including the 4541 character, 126 taxon matrix of O’Leary et al. (O’Leary et al. 2013) (including as updated by Velazco et al. (Velazco et al. 2022)), the 491 character, 112 taxon matrix of Luo et al. (Luo, Meng, et al. 2015), the Luo et al., 497 character, 114 taxon matrix of Luo et al. (Luo, Gatesy, et al. 2015), the 505 character, 117 taxon matrix of Han et al. (Han et al. 2017), the 538 character, 125 taxon matrix of Huttenlocker et al. (Huttenlocker et al. 2018), the 530 character, 84 taxon matrix of Krause et al. (Krause et al. 2020) (including as modified by Hoffman et al. (Hoffmann et al. 2020)), the 615 character, 135 taxon matrix of Wang and Wang (Wang and Wang 2023). Notably, the position of †*H. guimarotae* within crown *Mammalia* is unchanged regardless of the placement of problematic †*Haramiyida* and †*Gondwanatheria* among near-mammal mammaliaforms (Luo, Gatesy, et al. 2015; Han et al. 2017; Huttenlocker et al. 2018; Hoffmann et al. 2020; Krause et al. 2020; Wang and Wang 2023). For the purposes of this justification, we rely on the phylogenies presented in figure 1 of Hoffman et al. (Hoffmann et al. 2020). Because no character state optimizations were provided by Hoffman et al. (Hoffmann et al. 2020) for the placement of †*H. guimarotae* in crown *Mammalia*, we reran the matrix presented in the paper using TNT v 1.5 (Goloboff and Catalano 2016) to optimize apomorphies; we ran an initial Wagner search over 1000 replicates and default parameters for ratchet, tfuse, drift, and sectorial search followed by traditional bisection reconnection branch swapping over 100,000 trees. We summarized the topologies in a strict consensus and then recorded apomorphies of two nodes: crown *Mammalia* and *Theria*+†*Dryolestidae* (incl. †*H. guimarotae*). Phylogenetically optimized apomorphies uniting †*H. guimarotae* with crown *Mammalia* are: lumbar ribs of dorsal vertebrae synostosed to vertebrae to form transverse processes (11:1 in Hoffman et al. (Hoffmann et al. 2020)), internal sutures of ilium, ischium, and pubis fused in adults (58:1 in Hoffman et al. (Hoffmann et al. 2020)), patellar groove of femur present (74:1 in Hoffman et al. (Hoffmann et al. 2020)), distinct medial malleolus of tibia (77:1 in Hoffman et al. (Hoffmann et al. 2020)), calcaneo-astragal facet of calcaneum placed medial to calcaneo-fibular facet (101:1 in Hoffman et al. (Hoffmann et al. 2020)), os calcaris present (111:1 in Hoffman et al. (Hoffmann et al. 2020)), dentary condyle of mandible positioned above the level of the postcanine alveoli (346:1 in Hoffman et al. (Hoffmann et al. 2020)), lower ultimate premolar (p5) hypoconid present (403:1 in Hoffman et al. (Hoffmann et al. 2020)), ml-2, protoconid far more bulging than paraconid and metaconid (432:1 in Hoffman et al. (Hoffmann et al. 2020)), ml-2, metacristid (protocristid) crest between protoconid (cusp a) and metaconid (cusp c) transversely oriented relative to long axis of lower molars (433:2 in Hoffman et al. (Hoffmann et al. 2020)), m1, main cusps of trigonid aligned at an acute angle (435:1 in Hoffman et al. (Hoffmann et al. 2020)), ml-2, primary functional cusps not labiolingually compressed (436:0 in Hoffman et al. (Hoffmann et al. 2020)), m1-2, lingual root present (506:1 in Hoffman et al. (Hoffmann et al. 2020)), and the presence of prevallum/postvallid shearing (522:1 in Hoffman et al.(Hoffmann et al. 2020)). Phylogenetically optimized apomorphies uniting †*H. guimarotae* and †*Dryolestidae* with *Theria* are: fusion of the neural hemiarches of the atlas (2:1 in Hoffman et al. (Hoffmann et al. 2020)), styloid process of radius present (45:1 in Hoffman et al. (Hoffmann et al. 2020)), styloid process of ulna present (50:1 in Hoffman et al. (Hoffmann et al. 2020)), femur neck distinct and long (64:1 in Hoffman et al. (Hoffmann et al. 2020)), lesser trochanter of femur positioned on ventromedial or ventral side of shaft (68:1 in Hoffman et al. (Hoffmann et al. 2020)), postpalatine torus of palate present (158:1 in Hoffman et al. (Hoffmann et al. 2020)), quadrate ramus of alisphenoid absent (208:2 in Hoffman et al. (Hoffmann et al. 2020)), absence of a diastema distal to lower canine (356:0 in Hoffman et al. (Hoffmann et al. 2020)), four lower incisors in each quadrant (357:1 in Hoffman et al. (Hoffmann et al. 2020)), four lower premolars in each quadrant (380:1 in Hoffman et al. (Hoffmann et al. 2020)), lower ultimate premolar (p5) paraconid well-developed (401:1 in Hoffman et al. (Hoffmann et al. 2020)), interlocking distal postcanines/ml-2 present (425:1 in Hoffman et al. (Hoffmann et al. 2020)).

**Stratigraphic Horizon**: Guimarota coal mine, Portugal (Krebs 1991; Jäger et al. 2020; Luo and Martin 2023); Alcobaça Formation, Lower Kimmeridgian Stage of the Late Jurassic, 154.8-149.2 Ma (Gradstein et al. 2021).

**Equivalent Fossil Calibrations:** Highly dependent on phylogeny. Equivalent-age or older dryolestids and allotherians are known but occasionally placed outside the mammalian crown group. Refer to the cited phylogenetic analyses above. We chose †*Henkelotherium guimarotae* as the fossil calibration for crown mammals because it is reliably placed within the crown across multiple phylogenetic analyses and is known from a well-characterized cranium and postcranium, contrary to other mammals from the Jurassic with similar morphology (Krebs 1991; Jäger et al. 2020). However, if allotherians, and in particular haramiyidans, are crown group mammals, the age of crown *Mammalia* could be as old as the Triassic (see, for example, Huttenlocker et al. (Huttenlocker et al. 2018)).

**Fossil tip age**: 154.8 Ma. The node calibration was created such that 97.5% of the probability distribution fell before 154.8 Ma.

*†Deltatheridium pretrituberculare*

**Justification of Placement:** †*Deltatheridium pretrituberculare* calibrates the MRCA of crown *Marsupialia* (specifiers: *Monodelphis domestica*, *Macropus eugenii*) in our node-dating analyses. The placement of †*D. pretrituberculare* as a member of crown *Marsupialia* is supported by multiple parsimony and Bayesian phylogenetic analyses of morphological characters in a recent analysis incorporating new morphological data (Velazco et al. 2022). For the purposes of this justification, we rely on the phylogeny presented in figure 1 of Velazco et al. (Velazco et al. 2022). Phylogenetically optimized apomorphies uniting †*D. pretrituberculare* and crown *Theria* are: mandibular foramen positioned at or posterior to the midpoint of the ramus (1228:1 in Velazco et al. (Velazco et al. 2022)), pterygoid crest absent (1230:1 in Velazco et al. (Velazco et al. 2022)), Meckelian groove absent (1236:1 in Velazco et al. (Velazco et al. 2022)), Li2 spatulate in shape (1308:1 in Velazco et al. (Velazco et al. 2022)), Li3 spatulate in shape (1325:1 in Velazco et al. (Velazco et al. 2022)), upper canine Uc1 larger than lower canine Lc1 (1409:0 in Velazco et al. (Velazco et al. 2022)), talonid present on Lp2 (1492:1 in Velazco et al.(Velazco et al. 2022)), Lp3 absent (1499:1 in Velazco et al. (Velazco et al. 2022)), protoconid height on Lp4 subequal to height of p5 protoconid (1532:1 in Velazco et al. (Velazco et al. 2022)), degree of hypsodonty of UP1 VAR more that 1.0 (hypsodont, 1718:1 in Velazco et al. (Velazco et al. 2022)), UP2 crown with two or more clearly defined cusps (1735:1 in Velazco et al. (Velazco et al. 2022)), P4 height less than that of P5 but greater than half the height of P5 (1792:3 in Velazco et al. (Velazco et al. 2022)), UP4 crown with two or more clearly defined cusps (1794:1 in Velazco et al. (Velazco et al. 2022)), one premolar trenchant in penultimate premolar position (2024:1 in Velazco et al. (Velazco et al. 2022)), precingulid present on Lm1 (2042:1 in Velazco et al. (Velazco et al. 2022)), paraconid size on Lm1 moderate (approx. 75 percent the height of the protoconid; 2058:1 in Velazco et al. (Velazco et al. 2022)), protocristid transversely oriented relative to the longitudinal axis of Lm1 (2076:1 in Velazco et al. (Velazco et al. 2022)), cristid obliqua (=ectolophid) development on Lm1 very strong (2100:1 in Velazco et al. (Velazco et al. 2022)), precingulid present on Lm2 (2166:1 in Velazco et al. (Velazco et al. 2022)), entoconid in straight alignment with the paraconid and metaconid on Lm2 (2188:1 in Velazco et al. (Velazco et al. 2022)), protocristid transversely oriented relative to the longitudinal axis of Lm2 (2196:1 in Velazco et al. (Velazco et al. 2022)), hypoconulid buccally position to midline relative to the entoconid on Lm2 (2251:1 in Velazco et al. (Velazco et al. 2022)), 3 UM1 roots (2307:3 in Velazco et al. (Velazco et al. 2022)), M1 stylocone subequal to paracone (2344:1 in Velazco et al. (Velazco et al. 2022)), M1 preprotocrista on UM1 extends to the base of the paracone (2411:1 in Velazco et al. (Velazco et al. 2022)), area of UM3 relative to UM1 VAR very small (2582:0 in Velazco et al. (Velazco et al. 2022)), diastema present between UP1 and UP2 (2700:1 in Velazco et al. (Velazco et al. 2022)), supinator well-developed, extends proximal to the proximal opening of the entepicondylar foramen but does not approach the midshaft region (3033:1 in Velazco et al. (Velazco et al. 2022)), and unfused pubes (3312:0 in Velazco et al. (Velazco et al. 2022)).

**Stratigraphic Horizon**: Ukhaa Tolgod, Gobi Desert, Mongolia; Djadokhta Formation, precise age within the Campanian is controversial (Dingus et al. 2008), but we follow Velazco et al. (Velazco et al. 2022) in recognizing this age as early Campanian, 83.6 Ma (Gradstein et al. 2021).

**Equivalent Fossil Calibrations:** None. See Velazco et al. (Velazco et al. 2022). For the sensitivity analysis, †*Djarthia murgonensis*, a member of the total clade of *Australidelphia*, was used; this taxon is known from fossils dating to 54.6 Ma (Beck et al. 2008).

**Fossil tip age**: 83.6 Ma. The node calibration was created such that 97.5% of the probability distribution fell before 83.6 Ma.

*†Riostegotherium yanei*

**Justification of Placement:** †*Riostegotherium yanei* calibrates the MRCA of crown *Xenarthra* (specifiers: *Bradypus variegatus*, *Dasypus novemcinctus*) in our node-dating analyses. The placement of †*R. yanei* as a member of crown *Xenarthra* is supported by multiple morphological characters, including the presence of subrectangular osteoderms with characteristic microstructure (Anon 1998; Bergqvist et al. 2019). A phylogeny including this taxon is not yet available, but †*R. yanei* has been used to calibrate the xenarthran crown in multiple analyses (see Foley et al. (Foley et al. 2023)).

**Stratigraphic Horizon**: Fissure-fill deposits corresponding to the Itaboraian SALMA of the Rio de Janeiro State, Brazil (Bergqvist et al. 2019); we follow Foley et al. (Foley et al. 2023) in using a terminal Ypresian age.

**Equivalent Fossil Calibrations: None.**

**Fossil tip age**: 47.8 Ma. The node calibration was created such that 97.5% of the probability distribution fell before 47.8 Ma.

*†Eritherium azzouzorum*

**Justification of Placement:** †*Eritherium azzouzorum* calibrates the MRCA of crown *Afrotheria* (specifiers: *Loxodonta africana*, *Orycteropus afer*) in our node-dating analyses.This is a somewhat conservative placement since multiple phylogenetic analyses of morphological characters resolve †*Eritherium azzouzorum* as a stem-probiscidean, including the 143 character, 20 taxon matrix of Gheerbrant (Gheerbrant 2009) and the 109 character, 44 taxon matrix of Avilla and Mothé (Avilla and Mothé 2021). Other analyses of the latter matrix resolve this taxon on the stem of *Sirenia* (Kramarz and MacPhee 2022). Finally, the 209 character, 33 taxon matrix of Gheerbrant et al. (Gheerbrant et al. 2018) resolves †*E. azzouzorum* on the stem of *Sirenia*+*Probiscidea*. Because of this uncertainty in the placement of †*E. azzouzorum* among *Paenungulata* (*Probiscidea*, *Hyracoidea,* and *Sirenia*), which itself evades resolution using genomic data (Seiffert 2007; Bowman et al. 2023; Foley et al. 2023; Liu et al. 2024), we used this taxon to calibrate the MRCA at the base of *Afrotheria*. For the purposes of this justification, we rely on the phylogeny presented in figure 4 of Gheerbrant et al. (Gheerbrant et al. 2018).

Because no character state optimizations were provided by Gheerbrant et al. (Gheerbrant et al. 2018) for the placement of †*E. azzouzorum* in crown *Afrotheria* on the stem of *Probiscidea+Sirenia* (=*Tethytheria*), we reran the matrix presented in the paper using TNT v 1.5 (Goloboff and Catalano 2016) to optimize apomorphies; we ran an initial Wagner search over 1000 replicates and default parameters for ratchet, tfuse, drift, and sectorial search followed by traditional bisection reconnection branch swapping over 100,000 trees. We summarized the topologies in a strict consensus and then recorded apomorphies of two nodes: crown *Afrotheria* and Pan-*Tethytheria.* Phylogenetically optimized apomorphies uniting †*E. azzouzorum* and crown *Afrotheria* are: postmetacrista predominantly longitudinal (110:1 in Gheerbrant et al. (Gheerbrant et al. 2018)) and preparacrista oriented longitudinally (111:1 in Gheerbrant et al. (Gheerbrant et al. 2018)). Phylogenetically optimized apomorphies uniting †*E. azzouzorum* and *Tethytheria* are: preparacrista absent or weak (112:1 in Gheerbrant et al. (Gheerbrant et al. 2018)),and external auditory meatus elevated high above tooth row (181:1 in Gheerbrant et al. (Gheerbrant et al. 2018)).

**Stratigraphic Horizon**: Sidi Chennane quarries, Ouled Abdoun phosphate basin, Morocco; early Thanetian (Gheerbrant 2009). we follow Foley et al. (Foley et al. 2023) in using an age of 59.2 Ma.

**Equivalent Fossil Calibrations:** Highly dependent on phylogeny. Equivalent-age or older fossils potentially assignable to crown *Afrotheria* are known, but their placement is unclear. See cited phylogenetic studies in “Justification of Placement” section above.

**Fossil tip age**: 59.2 Ma. The node calibration was created such that 97.5% of the probability distribution fell before 59.2 Ma.

*†Acritoparamys atavus*

**Justification of Placement:** †*Acritoparamys atavus* calibrates the MRCA of crown *Rodentia* (specifiers: *Mus musculus*, *Hydrochoerus hydrochaeris*) in our node-dating analyses. The placement of †*A. atavus* as a member of crown *Rodentia* in the stem-sciuromorph clade †*Ischyromyidae* is supported by parsimony phylogenetic analyses of morphological characters, including the 106 character, 91 taxon matrix of Marivaux et al (Marivaux et al. 2004) and the 343 character, 45 taxon matrix of Vianey-Laud and Marivaux (Vianey-Liaud and Marivaux 2021). For this purposes of this justification, we rely on the phylogeny presented in figure 1 of Marivaux et al. (Marivaux et al. 2004), which unites this taxon with other early-diverging ‘ischyromiform’ rodents primarily on the basis of protrogomorphy and the presence of pauciserial enamel.

**Stratigraphic Horizon**: Bear Creek, Montana, USA; Fort Union Formation, Clarkfordian, 56.0 Ma (Korth 1994; Dawson and Beard 1996; Dawson et al. 2008; Casanovas-Vilar et al.).

**Equivalent Fossil Calibrations:** †*Paramys adamus* is an equivalent fossil calibration of similar placement in rodent phylogeny (within “†*Ischyromyidae*”) (Marivaux et al. 2004). †*Eliwourus topernawiensis* is a crown rodent from 29.5 Ma in Kenya, and is used as the calibration for crown *Rodentia* in the sensitivity analysis (Seiffert et al. 2025).

**Fossil tip age**: 56.0 Ma. The node calibration was created such that 97.5% of the probability distribution fell before 56.0 Ma.

*†Purgatorius janisae*

**Justification of Placement:** †*Purgatorius janisae* calibrates the MRCA of *Primates* and *Dermoptera* (specifiers: *Galeopterus variegatus*, *Homo sapiens*) in our node-dating analyses. The placement of †*Purgatorius* as a member of Pan-*Primates* is supported by parsimony phylogenetic analyses of morphological characters, including the 415 character, 85 taxon matrix of Chester et al. (Chester et al. 2015) and the 240 character, 30 taxon matrix of Chester et al. (Chester et al. 2017) For the purposes of this justification, we rely on the phylogeny presented in Figure S2 of Chester et al. (Chester et al. 2017) Phylogenetically optimized apomorphies uniting †*Purgatorius* spp. and *Primatomorpha* (*Dermoptera* + *Primates*) are: plantar pit on cuboid facet of calcaneum present (51:1 in Chester et al. (Chester et al. 2017)), m1 metastyle absent (163:0 in Chester et al. (Chester et al. 2017)), and m1, both conules present (165:0 in Chester et al. (Chester et al. 2017)). Phylogenetically optimized apomorphies uniting †*Purgatorius* spp. and Pan-*Primates* are: m1, precingulum present and not connected to postcingulum (156:0 in Chester et al. (Chester et al. 2017)), m1, entoconid notch absent (219:0 in Chester et al. (Chester et al. 2017)), m3 greater in length than m2 (226:1 in Chester et al. (Chester et al. 2017)), and lower molars that get progressively larger from m1 to m3 (229:1 in Chester et al. (Chester et al. 2017)). See Mantilla et al. (Wilson Mantilla et al. 2021) for comparisons of dental morphology across the earliest known †*Purgatorius* species.

**Stratigraphic Horizon**: ‘Harley’s Point’ UCMP locality V77087, Montana, USA; Tullock Member of the Fort Union Formation, early Puercan, lowermost Paleocene (Wilson Mantilla et al. 2021). We follow Foley et al. in using an age of 65.5 Ma (Foley et al. 2023).

**Equivalent Fossil Calibrations:** Several other species of †*Purgatorius.* See Mantilla et al. (Wilson Mantilla et al. 2021). †*Archicebus achilles* from the lower Eocene of China (54.8 Ma), is a probable crown primate that was used to calibrate the node of crown primates (this same node excluding *Galeopterus*) in the sensitivity analysis (Ni et al. 2013).

**Fossil tip age**: 65.5 Ma. The node calibration was created such that 97.5% of the probability distribution fell before 65.5 Ma.

*†Hesperocyon gregarious*

**Justification of Placement:** †*Hesperocyon gregarious* calibrates the MRCA of *Carnivora* (specifiers: *Panthera onca*, *Canis lupus*) in our node-dating analyses. The placement of †*Hesperocyon gregarious* as a member of *Carnivora* on the stem of *Canidae* is supported by parsimony phylogenetic analyses of morphological characters, including the 99 character, 48 taxon matrix of Wesley-Hunt and Flynn (Wesley-Hunt and Flynn 2005), the 98 character, 36 taxon matrix of Wesley-Hunt and Werdelin (Wesley-Hunt and Werdelin 2005), the 98 character, 50 taxon matrix of Tomiya (Tomiya 2011), the 123 character, 121 taxon matrix of Slater (Slater 2015), and the 108 character, 28 taxon matrix of Tomiya and Tseng (Tomiya and Tseng 2016). Because of instability in the placement of different lineages in *Carnivora* in these analyses, we chose to conservatively use †*H. gregarious* to calibrate the crown of *Carnivora* in our analyses. For the purposes of this justification, we rely on the phylogeny presented in figure 1 in Tomiya and Tseng (Tomiya and Tseng 2016). Phylogenetically optimized apomorphies uniting †*H. gregarious* to *Canoidea* in crown *Carnivora* are: infraorbital foramen above P4 anterior edge (4:1 in Tomiya and Tseng (Tomiya and Tseng 2016)), basioccipital lateral flange large (34:2 in Tomiya and Tseng (Tomiya and Tseng 2016)), M3 absent (53:1 in Tomiya and Tseng (Tomiya and Tseng 2016)), and M1 postprotocrista posteriorly directed (102:1 in Tomiya and Tseng (Tomiya and Tseng 2016)).

**Stratigraphic Horizon**: northeastern Colorado, USA; White River Formation. Also known from the Cypress Hills Formation, Saskatchewan, Canada, the Renova Formation, western Montana, USA; Chadron and Brule Formations, Colorado, Nebraska, North Dakota, South Dakota, and Wyoming, USA (Wang and Tedford 1996). Duchesnian to Whitneyan Land Mammal Stages (Wang and Tedford 1996); following Foley et al. (Foley et al. 2023) we use the midpoint of 37.71 Ma.

**Equivalent Fossil Calibrations:** Several in the genera †*Daphoenus* and †*Hesperocyon*. See Wesley-Hunt and Flynn (Wesley-Hunt and Flynn 2005), Benton and Donoghue (Benton and Donoghue 2007), and Slater (Slater 2015).

**Fossil tip age**: 37.71 Ma. The node calibration was created such that 97.5% of the probability distribution fell before 37.71 Ma.

†*Icaronycteris spp*.

**Justification of Placement:** †*Icaronycteris* spp. calibrates the MRCA of *Chiroptera* (specifiers: *Pteronotus parnellii*, *Megaderma lyra*) in our node-dating analyses. The placement of †*Icaronycteris spp.* as a member of *Chiroptera* in a polytomy with crown clades is supported by parsimony phylogenetic analyses of morphological characters, including the 568 character, 14 taxon matrix of Reitbergen et al. (Rietbergen et al. 2023), and the 699 character, 82 taxon matrix of Jones et al. (Jones et al. 2024). Several analyses have also recovered †*Icaronycteris* spp. on the stem of *Chiroptera*, but close to the crown clade compared to other Eocene bats. These phylogenies include the 50% majority rule tree in Jones et al. (Jones et al. 2024), phylogenies generated from the 172 character, 30 taxon matrix of Simmons and Geisler (Simmons and Geisler 1998), and the 2665 character, 30 taxon matrix of Hand et al. (Hand et al. 2023).

However, because several similarly aged Eocene bats of potential crown group affinity are known, and given the unstable relationships of extinct Eocene bats to the crown (Simmons and Geisler 1998; Hand et al. 2023; Rietbergen et al. 2023; Jones et al. 2024), we use †*Icaronycteris* spp. to calibrate the crown. For the purposes of this justification, we rely on the phylogeny presented in figure 2 in Jones et al. (Jones et al. 2024). Phylogenetically optimized apomorphies uniting †*Icaronycteris spp.* to the most inclusive clade containing these Green River bats and crown bat clades (node 19 in Jones et al. (Jones et al. 2024)) are (NB: some of these are variable in different species of †*Icaronycteris*): upper canine with flattened posterolabial surface (38:1 in Jones et al. (Jones et al. 2024)), m1-2 with lingual cingulum (75:1 in Jones et al. (Jones et al. 2024)), longitudinally oriented roots of p3 (177:0 in Jones et al. ^180,181^(Jones et al. 2024)), p3 slightly shorter than p4 (187:3 in Jones et al. (Jones et al. 2024)), p3 between 50% to 100% the size of p4 (188:2 in in Jones et al. (Jones et al. 2024)), p4 with one metaconid (203:1 in in Jones et al. (Jones et al. 2024)), and m3 hypoconulid small and low (280:1 in in Jones et al. (Jones et al. 2024)).

**Stratigraphic Horizon**: The oldest species of †*Icaronycteris* is from the American Fossil Quarry Wyoming, USA; Fossil Butte Member of the Green River Formation. This locality matches the age of Fossil Lake, which is minimally dated to 51.98 Ma (Smith et al. 2008; Rietbergen et al. 2023).

**Equivalent Fossil Calibrations:** Various, including the perhaps slightly (47.8 Ma) younger (though still Eocene) †*Tachypteron franzeni*, †*Witwatia sigei*, and †*Dizzya exsultans*, which have previously been used to constrain the age of *Chiroptera* (Foley et al. 2023) but are also unstable in phylogenetic position and are represented by fossils that are less character-rich than those of many species of †*Icaronycteris* (Jones et al. 2024). These other fossils were used in place of †*Icaronycteris* in the sensitivity analysis.

**Fossil tip age**: 51.98 Ma. The node calibration was created such that 97.5% of the probability distribution fell before 51.98 Ma.

*†Enaliarctos tedfordi*

**Justification of Placement:** †*Enaliarctos tedfordi* calibrates the MRCA of *Arctoidea* (specifiers: *Ailurus stylani*, *Ailuropoda melanoleuca*) in our node-dating analyses. The placement of †*E. tedfordi* in crown *Arctoidea* on the stem of *Pinnipedia* is supported by parsimony and Bayesian phylogenetic analyses of morphological characters, including the 52 character, 9 taxon matrix of Berta (Berta 1991) and the 204 character, 65 taxon matrix of Paterson et al. (Paterson et al. 2020). For the purposes of this justification, we rely on the phylogeny presented in figure 3 in Paterson et al. (Paterson et al. 2020). Phylogenetically optimized apomorphies uniting †*E. tedfordi* with Pan-*Pinnipedia* (=*Pinnipedomorpha* + *Puijila* clade in Paterson et al. (Paterson et al. 2020)) are: postorbital constriction present, long (3:2 in Paterson et al. (Paterson et al. 2020)), prominent pseudosylvian sulcus on braincase (36:1 in 1 in Paterson et al. (Paterson et al. 2020)), fenestra cochleae larger than oval window (70:1 in Paterson et al. (Paterson et al. 2020)), and m1 reduced relative to p4 (95:1 in Paterson et al. (Paterson et al. 2020)).

**Stratigraphic Horizon**: Beaver Creek, Lincoln County, Oregon; Yaquina Formation, Oligocene, 28.1 Ma (Prothero et al. 2001; Poust and Boessenecker 2018; Paterson et al. 2020; Foley et al. 2023).

**Equivalent Fossil Calibrations:** Potentially the pan-mustelids †*Mustelavus priscus*, †*Pseudobassaris riggsi*, the arctoid †*Amphicticeps shackelfordi*, and the pan-arctids †*Allocyon loganensis* and †*Phoberogale* sp. (Paterson et al. 2020).

**Fossil tip age**: 28.1 Ma. The node calibration was created such that 97.5% of the probability distribution fell before 28.1 Ma.

*†Ravenictis krausei*

**Justification of Placement:** †*Ravenictis krausei* calibrates the MRCA of *Ferae* (specifiers: *Ailurus stylani*, *Manis pentadactyla*) in our node-dating analyses. The identification of †*Ravenictis krausei* as a viverrid pan-carnivoran is based on the following morphological features: crown asymmetry, and the development of the protocone cingula (Fox and Youzwyshyn 1994). Wear patterns and the relative development of the cusps were also cited as support for a pan-carnivoran identity (Fox and Youzwyshyn 1994). We follow previous analyses (Foley et al. 2023) in using †*Ravenictis krausei* as a calibration for the split between *Carnivora* and its sister lineage *Pholidota*.

**Stratigraphic Horizon**: Saskatchewan, Canada; Rav W-1 Horizon, Ravenscrag Formation, Paleocene (Clarkfordian), 64.0 Ma (Fox and Youzwyshyn 1994; Foley et al. 2023).

**Equivalent Fossil Calibrations:** Various, including several slightly younger (62.0-56.0 Ma) stem-carnivorans in †*Viverridae* and †*Miacidae* (Fox and Youzwyshyn 1994; Wesley-Hunt and Flynn 2005; Tomiya 2011; Tomiya and Tseng 2016). This slightly younger age (56.0 Ma) was used for the sensitivity analysis.

**Fossil tip age**: 64.0 Ma. The node calibration was created such that 97.5% of the probability distribution fell before 64.0 Ma.

*†Mystacodon selensis*

**Justification of Placement:** †*Mystacodon selensis* calibrates the MRCA of *Cetacea* (specifiers: *Orcinus orca*, *Balaenoptera acutorostrata*) in our node-dating analyses. The placement of †*M. selensis* in crown *Cetacea* on the stem of *Mysticeti* is supported by parsimony and Bayesian phylogenetic analyses of morphological characters, including the 272 character, 38 taxon matrix of Lambert et al. (Lambert et al. 2017), the 391 character, 138 taxon dataset of Boessenecker et al. (Boessenecker et al. 2023), and the 282 character, 35 taxon matrix of Muizon et al. (Anon 2019). For the purposes of this justification, we rely on the phylogeny presented in figure 52 of Muizon et al.(Anon 2019). Phylogenetically optimized apomorphies uniting †*M. selensis* with Pan-*Mysticeti* in crown *Cetacea* are: maxilla roughly as high as wide or lower relative to width at transverse midpoint of section anteroposteriorly located at the level of posterior third of the bone length but anterior to antorbital process (5:1 in Muizon et al. (Anon 2019)), anteriormost portion of jugal broadly underlapped by maxilla (54:1 in Muizon et al. (Anon 2019)), presence of the maxillary infraorbital plate (57:1 in Muizon et al. (Anon 2019)), zygomatic process apex of squamosal closely apposed or situated ventral to postorbital process (102:1 in Muizon et al. (Anon 2019)), supraoccipital triangular in dorsal view (115:1 in Muizon et al. (Anon 2019)), external occipital crest restricted to anterior half of supraoccipital shield (117:1 in Muizon et al. (Anon 2019)), height of dentary does not taper ventrally at anterior extremity (226:1 in Muizon et al. (Anon 2019)), and humeral head oriented posteriorly to posteroproximally (265:1 in Muizon et al. (Anon 2019)).

**Stratigraphic Horizon**: Playa Media Luna locality, Pisco Basin, Peru; Yumaque Member of the Paracas Formation, Eocene (Priabonian), 36.4 Ma (Lambert et al. 2017; Anon 2019).

**Equivalent Fossil Calibrations:** None. The closest fossil calibration is the slightly younger †*Llanocetus denticrenatus*, which is another pan-mysticete that is closer to the crown clade than †*Mystacodon selensis* (Lambert et al. 2017; Fordyce and Marx 2018; Fordyce and Marx 2018; Anon 2019; Boessenecker et al. 2023).

**Fossil tip age**: 36.4 Ma. The node calibration was created such that 97.5% of the probability distribution fell before 36.4 Ma.

*†Megachirella watchleri*

**Justification of Placement:** †*Megachirella watchleri* calibrates the MRCA of crown *Lepidosauria* (specifiers: *Sphenodon punctatus*, *Python molurus*) in our node-dating analyses.

The placement of †*M. watchleri* within crown *Lepidosauria* is supported by parsimony and Bayesian phylogenetic analyses of morphological characters, including the 347 character, 129 taxon dataset of Simões et al. (Simões et al. 2018) (including its modifications in Bittencourt et al. (Bittencourt et al. 2020), Brownstein et al. (Chase D. Brownstein et al. 2023), Martinez et al. (Martínez et al. 2021), and Sobral et al. (Sobral et al. 2020)), the 348 character, 125 taxon matrix of Simões et al. (Simões et al. 2022), the 381 character, 116 taxon matrix of Ford et al. (Ford et al. 2021), the 377 character, 115 taxon matrix of Griffiths et al. (Griffiths et al. 2021), and the 131 character, 29 taxon matrix of Simões et al. (Simões et al. 2020) (including as modified by Freisem et al. (Freisem et al. 2024) The modified version of the matrix in Griffiths et al. (Griffiths et al. 2021)/ Ford et al. (Ford et al. 2021) presented in Tałanda et al. (Tałanda et al. 2022) recovers †*M. watchleri* within a clade of Early Triassic to Middle Jurassic pan-lepidosaurs that forms the sister to the crown group. However, the varied morphology of members of this clade and the incomplete nature of some of these specimens (see Griffiths et al. (Griffiths et al. 2021) and Ford et al. (Ford et al. 2021)) means that its monophyly should be viewed with caution, especially since different methodologies applying molecular constraints induce the collapse of this clade into a polytomy of lepidosaur total group lineages (see extended data figure 8b in Tałanda et al. (Tałanda et al. 2022)). In any case, equivalent calibrations exist (see below). For the purposes of this justification, we rely on the phylogeny presented in figure 2 of Simões et al. (Simões et al. 2018). Phylogenetically optimized apomorphies uniting †*M. wetmorei* with Pan-*Lepidosauria* are: posterior dentary teeth delimited by a labial wall only (212:0 in Simões et al. (Simões et al. 2018)), posterior maxillary teeth delimited by a labial wall only (213:0 in Simões et al. (Simões et al. 2018)), and presacral pleurocentra, midventral crest present (232:1 in Simões et al. (Simões et al. 2018)). †*M. wetmorei* is united with *Lepidosauria* based on a single phylogenetically optimized apomorphy: frontals fused (67:1 in Simões et al. (Simões et al. 2018)). Phylogenetically optimized apomorphies uniting †*M. wetmorei* with Pan-*Squamata* are: absence of squamosal anteroventral process (50:0 in Simões et al. (Simões et al. 2018)), lateral process of ectopterygoid absent (114:0 in Simões et al. (Simões et al. 2018)), alar crest of prootic present (142:1 in Simões et al. (Simões et al. 2018)), clavicles secondarily curved anteroposteriorly (287:1 in Simões et al. (Simões et al. 2018)), developed radial condyle on humerus present (310:0 in Simões et al. (Simões et al. 2018)), distal carpal 1 fused to metacarpal (324:1 in Simões et al. (Simões et al. 2018)).

**Stratigraphic Horizon**: Monte Prà della Vacca, Braies Dolomites, Bolzano, Italy (Renesto and Bernardi 2014); Dont Formation, Braies Group, Anisian Stage of the Early Triassic, approximately 242.0 Ma (Renesto and Bernardi 2014; Simões et al. 2018).

**Equivalent Fossil Calibrations:** †*Wirtembergia hauboldae* from the Ladinian (242.0 to 237.0 Ma)(Gradstein et al. 2021) Erfut Formation of Germany is the oldest known rhychocephalian and is only slightly younger than †*Megachirella watchleri* from the Anisian of Italy. †*Fraxinisaura rozynekae* is a lepidosaur that is also known from the Erfut Formation of Germany, but whose affinities to the lepidosaur crown clade remain unresolved (Schoch and Sues 2018; Ford et al. 2021; Griffiths et al. 2021; Chase D. Brownstein et al. 2023).

**Fossil tip age**: 242.0 Ma. The node calibration was created such that 97.5% of the probability distribution fell before 242.0 Ma.

*†Eoscincus ornatus*

**Justification of Placement:** †*Eoscincus ornatus* calibrates the MRCA of crown *Unidentata* (specifiers: *Hemicordylus capensis*, *Python molurus*) in our node-dating analyses. The placement of †*E. ornatus* within crown *Unidentata* is supported by parsimony and Bayesian phylogenetic analyses of morphological characters, including the 622 character, 119 taxon matrix of Brownstein et al. (Brownstein et al. 2022) and the 637 character, 154 taxon matrix of Meyer et al. (Meyer et al. 2023). For the purposes of this justification, we use the phylogeny presented in Figure S10 of Brownstein et al. (Brownstein et al. 2022). Phylogenetically optimized apomorphies uniting †*E. ornatus* with Pan-*Scincoidea* in crown *Squamata* are: surangular adductor fossa of the external face of mandible is deep and extends ventrally more than halfway down (399:1 in Brownstein et al. (Brownstein et al. 2022)). Absence of the lateral exposure of the jugal below the orbit due to obscuration by the maxilla jugal process (149:0 in Brownstein et al. (Brownstein et al. 2022)) is also found to be a character uniting †*E. ornatus* with Pan-*Scincoidea* in the analysis constrained based on molecular phylogenies. Phylogenetically optimized apomorphies uniting †*E. ornatus* with †*Paramacellodidae* in Pan-*Scincoidea* are: jugal broadly separated from prefrontal (144:0 in Brownstein et al. (Brownstein et al. 2022)), jugal medial ridge weakly developed such that the jugal is positioned lateral to the ectopterygoid at base in dorsal view (157:0 in Brownstein et al. (Brownstein et al. 2022)), 10 to 20 dentary teeth (421:2 in Brownstein et al. (Brownstein et al. 2022)), posterior teeth unicuspid (434:0 in in Brownstein et al. (Brownstein et al. 2022)), and dermal skull roof bones ornamented with slight rugosities about the frontoparietal suture (572:1 in Brownstein et al. (Brownstein et al. 2022)). **Stratigraphic Horizon**: DINO 317 Quarry, Dinosaur National Monument, Utah, USA; Brushy Basin Member of the Morrison Formation, Tithonian Stage of the Late Jurassic, approximately 145.0 Ma.(Brownstein et al. 2022)

**Equivalent Fossil Calibrations:** †*Microteras borealis*, known from the Tithonian Morrison Formation of Wyoming, is an equivalent fossil calibration (Brownstein et al. 2022).

**Fossil tip age**: 145.0 Ma. The node calibration was created such that 97.5% of the probability distribution fell before 145.0 Ma.

*†Chthonophis subterraneus*

**Justification of Placement:** †*Chthonophis subterraneus* calibrates the MRCA of *Lacertoidea* (specifiers: *Lacerta agilis*; *Salvator merianae*) in our node-dating analyses. The placement of †*C. subterraneus* within *Lacertoidea* as a member of Pan-*Amphisbaenia* is supported by parsimony phylogenetic analyses of morphological characters, including the 308 character, 51 taxon matrix of Longrich et al. (Longrich et al. 2015). For the purposes of this justification, we use the phylogeny presented in Figure S1 of Longrich et al. (Longrich et al. 2015). Because no character state optimizations were provided by Longrich et al. (Longrich et al. 2015) for the placement of †*C. subterraneus* within *Lacertoidea* as a member of Pan-*Amphisbaenia*, we reran the matrix presented in the paper using TNT v 1.5(Goloboff and Catalano 2016) to optimize apomorphies; we ran an initial Wagner search over 1000 replicates and default parameters for ratchet, tfuse, drift, and sectorial search followed by traditional bisection reconnection branch swapping over 100,000 trees. We summarized the topologies in a strict consensus and then recorded apomorphies of Pan-*Amphisbaenia*. Phylogenetically optimized apomorphies uniting †*C. subterraneus* with Pan-*Amphisbaenia* are: dentary symphysis with extensive development of a symphyseal facet extending caudally below the Meckelian groove (165:1 in Longrich et al. (Longrich et al. 2015)), Fusion of marginal teeth: unfused to each other (0); fused to each other (1). Marginal tooth crowns separated by large gaps (220: 1 in Longrich et al. (Longrich et al. 2015)), and eight dentary teeth present (231:2 in Longrich et al. (Longrich et al. 2015)).

**Stratigraphic Horizon**: Bug Creek Anthills, Fort Union Formation, Montana, USA; Maastrichtian Stage of Late Cretaceous to earliest Danian Stage of Paleocene (Longrich et al. 2015), 66.02 Ma (Gradstein et al. 2021).

**Equivalent Fossil Calibrations: None.**

**Fossil tip age**: 66.02 Ma. The node calibration was created such that 97.5% of the probability distribution fell before 66.02 Ma.

*†Priscagama gobiensis*

**Justification of Placement:** †*Priscagama gobiensis* calibrates the MRCA of *Iguania* (specifiers: *Pogona vitticeps*, *Anolis carolinensis*) in our node-dating analyses. The placement of †*P. gobiensis* within *Iguania* as a member of Pan-*Acrodonta* is supported by parsimony and Bayesian phylogenetic analyses of morphological characters, including the 610 character, 192 taxon matrix of Gauthier et al. (Gauthier et al. 2012), the 347 character, 129 taxon dataset of Simões et al.,(Simões et al. 2018) the 622 character, 119 taxon matrix of Brownstein et al. (Brownstein et al. 2022), and the 155 character, 133 taxon matrix of Scarpetta (Scarpetta 2024). For the purposes of this justification, we use the phylogeny presented in Figure S10 of Brownstein et al. (Brownstein et al. 2022). Because no character state optimizations were provided by Brownstein et al. (Brownstein et al. 2022) for the placement of †*P. gobiensis* within *Iguania* as a member of Pan-*Acrodonta*, we reran the matrix presented in the paper using TNT v 1.5 (Goloboff and Catalano 2016) to optimize apomorphies; we ran an initial Wagner search over 1000 replicates and default parameters for ratchet, tfuse, drift, and sectorial search followed by traditional bisection reconnection branch swapping over 100,000 trees. We summarized the topologies in a strict consensus and then recorded apomorphies of *Iguania* and Pan-*Acrodonta*. Phylogenetically optimized apomorphies uniting †*P. gobiensis* with *Iguania* are: nasals in contact dorsal to premaxillary nasal process (25:1 in Brownstein et al. (Brownstein et al. 2022)), postorbital contacts parietal ventrolaterally at frontoparietal suture (71:2 in Brownstein et al. (Brownstein et al. 2022)), and dentary coronoid process posteriorly terminates just behind level of coronoid apex (364:1 in Brownstein et al. (Brownstein et al. 2022)). Phylogenetically optimized apomorphies uniting †*P. gobiensis* with Pan-*Acrodonta* are: frontal interorbital width 28-34% of frontoparietal suture width (48:3 in Brownstein et al. (Brownstein et al. 2022)) and maxillary tooth row extends to posterior third of orbit (415:1 in Brownstein et al. (Brownstein et al. 2022)).

**Stratigraphic Horizon**: Campanian-Maastrichtian of the Gobi Desert of Mongolia and China;(Borsuk-Bialynicka and Moody 1984; Keqin and Lianhai 1995; Gao and Norell 2000) following Brownstein et al. (Brownstein et al. 2022) we use an age of 71.0 Ma.

**Equivalent Fossil Calibrations:** At least 10 other species of iguanian have been reported from the same Campanian units of the Gobi Desert. Among these, †*Ctenomastax parva*, †*Mimeosaurus crassus*, †*Phrynosomimus asper*, †*Isodontosaurus gracilis*, †*Zapsosaurus sceliphros*, and †*Polrussia mongoliensis* have been placed within the crown group on the stems of *Pleurodonta* and *Acrodonta* in multiple phylogenetic analyses (Gauthier et al. 2012; Longrich et al. 2012; Simões et al. 2018; Brownstein et al. 2022; Scarpetta 2024).

**Fossil tip age**: 71.0 Ma. The node calibration was created such that 97.5% of the probability distribution fell before 71.0 Ma.

*†Arcanosaurus ibericus*

**Justification of Placement:** †*Arcanosaurus ibericus* calibrates the MRCA of crown *Anguimorpha* (specifiers: *Heloderma charlesbogerti*, *Varanus komodoensis*) in our node-dating analyses. The identification of †*A. ibericus* as a member of Pan-*Varanidae* in *Anguimorpha* is based on the following morphological features: cervical vertebrae with a hypapophysis originating from the centrum with individualized intercentral elements, obliquity of the axis of the cotyle–condyle system, articulated caudal chevron intercentra present (Houssaye et al. 2013). **Stratigraphic Horizon**: Viajete site, Salas de los Infantes, Cameros Basin, Iberian Peninsula, Burgos, Spain; Castrillo de la Reina Formation, Barremian to Aptian Stages of the Early Cretaceous.(Houssaye et al. 2013) We use an age of 121.4 Ma, which represents the boundary for these two stages.(Gradstein et al. 2021)

**Equivalent Fossil Calibrations:** †*Dorsetisaurus* spp. is a crown squamate known from fossils as old as the Tithonian Stage of the Late Jurassic (∼145 Ma) that has variously been placed in Pan-*Anguimorpha* or within the crown group (Evans et al. 2006; Conrad 2008).

**Fossil tip age**: 121.4 Ma. The node calibration was created such that 97.5% of the probability

*†Messelopython freyi*

**Justification of Placement:** †*Messelopython freyi* calibrates the MRCA of crown *Pythonomorpha* (specifiers: *Python molurus*, *Naja naja*) in our node-dating analyses. The placement of †*M. freyi* within *Pythonomorpha* as a member of Pan-*Pythonidae* (and indeed, the pythonid crown group) is supported by parsimony and Bayesian phylogenetic analyses of morphological characters, including the 785 character, 90 taxon matrix of Zaher and Smith,(Zaher and Smith 2020) the 264 character, 52 taxon matrix of Onary et al. (Onary et al. 2021), the 240 character, 50 taxon matrix of Palci et al. (Palci et al. 2024), and the 240 character, 51 taxon matrix of Croghan et al. (Croghan et al. 2024). For the purposes of this justification, we use the phylogeny presented in figure 2a of Zaher and Smith (Zaher and Smith 2020). Because no character state optimizations were provided by Zaher and Smith (Zaher and Smith 2020) for the placement of †*M. freyi* within *Pythonomorpha* as a member of Pan-*Pythonidae*, we reran the matrix presented in the paper using TNT v 1.5 (Goloboff and Catalano 2016) to optimize apomorphies; we ran an initial Wagner search over 1000 replicates and default parameters for ratchet, tfuse, drift, and sectorial search followed by traditional bisection reconnection branch swapping over 100,000 trees. We summarized the topologies in a strict consensus and then recorded apomorphies of Pan-*Pythonidae*. Phylogenetically optimized apomorphies uniting †*Messelopython freyi* with Pan-*Pythonidae* include: quadrate lateral head off crest borders ventral fossa (231:1 in Zaher and Smith (Zaher and Smith 2020)), palatine ventromedial extension from maxillary process present but does not extend ventromedially (296:0 in Zaher and Smith (Zaher and Smith 2020)), palatine choanal process forms a short vertical or horizontal lamina that does not reach the vomer (312:2 in Zaher and Smith (Zaher and Smith 2020)), and posterior border of the palatine process of the maxilla inconspicuous or rounded, not projecting posteromedially (771:0 in Zaher and Smith (Zaher and Smith 2020)).

**Stratigraphic Horizon**: Messel Pit, Germany; Middle Messel Formation, which is at minimum the upper bound of the Geiseltalian Mammal Stage at 46.5 Ma (Mertz and Renne 2005).

**Equivalent Fossil Calibrations:** Several, including the booids †*Messelophis variatus*, †*Rieppelophis ermannorum,* †*Eoconstrictor fischeri*, †*E. spinifer*, †*E. barnesi*, and †*Rageryx schmidi* (Zaher and Smith 2020; Croghan et al. 2024; Palci et al. 2024).

**Fossil tip age**: 46.5 Ma. The node calibration was created such that 97.5% of the probability

*†Notoemys oxfordiensis*

**Justification of Placement:** †*Notoemys oxfordiensis* calibrates the MRCA of crown *Testudines* (specifiers: *Podocnemis expansa*, *Chelydra serpentina*) in our node-dating analyses. The placement of †*N. oxfordiensis* within *Testudines* in Pan-*Pleurodira* is supported by parsimony phylogenetic analyses of morphological characters, including the 245 character, 100 taxon matrix of Conde et al. (López-Conde et al. 2017). For the purposes of this justification, we rely on the phylogeny presented in Figure S6 of Conde et al. (López-Conde et al. 2017).

Phylogenetically optimized characters uniting †*N. oxfordiensis* with Pan-*Pleurodira* are: distinct anal notch of xiphiplastron present (164:1 in Conde et al. (López-Conde et al. 2017)), gular, only one scute (168:1 in Conde et al. (López-Conde et al. 2017)), and ischium attached to plastron by a broad suture (225:1 in Conde et al. (López-Conde et al. 2017)).

**Stratigraphic Horizon**: Viñales, Cuba; Jagua Vieja Member of the Jagua Formation, Oxfordian (de la Fuente and Iturralde-Vinent 2001), 161.5 to 154.8 Ma (Gradstein et al. 2021).

**Equivalent Fossil Calibrations:** None. Several other species of †*Notoemys* do occur in slightly younger geological units (López-Conde et al. 2017).

**Fossil tip age**: 154.8 Ma. The node calibration was created such that 97.5% of the probability distribution fell before 154.8 Ma.

*†Kappachelys okurai*

**Justification of Placement:** †*Kappachelys okurai* calibrates the MRCA of *Trionychia* (specifiers: *Apalone spinifera*, *Carettochelys insculpta*) in our node-dating analyses. The placement of †*K. okurai* within *Trionychia* on the stem (but not crown) of *Trionychidae* is supported by morphological characters, including: 20 or fewer peripheral bones, partial loss of keratinous scutes in the carapace, and loss of the costal-peripheral suture (Hirayama et al. 2013; 2017).

**Stratigraphic Horizon**: Oarashidani locality, Shiramine area, Hakusan City, Ishikawa Prefecture, Honshu, Japan; lower member of the Akaiwa Formation, Tetori Group (2017). We use 131.7 Ma, which is the upper bound for the age of the perhaps contemporaneous lower member of the Kuwajima Formation (2017).

**Equivalent Fossil Calibrations:** Unclear; some fossils from China and Japan may belong to trionychids of similar age (2017).

**Fossil tip age**: 131.7 Ma. The node calibration was created such that 97.5% of the probability distribution fell before 131.7 Ma.

*†“Trachemys” antiqua*

**Justification of Placement:** †“*Trachemys*” *antiqua* calibrates the MRCA of *Emydidae* (specifiers: *Cuora amboinensis*, *Mauremys reevesii*) in our node-dating analyses. The placement of †“*T.*” *antiqua* in *Emydidae* is supported by phylogenetic analysis of morphological characters, including the 170 character, 125 taxon matrix of Vlachos (Vlachos 2018). For the purposes of this justification, we rely on the phylogeny presented in figure 1 of Vlachos (Vlachos 2018).

Phylogenetically optimized apomorphies uniting †“*T.*” *antiqua* with *Emydidae* are: anterior musk duct foramina absent (134:1 in Vlachos (Vlachos 2018)), posterior musk duct foramina absent (135:1 in Vlachos (Vlachos 2018)).

**Stratigraphic Horizon**: Various from North Dakota, South Dakota, Colorado, and Nebraska, USA; Chadron and White River formations, Priabonian Stage of the Eocene to Brule Formation, Rupelian Stage of the Oligocene (Vlachos 2018). We follow Joyce et al. (Joyce et al. 2013) in using an age of 32.0 Ma for this taxon.

**Equivalent Fossil Calibrations: None.**

**Fossil tip age**: 32.0 Ma. The node calibration was created such that 97.5% of the probability distribution fell before 32.0 Ma.

*†Protorosaurus spleneri*

**Justification of Placement:** †*Protorosaurus spleneri* calibrates the MRCA of *Archelosauria* (specifiers: *Alligator mississippiensis*, *Chelydra serpentina*) in our node-dating analyses. The placement of †*P. spleneri* within *Archelosauria* as a member of Pan-*Archosauria* is supported by parsimony and Bayesian phylogenetic analyses of morphological characters, including the 144 character, 17 taxon matrix of Gottmann-Quesada and Sander,(Gottmann-Quesada and Sander 2009) the 219 character, 40 taxon matrix of Ezcurra et al. (Ezcurra et al. 2014), the 822 character, 157 taxon matrix of Ezcurra et al. (Ezcurra et al. 2020), and the 307 character, 42 taxon matrix of Spiekman et al. (Spiekman et al. 2021). For the purposes of this justification, we rely on the phylogeny presented in figure 33 of Spiekman et al. (Spiekman et al. 2021). Phylogenetically optimized apomorphies supporting placement of †*P. spleneri* within *Archelosauria* in Pan-*Archosauria* are: maxilla alveolar margin concave in lateral view (25:1 in Spiekman et al. (Spiekman et al. 2021)), medially expanded supratemporal fossa resulting in mediolaterally narrow parietal table (83:1 in Spiekman et al. (Spiekman et al. 2021)), anterior portion of surangular-angular suture appears anteroposteriorly concave ventrally in lateral view (158:0 in Spiekman et al. (Spiekman et al. 2021)), retroarticular process upturned (163:1 in Spiekman et al. (Spiekman et al. 2021)), diapophyses and parapophyses of anterior to mid-postaxial cervical vertebrae on different processes nearly touch (188:2 in Spiekman et al. (Spiekman et al. 2021)), anterior margin of neural spine of anterior to mid-postaxial cervical vertebrae anterodorsally inclined at more than 60° from horizontal plane (194:1 in Spiekman et al. (Spiekman et al. 2021)), long transverse processes of mid-dorsal vertebrae (207:1 in Spiekman et al. (Spiekman et al. 2021)), shallow fossa lateral to base of neural spine of each dorsal vertebra (209:1 in Spiekman et al. (Spiekman et al. 2021)), coracoid moderately expanded posteriorly in lateral view (232:1 in Spiekman et al. (Spiekman et al. 2021)), proximal portion of interclavicle subtriangular to diamond-shaped (237:0 in Spiekman et al. (Spiekman et al. 2021)), weak (<35°) torsion between proximal and distal ends of humerus (241:1 in Spiekman et al. (Spiekman et al. 2021)), and entepicondylar foramen of humerus absent (246:1 in Spiekman et al. (Spiekman et al. 2021)).

**Stratigraphic Horizon**: Middridge and Quarrington Quarries, Durham, England, and various localities in central Germany (Gottmann-Quesada and Sander 2009; Spiekman et al. 2021); Marl Slate and Kupferschiefer Formation, Wuchiapingian Stage of Permian. We use an age of 256.0 Ma (Simões et al. 2018).

**Equivalent Fossil Calibrations:** None, although several ‘protorosaurian’-grade pan-archosaurs of unclear phylogenetic placement are known from the latest Permian, as are the earliest archosauriforms (Ezcurra et al. 2014; Ezcurra et al. 2014; Bernardi et al. 2015; Spiekman et al. 2021).

**Fossil tip age**: 256.0 Ma. The node calibration was created such that 97.5% of the probability distribution fell before 256.0 Ma.

*†Teleocrater rhadinus*

**Justification of Placement:** †*Teleocrater rhadinus* calibrates the MRCA of *Archosauria* (specifiers: *Alligator mississippiensis*, *Columbia livia*) in our node-dating analyses. The placement of †*T. rhadinus* within *Archosauria* and in Pan-*Aves* is supported by parsimony and Bayesian analyses of morphological characters, including the 606 character, 86 taxon matrix of Nesbitt et al. (Nesbitt et al. 2017), the 822 character, 157 taxon matrix of Ezcurra et al. (Ezcurra et al. 2020), and the 823 character, 159 taxon matrix of Nesbitt et al. (Nesbitt et al. 2023). For the purposes of this justification, we rely on the phylogeny presented in figure 21 of Nesbitt et al. (Nesbitt et al. 2023). Because no character state optimizations were provided by Nesbitt et al. (Nesbitt et al. 2023) for the placement of †*T. rhadinus* within *Archosauria* as a member of Pan-*Aves*, we reran the matrix presented in the paper using TNT v 1.5 (Goloboff and Catalano 2016) to optimize apomorphies; we ran an initial Wagner search over 1000 replicates and default parameters for ratchet, tfuse, drift, and sectorial search followed by traditional bisection reconnection branch swapping over 100,000 trees. We summarized the topologies in a strict consensus and then recorded apomorphies of *Archosauria*. Phylogenetically optimized apomorphies supporting placement of †*T. rhadinus* within *Archosauria* are: antorbital fossa on the lateral surface of the maxilla present on the horizontal process (= posterior process) of the maxilla ventral to the antorbital fenestra, but not reaching the posteroventral corner of the maxillary contribution to the border of the opening (54:2 in Nesbitt et al. (Nesbitt et al. 2023)), fibula transverse width at mid-length distinctly narrower than transverse width of the tibia (529:1 in Nesbitt et al. (Nesbitt et al. 2023)), fibula distal end in lateral view rounded or flat (symmetrical; 532:1 in Nesbitt et al. (Nesbitt et al. 2023)), ventral notch between main body of calcaneum and the calcaneal tuber present (552:1 in Nesbitt et al. (Nesbitt et al. 2023)), and articular facets for the fibula and astragalus on calcaneum separated (555:1 in Nesbitt et al. (Nesbitt et al. 2023)).

**Stratigraphic Horizon**: Ruhuhu Basin, Tanzania; Lifua Member, Manda Beds;(Nesbitt et al. 2017) Anisian, 247.2 to 242.0 Ma (Gradstein et al. 2021).

**Equivalent Fossil Calibrations:** The aphanosaurs †*Dongusuchus efremovi* and †*Yarasuchus deccanensis* are equivalent calibrations (Nesbitt et al. 2017; Nesbitt et al. 2023).

**Fossil tip age**: 242.0 Ma. The node calibration was created such that 97.5% of the probability distribution fell before 242.0 Ma.

*†Deinosuchus rugosus*

**Justification of Placement:** †*Deinosuchus rugosus* calibrates the most recent common ancestor of *Crocodylia* (specifiers: *Alligator mississippiensis*, *Crocodylus porosus*) in our node-dating analyses. The placement of †*D. rugosus* as a member of crown *Crocodylia* on the stem *Caimaninae* is supported by parsimony analyses of morphological characters, including the 163 character, 83 taxon matrix of Cossette and Brochu (Cossette and Brochu 2020). The use of †*D. rugosus* as a calibration point for the crocodylian crown clade is conservative; we employed a conservative approach due to the unstable placement of several putatively older crown crocodylian species in phylogenetic analyses of morphological characters (Brochu 1997; Bona et al. 2018; Lee and Yates 2018; Mateus et al. 2019; Melstrom and Irmis 2019; Rio and Mannion 2021; Darlim et al. 2022). For the purposes of this justification, we rely on the phylogeny presented in figure 28 of Cossette and Brochu.(Cossette and Brochu 2020). Phylogenetically optimized apomorphies uniting †*D. rugosus* (=*schwimmeri*) and equivalent-aged gobidontan crocodylians with Pan-*Alligatoridae* are: subequal anterior processes of surangular (54:1 in Cossette and Brochu (Cossette and Brochu 2020)), foramen aerum of articular inset from retroarticular process margin (63:1 in Cossette and Brochu (Cossette and Brochu 2020)), all dentary teeth occlude lingual to maxillary teeth (82:0 in Cossette and Brochu (Cossette and Brochu 2020)), high quadratojugal spine sits between posterior and superior infratemporal fenestra angles (121:1 in Cossette and Brochu (Cossette and Brochu 2020)).

**Stratigraphic Horizon**: The oldest definitive occurrences of †*Deinosuchus rugosus* are from the Ellisdale Site, Monmouth County, New Jersey (Schwimmer 2002; Milàn et al. 2010; Brownstein 2018; Brownstein 2019) (basal Marshalltown Formation, Campanian, between 75.7 and 71.2 Ma, though possibly older)(Miller et al. 2004; Denton 2022) and various localities corresponding to the Tar Heel Formation of North Carolina (Campanian, 78.7-74.5 Ma)(Schwimmer 2002; Cossette and Brochu 2020; Anon). Equivalently old fossils of †*Deinosuchus* are also known from western North America (Schwimmer 2002; Mohler et al. 2021).

**Equivalent Fossil Calibrations:** Various, dependent on phylogenetic topology (see earlier). **Fossil tip age**: 75.7 Ma. The node calibration was created such that 97.5% of the probability distribution fell before 75.7 Ma.

*†Asteriornis maastrichtensis*

**Justification of Placement:** †*Asteriornis maastrichtensis* calibrates the MRCA of *Neognathae* (specifiers: *Columbia livia*, *Gallus gallus*) in our node-dating analyses. The placement of †*A. maastrichtensis* within crown *Neognathae* as a member of Pan-*Galloanserae* is supported by parsimony and Bayesian phylogenetic analyses of morphological characters, including the 297 character, 39 taxon matrix of Field et al. (Field et al. 2020), including as modified by Houde et al. (Houde et al. 2023) and Crane et al. (Crane et al. 2024) and the 234 character, 49 taxon matrix of Benito et al. (Benito, Kuo, et al. 2022) (as modified from Benito et al. (Benito, Chen, et al. 2022) and thus Torres et al. (Torres et al. 2021)). For the purposes of this justification, we rely on the phylogeny presented in figure 5 of Crane et al. (Crane et al. 2024). Crane et al. (Crane et al. 2024) find †*A. maastrichtensis* to be a possible pan-galliform in agreement with some trees resolved by Field et al. (Field et al. 2020); however, we conservatively use this taxon to calibrate the MRCA of *Neognathae* rather than *Galloanserae* given its placement on the stem of *Galloanserae* in other analyses (Benito, Chen, et al. 2022; Benito, Kuo, et al. 2022).

Phylogenetically optimized apomorphies uniting †*A. maastrichtensis* with Pan-*Neognathae* are: poorly define impression for cranial cruciate ligament (221:1 in Crane et al. (Crane et al. 2024)). Phylogenetically optimized characters uniting †*A. maastrichtensis* with *Neognathae* are: subcapitular tubercle on quadrate present (50:1 in Crane et al. (Crane et al. 2024)).

Phylogenetically optimized characters uniting †*A. maastrichtensis* with *Galloanserae* are: long, narrow, and dorsally oriented medial mandibular process (68:1 in Crane et al. (Crane et al. 2024)).

**Stratigraphic Horizon**: CBR-Romontbos Quarry, Eben-Emael, Liège, Belgium(Field et al. 2020); Maastricht Formation, late Maastrichtian, between approximately 69.5 and 66.5 Ma (Kroth et al. 2024). Field et al. (Field et al. 2020) reported that the Valkenburg Member of the Maastricht Formation had yielded holotype and only known specimen of †*Asteriornis maastrichtensis*. However, the division of the Maastricht Formation into six members has recently been challenged (Kroth et al. 2024). Because the Valkenburg Member generally corresponds to lower horizons in the Maastricht Formation, we use an age of 68.0 Ma, which is approximately the midpoint of the Cretaceous portion of this unit (see (Kroth et al. 2024)).

**Equivalent Fossil Calibrations: None.**

**Fossil tip age**: 68.0 Ma. The node calibration was created such that 97.5% of the probability distribution fell before 68.0 Ma.

*†Conflicto antarcticus*

**Justification of Placement:** †*Conflicto antarcticus* calibrates the MRCA of crown *Galloanserae* (specifiers: *Anas platyrhynchos*, *Gallus gallus*) in our node-dating analyses. The placement of †*C. antarcticus* in crown *Galloanserae* in Pan-*Anseriformes* is supported by Bayesian and parsimony phylogenetic analyses of morphological characters, including 290 character, 51 taxon matrix of Tambussi et al. (Tambussi et al. 2019), the 297 character, 39 taxon matrix of Field et al. (Field et al. 2020), including as modified by Houde et al. (Houde et al. 2023) and Crane et al. (Crane et al. 2024), the 719 character, 41 taxon matrix of Musser and Clarke (Musser and Clarke 2024), and the 100 character, 34 taxon matrix of McInerney et al. (McInerney et al. 2024). For the purposes of this justification, we rely on the phylogeny presented in figure 14 of Tambussi et al. (Tambussi et al. 2019). Phylogenetically optimized apomorphies uniting †*Conflicto antarcticus* with Pan-*Anseriformes* are: caudally divergent borders of premaxilla (3:0 in Tambussi et al. (Tambussi et al. 2019)), processus basipterygoidei on sides of rostrum parasphenoidale anterior to caudal end of rostrum (28:2 in Tambussi et al. (Tambussi et al. 2019)), fonticuli present on occipital region of skull (30:1 in Tambussi et al. (Tambussi et al. 2019)), distinct foramen pneumaticum caudomediale present on processus oticus of the quadrate (51:1 in Tambussi et al. (Tambussi et al. 2019)), long, narrow interramal region on mandibular rami (69:1 in Tambussi et al. (Tambussi et al. 2019)), processus ventralis of the axis pronounced (75:1 in Tambussi et al. (Tambussi et al. 2019)), foramen enclosed by processus transversus and processus articularis caudalis of third cervical vertebra absent (76:0 in Tambussi et al. (Tambussi et al. 2019)), os metacarpale minor extends dorsal to ventral rim of trochlea carpalis (159:0 in Tambussi et al. (Tambussi et al. 2019)), length of fornix between the os metacarpale minor and major exceeds width (171:1 in Tambussi et al. (Tambussi et al. 2019)), sulcus patellaris of femur narrow and deep (213:1 in Tambussi et al. (Tambussi et al. 2019)), and pronounced medial displacement of the condylus medialis compared to facies medialis on tibiotarsus (244:1 in Tambussi et al. (Tambussi et al. 2019)).

**Stratigraphic Horizon**: Southern part of Seymour Island, near Punta Pingüino on the Weddel Sea, Antarctica; López de Bertodano Formation, Danian Stage of the Paleogene, 64.0–61.1 Ma (Tambussi et al. 2019).

**Equivalent Fossil Calibrations: None.**

**Fossil tip age**: 61.1 Ma. The node calibration was created such that 97.5% of the probability distribution fell before 61.1 Ma.

†*Eogrus* spp.

**Justification of Placement:** †*Eogrus* spp. calibrates the MRCA of *Paleognathae* (specifiers: *Struthio camelus*, *Tinamus guttatus*) in our node-dating analyses. The placement of †*Eogrus* spp. in crown *Paleognathae* as a member of Pan-*Struthio* is supported by the following characters, including the presence of shortened pedal phalanges II-2 through II-4, presence of a tubercle on the distal end of the tibiotarsus, an elongated tarsometatarsus, and the reduction of the trochlea of metatarsal II such that it is smaller than the trochlea of metatarsal IV (Mayr and Zelenkov 2021). **Stratigraphic Horizon**: Various localities, Shara Murun region, China; Ulan Shireh and Irdin Manha formations, middle Eocene (Russell 1987; Clarke et al. 2005; Mayr and Zelenkov 2021). We assign an age of 47.8 Ma following the age of the oldest †*Eogrus* fossils in figure 7 of Mayr and Zelenkov (Mayr and Zelenkov 2021).

**Equivalent Fossil Calibrations:** Dependent on phylogeny. Members of the †*Palaeotididae* and †*Lithornithidae* are older than †*Eogrus* but have highly unstable positions in paleognath phylogeny relative to the crown group (Mayr 2015; Nesbitt and Clarke 2016; Worthy et al. 2017; Field et al. 2020; Houde et al. 2023; McInerney et al. 2024; Musser and Clarke 2024).

**Fossil tip age**: 47.8 Ma. The node calibration was created such that 97.5% of the probability distribution fell before 47.8 Ma.

*†Tsidiiyazhi abini*

**Justification of Placement:** †*Tsidiiyazhi abini* calibrates the MRCA of Pan-*Coliiformes* with other *Neoaves* (specifiers: *Colius striatus*, *Cuculus canorus*). This is a conservative fossil calibration that results from (1) the lack of Cretaceous to earliest Danian neoavian fossils assignable to that clade based on phylogenetic analyses of morphological characters, and (2) the high degree of uncertainty surrounding the relationships of neoavian orders in our phylogeny that causes the topology presented in Figure 1 to disagree with previous analyses of avian relationships (Hackett et al. 2008; Jarvis et al. 2014; Prum et al. 2015; Reddy et al. 2017; Kimball et al. 2019; Kuhl et al. 2021; Berv et al. 2024; Stiller et al. 2024). We refer the reader to recently published phylogenies of *Neoaves* based on larger avian-specific loci sets for resolution of backbone neoavian relationships (Stiller et al. 2024). The placement of †*T. abini* in *Neoaves* as a member of Pan-*Coliiformes* is supported by parsimony analyses of morphological characters, including the 111 character, 48 taxon matrix of Ksepka et al. (Ksepka et al. 2017) and the 146 character, 62 taxon matrix of Ksepka et al. (Ksepka et al. 2019) For the purposes of this justification, we rely on the phylogeny presented in figure 2 of Ksepka et al. (Ksepka et al. 2019). Because no character state optimizations were provided by Ksepka et al. (Ksepka et al. 2019) for the placement of †*T. abini* in *Neoaves* as a member of Pan-*Coliiformes*, we reran the matrix presented in the paper using TNT v 1.5 (Goloboff and Catalano 2016) to optimize apomorphies; we ran an initial Wagner search over 1000 replicates and default parameters for ratchet, tfuse, drift, and sectorial search followed by traditional bisection reconnection branch swapping over 100,000 trees. We summarized the topologies in a strict consensus and then recorded apomorphies of Pan-*Coliiformes.* Phylogenetically optimized apomorphies uniting †*T. abini* with Pan-*Coliiformes* are: processus acrocoracoideus of coracoid with hooked outline (49:1 in Ksepka et al. (Ksepka et al. 2019)) and crista trochanteris of femur does not surpass level of the femoral head (96:1 in Ksepka et al. (Ksepka et al. 2019)).

**Stratigraphic Horizon**: NMMNH locality L-6898, West Flank of Torreon Wash, San Juan Basin, Sandoval County, New Mexico, USA; Ojo Encino Member, Nacimiento Formation (Ksepka et al. 2017), between 62.221–62.517 Ma (Ogg et al. 2016; Ksepka et al. 2017).

**Equivalent Fossil Calibrations: None.**

**Fossil tip age**: 62.221 Ma. The node calibration was created such that 97.5% of the probability distribution fell before 62.221 Ma.

**†Waimanu manneringi**

**Justification of Placement:** †*Waimanu manneringi* calibrates the MRCA of *Pygoscelis adeliae* and *Phalacrocorax brasilianus* in our node-dating analyses. The placement of †*W. manneringi* in Pan-*Spheniscidae* is supported by Bayesian and parsimony phylogenetic analyses of morphological characters, including the 148 character, 49 taxon matrix of Slack et al.(Slack et al. 2006), the 181 character, 50 taxon matrix of Ksepka et al. (Ksepka et al. 2006), the 75 character, 50 taxon matrix of Mayr et al. (Mayr, De Pietri, et al. 2017), the 274 character, 66 taxon matrix of Thomas et al. (Thomas et al. 2020), the 281 character, 74 taxon matrix of Cole et al. (Cole et al. 2022), the 279 character, 66 taxon matrix of Ksepka et al. (Ksepka et al. 2023), and the 281 character, 70 taxon matrix of Thomas et al.(Thomas et al. 2023) (including as modified by Mayr et al. (Mayr, Pietri, et al. 2017)). For the purposes of this justification, we rely on Figure S3 of Slack et al. (Slack et al. 2006). Phylogenetically optimized characters uniting †*W. manneringi* with Pan-*Sphenisciformes* are: thoracic vertebrae variably not heterocoelous (57:0 in Slack et al. (Slack et al. 2006)), synsacrum composed of 11 to 12 ankylozed vertebrae (91:1 in Slack et al. (Slack et al. 2006)), and poorly developed hypotarsal crests and grooves on tarsometatarsus (103:0 in Slack et al. (Slack et al. 2006)).

**Stratigraphic Horizon**: Waipara River, New Zealand; Basal Waipara Greensand, Paleocene, 60.5–61.6 Ma (Slack et al. 2006).

**Equivalent Fossil Calibrations:** None, though several other stem-penguins are known from slightly younger horizons in the Waipara Greensand (Slack et al. 2006; Mayr, Pietri, et al. 2017). **Fossil tip age**: 60.5 Ma. The node calibration was created such that 97.5% of the probability distribution fell before 60.5 Ma.

*†Protopsephurus liui*

**Justification of Placement:** †*Protopsephurus liui* calibrates the MRCA of *Acipenseriformes* (*Polyodon spathula*, *Acipenser ruthenus*) in our node-dating analyses. The placement of †*P. liui* in *Acipenseriformes* on the stem of *Polyodontidae* is supported by parsimony phylogenetic analyses of morphological characters, including the 44 character, 10 taxon matrix of Grande et al. (Grande et al. 2002) and the 62 character, 18 taxon matrix of Hilton et al. (Hilton et al. 2011). For the purposes of this justification, we rely on the phylogeny presented in figure 20 of Grande et al. (Grande et al. 2002). Phylogenetically optimized characters supporting the placement of †*P. liui* as a member of Pan-*Polyodontidae* are: body scales tiny and non-overlapping or absent (4:1 in Grande et al. (Grande et al. 2002)), dorsal end of quadratojugal with anterior projection (9:1 in Grande et al. (Grande et al. 2002)), elongate dorsal median rostral bones present (21:1 in Grande et al. (Grande et al. 2002)), well developed anterior and posterior divisions of fenestra longitudinalis present (24:1 in Grande et al. (Grande et al. 2002)), stellate bones supporting paddle present (26:1 in Grande et al. (Grande et al. 2002)), and parietals extend posterior to posttemporals (32:1 in Grande et al. (Grande et al. 2002)).

**Stratigraphic Horizon**: Songzhangzi and Fanzhangzi area, Lingyuan City, western Liaoning Province, China; Dawangzhangzi Beds, Yixian Formation, Jehol Group, Aptian-Albian (Hilton et al. 2011). We use an age of 121.4 Ma, which represents the boundary for these two stages (Gradstein et al. 2021).

**Equivalent Fossil Calibrations: None.**

**Fossil tip age**: 121.4 Ma. The node calibration was created such that 97.5% of the probability distribution fell before 121.4 Ma.

*†Watsonulus eugnathoides*

**Justification of Placement:** †*Watsonulus eugnathoides* calibrates the MRCA of *Holostei* (specifiers: *Amia calva*, *Lepisosteus osseus*) in our node-dating analyses. The placement of †*W. eugnathoides* in *Holostei* as a member of Pan-*Amia* is supported by Bayesian and parsimony analyses of morphological characters, including the 90 character, 37 taxon matrix of López-Arbarello (López-Arbarello 2012), the 96 character, 28 taxon matrix of Xu et al. (Xu et al. 2014), the 265 character, 93 taxon matrix of Giles et al. (Giles et al. 2017), the 339 character, 99 taxon matrix of López-Arbarello and Sferco (López-Arbarello and Sferco 2018), the 275 character, 97 taxon dataset of Argyriou et al. (Argyriou et al. 2018), the 224 character, 60 taxon dataset of Xu (Xu 2019), and the 300 character, 117 taxon matrix of Argyriou et al. (Argyriou et al.). For the purposes of this justification, we rely on the phylogeny presented in figure 11 of Argyriou et al. (Argyriou et al.). Phylogenetically optimized apomorphies uniting †*W. eugnathoides* with Pan-*Amia* are: posterior maxillary notch present (75:1 in Argyriou et al. (Argyriou et al.)), single supramaxilla (76:1 in Argyriou et al. (Argyriou et al.)), accessory hyoid element involved in jaw joint (97:1 in Argyriou et al. (Argyriou et al.)), foramen for abducens nerve (VI) level with optic foramen (II) (135:1 in Argyriou et al. (Argyriou et al.)), anterior ossification of ceratohyal unconstructed (220:0), and jaw articulation posterior to orbit (280:0 in Argyriou et al. (Argyriou et al.)).

**Stratigraphic Horizon**: Ambilombe, Madagascar; Sakamena Formation, Early Triassic, Induan (Olsen 1984). We use an age of 251.2 Ma, which is the base of the Induan (Gradstein et al. 2021), following previous studies (Benton et al. 2015).

**Equivalent Fossil Calibrations: None.**

**Fossil tip age**: 251.2 Ma. The node calibration was created such that 97.5% of the probability distribution fell before 251.2 Ma.

*†Atractosteus falipoui*

**Justification of Placement:** †*Atractosteus falipoui* calibrates the MRCA of *Lepisosteidae* (specifiers: *Atractosteus tropicus*, *Lepisosteus osseus*) in our node-dating analyses. The placement of †*A. falipoui* in crown *Lepisosteidae* as a member of Pan-*Atractosteus* is supported by Bayesian and parsimony analyses of morphological characters, including the 105 character, 32 taxon matrix of Grande (Grande 2010) (including as modified by Brito et al. (Brito et al. 2017) and Brownstein and Lyson) (Brownstein and Lyson 2022), and the 104 character, 38 taxon matrix of Brownstein et al. (Chase Doran Brownstein et al. 2023). For the purposes of this justification, we rely on the phylogeny presented in supplementary figure 3 of Brito et al. (Brito et al. 2017). Phylogenetically optimized apomorphies that unite †*A. falipoui* with Pan-*Atractosteus* are: collective shape of laterally expanded vomerine heads form a shape roughly like an equilateral triangle (40:1 in Grande (Grande 2010)/Brito et al. (Brito et al. 2017)), tooth plates associated with second and third hypobranchials (80:1 in Grande (Grande 2010)/Brito et al. (Brito et al. 2017)), and anterior end of first coronoid curves medially and expands broadly to a flat symphysis (104:1 in Grande (Grande 2010)/Brito et al. (Brito et al. 2017)).

**Stratigraphic Horizon**: Various localities, Kem Beds, Morocco; Auoufous Formation, Cenomanian (Grande 2010). We use an age of 97.2 Ma following previous studies (Brownstein and Lyson 2022; Chase Doran Brownstein et al. 2023).

**Equivalent Fossil Calibrations: None.**

**Fossil tip age**: 97.2 Ma. The node calibration was created such that 97.5% of the probability distribution fell before 97.2 Ma.

*†Ichthyemidion vidali*

**Justification of Placement:** †*Ichthyemidion vidali* calibrates the MRCA of *Elopomorpha* (specifiers: *Megalops atlanticus*, *Anguilla anguilla*) in our node-dating analyses. The placement of †*I. vidali* in *Elopomorpha* on the stem of *Elopiformes* is supported by Bayesian and parsimony analyses, including the morphological + molecular dataset of Dornburg et al. (Dornburg et al. 2015), the 74 character, 27 taxon dataset of Figueiredo et al. (Hernández-Guerrero et al. 2021), and the 75 character, 30 taxon dataset of Hernández-Guerrero et al. (Hernández-Guerrero et al. 2021). Given the considerable uncertainty surrounding the ages and phylogenetic positions of Late Jurassic to Early Cretaceous members of *Elopomorpha* and its constituent clades (Arratia 2000; de Figueiredo et al. 2012; Dornburg et al. 2015; Davesne et al. 2021), this is a conservative fossil calibration. For the purposes of this justification, we rely on the phylogeny presented in figure 6 of Hernández-Guerrero et al. (Hernández-Guerrero et al. 2021). Because no character state optimizations were provided by Hernández-Guerrero et al. (Hernández-Guerrero et al. 2021) for the placement of †*I. vidali* in *Elopomorpha* as a member of Pan-*Elopiformes*, we reran the matrix presented in the paper using TNT v 1.5 (Goloboff and Catalano 2016) to optimize apomorphies; we ran an initial Wagner search over 1000 replicates and default parameters for ratchet, tfuse, drift, and sectorial search followed by traditional bisection reconnection branch swapping over 100,000 trees. We summarized the topologies in a strict consensus and then recorded apomorphies of *Elopomorpha*, Pan-*Elopiformes*, and the clade containing †*I. vidali* + *Elopiformes*. Phylogenetically optimized apomorphies that unite †*I. vidali* with *Elopomorpha* are: compound neural arch over Pu1 and U1 present (62:1 in Hernández-Guerrero et al. (Hernández-Guerrero et al. 2021)), fringing fulcra present (62:1 in Hernández-Guerrero et al. (Hernández-Guerrero et al. 2021)), and pelvic splint present (69:1 in Hernández-Guerrero et al. (Hernández-Guerrero et al. 2021)). Phylogenetically optimized apomorphies that unite †*I. vidali* with Pan-*Elopiformes* are: elongate jaw as bearing numerous villiform teeth (37:1 in Hernández-Guerrero et al. (Hernández-Guerrero et al. 2021)). Phylogenetically optimized apomorphies that unite †*I. vidali* and *Elopiformes* are: preopercle posteroventrally expanded (33:1 in Guerrero et al. (Hernández-Guerrero et al. 2021)).

**Stratigraphic Horizon**: El Montsec, Lérida Province, Spain; Berriasian–Barremian (Dornburg et al. 2015). We use an age of 125.0 Ma, which is the traditional upper bound of the Barremian (Gradstein et al. 2021).

**Equivalent Fossil Calibrations:** Dependent on the phylogenetic resolution of fossils placed stemward in Pan-*Elopiformes* in the phylogeny presented by Hernández-Guerrero et al. (Hernández-Guerrero et al. 2021).

**Fossil tip age**: 125.0 Ma. The node calibration was created such that 97.5% of the probability distribution fell before 125.0 Ma.

*†Laeliichthys ancestralis*

**Justification of Placement:** †*Laeliichthys ancestralis* calibrates the MRCA of *Osteoglossiformes* (specifiers: *Pantodon buchholzi*, *Osteoglossum bicirrhosum*) in our node-dating analyses. The placement of †*L. ancestralis* in crown *Osteoglossiformes* is supported by Bayesian and parsimony analyses of morphological characters, including the 87 character, 28 taxon matrix of Brito et al.,(Brito et al. 2020) the 96 character, 34 taxon matrix of Capobianco et al. (Capobianco et al. 2025), and the 96 character, 56 taxon matrix of Capobianco and Friedman (Capobianco and Friedman 2024). For the purposes of this justification, we rely on the phylogeny presented in Figure S1 of Brito et al. (Brito et al. 2020). Phylogenetically optimized apomorphies that unite †*L. ancestralis* with crown *Osteoglossiformes* are: supratemportal commissure passes through the parietals (10:1 in Brito et al. (Brito et al. 2020)), dorsal arm of posttemporal less than 1.5x as long as ventral arm (57:0 in Brito et al. (Brito et al. 2020)), and epurals absent (68:2 in Brito et al.(Brito et al. 2020)).Phylogenetically optimized apomorphies that unite †*L. ancestralis* with Pan-*Notopteridae* are: extrascapular reduced and irregularly shaped (2:1 in Brito et al. (Brito et al. 2020)), frontal anterior margin width equal to posterior margin width (3:1 in Brito et al. (Brito et al. 2020)), nasal bones meet at midline (6:2 in Brito et al. (Brito et al. 2020)), subopercle absent (35:2 in Brito et al. (Brito et al. 2020)), posterior portion of maxilla lies on angular (39:0 in Brito et al. (Brito et al. 2020)), and first infrapharyngobranchial present (53:1 in Brito et al. (Brito et al. 2020)).

**Stratigraphic Horizon**: Quarries of São José do Geribá farmstead, near Patos de Minas Sanfranciscana Basin, Minas Gerais, Brazil (Brito et al. 2020); Upper Quiricó Formation, Barremian-Aptian (Brito et al. 2020; Celerino de Carvalho and Santucci 2024), which range from 125.77 to 113.0 Ma (Gradstein et al. 2021). We use an age of 113.0 Ma.

**Equivalent Fossil Calibrations: None.**

**Fossil tip age**: 113.0 Ma. The node calibration was created such that 97.5% of the probability distribution fell before 113.0 Ma.

*†Tischlingerichthys viohli*

**Justification of Placement:** †*Tischlingerichthys viohli* calibrates the MRCA of *Otocephala* (specifiers: *Clupea harengus*, *Danio rerio*) in our node-dating analyses. The placement of †*T. viohli* in crown *Otocephala* is supported by parsimony analyses of morphological characters, including the 74 character, 27 taxon dataset of Figueiredo et al. (de Figueiredo et al. 2012) and the 75 character, 31 taxon matrix of L-Recinos et al. (L-Recinos et al. 2023). For the purposes of this justification, we rely on the phylogeny presented in figure 15 of L-Recinos et al. (L-Recinos et al. 2023). Because no character state optimizations were provided by L-Recinos et al. (L-Recinos et al. 2023) for the placement †*T. viohli* in crown *Otocephala* as a member of Pan-*Ostariophysi*, we reran the matrix presented in the paper using TNT v 1.5 (Goloboff and Catalano 2016) to optimize apomorphies; we ran an initial Wagner search over 1000 replicates and default parameters for ratchet, tfuse, drift, and sectorial search followed by traditional bisection reconnection branch swapping over 100,000 trees. We summarized the topologies in a strict consensus and then recorded apomorphies of *Otocephala* and Pan-*Ostariophysi*.

Phylogenetically optimized apomorphies that unite †*T. viohli* with crown *Otocephala* are: retroarticular excluded from jaw joint (42:1 in L-Recinos et al. (L-Recinos et al. 2023)). Phylogenetically optimized apomorphies that unite †*T. viohli* with Pan-*Ostariophysi* are: parietal shorter than wide (2:1 in L-Recinos et al. (L-Recinos et al. 2023)), preopercle posteroventrally expanded (33:1 in L-Recinos et al. (L-Recinos et al. 2023)), ventral margin of quadrate horizontal (39:1 in L-Recinos et al. (L-Recinos et al. 2023)), and first preural and first ural fused (59:1 in L-Recinos et al. (L-Recinos et al. 2023)).

**Stratigraphic Horizon**: Mühlheim, Bavaria, Germany; Mörnsheim Formation, lower Tithonian (Arratia 2000). We use an age of 151.5 Ma, which is the maximum upper bound on the age of the Tithonian (Gradstein et al. 2021).

**Equivalent Fossil Calibrations: None.**

**Fossil tip age**: 151.5 Ma. The node calibration was created such that 97.5% of the probability distribution fell before 151.5 Ma.

*†Rubiesichthys gregalis*

**Justification of Placement:** †*Rubiesichthys gregalis* calibrates the MRCA of *Ostariophysi* (specifiers: *Chanos chanos*, *Danio rerio*) in our node-dating analyses. The placement of †*R. gregalis* in *Ostariophysi* as a member of *Gonorynchiformes* is supported by Bayesian and parsimony analyses of morphological characters, including the 94 character, 22 taxon matrix of Grande and Poyato-Ariza (GRANDE and POYATO-ARIZA 1999), the 106 character, 14 taxon matrix of Ribiero et al. (Ribeiro et al. 2018), the 130 character, 22 taxon matrix of Grande et al. (Diogo 2009) (including as modified by Amaral et al. (Amaral et al. 2013)), and the 128 character, 20 taxon matrix used with molecular data by Near et al. (Near et al. 2014). For the purposes of this justification, we rely on the phylogeny presented in figure 7.10 of Grande et al. (Diogo 2009). Phylogenetically optimized apomorphies that unite †*R. gregalis* with *Gonorynchiformes* are: orbitosphenoid absent (1:1 in Grande et al. (Diogo 2009)), parietals completely separated by supraoccipital (13:2 in Grande et al. (Diogo 2009)), reduced, flat, and blade-like parietals (15:1 in Grande et al. (Diogo 2009)), teeth absent in premaxilla, maxilla, dentary (20:1 in Grande et al. (Diogo 2009)), premaxillary ascending process absent (24:1 in Grande et al. (Diogo 2009)), and rib on third vertebral centrum wide and short (82:1 in Grande et al. (Diogo 2009)).

**Stratigraphic Horizon**: El Montsec and Las Hoyas, Lérida and Cuenca Provinces, Spain (Poyato-Ariza 1996a; Poyato-Ariza 1996b); Berriasian–Barremian (Dornburg et al. 2015). El Montsec is Barriasian through lower Barremian, whereas Las Hoyas is Barremian (Gil-Delgado et al. 2023; Marugán-Lobón et al. 2023). We use an age of 121.4 Ma, which is the upper bound of the Barremian (Gradstein et al. 2021).

**Equivalent Fossil Calibrations:** †*Gordichthys conquensis*, also from the Early Cretaceous of Spain, is a Pan-*Chanos* and is an equivalent fossil calibration (Poyato-Ariza 1996a).

**Fossil tip age**: 121.4 Ma. The node calibration was created such that 97.5% of the probability distribution fell before 121.4 Ma.

*†Santanichthys diasii*

**Justification of Placement:** †*Santanichthys diasii* calibrates the MRCA of *Otophysi* (specifiers: *Corydoras julii*, *Danio rerio*) in our node-dating analyses. The placement of †*S. diasii* in *Otophysi* is supported by the following characters: basisphenoid absent, dermopalatine absent, expanded dorsomedial portion of anterior neural arches, the expansion of the anterior supraneural, modification of the first neural arch into the scaphium, modification of the second neural arch into the intercalarium, first four centra shortened, and modification of the third centrum rib parapophysis into the tripus (Filleul and Maisey 2004).

**Stratigraphic Horizon**: Araripe Basin, northeastern Brazil; Santana Formation, Albian Stage of the Early Cretaceous.(Filleul and Maisey 2004) Following previous studies, we use an age of 112.5 Ma (Chase Doran Brownstein et al. 2023; Brownstein and Near 2024).

**Equivalent Fossil Calibrations:** None.

**Fossil tip age**: 112.5 Ma. The node calibration was created such that 97.5% of the probability distribution fell before 112.5 Ma.

*†Wilsonium brevipinne*

**Justification of Placement:** †*Wilsonium brevipinne* calibrates the MRCA of *Cypriniformes* (specifiers: *Gyrinocheilus aymonieri*, *Danio rerio*) in our node-dating analyses. The placement of †*W. brevipinne* in *Cypriniformes* as a member of Pan-*Catostomidae* is supported by parsimony analysis of morphological characters, including the 83 character, 71 taxon matrix of Liu et al (Liu 2021). For the purposes of this justification, we rely on the phylogeny presented in figure 6 of Liu et al (Liu 2021). Phylogenetically optimized apomorphies that unite †*W. brevipinne* with Pan-*Catostomidae* are: pharyngeal toothplate falcate (Liu 2021).

**Stratigraphic Horizon**: North Fork of the Similkameen River, Pleasant Valley, British Columbia, Canada; Allenby Formation, Eocene (Liu 2021). Following Bagley et al. (Bagley et al. 2018), we use an age of 48.88 Ma for this taxon.

**Equivalent Fossil Calibrations:** Various pan-catostomids, see Bagley et al. (Bagley et al. 2018). **Fossil tip age**: 48.88 Ma. The node calibration was created such that 97.5% of the probability distribution fell before 48.88 Ma.

†*Siluroidei* indet.

**Justification of Placement:** Indeterminate siluroid fossils calibrate the MRCA of *Siluriformes* (specifiers: *Corydoras julii*, *Silurus asotus*) in our node-dating analyses. The placement of these fossils, which include the apomorphic pectoral fin spines of catfishes, in the crown clade as potential representatives of living catfish families is supported by the presence of large, compressed, and hooked posterior dentations and the absence of anterior dentations (Alves et al. 2019). Although Alves et al. (Alves et al. 2019) suggested affinities with several families of siluroid siluriform for these elements, we conservatively use them to calibrate the catfish crown group as they were not included in a phylogenetic analysis.

**Stratigraphic Horizon**: Santo Anastácio, São Paulo State, Brazil; Adamantina Formation, Turonian-Santonian Stages of the Late Cretaceous (Alves et al. 2019), which range from 93.9 to 83.6 Ma (Alves et al. 2019).

**Equivalent Fossil Calibrations:** Various isolated remains, see Alves et al. (Alves et al. 2019) for a review.

**Fossil tip age**: 83.6 Ma. The node calibration was created such that 97.5% of the probability distribution fell before 83.6 Ma.

*†Cretazeus rinaldi*

**Justification of Placement:** †*Cretazeus rinaldi* calibrates the MRCA of *Zeiogadaria* (specifiers: *Gadus morhua*, *Zeus faber*) in our node-dating analyses. The placement of †*C. rinaldi* in *Zeiogadaria* as a member of Pan-*Zeiformes* is supported by parsimony analyses of morphological characters, including the 107 character, 43 taxon matrix of Tyler and Santini (Tyler and Santini 2005) (including as modified by Davesne et al. (Davesne et al. 2017)) and the combined dataset (including 105 characters and 40 taxa) of Grande et al. (Grande et al. 2018).

For the purposes of this justification, we rely on the phylogeny presented in figure 7 of Tyler and Santini (Tyler and Santini 2005). Phylogenetically optimized apomorphies uniting †*C. rinaldi* with Pan-*Zeiformes* are: multiple serrations present on lower border of dentary (22:2 in Tyler and Santini (Tyler and Santini 2005)), parahypural slightly removed from and does not embrace urostylar centrum (52:1 in Tyler and Santini (Tyler and Santini 2005)), absence of uroneural (54:1 in Tyler and Santini (Tyler and Santini 2005)), and two anal fin spines present (100:2 in Tyler and Santini (Tyler and Santini 2005)).

**Stratigraphic Horizon**: Nardò, Italy; Santonian-Maastrichtian Stages of the Late Cretaceous (Tyler and Santini 2005; Santaquiteria et al. 2021). Although traditionally considered to be Campanian-Maastrichtian, there is some evidence that the fossils at Nardò are actually older, from the Santonian (Santaquiteria et al. 2021). As such, we assign a basal Campanian age of 83.0 Ma to this fossil (Santaquiteria et al. 2021). Note that we assign an age of 86.0 Ma to †*Gasterorhamposus zuppichini*, as fossils of that taxon, which also appears at Nardò, may extend into the Santonian (Santaquiteria et al. 2021).

**Equivalent Fossil Calibrations:** None.

**Fossil tip age**: 83.0 Ma. The node calibration was created such that 97.5% of the probability distribution fell before 83.0 Ma.

*†Iridopristis parrisi*

**Justification of Placement:** †*Iridopristis parrisi* calibrates the MRCA of *Beryciformes* (specifiers: *Beryx splendens*, *Sargocentron rubrum*) in our node-dating analyses. The placement of †*Iridopristis parrisi* in *Beryciformes* as a member of Pan-*Holocentridae* is supported by Bayesian and parsimony analysis of morphological characters, including the 82 character, 18 taxon matrix of Andrews et al. (Andrews et al. 2023). Phylogenetically optimized apomorphies that unite †*I. parrisi* with Pan-*Holocentridae* are: transverse crest present on supraoccipital (2:1 in Andrews et al. (Andrews et al. 2023)), separate opening present for orbital branch of supraorbital sensory canal (3:1 in Andrews et al. (Andrews et al. 2023)), ventrolateral wing-like expansion of parasphenoid present (6:1 in Andrews et al. (Andrews et al. 2023)), anterior ceratohyal imperforate (8:1 in Andrews et al. (Andrews et al. 2023)), presence of deep notches along ventral margin of anterior ceratohyal (9:1 in Andrews et al. (Andrews et al. 2023)), alveolar platform near symphysis of dentary expanded and overhangs lateral margin of bone (27:1 in Andrews et al. (Andrews et al. 2023)), and presence of a concave premaxillary tooth gap at premaxillary median margin ventral to the premaxillary ascending process (28:1 in Andrews et al. (Andrews et al. 2023)).

**Stratigraphic Horizon**: Inversand Quarry (now the Edelman Fossil Park), Sewell, New Jersey, USA; Main Fossiliferous Layer, Hornerstown Formation, earliest Danian, Paleocene (Andrews et al. 2023), approximately 66.02 Ma (Gradstein et al. 2021).

**Equivalent Fossil Calibrations:** None.

**Fossil tip age**: 66.02 Ma. The node calibration was created such that 97.5% of the probability distribution fell before 66.02 Ma.

*†Gasterorhamposus zuppichini*

**Justification of Placement:** †*Gasterorhamposus zuppichini* calibrates the MRCA of *Syngnathiformes* (specifiers: *Aulostomus maculatus*, *Hippocampus erectus*) in our node-dating analyses. The placement of †*G. zuppichini* in *Syngnathiformes* is supported by parsimony and Bayesian analyses of morphological characters, including the 101 character, 36 taxon dataset of Brownstein (C D Brownstein 2023). For the purposes of this justification, we rely on the phylogeny presented in Figure S2 of Brownstein (C D Brownstein 2023). Phylogenetically optimized apomorphies that unite †*G. zuppichini* with *Syngnathiformes* are: infraorbital bones do not form a complete ventral ring (6:1 in Brownstein (C D Brownstein 2023)), parietals absent (9:1 in Brownstein (C D Brownstein 2023)), preopercle articular socket for interhyal present (34:1 in Brownstein (C D Brownstein 2023)), and pelvic spines absent (77:1 in Brownstein (C D Brownstein 2023)).

**Stratigraphic Horizon**: Porto Selvaggio, Lecce province, Italy; “Calcari di Melissano”, Santonian-Campanian (Santaquiteria et al. 2021). We use a Santonian age here (86.0 Ma).

**Equivalent Fossil Calibrations:** None.

**Fossil tip age**: 86.0 Ma. The node calibration was created such that 97.5% of the probability distribution fell before 86.0 Ma.

*†Uylyaichthys eugeniae*

**Justification of Placement:** †*Uylyaichthys eugeniae* calibrates the MRCA of *Carangioidei* (specifiers: *Caranx ignobilis*, *Coryphaena hippurus*) in our node-dating analyses. The placement of †*Uylyaichthys eugeniae* in *Carangioidei* as a member of *Carangidae* is supported by the following morphological characters: gap between last two anal fin spines (Prokofiev 2002).

**Stratigraphic Horizon**: Uylya-Kushlyuk locality, 2 km northeast of Uylya-Kushlyuk village, Turkmenistan; Danata Formation, uppermost Thanetian-lowermost Ypresian (Paleocene to Eocene) (Prokofiev 2002; Ghezelayagh et al. 2022). Following Ghezelayagh et al. (Ghezelayagh et al. 2022), we use an age of 55.8 Ma.

**Equivalent Fossil Calibrations:** †*Archaeus oblongus* and †*Trachicaranx tersus*, both from the same geological unit as †*Uylyaichthys eugeniae*, are equivalent fossil calibrations (Prokofiev 2002).

**Fossil tip age**: 55.8 Ma. The node calibration was created such that 97.5% of the probability distribution fell before 55.8 Ma.

*†Eocoelopoma portentosum*

**Justification of Placement:** †*Eocoelopoma portentosum* calibrates the MRCA of *Scombriformes* (specifiers: *Nomeus gronovii*, *Thunnus albacares*) in our node-dating analyses. The placement of †*E. portentosum* in *Scombriformes* as a member of *Scombroidea* is supported by the following morphological characters: premaxillary fangs absent, caudal hypurostegy present, two epurals present, and single uroneural present (Friedman et al. 2019).

**Stratigraphic Horizon**: Uylya-Kushlyuk locality, 2 km northeast of Uylya-Kushlyuk village, Turkmenistan; Danatinsk Formation, Thanetian to Ypresian (Friedman et al. 2019). We use an age of 54.17 Ma following previous studies (Monsch and Bannikov 2011).

**Equivalent Fossil Calibrations:** Various, see Monsch and Bannikov (Monsch and Bannikov 2011) for a review of Paleogene scombroids from northern Eurasia.

**Fossil tip age**: 54.17 Ma. The node calibration was created such that 97.5% of the probability distribution fell before 54.17 Ma.

*†Phyllopharyngodon longipinnis*

**Justification of Placement:** †*Phyllopharyngodon longipinnis* calibrates the node representing the MRCA of *Labridae* (specifiers: *Thalassoma bifasciatum*, *Scarus ghobban*). The placement of †*P. longipinnis* within *Labridae* is supported by the following morphological characters: single, posteriorly oriented supraneural; fused pharyngeal jaw present; scales cycloid (Bellwood 1990). The placement of †*P. longipinnis* in Pan-*Hypsigenyinae* is supported by the following morphological character: phyllodonty (Bellwood 1990).

**Stratigraphic Horizon**: Monte Bolca, near Verona, Italy; Early-late Ypresian, Eocene, Paleogene 48.96 to 48.5 Ma (Friedman and Carnevale 2018).

**Equivalent Fossil Calibrations**: †*Bellwoodilabrus landinii*, also from Monte Bolca, is an equivalent fish calibration (Bannikov and Carnevale 2010).

**Fossil tip age**: 48.5 Ma. The node calibration was created such that 97.5% of the probability distribution fell before 48.5 Ma.

*†Caruso brachysomus*

**Justification of Placement:** †*Caruso brachysomus* calibrates the node representing the MRCA of *Lophioidei* (specifiers: *Antennarius striatus*, *Lophius piscatorius*) in our node-dating analyses. For a detailed discussion of morphological characters uniting †*C. brachysomus* with crown *Lophioidei* and *Lophiidae*, see Brownstein et al. (Brownstein, Zapfe, et al. 2024).

**Stratigraphic Horizon**: Monte Bolca, near Verona, Italy: early-late Ypresian, Eocene, Paleogene 48.96 to 48.5 Ma (Friedman and Carnevale 2018).

**Equivalent Fossil Calibrations**: †*Eosladenia caucasica* from the Middle Eocene of the northern Caucasus (A.f 2004), the Monte Bolca antennariids †*Eophryne barbutii* (Carnevale and Pietsch 2009), †*Histionotophorus bassani*, and †*Orrichthys longimanus* (Carnevale and Pietsch 2010), and the Monte Bolca ogcocephalid †*Tarkus squirei* (Carnevale and Pietsch 2011) are all equivalent fossil calibrations.

**Fossil tip age**: 48.5 Ma. The node calibration was created such that 97.5% of the probability distribution fell before 48.5 Ma.

*†Moclaybalistes danekrus*

**Justification of Placement:** †*Moclaybalistes danekrus* calibrates the MRCA of *Tetraodontoidei* (specifiers: *Mola mola*, *Pseudobalistes fuscus*) in our node-dating analyses. The placement of †*M. danekrus* in *Tetraodontoidei* is supported by parsimony analyses of morphological characters, including the 210 character, 56 taxon dataset of Santini and Tyler (SANTINI and TYLER 2003). For the purposes of this justification, we rely on the phylogeny presented in figure 4a of Santini and Tyler (SANTINI and TYLER 2003). Phylogenetically optimized apomorphies that unite †*M. danekrus* with *Tetraodontoidei* are: infraorbitals absent (24:1 in Santini and Tyler (SANTINI and TYLER 2003)), sensory groove on dentary absent (35:1 in Santini and Tyler (SANTINI and TYLER 2003)), mouth gape moderate to small (39:1 in Santini and Tyler (SANTINI and TYLER 2003)), skull bone lateral line canals absent (66:1 in Santini and Tyler (SANTINI and TYLER 2003)), thick caniniform teeth (69:3 in Santini and Tyler (SANTINI and TYLER 2003)), second dorsal spine elongated (159:2 in Santini and Tyler (SANTINI and TYLER 2003)), first anal fin pterygiophore positioned along posterior edge of first caudal haemal spine (164:1 in Santini and Tyler (SANTINI and TYLER 2003)), single anal fin pterygiophore in first interhaemal space (166:1 in Santini and Tyler (SANTINI and TYLER 2003)), procurrent caudal fin rays absent (178:1 in Santini and Tyler (SANTINI and TYLER 2003)), and 10 to 14 anal fin rays present (209:2 in Santini and Tyler (SANTINI and TYLER 2003)).

**Stratigraphic Horizon**: Various localities, Denmark; Fur Formation, Ypresian Stage of the Eocene, 56.0 Ma (Benton and Donoghue 2007).

**Equivalent Fossil Calibrations:** None.

**Fossil tip age**: 56.0 Ma. The node calibration was created such that 97.5% of the probability distribution fell before this 56.0 Ma.

## II. Supplementary Results and Discussion

### General Observations

We conducted phylogenetic analyses of the exon dataset at different levels of completeness (all 1105 exons, and a 75% complete matrix) using both multispecies coalescent (Zhang et al. 2018) and concatenation approaches to explore how alternative strategies affected inference of the jawed vertebrate tree. Generally, we inferred similar phylogenies across different sequence matrix completeness levels and analytical methods (Extended Data Figure 1). However, phylogenies inferred from the concatenated matrix inferred widely-accepted relationships, such as monophyly of *Syngnathiformes* and the placement of *Leiognathidae* within *Eupercaria*, that the ASTRAL-III multispecies coalescent phylogenies failed to resolve. Among some of the deepest divergences in jawed vertebrates, we also observe incongruence across trees inferred using different methodologies. For example, salamanders (*Caudata*) and caecilians (*Gymnophiona*) are consistently inferred as sister clades in phylogenies built using ASTRAL-III, whereas the conventional arrangement of amphibian relationships in which frogs (*Anura*) are sister to *Caudata* is recovered in phylogenies inferred from the concatenated sequences under maximum likelihood (Extended Data Figure 1). These and other conflicts are discussed in the following sections. Generally, gene and site concordance factors, bootstrap supports, and branch lengths (in substitutions per site) show similar linear relationships across analyses of the exon matrix at different completeness levels (Extended Data Figure 2). Across the jawed vertebrate phylogenies that we infer, coalescent support values, anomalous branch counts, and gene and site concordance factors do not substantially decrease in value across different matrix completeness levels (Extended Data Figure 3). Jawed vertebrate clades sampled in our phylogeny show clear patterns in time; a pronounced drop in nodal support and branch lengths (both in substitutions per site and in millions of years) is associated with the Cretaceous-Paleogene Mass Extinction (Figure 3; Figure 4; Extended Data Figure 7). Heirarchical clustering of jawed vertebrate node ages (nodes sampled are all older than 56 Ma) also shows a clear split between nodes dating to before the Late Cretaceous and from the Late Cretaceous onward, suggesting a signature of divergence associated with the Cretaceous Terrestrial Revolution and Cretaceous-Paleogene Mass Extinction.

### Resolution of Earliest Divergences in Sarcopterygii

Our phylogenetic analyses highlight several regions of uncertainty in early jawed vertebrate relationships. We highlight two key results: the resolution of a monophyletic group containing coelacanths (*Latimeria*) and lungfishes *(Dipnoi*) that forms the sister clade to *Tetrapoda*, and the uncertainty surrounding the living sister clade to other amphibians. We consistently resolve a lungfish-coelacanth sister lineage relationship with strong nodal support across all phylogenetic analyses (Figure 2; Extended Data Figure 1). This result contrasts with many phylogenomic analyses of sarcopterygians built using genome-scale data (Chen et al. 2015; Irisarri and Meyer 2016; Irisarri et al. 2017; Hime et al. 2020; Meyer et al. 2021; Wang et al. 2021; Schartl et al. 2024). However, we note that only one of these analyses fully sampled all major early divergences in jawed vertebrates, including the four oldest-diverging clades in ray-finned fishes (*Actinopterygii*): *Polypteridae*, *Acipenseriformes*, *Holostei*, and *Oseanacephala*,(Chen et al. 2015) and even this study does not sample *Amia*, which diverged from the only other living holosteans, gars (*Lepisosteidae*), at least 247.2 million years ago (given the position of †*Watsonulus eugnathoides* on the stem of *Amia*; see fossil calibration justification). This result is also not due to uncertain resolution of conflict among genes and sites, as gene and site concordance factors estimated for the coelacanth-lungfish node are comparable to or higher than widely accepted clades, including many ray-finned fish orders, laurasiatherian mammals, lepidosaurs, and archosaurs (Figure 2). Our result highlights that the classic question of what lobe-finned fish clade is the living sister to tetrapods (Brinkmann et al. 2004; Shan and Gras 2011; Irisarri and Meyer 2016; Takezaki and Nishihara 2017; Meyer et al. 2021) is arguably still unresolved and suggests that capturing all deep divergences in jawed vertebrates affects inference of this region of the phylogeny. We also note that the rapid successive divergences of coelacanths, lungfishes, and tetrapods suggested by the fossil record and supported by molecular phylogenies (Zhu et al. 2001; Irisarri et al. 2017; Wang et al. 2021; Cui et al. 2022; Schartl et al. 2024)(Figure 1) also provide a mechanism by which factors like incomplete lineage sorting and introgression interfere with phylogenetic resolution. In our main phylogeny (Figure 1), the age of crown *Sarcopterygii* is 444.02 Ma (95% highest posterior density, HPD: 413.95, 467.73 Ma) and the age of the lungfish-coelacanth split is 417.71 Ma (95% HPD: 383.17, 445.54 Ma), which is 270 kya younger than 97.5% of the prior distribution as given by the fossil †*Youngolepis praecursor* (see earlier), though the 95% highest posterior density interval overlaps considerably with this prior specification.

### The Earliest Divergence in Amphibia

We find uncertain relationships among the three living lineages of amphibians. As noted above, the resolution of two alternative hypotheses of living amphibian relationships, the *Procera* (*Anura*, (*Caudata*, *Gymnophiona*)) and the *Batrachia* (*Gymnophiona*, (*Anura*, *Caudata*)) differs according to whether a multispecies coalescent or a concatenated approach is employed (Extended Data Figure 1). Either hypothesis is supported by bootstrap or coalescent values of 100% in trees in which *Batrachia* or *Procera* is resolved (Figure 2, Extended Data Figure 1).

Different analyses of genome scale data have highlighted inadvertent paralog inclusion and considerable gene discordance as causes for this uncertainty.(Siu-Ting et al. 2019; Hime et al. 2020) Our results are generally supportive of these hypothesized drivers of uncertain relationships among the three major living clades of amphibians. Our inference of Permo-Triassic divergences of these three lineages also supports the more recent timescale of amphibian origination proposed by Hime et al. (Hime et al. 2020). In our main time tree, we estimate an age of 263.19 Ma (95% HPD: 239.87, 290.99 Ma) for crown *Amphibia* and an age of 249.94 Ma (95% HPD: 195.86, 283.77 Ma) for crown *Batrachia*. These estimates, which suggest that the amphibian crown diversified over a period of approximately 15 million years in the early Triassic, should be viewed as conservatively young since we use a conservative age for the calibration of crown *Batrachia* (see above). In any case, they support either a latest Permian or early Triassic origin of the three living amphibian clades, which is entirely concordant with the amphibian fossil record (Evans and Borsuk-Bialynicka 1998; Anderson et al. 2008; Schoch et al. 2020; Jones et al. 2022; Kligman et al. 2023).

### Uncertainty in Mammal Order-Level Relationships and the Age of Placentals

The relationships among placental mammals, as well as their timescale of evolution, remain debated, with the principal issues concerning what clade represents the living sister to all other placentals (dos Reis et al. 2012; dos Reis et al. 2014; Tarver et al. 2016; Esselstyn et al. 2017; Álvarez-Carretero et al. 2022), the relationships among laurasiatherians (Tarver et al. 2016; Esselstyn et al. 2017; Foley et al. 2023), and whether placentals represent an earlier Paleogene diversification (Springer et al. 2003; Wible et al. 2007; Meredith et al. 2011; O’Leary et al. 2013; dos Reis et al. 2014; Phillips and Fruciano 2018; Velazco et al. 2022; Budd and Mann 2023; Foley et al. 2023). Our time-calibrated phylogenies unambiguously infer that the earliest divergences among crown placentals took place during the Late Cretaceous, between 100 and 80 million years ago (Figure 1, Extended Data Figure 4, Extended Data Figure 6). Our main time tree places the MRCA of crown placentals at 111.65 Ma (95% HPD: 92.7, 136.51 Ma). Our ASTRAL-III species trees and our concatenated maximum likelihood phylogenies disagree on what clade forms the living sister to other placental mammals; the former resolve a clade, *Atlantogenata*, containing both *Afrotheria* and *Xenarthra*, whereas the latter resolve *Xenarthra* as the living sister to all other mammals. Our time calibrated phylogeny indicates that the unclear resolution of the living sister lineage of other placentals may be a consequence of a rapid diversification of placentals in the Aptian, as the clade (*Epitheria*) containing *Afrotheria* and *Boreoeutheria* appears at 111.58 Ma (95% HPD: 92.7, 136.51 Ma). Similarly rapid diversification took place in neoavian birds after the Cretaceous-Paleogene Mass Extinction and essentially renders several regions of neoavian phylogeny unresolvable (Hackett et al. 2008; Jarvis et al. 2014; Prum et al. 2015; Reddy et al. 2017; Kuhl et al. 2021; Stiller et al. 2024). As such, resolving the basal split among crown placentals might be nearly impossible even with genomic data and require mammal-specific analyses to maximize the number of orthologous genes available.(Tarver et al. 2016; Esselstyn et al. 2017) Similarly, we are unable to robustly resolve the relationships among four major groups of laurasiatherians: bats (*Chiroptera*), whales, hippos, pigs, bovids, giraffes, camels, and relatives (*Cetoartiodactyla*), horses, tapirs, rhinoceroses, and relatives (*Perissodactyla*), and a clade containing pangolins and carnivorans (*Ferae*). The relationships of these lineages remain unresolved in other phylogenomic analyses of placentals (dos Reis et al. 2012; Tarver et al. 2016; Esselstyn et al. 2017; Foley et al. 2023). It is notable, however, that with the exception of the placement of bats as sister to *Cetoartiodactyla* rather than as the sister clade to *Cetoartiodactyla* + *Ferae* + *Perissodactyla*, our topology is concordant with previous analyses in recovering a sister relationship between *Ferae* and *Perissodactyla* (dos Reis et al. 2012; Tarver et al. 2016; Esselstyn et al. 2017; Álvarez-Carretero et al. 2022; Foley et al. 2023). As in previous analyses, our result supports a ‘soft explosive’ model of placental mammal diversification in which diversification among most orders occurred in the Cretaceous and diversification of placental order crown clades occurred in the Paleocene (Meredith et al. 2011; dos Reis et al. 2014; Phillips and Fruciano 2018; Upham et al. 2021; Álvarez-Carretero et al. 2022; Budd and Mann 2023; Foley et al. 2023).

### Resolution of Lepidosaur and Archelosaur Relationships

One classically (Estes et al. 1988; Lee 1993; Gauthier et al. 2012; Hedges 2012; Bever et al. 2015) controversial region of jawed vertebrate phylogeny that has been rather conclusively resolved using genomic data is the phylogeny of living reptile clades. As in previous analyses, our phylogenetic analyses all strongly support the placement of turtles as sister to archosaurs (birds and crocodylians) to form *Archelosauria* (Iwabe et al. 2005; Chiari et al. 2012; Crawford et al. 2012; Hedges 2012; Wang et al. 2013) and the placement of iguanians deep within the living lineages of crown squamates to form a clade with snakes and anguimorphs (*Toxicofera*) (Townsend et al. 2004; Vidal and Hedges 2005; Crawford et al. 2012; Pyron et al. 2013; Zheng and Wiens 2016; Streicher and Wiens 2017; Simões et al. 2018; Burbrink et al. 2020; Singhal et al. 2021; Title et al. 2024). Resolution of these key deep divergences in reptiles is arguably one of the great accomplishments of the phylogenomic program in jawed vertebrates. However, the relationships among the tree major lineages of toxicoferans conflict among our analyses (Figure 2; Extended Data Figure 1; Extended Data Figure 4), as in previous studies, and the resolution of the living sister to snakes therefore remains an open question (Townsend et al. 2004; Vidal and Hedges 2005; Crawford et al. 2012; Pyron et al. 2013; Zheng and Wiens 2016; Streicher and Wiens 2017; Simões et al. 2018; Burbrink et al. 2020; Singhal et al. 2021; Title et al. 2024). Also notable is the latest Permian age of crown *Lepidosauria* estimated in our node-dating analyses (median MRCA age: 263.27 Ma, 95% HPD: 241.13, 285.25 Ma; Figure 1, Extended Data Figure 4, Extended Data Figure 6), which contrasts with the earliest Triassic origin of the lepidosaur crown group found in a number of studies using genomic data (Jones et al. 2013; Burbrink et al. 2020) but is congruent with several time-calibrated phylogenies built using morphological characters (Simões et al. 2018; Brownstein et al. 2022).

### Crown Bird Diversification and Phylogenetic Discordance

The most poorly resolved portion of jawed vertebrate phylogeny in our analyses is undisputably *Neoaves* (Figure 1, Figure S1, Extended Data Figure 4, Extended Data Figure 7). The backbone of *Neoaves* is supported by the lowest average gene and site concordance factors among post-Cretaceous radiations in our analyses (Extended Data Figure 7), making *Neoaves* an unusually difficult problem. Over the past decade, nearly a dozen phylogenomic analyses have failed to provide a clear resolution of the relationships of crown birds (Hackett et al. 2008; Jarvis et al. 2014; Claramunt and Cracraft 2015; Prum et al. 2015; Reddy et al. 2017; Kimball et al. 2019; Kuhl et al. 2021; Berv et al. 2024; Stiller et al. 2024). This problem is attributable to a combination of phenomena that occurred in birds during the Cretaceous-Paleogene transition, including strong selection for certain ecologies and life histories (Berv and Field 2018; Field et al. 2018; Berv et al. 2024), a rapid episode of body size reduction (Berv and Field 2018) suppressed recombination across long segments of the avian genome (Mirarab et al. 2024), and rapid successive divergences associated with the initial adaptive radiation of *Neoaves* (Figure 1, Figure 2, Figure 3, Extended Data Figure 4, Extended Data Figure 7, Extended Data Figure 8)(Prum et al. 2015; Stiller et al. 2024). Our phylogeny supports the hypothesis that, as with placental mammals, a huge proportion of orthologous genes are needed to resolve avian phylogeny (Stiller et al. 2024); this arguably requires focusing on neoavians to the exclusion of other jawed vertebrates. As with other analyses, our results suggest that only three or four major bird clades survived the K-Pg: *Paleognathae*, *Galloanserae*, and one or two lineages of *Neoaves* (*Mirandornithes*, all others)(Jarvis et al. 2014; Prum et al. 2015; Berv et al. 2024; Stiller et al. 2024). In our analyses, the origin of *Aves* is estimated to lie in the Late Cretaceous, 97.42 Ma (95% HPD: 83.15, 132.22 Ma), and the origin of *Neognathae* (*Neoaves* + *Galloanserae*) lies within the Campanian, 78.88 Ma (95% HPD: 73.32, 84.54 Ma). The relationships of most major neoavian clades remain unresolved in our analysis, and we emphasize that the phylogeny of *Aves* presented in this manuscript is less robust than previously inferred phylogenies that have focused on this clade. Odd relationships, such as the paraphyly of lineages in *Strisores* and *Coraciimorphae*, inferred in our phylogenies are probably caused by the absence of informative sequences for avian phylogeny included in our dataset, an inference that is supported by the low gene and site concordance factors for nodes across *Neoaves*.

### Early-Diverging Actinopterygian Phylogeny

As in previous studies, we find that four major lineages comprise the earliest divergences among ray-finned fishes (*Actinopterygii*): bichirs and Reedfish (*Polypteridae*), sturgeons and paddlefishes (*Acipenseriformes*), gars and bowfins (*Holostei*), and all other ray-finned fishes (*Teleostei*)(Grande 2010; Near et al. 2012; Betancur-R et al. 2013; Braasch et al. 2016; Hughes et al. 2018; Du et al. 2020; Bi et al. 2021a; Thompson et al. 2021; Mallik et al. 2025). Both analyses in which the multispecies coalescent or concatenation approaches are employed resolve *Polypteridae*, *Acipenseriformes*, and *Holostei* as the successive sister lineages to *Teleostei*; however, some ASTRAL-III trees unite *Polypteridae* and *Acipenseriformes* to the exclusion of other actinopterygians (Extended Data Figure 1). Uncertainty over the earliest divergence in ray-finned fishes has mainly centered around the conflict between morphological and molecular data, in which the latter unites *Polypteridae* and *Acipenseriformes* (Argyriou et al. 2018; Argyriou et al.) or find them in a polytomy including fossils of early actinopterygians (Giles et al. 2017).One previous study of actinopterygian genomes did find only moderate (bootstrap < 100) support for *Actinopteri*, the clade uniting *Acipenseriformes*, *Holostei*, and *Teleostei* (Bi et al. 2021a); like ours, this study sampled several non-ray-finned fish outgroups (Bi et al. 2021b). The ages of these successive divergences are even less clear, as excluding representatives two of these lineages (*Acipenseriformes*, *Holostei*) notable for their exceptionally slow rates of genome sequence evolution (Braasch et al. 2016; Takezaki 2018; Brownstein, MacGuigan, et al. 2024) shift the crown ages of *Actinopterygii* and *Teleostei* forward in time by up to 50 million years (Figure 5; Extended Data Figure 9, Extended Data Figure 10). The exceptionally rapid rates of molecular evolution in teleost fishes (Brownstein, MacGuigan, et al. 2024), which appear to slip towards the root of the ray-finned fish crown clade in our analyses (Extended Data Figure 9, Extended Data Figure 10), probably also play a role in uncertainty surrounding the ages of these divergences, which conflict with the much younger ages of *Actinopterygii* and *Teleostei* implied by the fossil record (Hurley et al. 2006; Near et al. 2012; Friedman 2022). Our results suggest that, in the absence of additional fossil discoveries, the ages of these clades, which contain nearly half of living vertebrate diversity, are unknowable owing to biological variation in the tempo of genome evolution. In contrast, our analyses contribute to consilience around the earliest divergence in teleosts. Recently, some analyses of whole genomes using traditional phylogenetic methodology (Bian et al. 2016; Hao et al. 2020; Takezaki 2021) and synteny analysis (Parey et al. 2023) resolved a clade containing mooneyes and bonytongues (*Osteoglossomorpha*) and eels, tarpons, bonefishes, and halosaurs (*Elopomorpha*) that forms the sister clade to all other teleosts (*Clupeocephala*). However, studies of legacy nuclear and mitochondrial markers (Near et al. 2012; Betancur-R et al. 2013) and at least some analyses of genome-wide sequence data(Hughes et al. 2018) resolve these lineages as successive sister clades to *Clupeocephala.* Our analyses, which use the same exon set as Hughes et al.,(Hughes et al. 2018) unambiguously resolve a sister lineage relationship between *Elopomorpha* and *Osteoglossomorpha*, a result also obtained in another recent analysis of these loci (Hughes et al. 2021). Thus, across genomic datasets, there is little to no conflict over the split at the base of teleosts.

### Ostariophysan Relationships and the Recognition of Cithariniformes

Another problem in ray-finned fish phylogenetics that our results contribute towards resolving are the relationships of the major freshwater fish clades that form the clade *Ostariophysi*.

Analyses of genomic markers, including ultraconserved elements (Chakrabarty et al. 2017; Melo et al. 2022) and exons (Hughes et al. 2021), find the traditional *Characiformes* to be paraphyletic, with the *Cithariniformes* sensu Near and Thacker (Near and Thacker 2024) forming the sister lineage to a clade containing catfishes (*Siluriformes*) and *Characiformes* sensu lato. However, some studies of exon loci regard the traditional placement of *Cithariniformes* as the sister to other *Characiformes* to be more likely (Arcila et al. 2017; Hughes et al. 2018; Betancur-R. et al. 2019). Our analyses unambiguously reject the monophyly of traditional *Characiformes* and place *Cithariniformes* sister to a clade containing *Siluriformes*, *Characiformes* sensu lato, and *Gymnotiformes*, which contains the Neotropical electric fishes (Figure 1, Figure 2). Although the clade containing *Siluriformes* and *Characiformes* sensu lato to the exclusion of other otophysans is always supported by high bootstrap, coalescent, gene, and site concordance factors, the placement of *Gymnotiformes* as the sister lineage to this clade is only moderately supported (coalescent value < 1) in some analyses (Figure 2). As has the divergence at the base of teleosts (Parey et al. 2023), resolving the relationships of order-level clades in *Ostariophysi* would probably benefit from an analysis of genome structural conservation.

### The Black Box of Euteleost Relationships

A problem in vertebrate phylogeny whose lack of resolution has decidedly gone rather unnoticed is the relationships among the major lineages of euteleosts; these differ across essentially every analysis of ray-finned fish phylogeny (Near et al. 2012; Betancur-R et al. 2013; Hughes et al. 2018; Straube et al. 2018; Hughes et al. 2021). Although we find support for several key clades, including the *Stomiati*, containing the dragonfishes and hatchetfishes (*Stomiiformes*) and smelts (*Osmeriformes*), and the *Salmoniformes*, containing the salmons, charrs, and trouts (*Salmonidae*) and mudminnows and pikes (*Esocidae*), we fail to consistently resolve the positions of marine smelts and barreleyes (*Osmeriformes*) or the galaxiids (*Galaxiidae*) among euteleosts (Figure 2; Extended Data Figure 1). Given that the placement of these lineages might have major implications for reconstructing the ancestral habitat (deep marine, shallow marine, or freshwater)(Miller et al. 2022) of ray-finned fishes, we highlight this region of the tree as ripe for investigation.

### Phylogenetic Resolution of Acanthomorpha

Our analyses provide a degree of consilience around the relationships of spiny-rayed fishes (*Acanthomorpha*). Resolution of this region of ray-finned fish phylogeny is recognized as an important milestone of phylogenomics (Dornburg and Near 2021; Near and Thacker 2024). As in several recent studies of genome-wide markers (Alfaro et al. 2018; Hughes et al. 2018; Hughes et al. 2021; Ghezelayagh et al. 2022), we find that four lineages form the successive four or five divergences among acanthomorphs: beardfishes (*Polymixia*), opahs and oarfishes (*Lampriformes*), codfishes, granadiers, Tube-Eye, oreos, and John Dories (*Zeiogadaria*), and Pirate Perch, trout-perches, and North American cavefishes (*Percopsiformes*)(Figure 1). The positions of *Lampriformes*, *Percopsiformes,* and *Polymixia* change depending on analytical method employed and exon sample used in our analyses (see Figure 2), underscoring the uncertain positions of these lineages (Alfaro et al. 2018; Hughes et al. 2018; Ghezelayagh et al. 2022). Next to diverge are an oddball clade containing flashlightfishes, porcupinefishes, roughies, and fangtooths (*Trachichthyiformes*), the squirrelfishes and soldierfishes (*Holocentridae)*, and the alfonsinos, whalefishes, gibberfishes, and bigscales (*Berycioidei*); the latter two form a clade (*Beryciformes*) in all of our phylogenies produced using concatenated exons and in some of our phylogenies produced using the multispecies coalescent in ASTRAL-III. Although phylogenies that we generated using concatenated exon sequences are consistent with the hypothesis that *Trachichthyiformes* is the sister lineage to *Beryciformes* and other acanthomorphs (*Percomorpha*), as supported by analyses of ultraconserved elements (Ghezelayagh et al. 2022; Brownstein et al. 2025), anchored hybrid enrichment loci (Dornburg et al. 2017), and exons (Hughes et al. 2018; Musilova et al. 2019; Hughes et al. 2021) (though see (Hughes et al. 2018)), ASTRAL-III phylogenies that we generated find that these three lineages form a clade sister to *Percomorpha*. We fail to recover *Holocentridae* as the sister lineage to *Percomorpha*, in contrast to a previous study of ray-finned fish phylogeny using the same exon loci (Hughes et al. 2018). We consistently find that cusk eels and brotulas (*Ophidiiformes*) and toadfishes *(Batrachoididae*) are the first two lineages of percomorphs to diverge. The relationships of the remaining percomorph lineages break down in the ASTRAL-III topologies, owing largely to the low level of concordance among gene trees (Figure 2). However, the phylogenies built from the maximum likelihood analyses of the concatenated exon sequence set unite seahorses, trumpetfishes, goatfishes, gurnards, and searobins (*Syngnathiformes*) and tunas, mackerels, swallowers, pomfrets, and driftfishes (*Scombriformes*) in a clade that forms the third major divergence in *Percomorpha* (Figure 1). The remaining order-level clades of percomorphs are organized into two major lineages. One, *Eupercaria*, comprises the *Perciformes* (sea basses and basslets, groupers, Antarctic icefishes, darters, sculpins, scorpionfishes, rockfishes, snailfishes, eelpouts, sticklebacks, searobins, and relatives), *Gerreidae* (mojarras), *Labriformes* (wrasses, parrotfishes, stargazers, sandlances, southern sandfishes, sandperches, and torrentfishes), *Centrarchiformes* (sunfishes, black basses, temperate perches, flagtails, knifejaws, kelpfishes, morwongs, and relatives), and *Acanthuriformes* (butterflyfishes, surgeonfishes, pufferfishes, triggerfishes, Mola mola, spadefishes, anglerfishes, drums, rabbitfishes, ponyfishes, Moorish Idol, angelfishes, snappers, grunts, and relatives) (Hughes et al. 2018; Ghezelayagh et al. 2022; Near and Thacker 2024). As in previous studies, *Perciformes* is the sister lineage to the rest of these clades and the position of *Gerreidae* is highly unstable (Alfaro et al. 2018; Hughes et al. 2018; Musilova et al. 2019; Hughes et al. 2021; Ghezelayagh et al. 2022). The other consists of four order-level clades: *Carangiformes* (jacks, trevallies, swordfishes, remoras, Mahi Mahi, flatfishes, flounders, soles, archerfishes, threadfishes, snooks, barracudas, and giant perches), *Synbranchiformes* (swamp eels, betta fishes, and climbing perches), *Atheriniformes* (New World silversides, flying fishes, halfbeaks, ricefishes, pupfishes, killifishes, and relatives), and *Blenniiformes* (blennies, cichlids, damselfishes and anemonefishes, mullets, surfperches, and relatives) (Hughes et al. 2018; Ghezelayagh et al. 2022; Near and Thacker 2024). As in previous studies, we find a Jurassic age for the origin of *Acanthomorpha* and its initial divergences (Figure 1, Extended Data Figure 5), followed by the diversification of most percomorph orders and their constituent lineages throughout the Cretaceous and into the Paleocene (Alfaro et al. 2018; Hughes et al. 2018; Musilova et al. 2019; Ghezelayagh et al. 2022). Along with birds and mammals, acanthomorph fishes show a high degree of phylogenetic discordance associated with diversification occurring around the Cretaceous-Paleogene Mass Extinction (Figure 3, Figure 4, Extended Data Figure 7).

## IV. Supplementary Tables

**Table S1.**
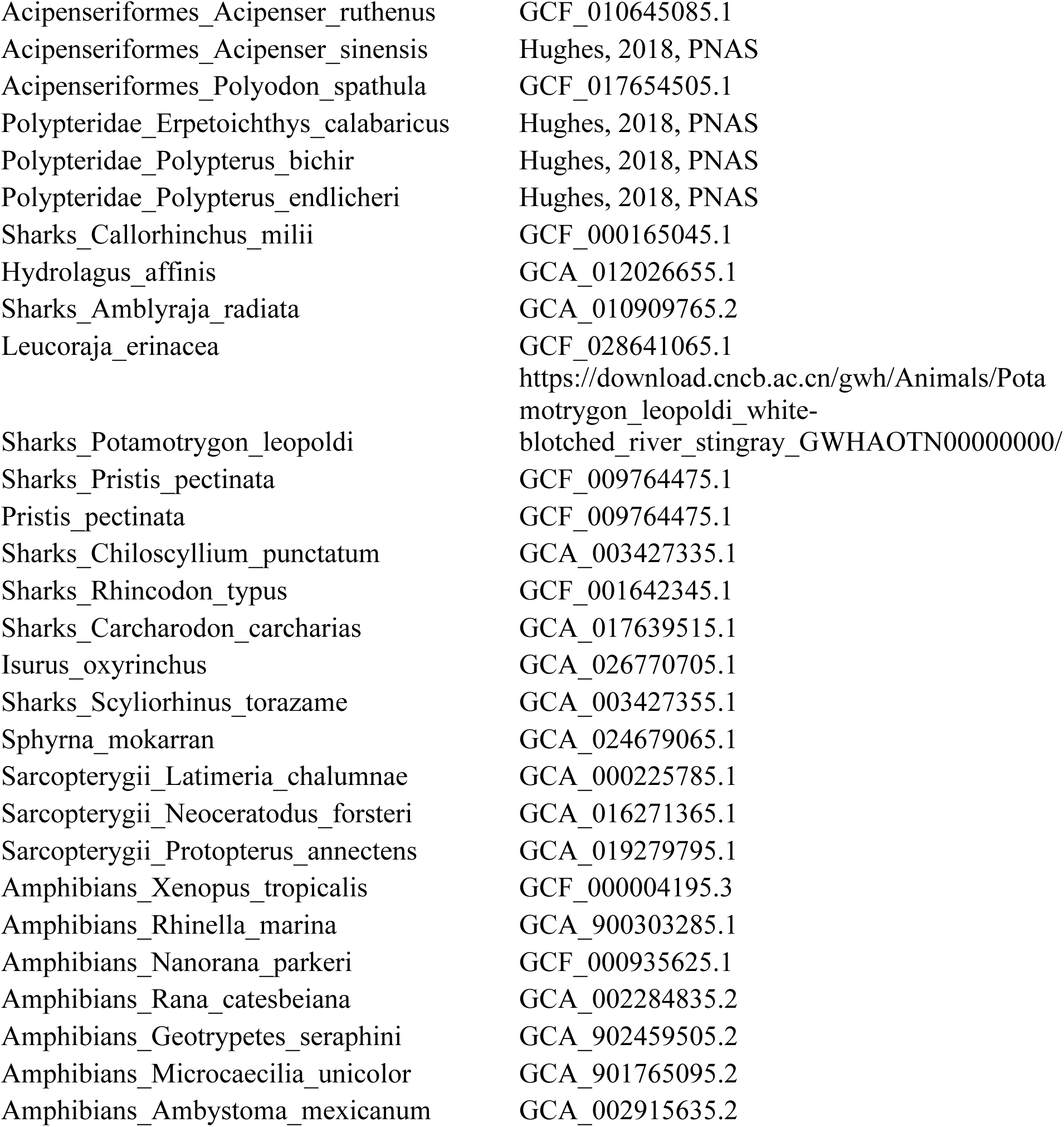
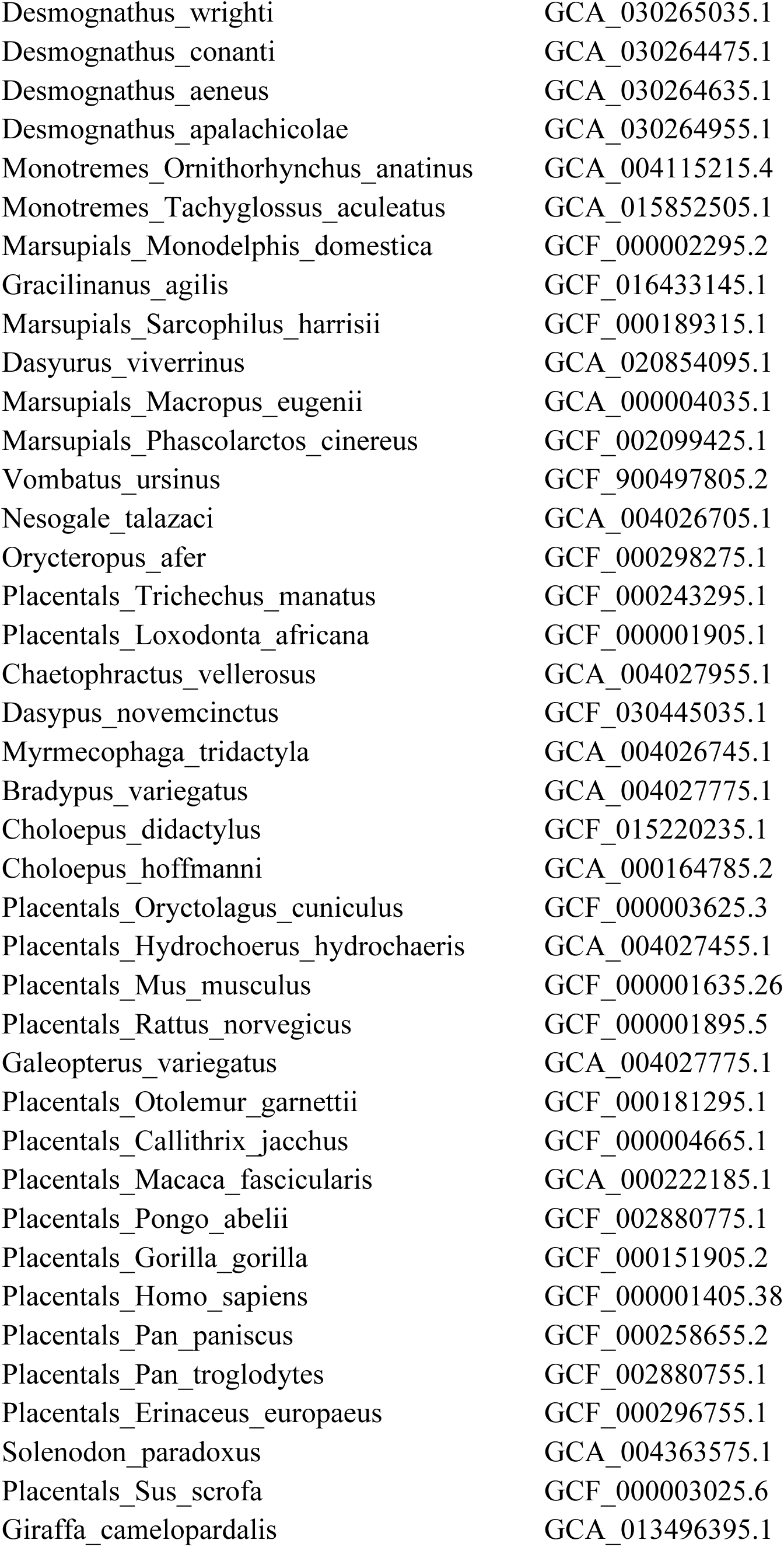
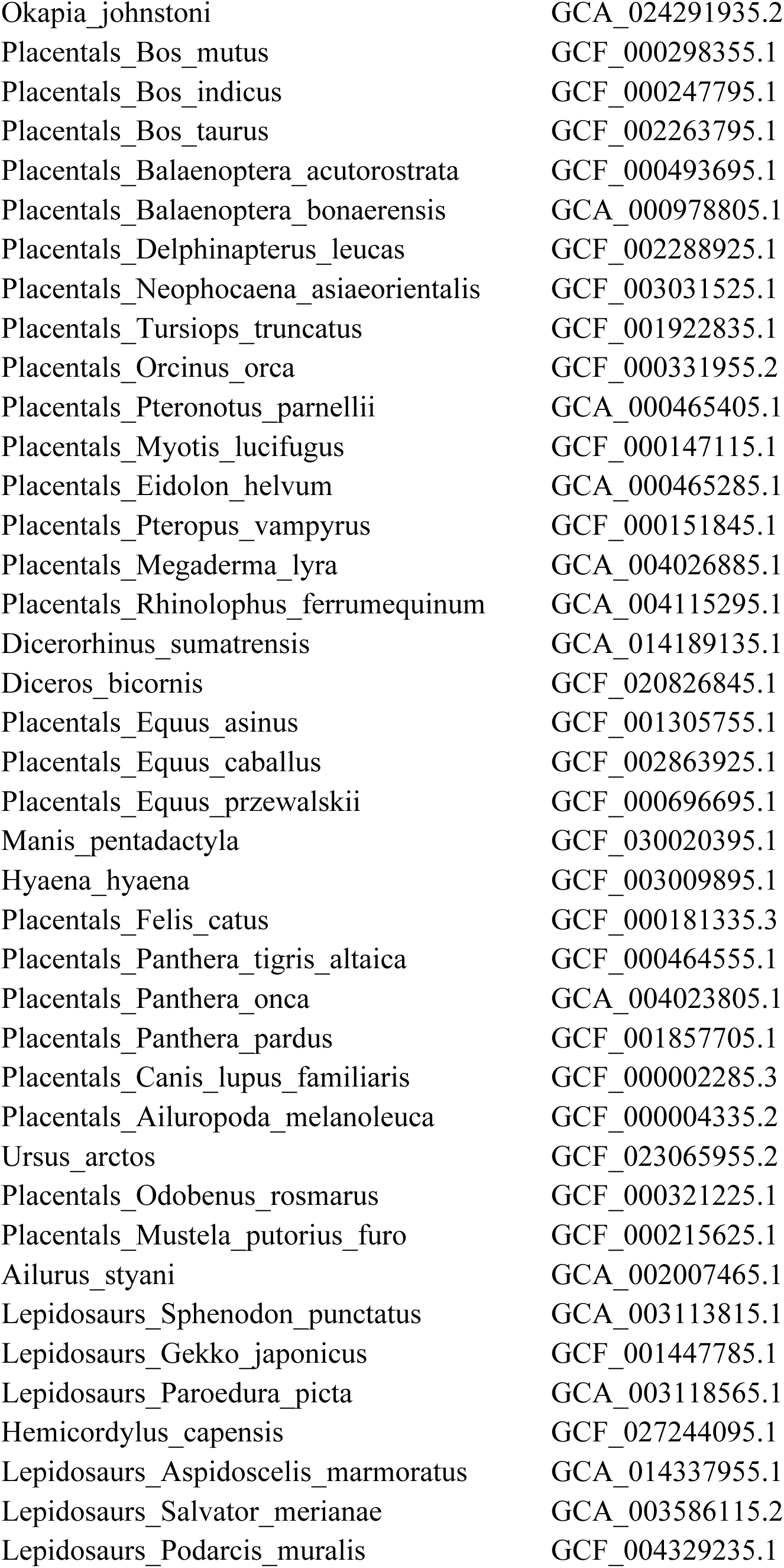
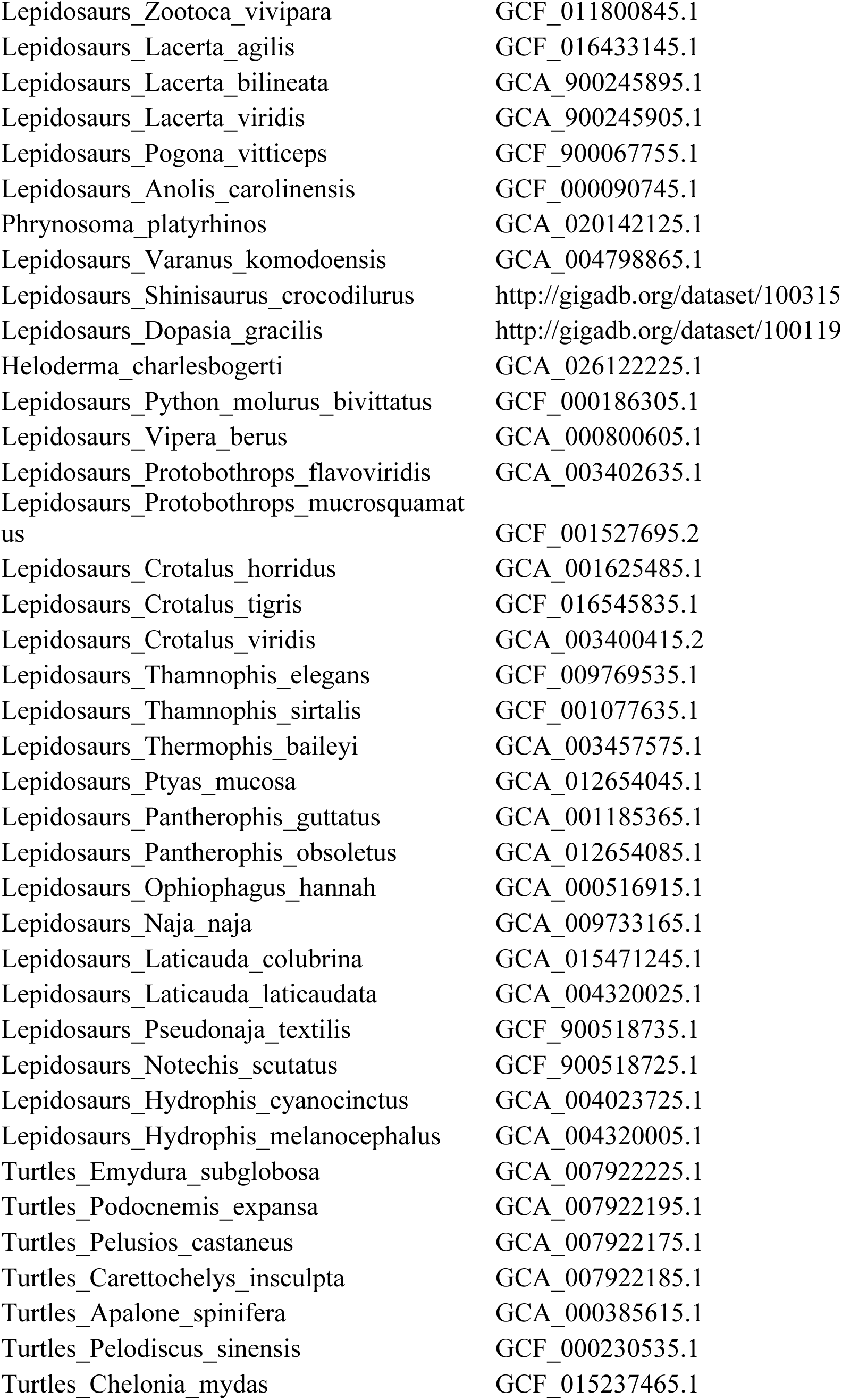
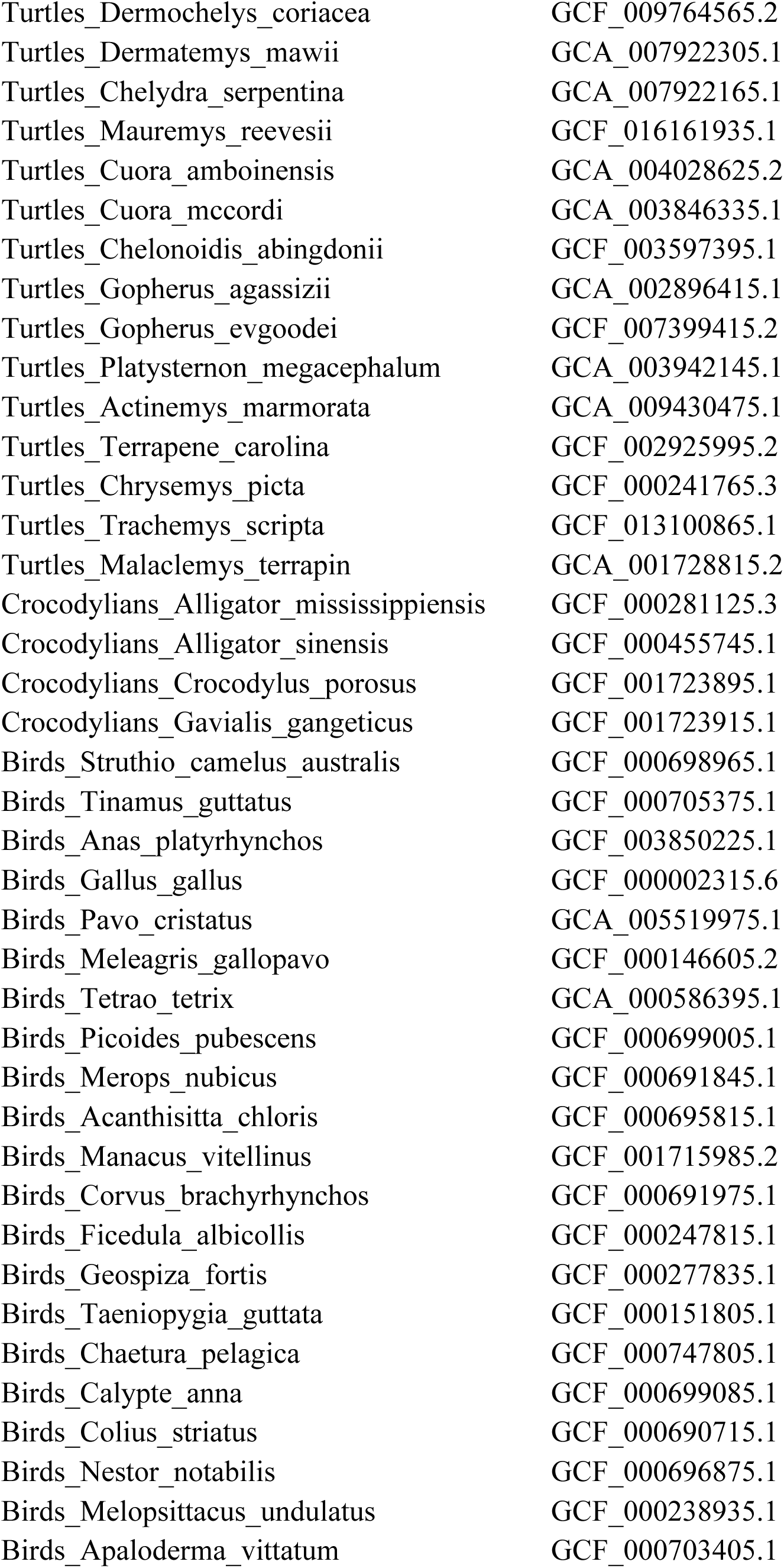
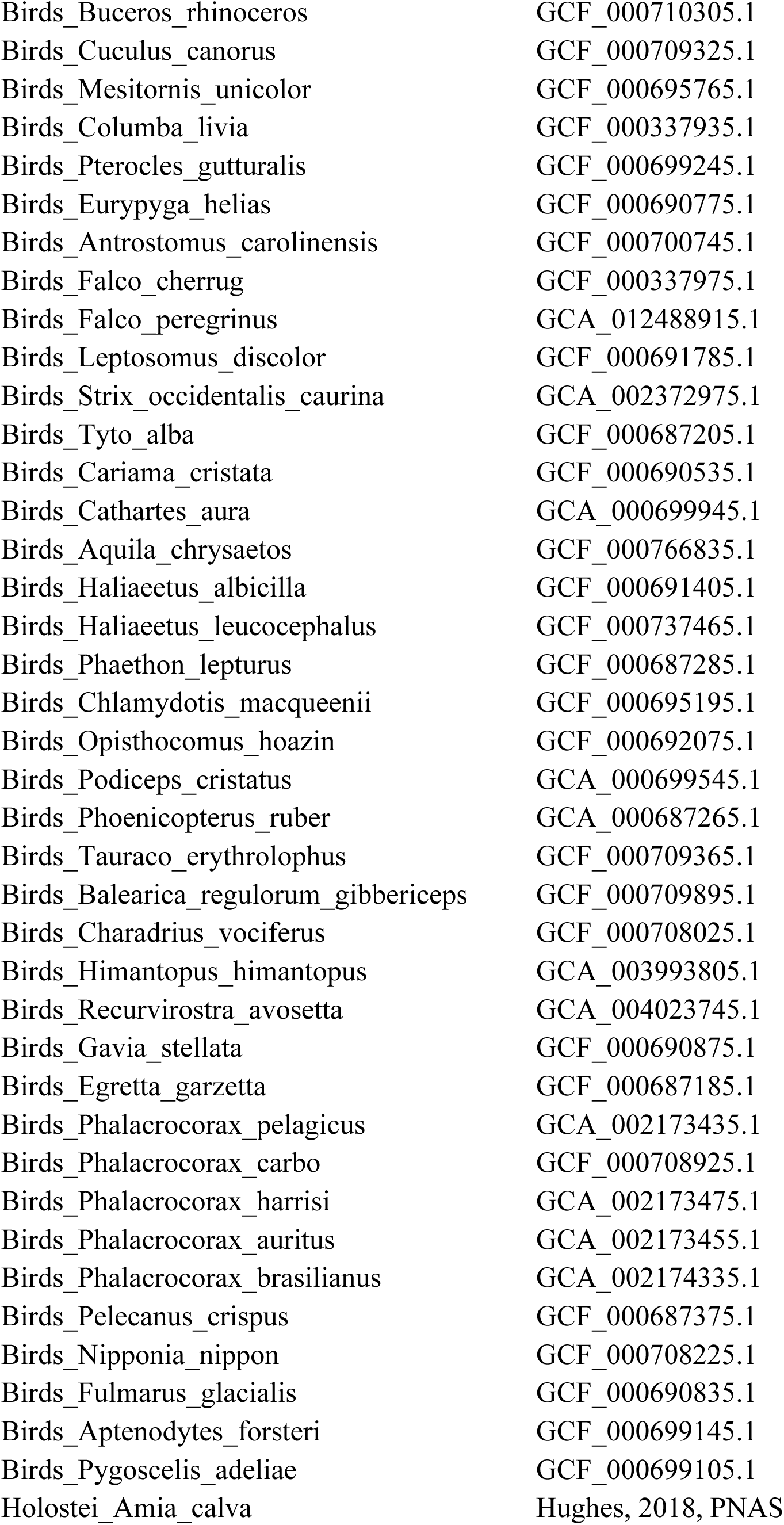
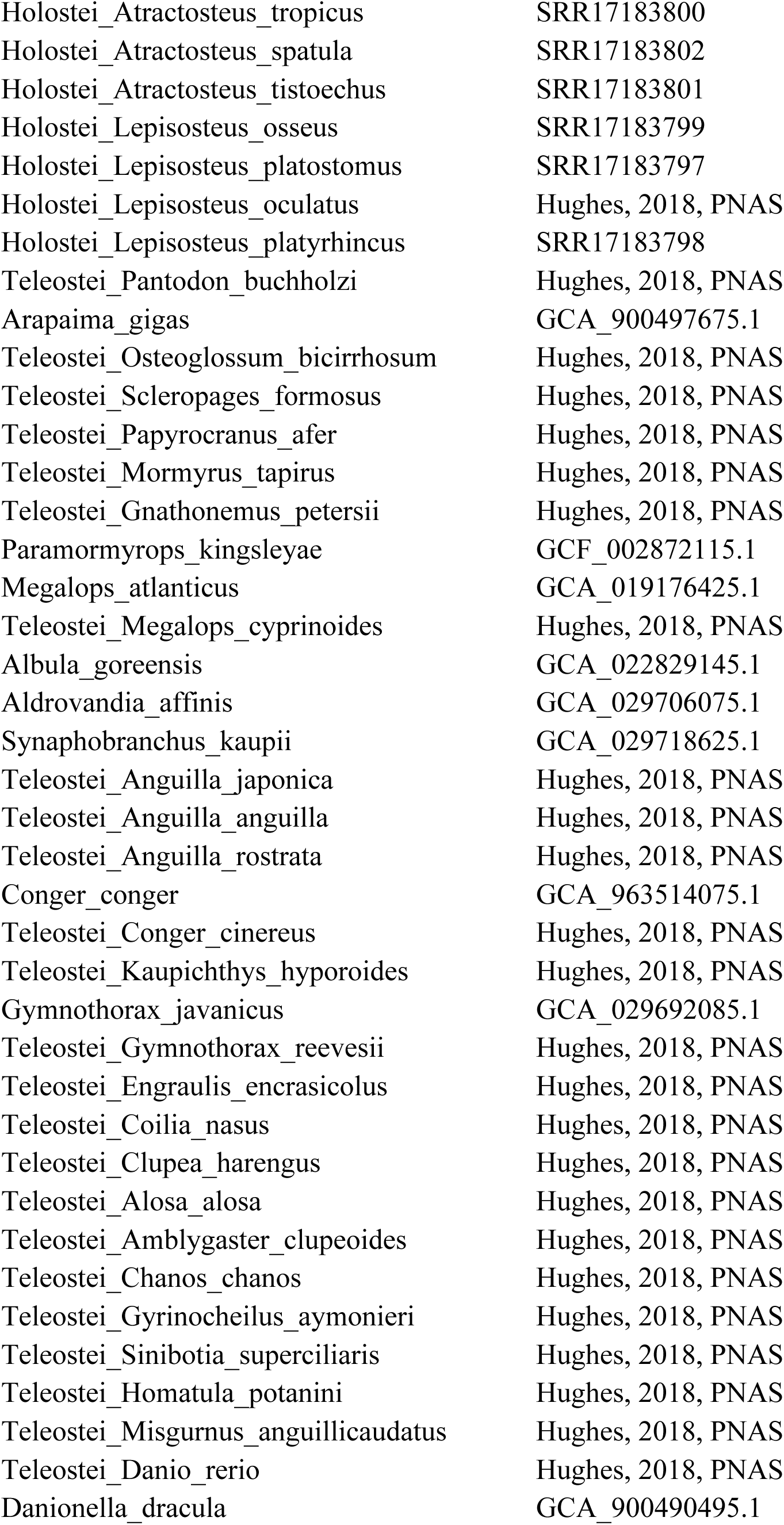
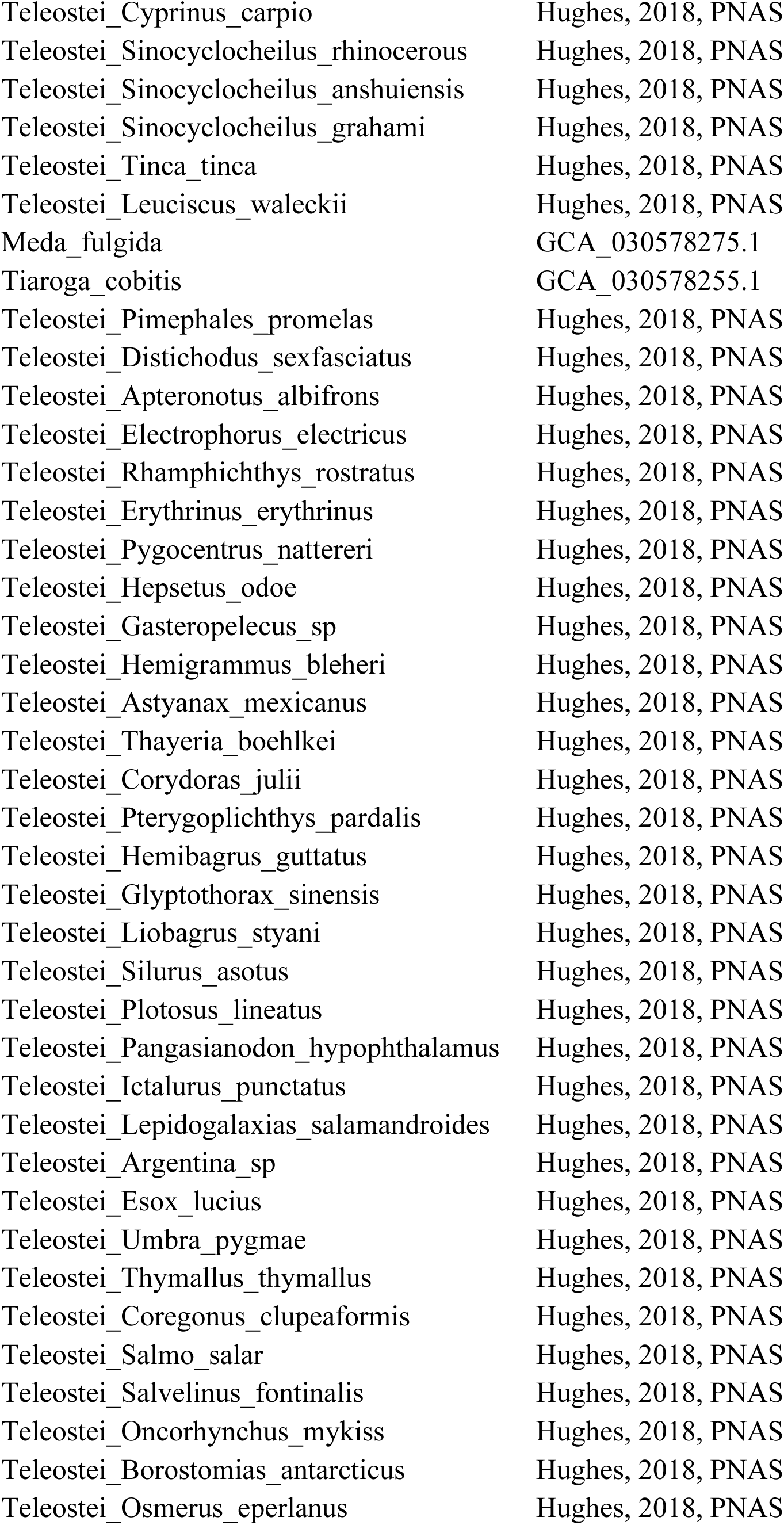
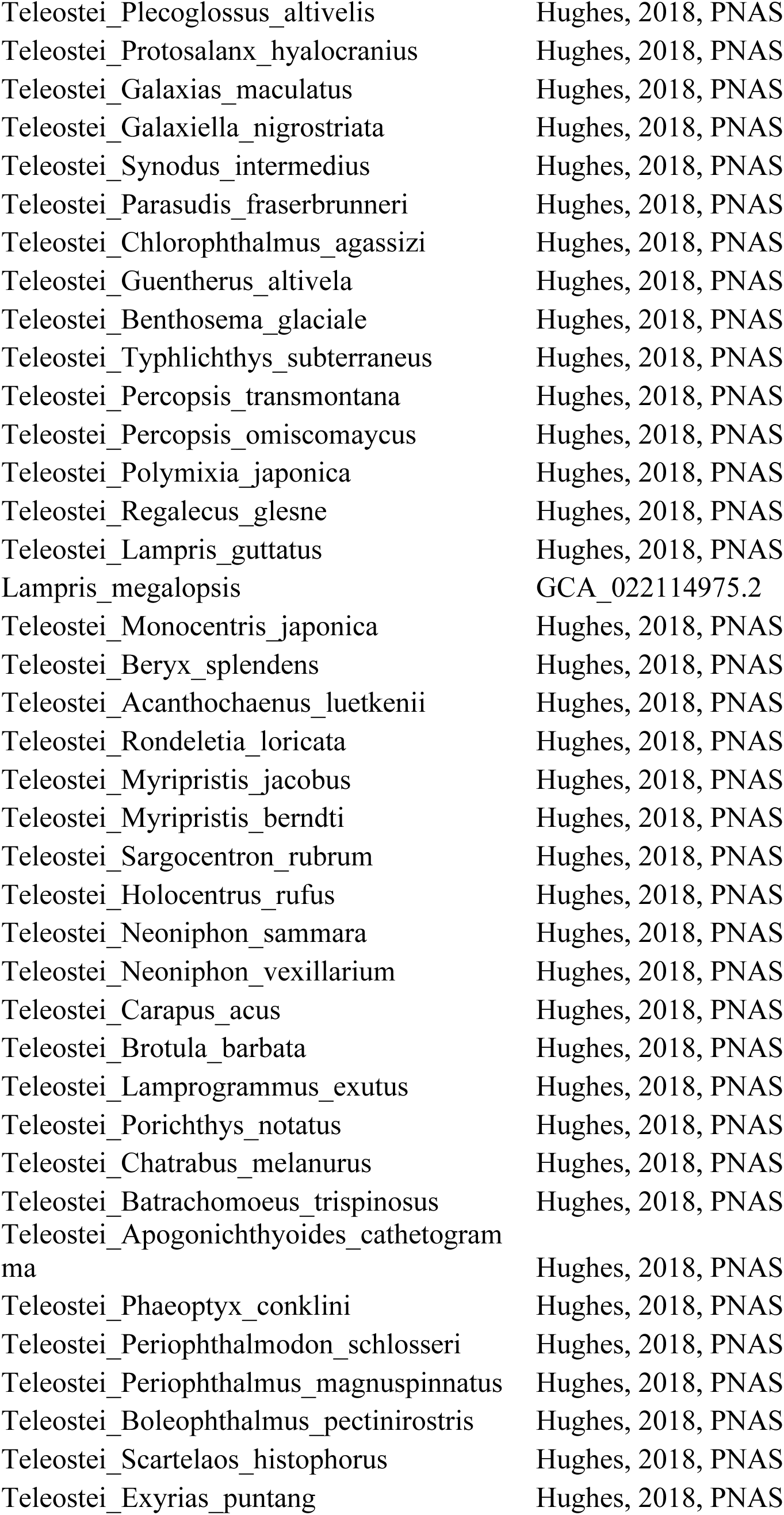
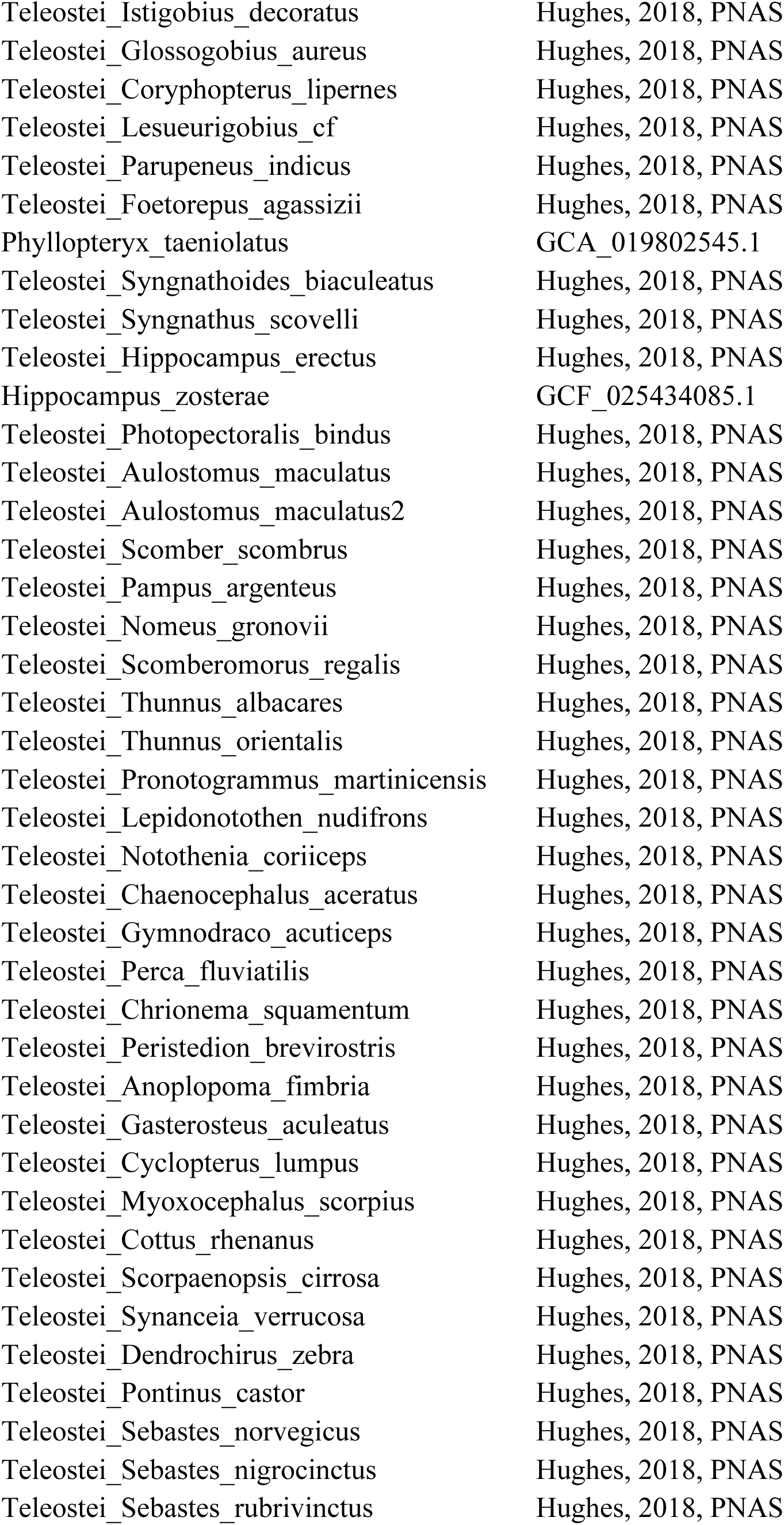
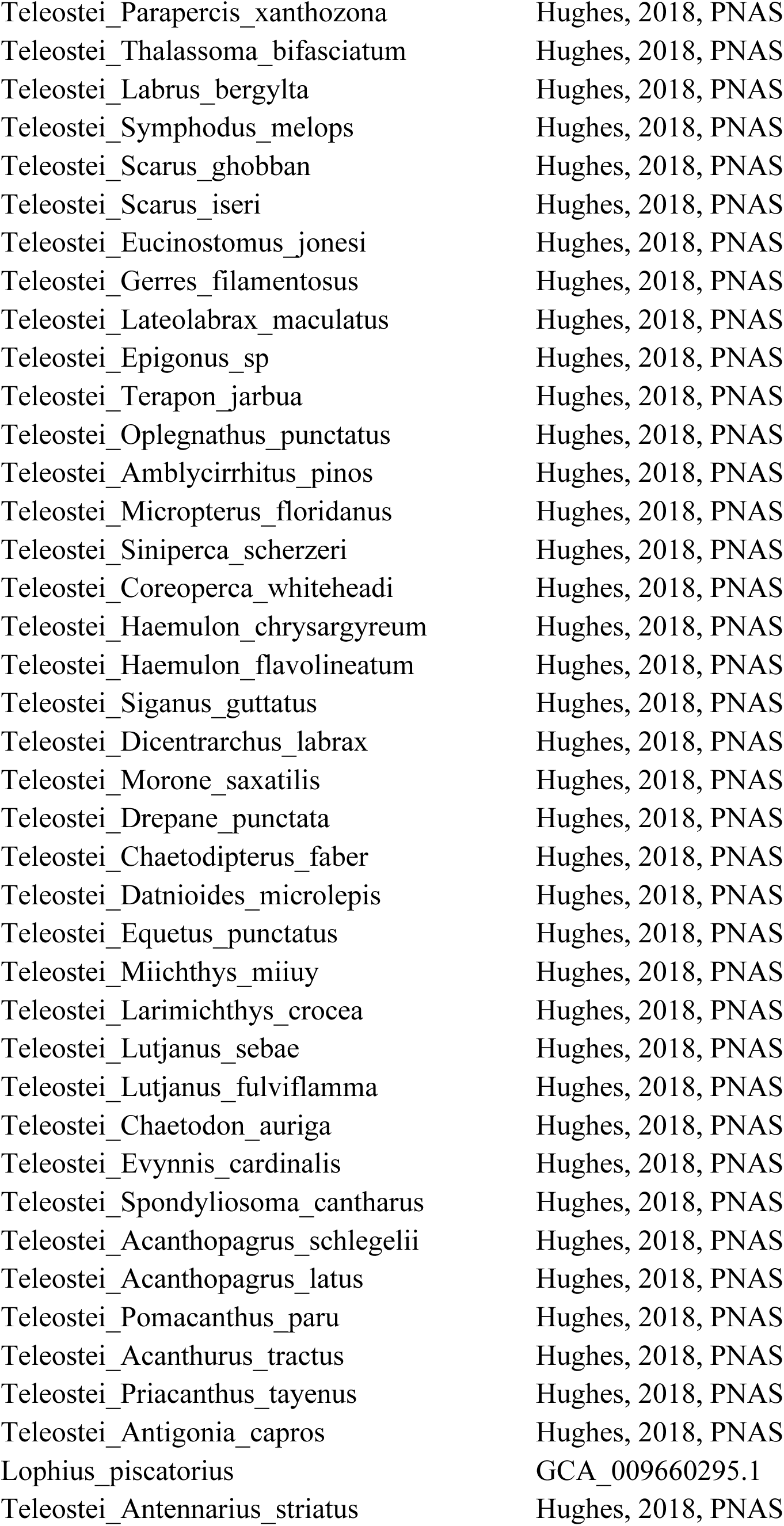
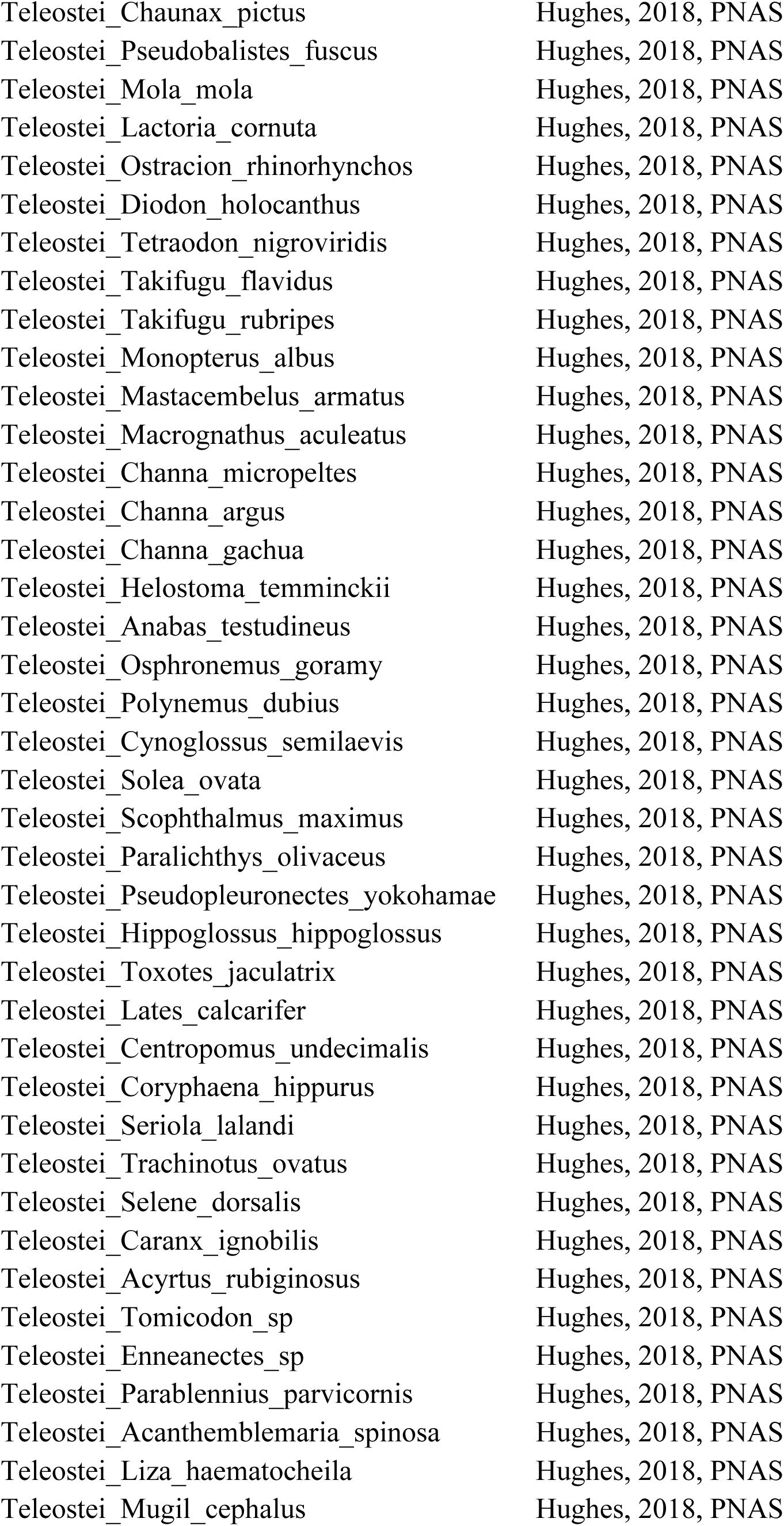
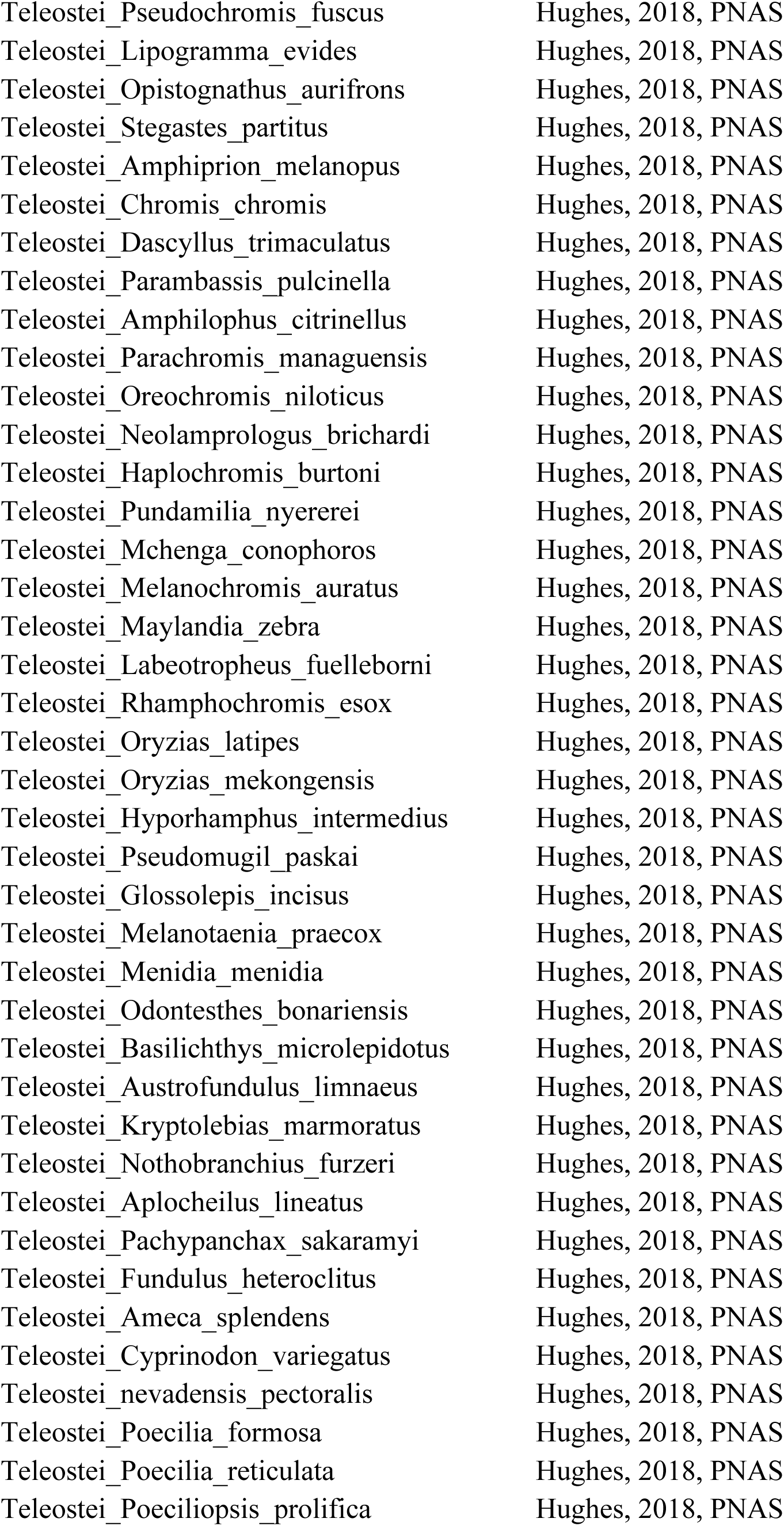
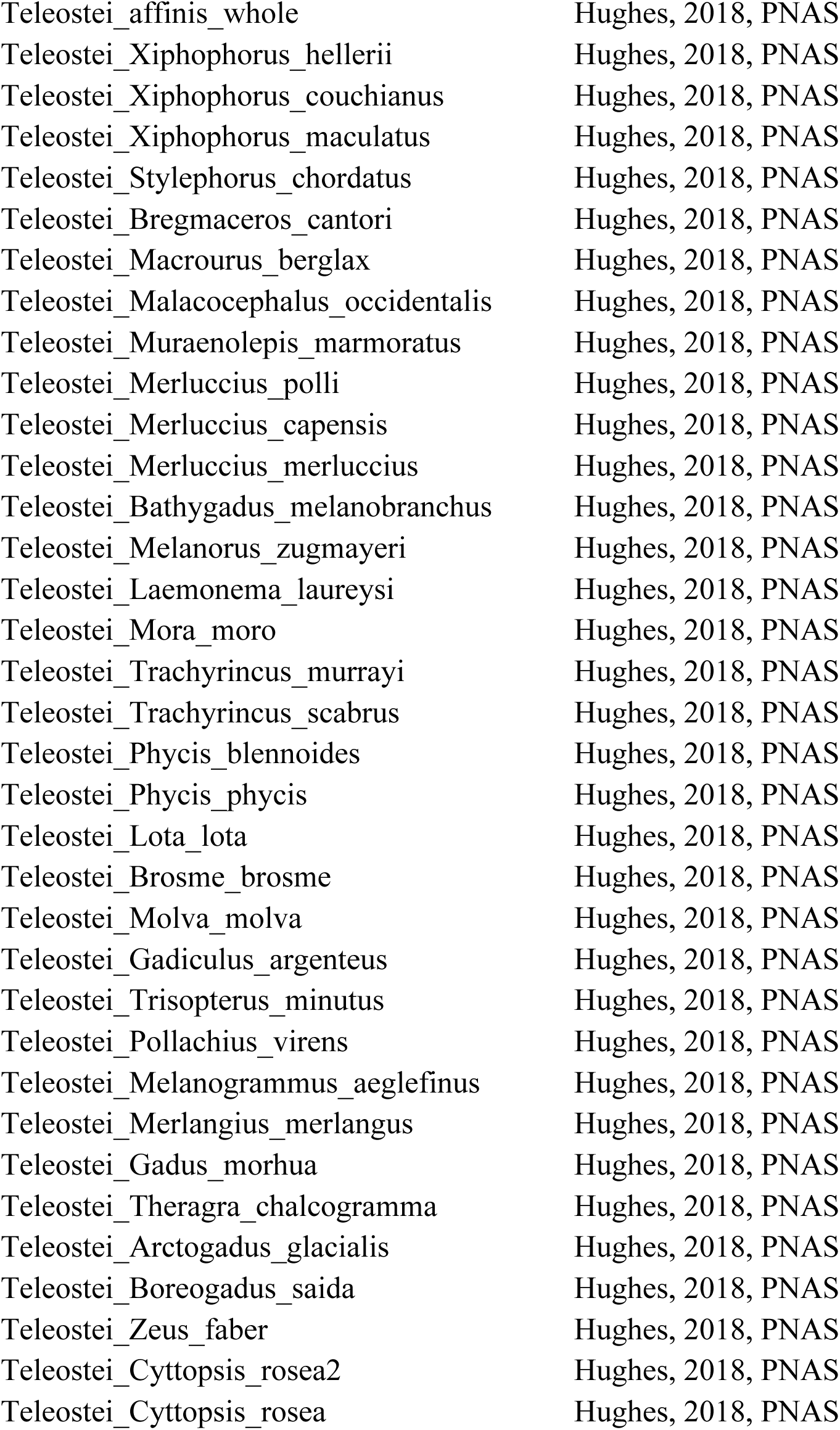
Genome Resources.

**Table S2.**
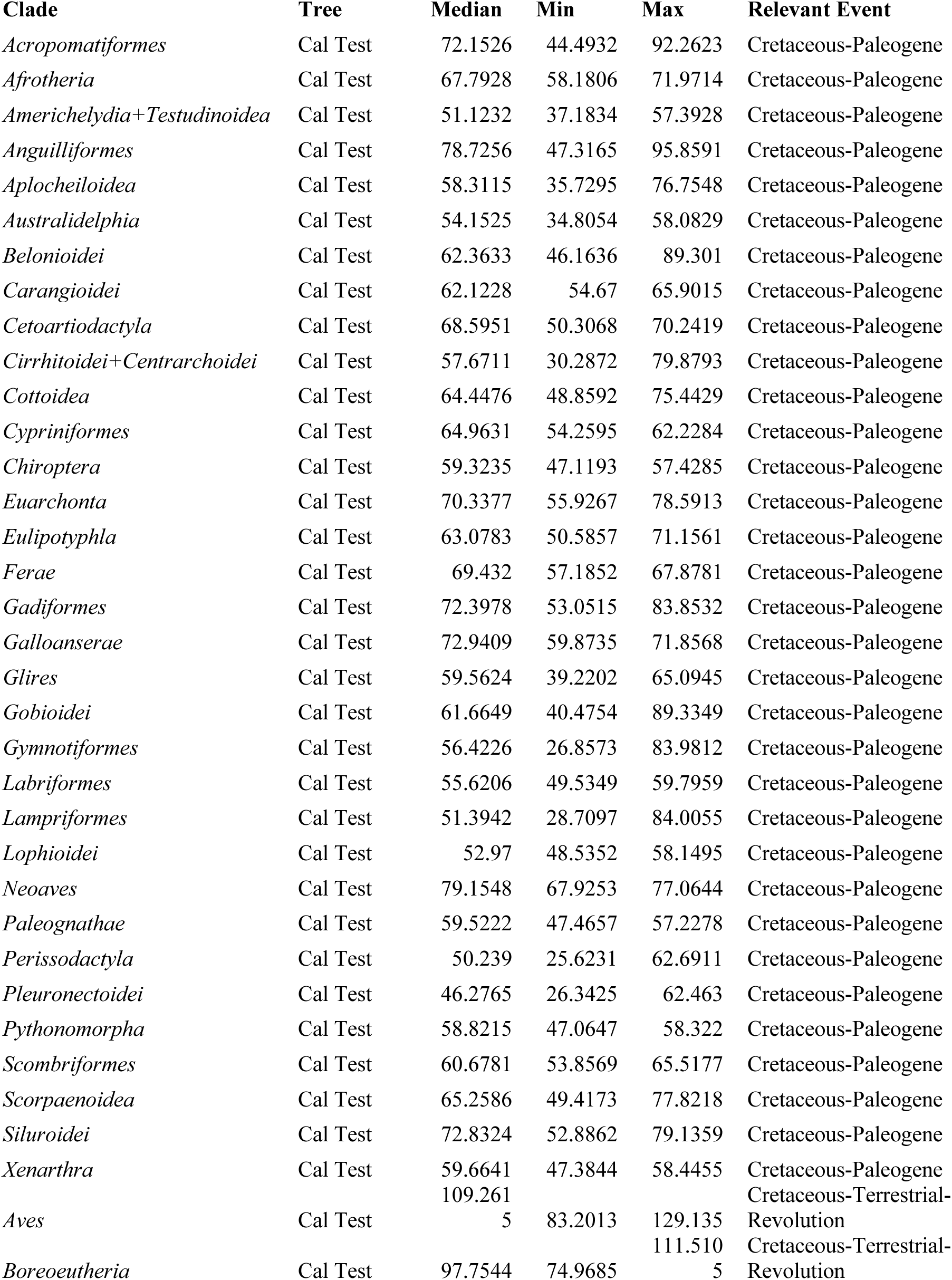
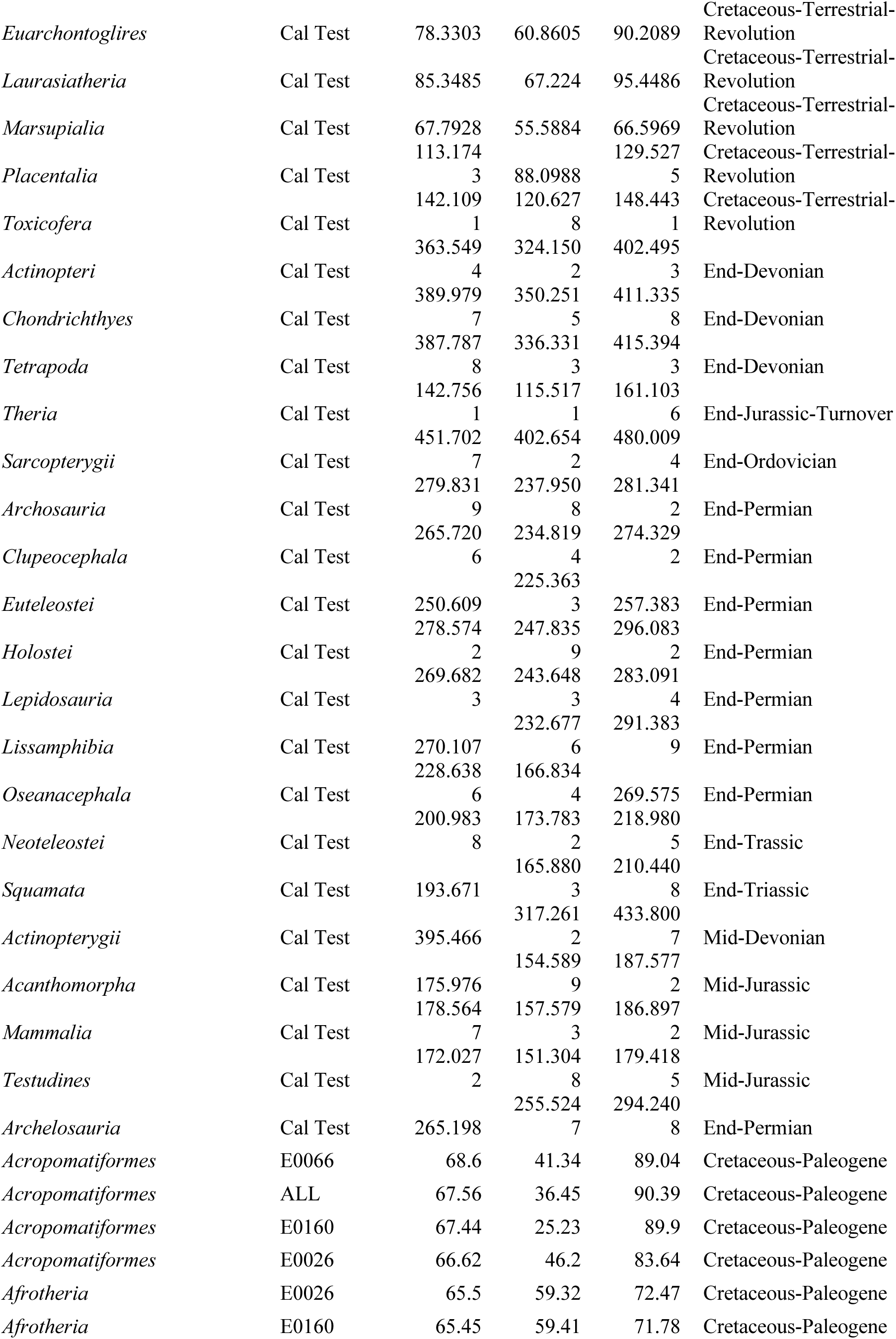
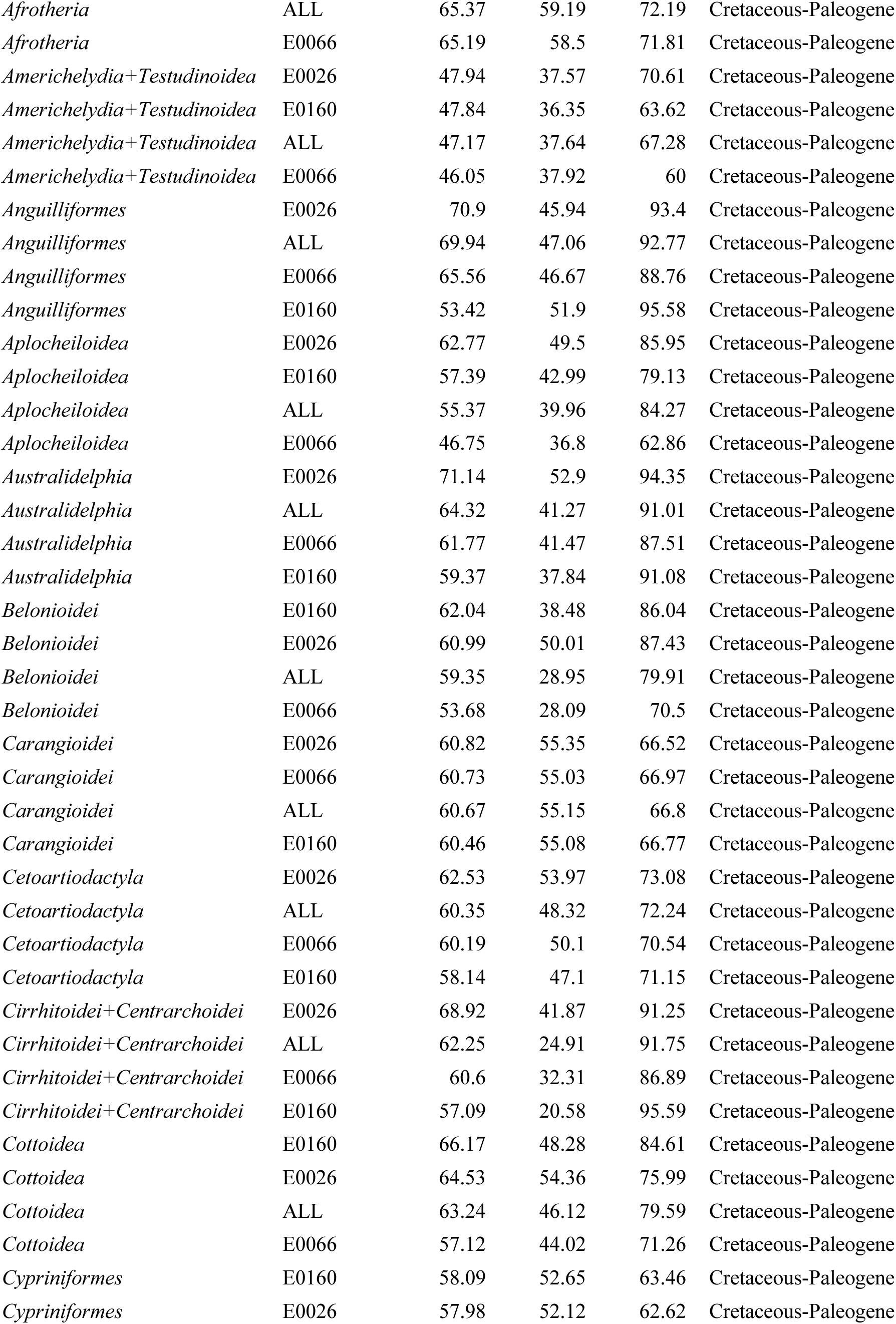
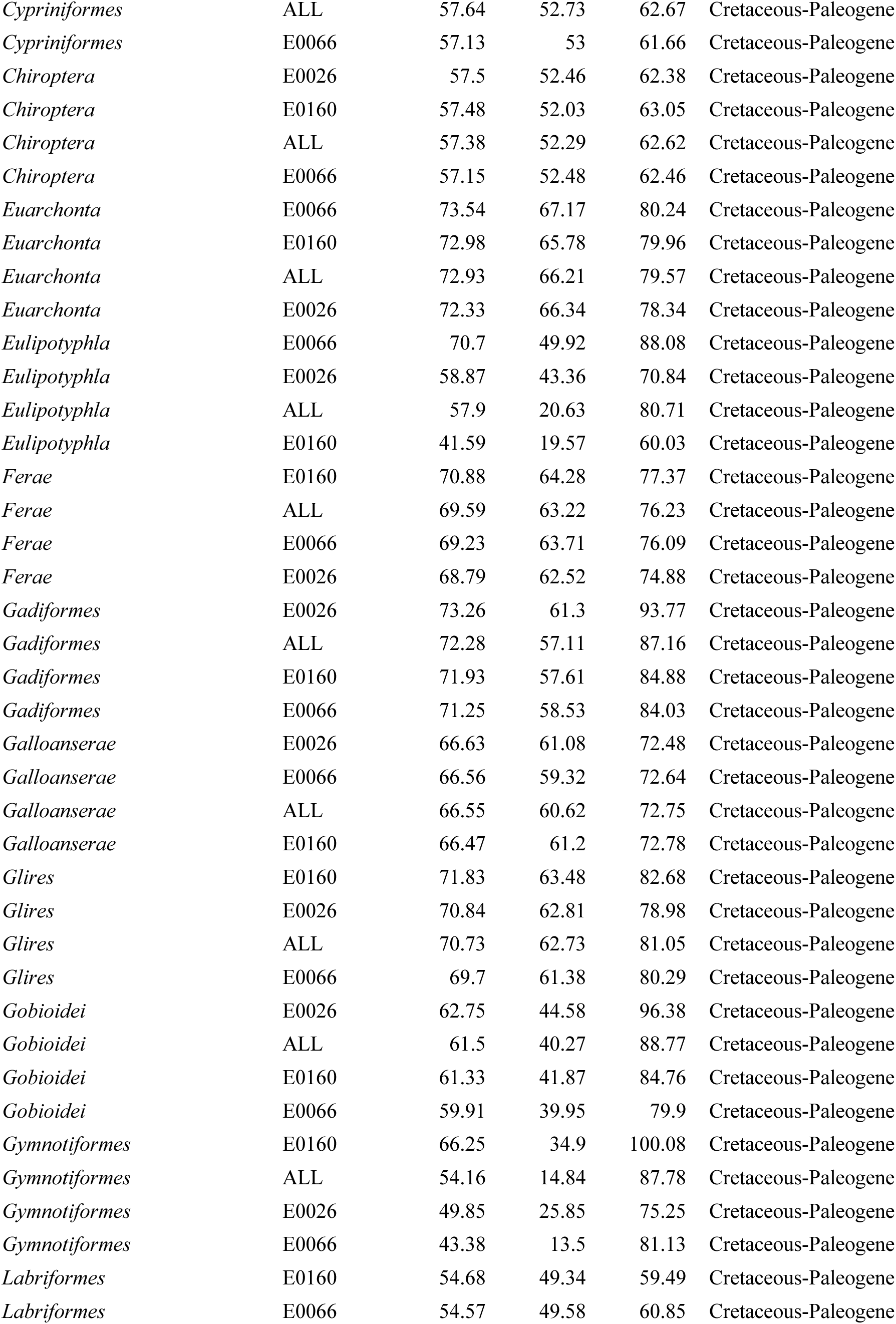
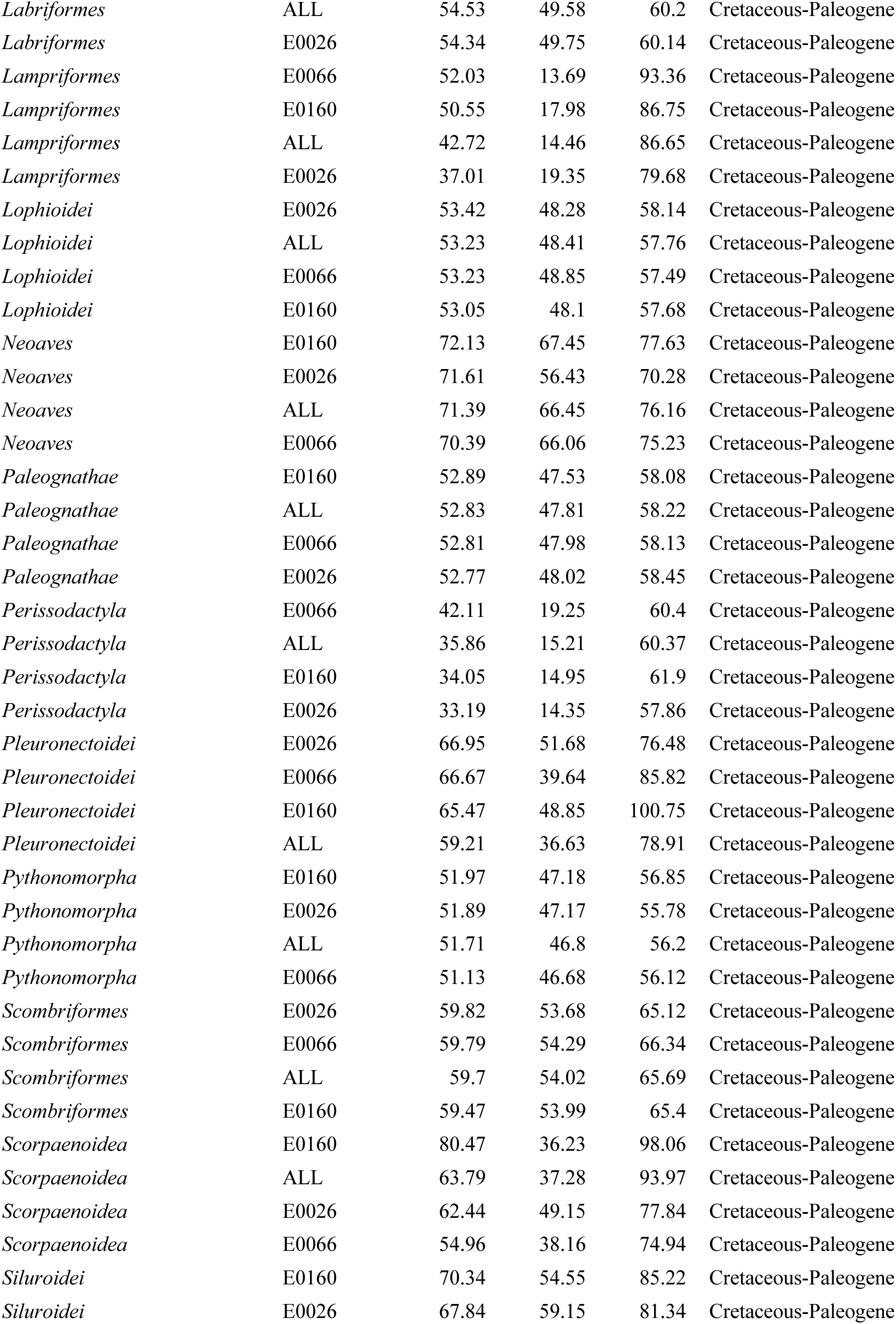
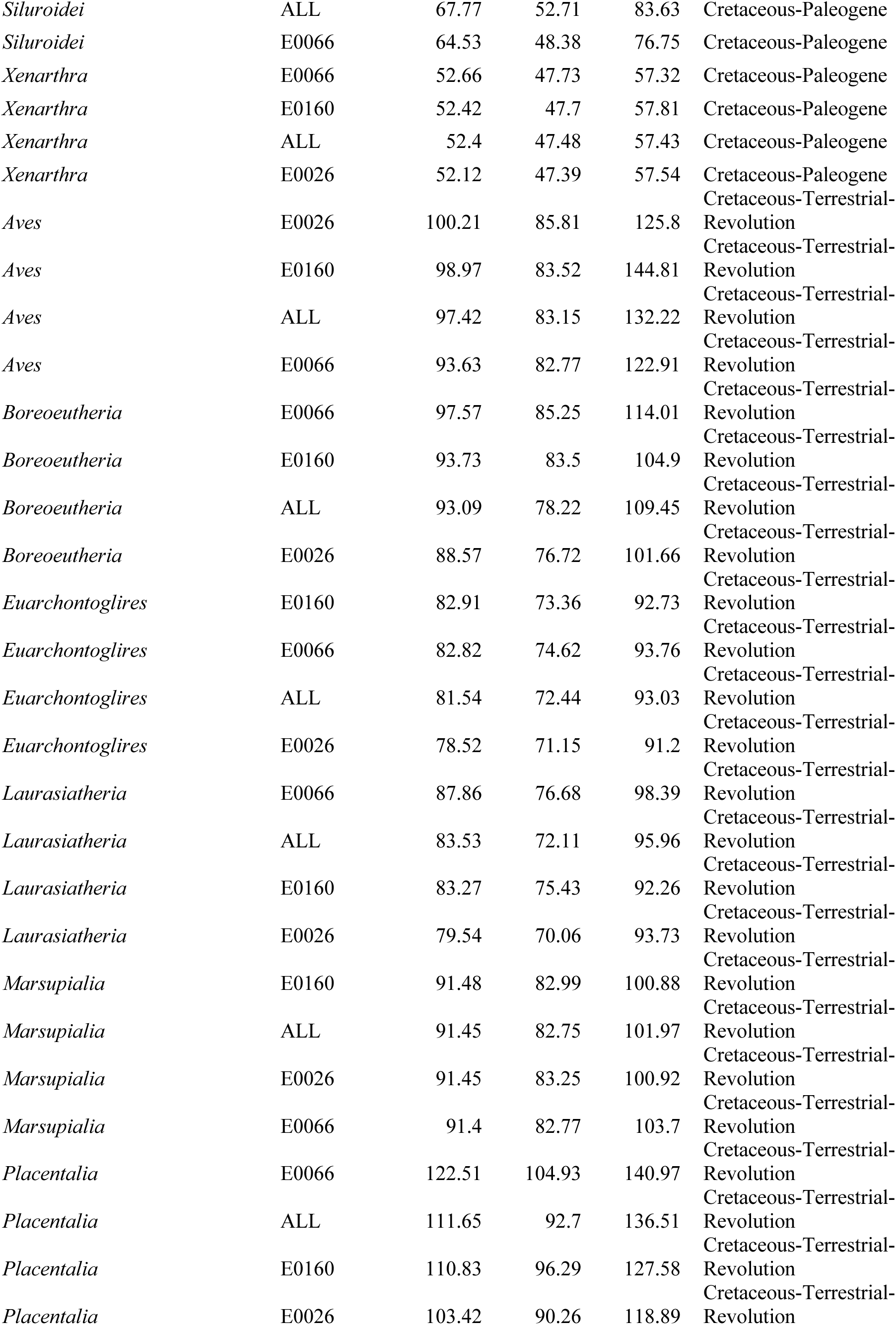
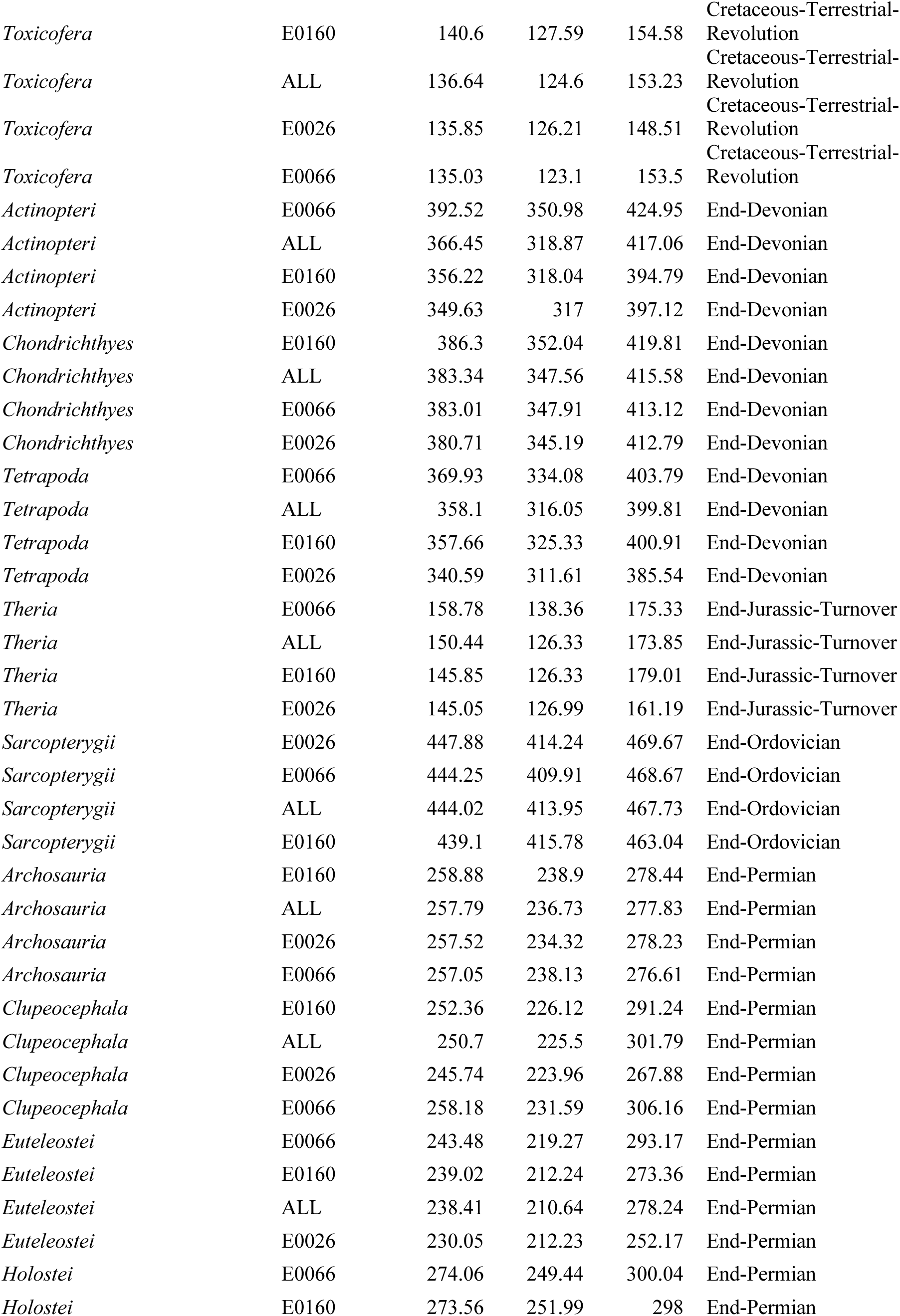
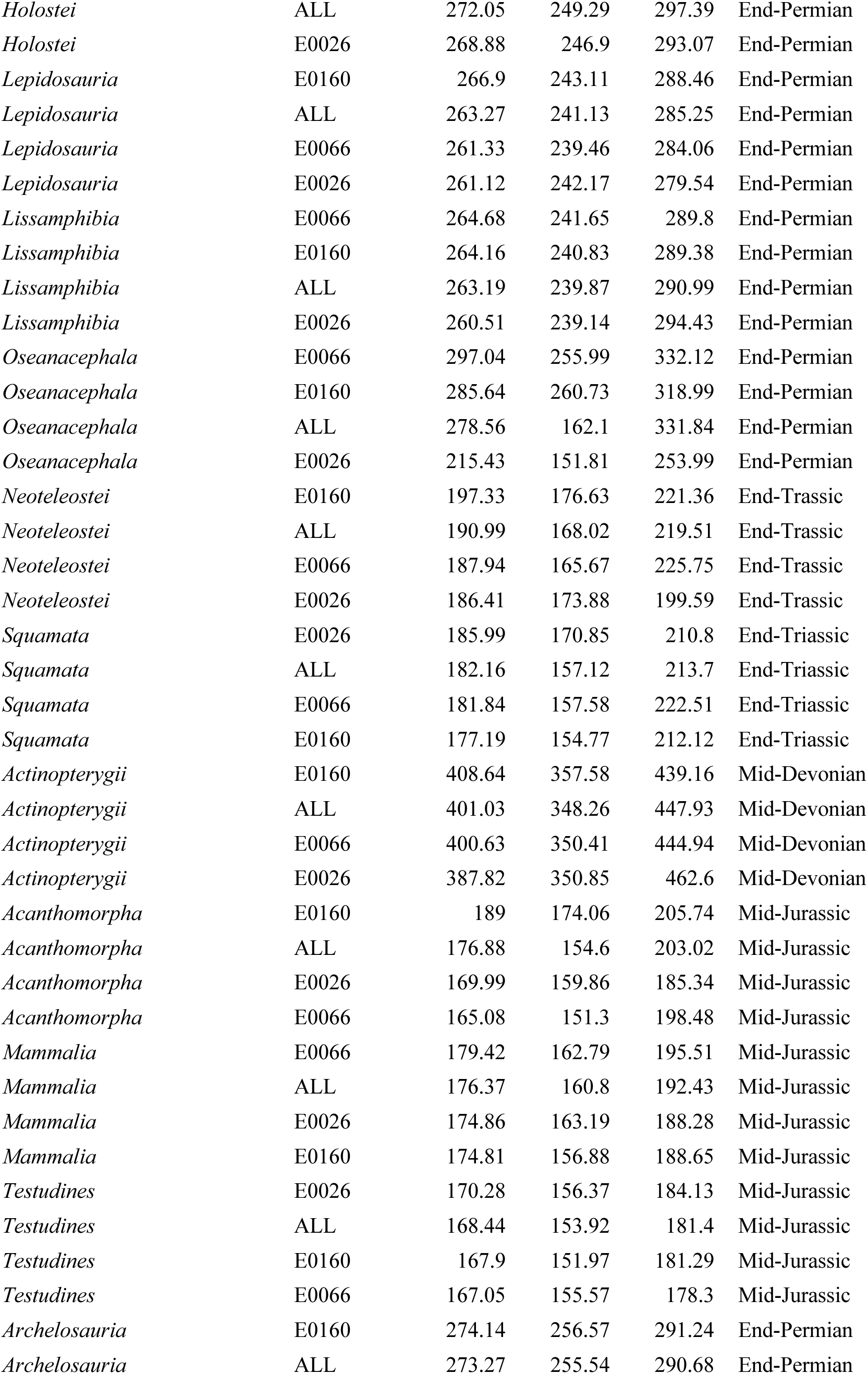
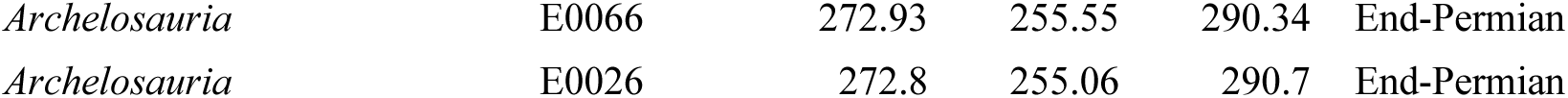
Comparison of divergence time estimates for key clades estimated from BEAST2 node dating analyses of separate and combined exon sets. Mean and 95% HPDs (Min, Max) are shown. Note: *Cetoartiodactyla* and *Eulipotypha* may not fully sample MRCA of these clades. ALL, all three sets; E0066, set 1; E0160, set 2; E0026, set 3.

**Table S3.** Input Prior Bounds for Full Analysis. Priors on individual nodes in the analysis of the ‘ALL’ dataset.

| Clade | Mean | Offset | 2.5% Q | 5% Q | Median | 95% Q | 97.5% Q | Distribution Type |
| --- | --- | --- | --- | --- | --- | --- | --- | --- |
| *Acipenseriformes* | 133.5 | 0.05 | 121 | 123 | 133 | 145 | 147 | Lognormal, Real Time |
| *Archelosauria* | 283 | 0.05 | 256 | 260 | 283 | 307 | 312 | Lognormal, Real Time |
| *Archosauria* | 267 | 0.05 | 242 | 246 | 267 | 290 | 294 | Lognormal, Real Time |
| *Arctoidea* | 31 | 0.05 | 28.1 | 28.5 | 31 | 33.6 | 34.1 | Lognormal, Real Time |
| *Beryciformes* | 72.9 | 0.05 | 66 | 67.1 | 72.8 | 79.1 | 80.3 | Lognormal, Real Time |
| *Carnivora* | 41.6 | 0.05 | 37.7 | 38.3 | 41.5 | 45.1 | 45.8 | Lognormal, Real Time |
| *Cetacea* | 40.2 | 0.05 | 36.4 | 37 | 40.1 | 43.6 | 44.3 | Lognormal, Real Time |
| *Chiroptera* | 57.4 | 0.05 | 52 | 52.8 | 57.3 | 62.2 | 63.2 | Lognormal, Real Time |
| *Chondrichthyes* | 397 | 0.05 | 359 | 365 | 397 | 430 | 437 | Lognormal, Real Time |
| *Cypriniformes* | 53.9 | 0.05 | 48.8 | 49.6 | 53.8 | 58.4 | 59.4 | Lognormal, Real Time |
| *Dipnoi* | 154.3 | 0.05 | 140 | 142 | 154 | 167 | 170 | Lognormal, Real Time |
| *Elopomorpha* | 138 | 0.05 | 125 | 127 | 138 | 150 | 152 | Lognormal, Real Time |
| *Emydidae* | 35.3 | 0.05 | 32 | 32.5 | 35.3 | 38.3 | 38.9 | Lognormal, Real Time |
| *Ferae* | 70.7 | 0.05 | 64 | 65 | 70.6 | 76.7 | 77.9 | Lognormal, Real Time |
| *Galloanserae* | 67.5 | 0.05 | 61.1 | 62.1 | 67.4 | 73.2 | 74.4 | Lognormal, Real Time |
| *Holostei* | 276.9 | 0.05 | 251 | 255 | 277 | 300 | 305 | Lognormal, Real Time |
| *Labridae-Parapercis* | 53.6 | 0.05 | 48.5 | 49.3 | 53.5 | 58.1 | 59 | Lognormal, Real Time |
| *Lepidosauria* | 267 | 0.05 | 242 | 246 | 267 | 290 | 294 | Lognormal, Real Time |
| *Lepisosteidae* | 107.3 | 0.05 | 97.2 | 98.7 | 107 | 116 | 118 | Lognormal, Real Time |
| *Lissamphibia* | 262 | 0.05 | 237 | 241 | 262 | 284 | 289 | Lognormal, Real Time |
| *Lophioidei* | 53.6 | 0.05 | 48.5 | 49.3 | 53.5 | 58.1 | 59 | Lognormal, Real Time |
| *Mammalia* | 170.5 | 0.05 | 154 | 157 | 170 | 185 | 188 | Lognormal, Real Time |
| *Marsupialia* | 92.3 | 0.05 | 83.6 | 84.9 | 92.2 | 100 | 102 | Lognormal, Real Time |
| *Ostariophysi* | 134 | 0.05 | 121 | 123 | 134 | 145 | 148 | Lognormal, Real Time |
| *Osteichthyes* | 467 | 0.05 | 423 | 430 | 466 | 506 | 514 | Lognormal, Real Time |
| *Osteoglossomorpha* | 125 | 0.05 | 113 | 115 | 125 | 136 | 138 | Lognormal, Real Time |
| *Otocephala* | 166.8 | 0.05 | 151 | 153 | 167 | 181 | 184 | Lognormal, Real Time |
| *Otophysi* | 123.6 | 0.05 | 112 | 114 | 123 | 134 | 136 | Lognormal, Real Time |
| *Pan-Paenungulata* | 65.4 | 0.05 | 59.2 | 60.2 | 65.3 | 70.9 | 72 | Lognormal, Real Time |
| *Pan-Acrodonta* | 78.4 | 0.05 | 71 | 72.1 | 78.3 | 85 | 86.4 | Lognormal, Real Time |
| *Pan-Caimaninae* | 83.6 | 0.05 | 75.7 | 76.9 | 83.5 | 90.7 | 92.1 | Lognormal, Real Time |
| *Pan-Coliiformes* | 68.7 | 0.05 | 62.2 | 63.2 | 68.6 | 74.5 | 75.7 | Lognormal, Real Time |
| *Pan-Galloanserae* | 75.1 | 0.05 | 68 | 69.1 | 75 | 81.4 | 82.7 | Lognormal, Real Time |
| *Pan-Lacertidae* | 72.9 | 0.05 | 66 | 67.1 | 72.8 | 79.1 | 80.3 | Lognormal, Real Time |
| *Pan-Primates* | 72.3 | 0.05 | 65.5 | 66.5 | 72.2 | 78.4 | 79.6 | Lognormal, Real Time |
| *Pan-Pythonidae* | 51.3 | 0.05 | 46.5 | 47.2 | 51.2 | 55.6 | 56.5 | Lognormal, Real Time |
| *Pan-Scincoidea* | 160 | 0.05 | 145 | 147 | 160 | 173 | 176 | Lognormal, Real Time |
| *Pan-Spheniscidae* | 66.8 | 0.05 | 60.5 | 61.4 | 66.7 | 72.4 | 73.6 | Lognormal, Real Time |
| *Pan-Struthio* | 52.8 | 0.05 | 47.8 | 48.6 | 52.7 | 57.3 | 58.2 | Lognormal, Real Time |
| *Pan-Varanidae* | 133.5 | 0.05 | 121 | 123 | 133 | 145 | 147 | Lognormal, Real Time |
| *Pan-Zeiformes* | 91.7 | 0.05 | 83 | 84.4 | 91.6 | 99.4 | 101 | Lognormal, Real Time |
| *Pipoidea-Neobatrachia* | 163 | 0.05 | 148 | 150 | 163 | 177 | 180 | Lognormal, Real Time |
| *Rodentia* | 61.8 | 0.05 | 56 | 56.8 | 61.7 | 67 | 68.1 | Lognormal, Real Time |
| *Pan-Dipnoi* | 462 | 0.05 | 418 | 425 | 461 | 501 | 509 | Lognormal, Real Time |
| *Scombriformes* | 59.9 | 0.05 | 54.2 | 55.1 | 59.8 | 65 | 66 | Lognormal, Real Time |
| *Siluriformes* | 92.2 | 0.05 | 83.5 | 84.8 | 92.1 | 100 | 102 | Lognormal, Real Time |
| *Syngnathiformes* | 95 | 0.05 | 86 | 87.4 | 94.9 | 103 | 105 | Lognormal, Real Time |
| *Testudines* | 170.5 | 0.05 | 154 | 157 | 170 | 185 | 188 | Lognormal, Real Time |
| *Tetraodontoidei* | 61.8 | 0.05 | 56 | 56.8 | 61.7 | 67 | 68.1 | Lognormal, Real Time |
| *Trionychia* | 145 | 0.05 | 131 | 133 | 145 | 157 | 160 | Lognormal, Real Time |
| *Xenarthra* | 52.2 | 0.05 | 47.3 | 48 | 52.1 | 56.6 | 57.5 | Lognormal, Real Time |
| *Pan-Carangidae* | 61.6 | 0.05 | 55.8 | 56.7 | 61.5 | 66.8 | 67.9 | Lognormal, Real Time |

**Table S4.** Comparison of divergence time estimates for vertebrate clades estimated in the taxon exclusion rate test experiments. Mean and 95% HPDs (Min, Max) are shown. Abbreviatons: Subset, No Cal. On LF= subset analysis without node calibrations on living fossil lineages. Subset, LF with Cal. = subset analysis with node calibrations on living fossils. Subset, without LF = subset analysis excluding living fossil lineages. All mean rates (substitutions per site per Ma) are log10 transformed.

| Clade | Analysis | Median | Min | Max | Rate_Root | Rate_Min | Rate_Max | Rate_Root_Log | Rate_Min_Log | Rate_Max_Log |
| --- | --- | --- | --- | --- | --- | --- | --- | --- | --- | --- |
| *Teleostei* | Full Tree (Figure 1) | 284.12 | 242.11 | 326.26 | 0.002556 | 0.001046 | 0.006375 | -2.5924392 | -2.9804683 | -2.1955198 |
| *Actinopteri* | Full Tree (Figure 1) | 366.67 | 318.87 | 417.06 | 0.001167 | 0.000468 | 0.00272 | -2.9329291 | -3.3297541 | -2.5654311 |
| *Neopterygii* | Full Tree (Figure 1) | 322.92 | 279.66 | 370.27 | 0.001791 | 0.000662 | 0.004479 | -2.7469044 | -3.179142 | -2.3488189 |
| *Holostei* | Full Tree (Figure 1) | 272.79 | 249.29 | 297.39 | 0.000569 | 0.000216 | 0.00165 | -3.2448877 | -3.6655462 | -2.7825161 |
| *Lepisosteidae* | Full Tree (Figure 1) | 104.51 | 95.03 | 114.56 | 0.000594 | 0.000484 | 0.000719 | -3.2262136 | -3.3151546 | -3.1432711 |
| *Acipenseriformes* | Full Tree (Figure 1) | 131.54 | 118.87 | 144.76 | 0.000442 | 0.000308 | 0.00116 | -3.3545777 | -3.5114493 | -2.935542 |
| *Actinopterygii* | Full Tree (Figure 1) | 399.56 | 348.26 | 417.06 | 0.001425 | 0.000575 | 0.002946 | -2.8461851 | -3.2403322 | -2.5307673 |
| *Dipnoi* | Full Tree (Figure 1) | 154.07 | 139.35 | 168.66 | 0.0004 | 0.000323 | 0.000483 | -3.39794 | -3.4907975 | -3.3160529 |
| *Teleostei* | Subset, No Cal. On LF | 228.34 | 187.45 | 439.13 | 0.001685 | 0.000313 | 0.020984 | -2.7734001 | -3.5044557 | -1.6781117 |
| *Actinopteri* | Subset, No Cal. On LF | 327.55 | 291.68 | 442.71 | 0.001814 | 0.000533 | 0.064645 | -2.7413627 | -3.2732728 | -1.1894651 |
| *Neopterygii* | Subset, No Cal. On LF | 284.57 | 258.9 | 440.96 | 0.002516 | 0.000858 | 0.066502 | -2.5992894 | -3.0665127 | -1.1771653 |
| *Holostei* | Subset, No Cal. On LF | 257.73 | 116.06 | 282.36 | 0.001394 | 0.000098 | 0.024633 | -2.8557372 | -4.0087739 | -1.6084827 |
| *Lepisosteidae* | Subset, No Cal. On LF | 9.8 | 2.16 | 31.72 | 0.000392 | 0.000323 | 0.000991 | -3.4067139 | -3.4907975 | -3.0039263 |
| *Acipenseriformes* | Subset, No Cal. On LF | 22.92 | 5.01 | 52.89 | 0.000369 | 0.000237 | 0.002368 | -3.4329736 | -3.6252517 | -2.6256183 |
| *Actinopterygii* | Subset, No Cal. On LF | 378.68 | 327.49 | 443.66 | 0.001955 | 0.000013 | 0.018144 | -2.7088532 | -4.8860566 | -1.741267 |
| *Dipnoi* | Subset, No Cal. On LF | 137.15 | 35.55 | 257.7 | 0.000398 | 0.000273 | 0.000708 | -3.4001169 | -3.5638374 | -3.1499667 |
| *Teleostei* | Subset, LF with Cal. | 249.25 | 197.4 | 313.5 | 0.001909 | 0.00052 | 0.005589 | -2.7191941 | -3.2839967 | -2.2526659 |
| *Actinopteri* | Subset, LF with Cal. | 321.82 | 274.68 | 369.55 | 0.001509 | 0.0004 | 0.005112 | -2.8213108 | -3.39794 | -2.2914092 |
| *Neopterygii* | Subset, LF with Cal. | 292.11 | 263.66 | 331.31 | 0.003521 | 0.000903 | 0.013466 | -2.453334 | -3.0443123 | -1.8707614 |
| *Holostei* | Subset, LF with Cal. | 261.69 | 48.63 | 287.09 | 0.001202 | 0.000211 | 0.198062 | -2.9200955 | -3.6757175 | -0.7031988 |
| *Lepisosteidae* | Subset, LF with Cal. | 101.91 | 47.4 | 112.05 | 0.000611 | 0.000407 | 0.094379 | -3.2139588 | -3.3904056 | -1.0251246 |
| *Acipenseriformes* | Subset, LF with Cal. | 132.33 | 119.27 | 145.81 | 0.000555 | 0.000344 | 0.000742 | -3.255707 | -3.4634416 | -3.1295961 |
| *Actinopterygii* | Subset, LF with Cal. | 377.23 | 321.05 | 443.73 | 0.00202 | 0.000008 | 0.009097 | -2.6946486 | -5.09691 | -2.0411018 |
| *Dipnoi* | Subset, LF with Cal. | 121.52 | 21.02 | 303.91 | 0.000391 | 0.000262 | 0.000964 | -3.4078232 | -3.5816987 | -3.015923 |
| *Teleostei* | Subset, without LF | 220.93 | 191.9 | 252.2 | 0.001819 | 0.000918 | 0.00346 | -2.7401673 | -3.0371573 | -2.4609239 |
| *Actinopterygii* | Subset, without LF | 312.28 | 257.64 | 376.84 | 0.001135 | 0.000514 | 0.002419 | -2.9450041 | -3.2890369 | -2.6163641 |
| *Dipnoi* | Subset, without LF | 145.66 | 56.31 | 227.41 | 0.000409 | 0.000293 | 0.000598 | -3.3882767 | -3.5331324 | -3.2232988 |

---

**Figure S1.**
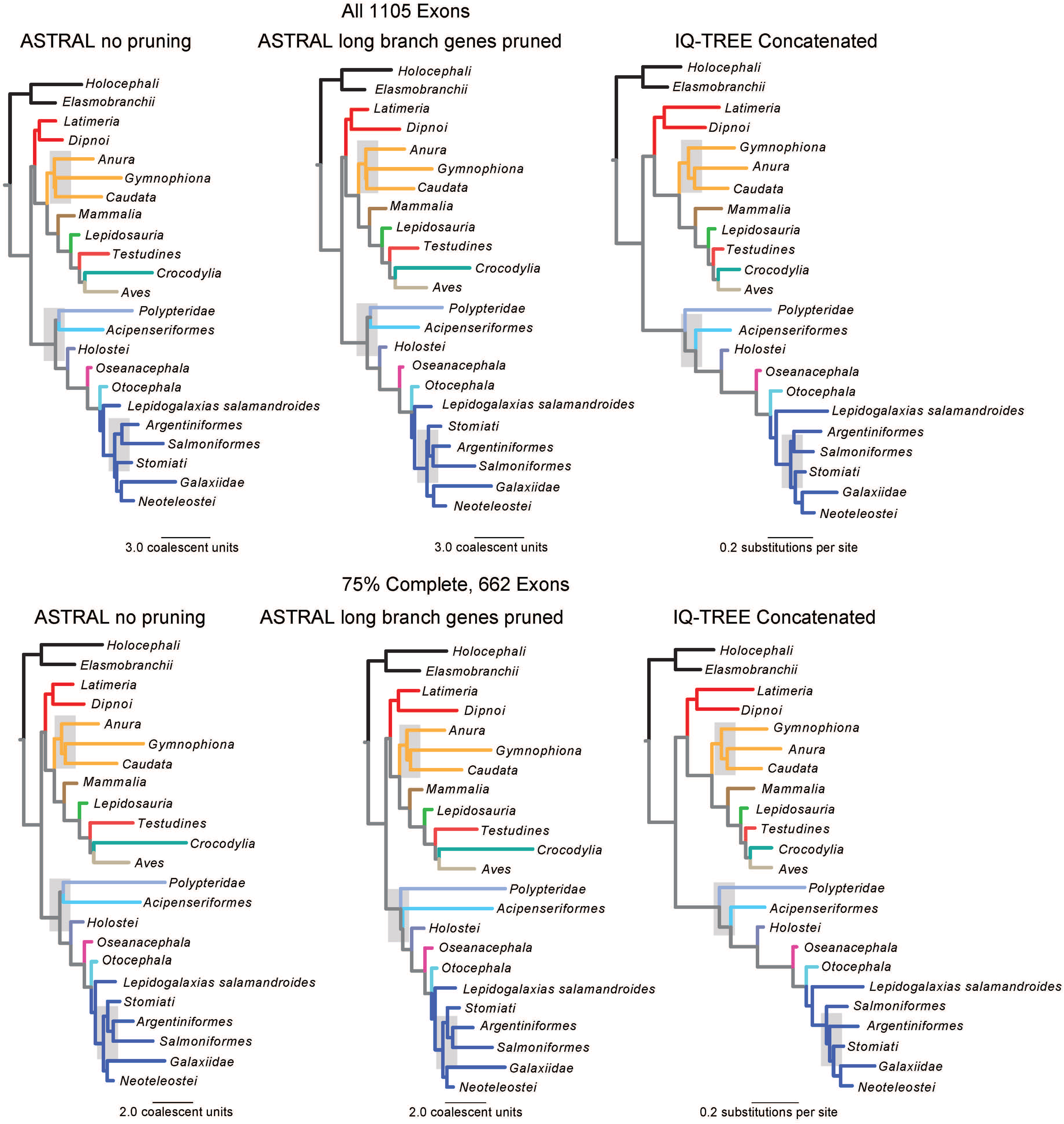
Comparison of phylogenetic trees. Simplified phylogenies of jawed vertebrates comparing the topologies recovered using ASTRAL-III or maximum likelihood analysis of concatenated sequence data for the whole 1105 exon dataset (top row) and the 75% complete matrix (bottom row). Clades are color-coded according to their parent lineages: *Chondrichthyes* in black, non-tetrapod sarcopterygians in red, *Amphibia* in yellow, *Euteleostei* in dark blue.

**Figure S2.**
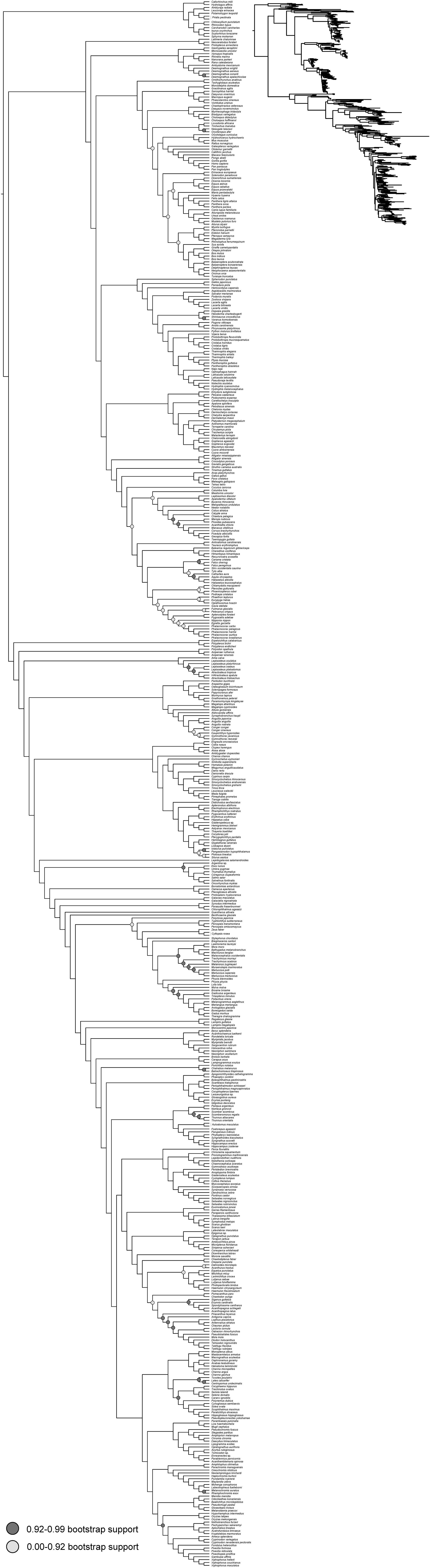
Complete concatenated phylogenies. Phylogeny of 540 jawed vertebrates recovered using maximum likelihood analysis of concatenated sequence data for the whole 1105 exon dataset.

**Figure S3.**
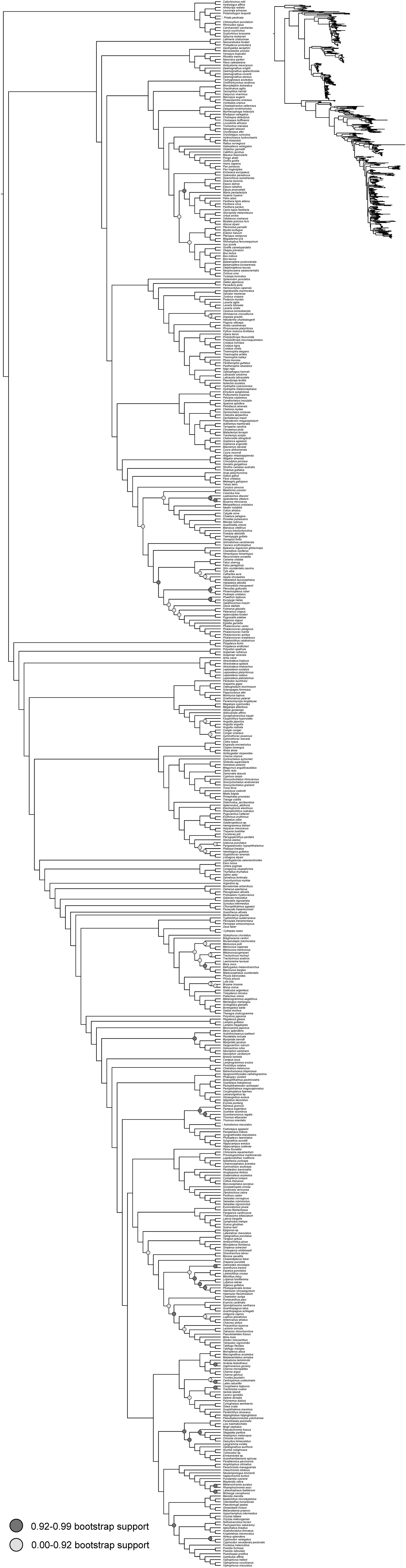
Complete concatenated phylogenies. Phylogeny of 540 jawed vertebrates recovered using maximum likelihood analysis of concatenated sequence data for the 662 (75% complete matrix) exon dataset.

**Figure S4.**
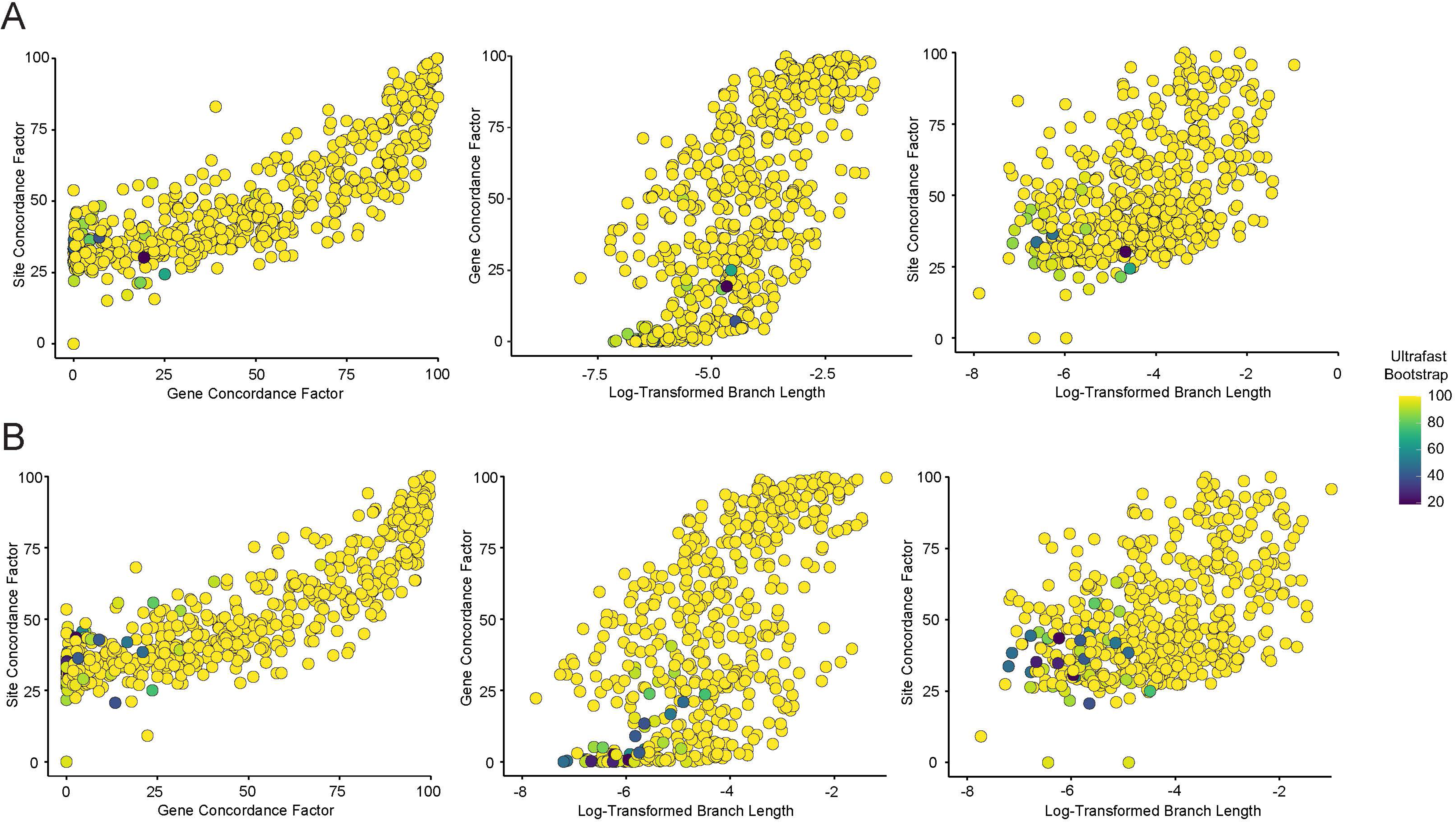
Relationships among gene and site concordance factors, bootstrap support values, and branch lengths. Scatterplots show relationships between support values and branch lengths for the whole 1105 exon dataset (A) and the 75% complete matrix (B).

**Figure S5.**
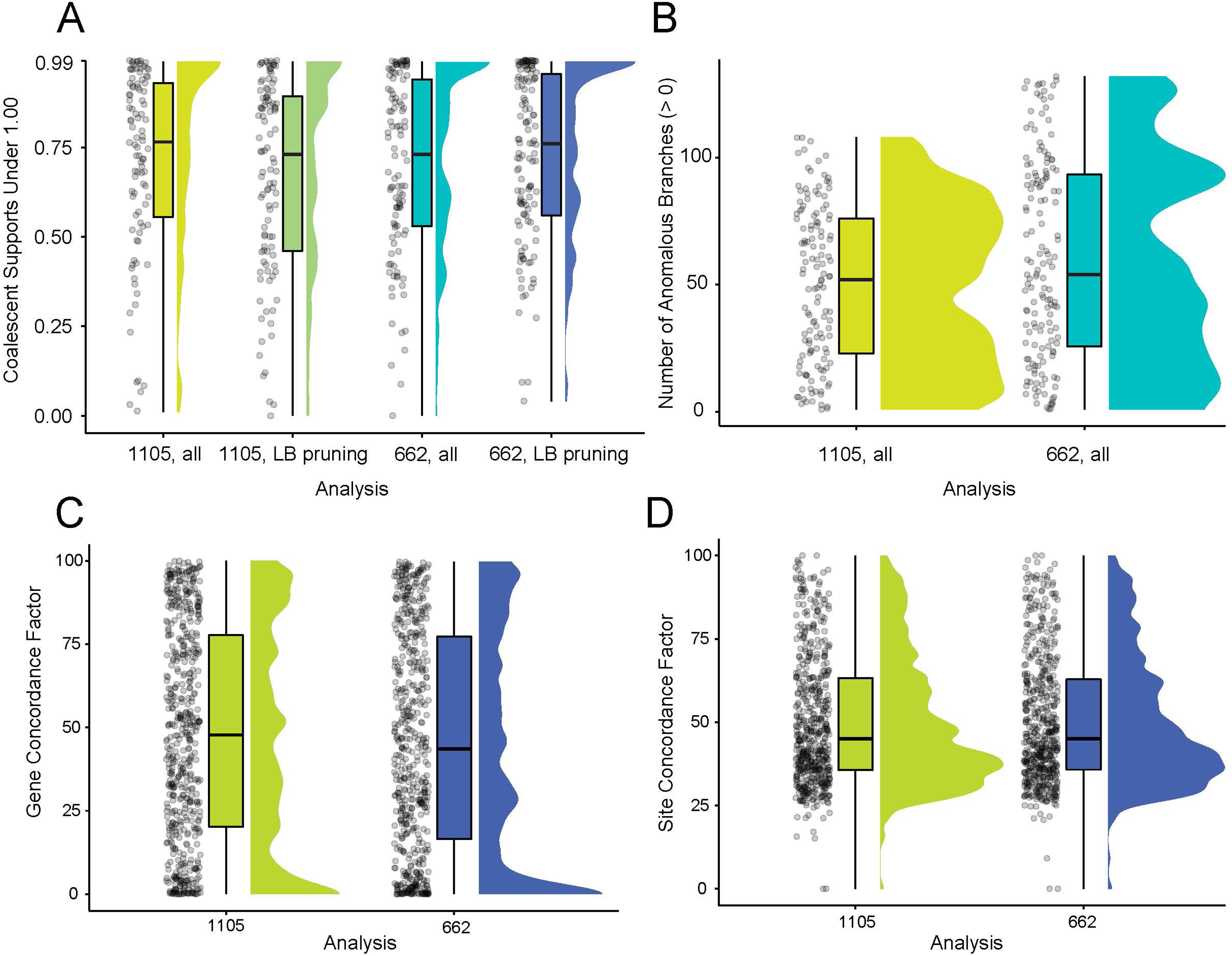
Summary of nodal support across phylogenies. Violin plots show (A) coalescent support values under 1.0 across different ASTRAL-III species trees, (B) minimum number of anomalous branches for nodes in anomaly zones in the two analyzed ASTRAL-III species trees, and (C) gene and (D) site concordance factors across the phylogenies inferred via maximum likelihood analyses of concatenated exon sequences. 1105 = all exons, 662 = 75% complete matrix.

**Figure S6.**
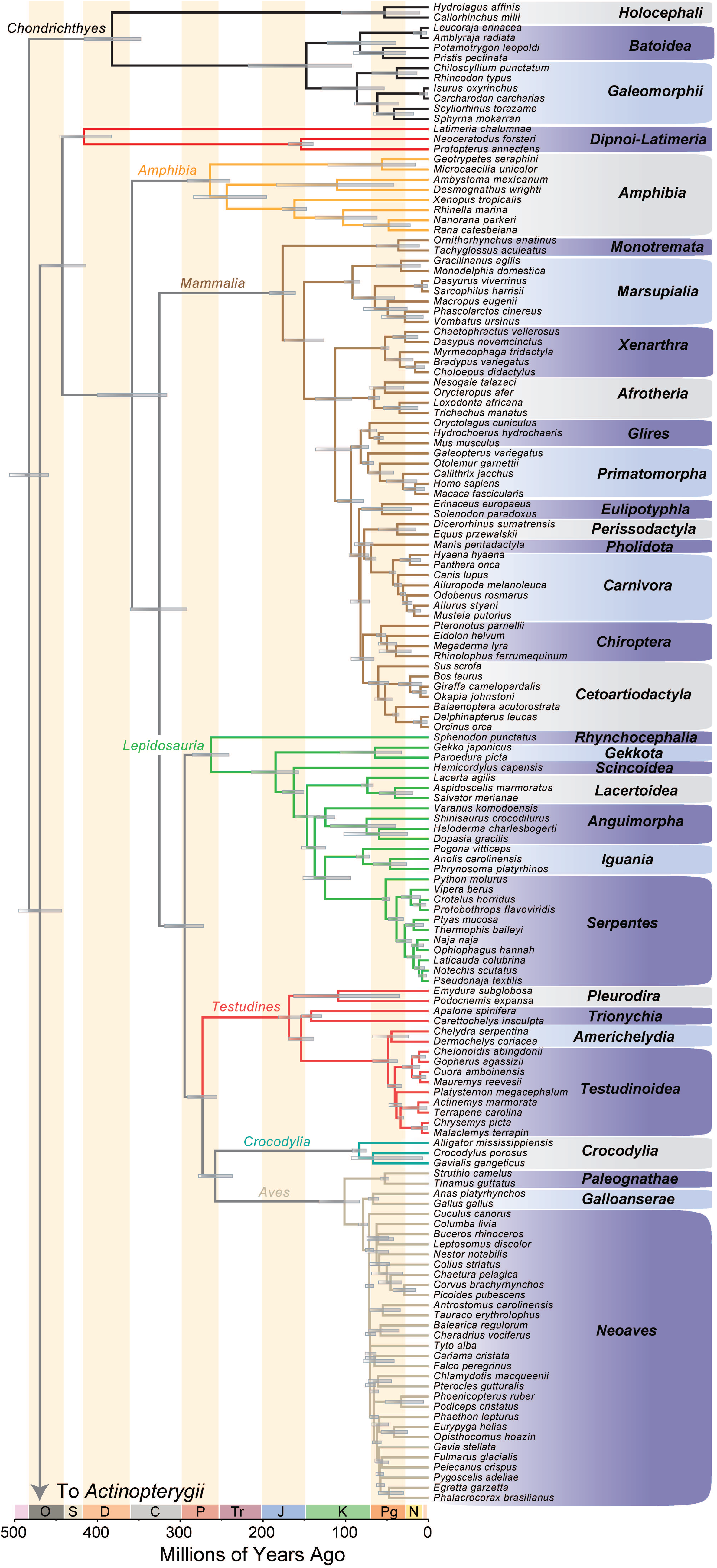
Main time-calibrated phylogeny of jawed vertebrates, Part I. Phylogeny of jawed vertebrates (*Gnathostomata*) estimated under a Bayesian node-dating approach, showing the relationships of *Chondrichthyes* and *Sarcopterygii*. Bars at nodes represent 95% highest posterior density intervals for divergence times. Abbreviations: O, Ordovician; S, Silurian; D, Devonian; C, Carboniferous; P, Permian; Tr, Triassic; J, Jurassic; K, Cretaceous; Pg, Paleogene; N, Neogene.

**Figure S7.**
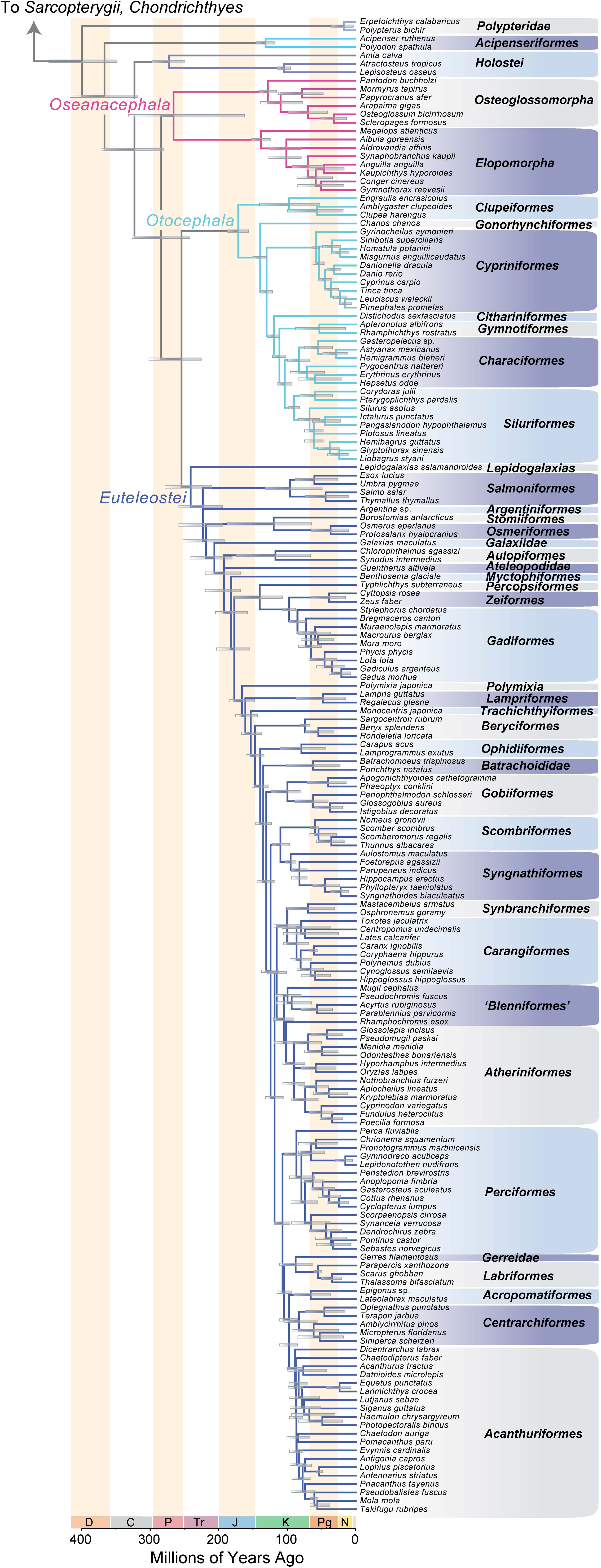
Main time-calibrated phylogeny of jawed vertebrates, Part II. Phylogeny of jawed vertebrates (*Gnathostomata*) estimated under a Bayesian node-dating approach, showing the relationships of *Actinopterygii.* Bars at nodes represent 95% highest posterior density intervals for divergence times. Abbreviations: O, Ordovician; S, Silurian; D, Devonian; C, Carboniferous; P, Permian; Tr, Triassic; J, Jurassic; K, Cretaceous; Pg, Paleogene; N, Neogene.

**Figure S8.**
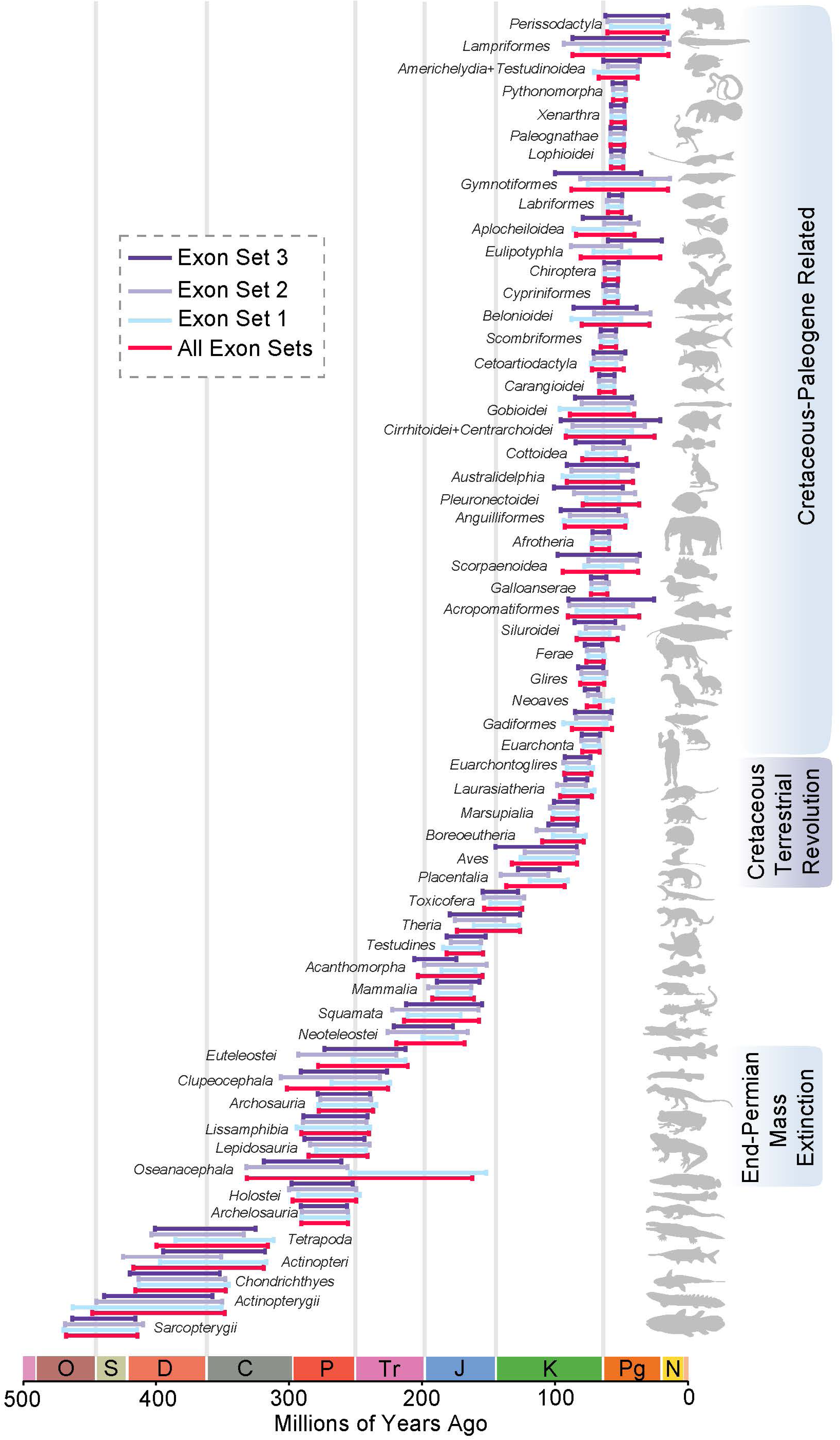
Comparison of estimated divergence times. Plot shows estimated median ages and 95% HPD intervals for selected clades of jawed vertebrates found in the time tree made by pooling all three sets of 50 exons and from trees built from each exon set. Abbreviations: O, Ordovician; S, Silurian; D, Devonian; C, Carboniferous; P, Permian; Tr, Triassic; J, Jurassic; K, Cretaceous; Pg, Paleogene; N, Neogene.

**Figure S9.**
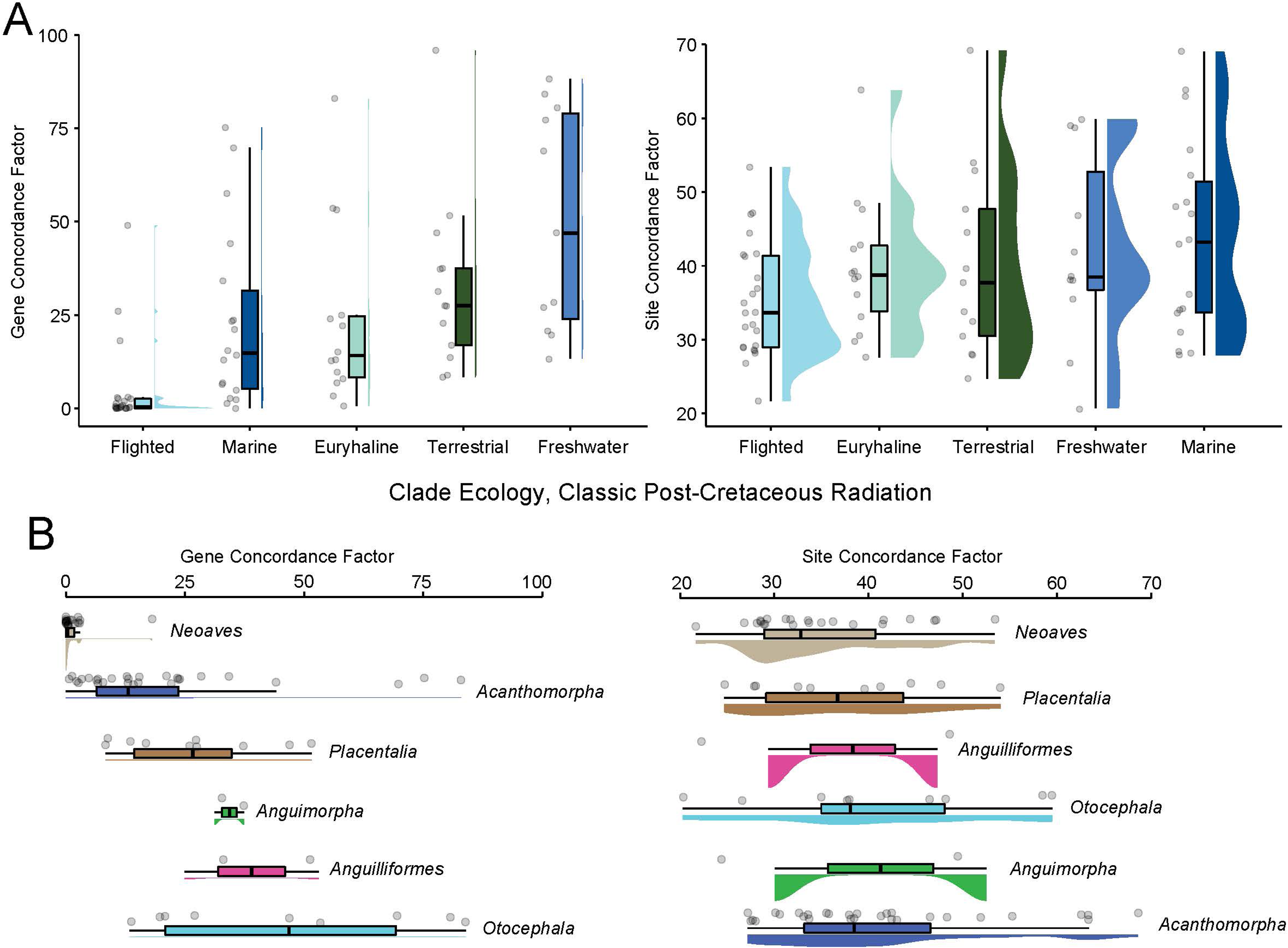
Signature of the Cretaceous-Paleogene Mass Extinction on Jawed Vertebrate Phylogenetic Resolution. Violin plots show gene and site concordance factors for nodes that appear within 10 million years of the Cretaceous-Paleogene boundary (66.02 Ma)(Gradstein et al. 2021) and grouped according to (A) ecology and (B) clade.

**Figure S10.**
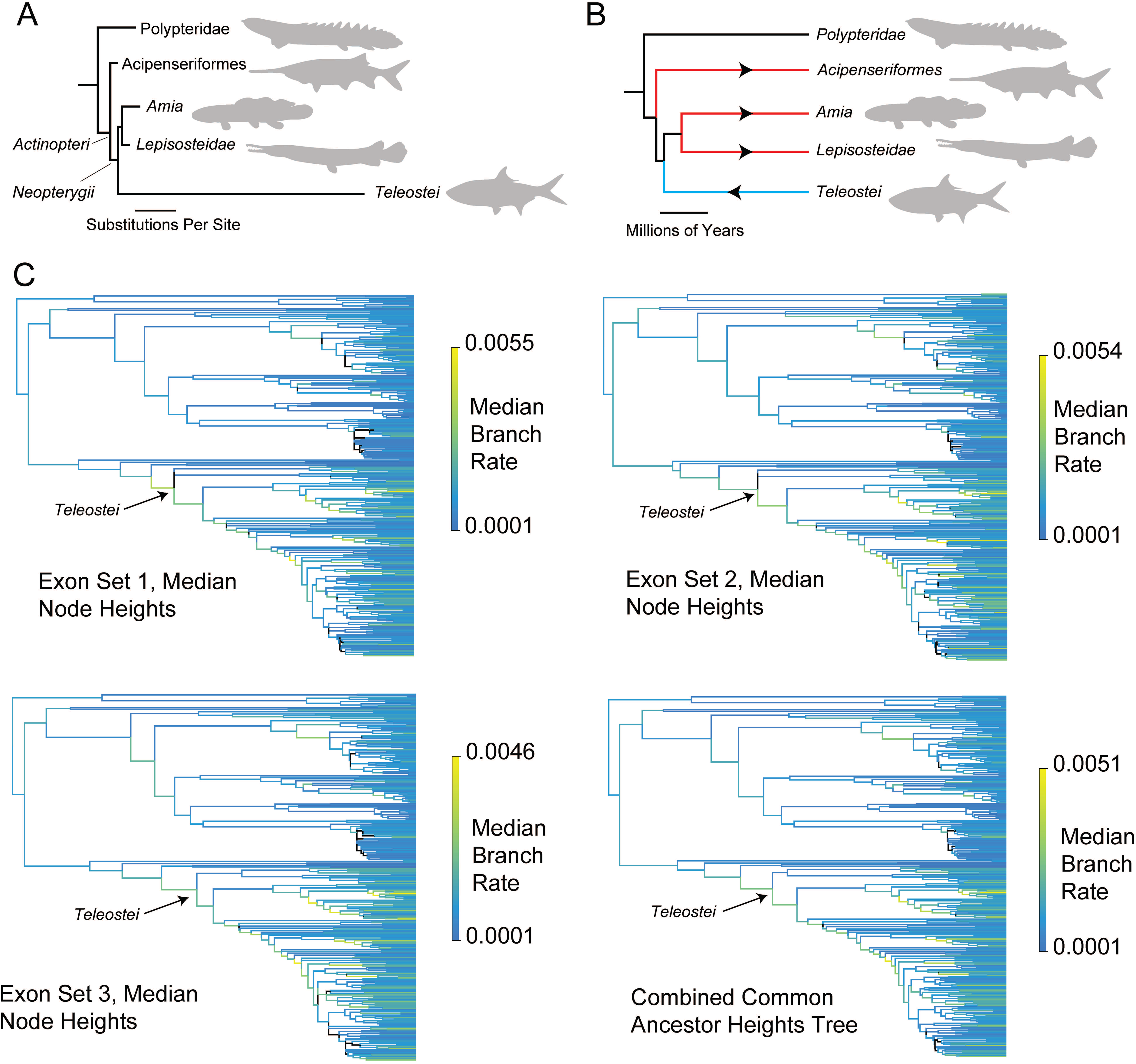
Rates of Jawed Vertebrate Evolution Part I. Hypothesis of how variation in rates of nucleotide substitution (A) can distort the inferred timescale (B) of ray-finned fish evolution. (C) shows the main time-calibrated phylogeny of jawed vertebrates (see Figure 1) and trees built by annotating the target tree using posterior tree sets from analyses of each exon set, with branches colored according to their inferred substitution rate values (substitutions per site million years. Note the very high backbone rates associated with *Actinopterygii*.

**Figure S11.**
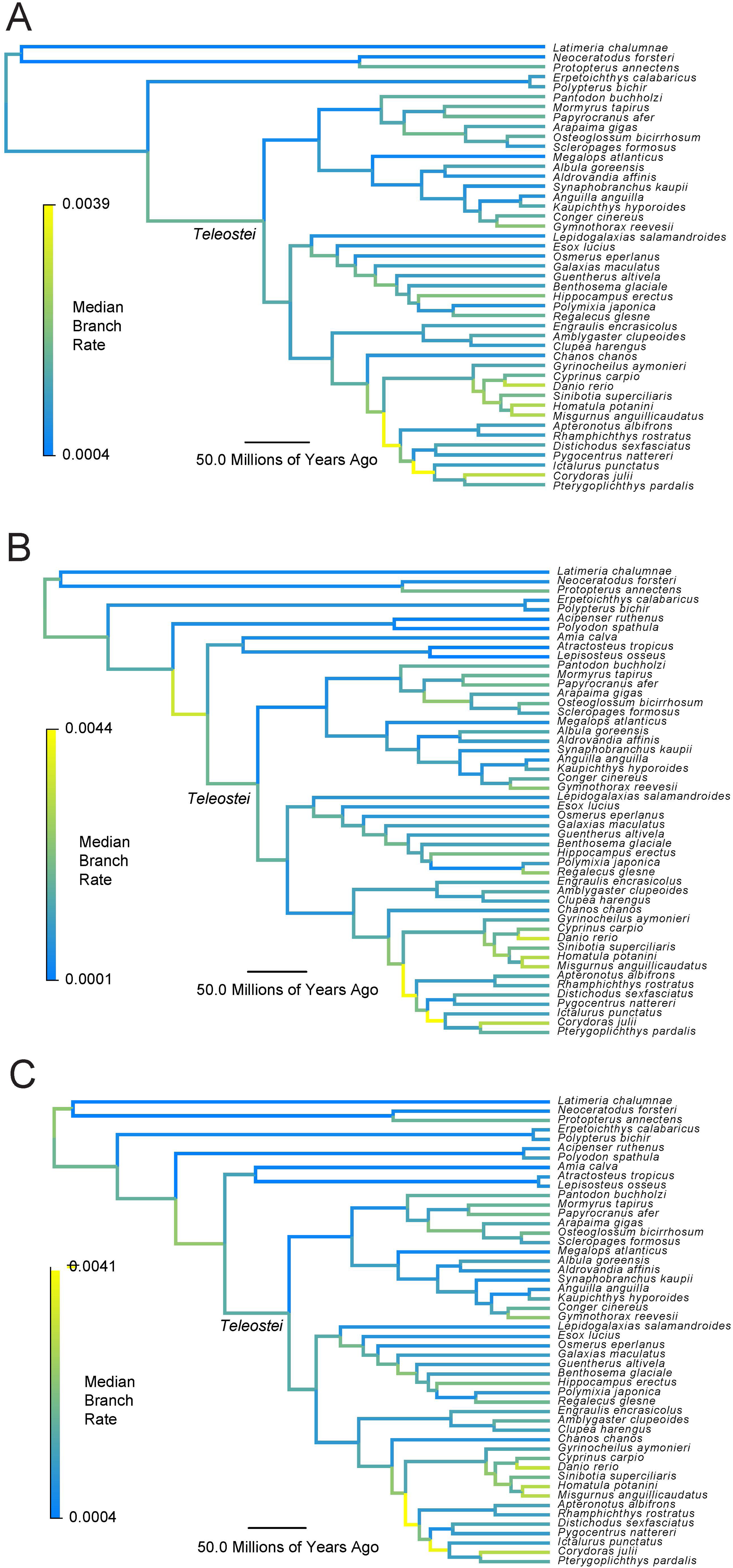
Rates of Jawed Vertebrate Evolution Part II. Panel (A) shows the test analysis in which living fossil lineages (*Acipenseriformes*, *Holostei*) were excluded (see Figure 5, Methods), panel (B) shows the full reduced sampling time-calibrated phylogeny of jawed vertebrates (see Figure 5), and panel (C) shows the full reduced sampling time-calibrated phylogeny of jawed vertebrates, but with no fossil calibrations for living fossil clades (see Figure 5).

## References

Alda, F., W. B. Ludt, D. J. Elías, C. D. McMahan, and P. Chakrabarty. 2021. Comparing Ultraconserved Elements and Exons for Phylogenomic Analyses of Middle American Cichlids: When Data Agree to Disagree. Genome Biol. Evol. 13:evab161.

Alfaro, M. E., B. C. Faircloth, R. C. Harrington, L. Sorenson, M. Friedman, C. E. Thacker, C. H. Oliveros, D. Černý, and T. J. Near. 2018. Explosive diversification of marine fishes at the Cretaceous–Palaeogene boundary. Nat. Ecol. Evol. 2:688–696. Nature Publishing Group.

Alroy, J. 1999. The Fossil Record of North American Mammals: Evidence for a Paleocene Evolutionary Radiation. Syst. Biol. 48:107–118.

Álvarez-Carretero, S., A. U. Tamuri, M. Battini, F. F. Nascimento, E. Carlisle, R. J. Asher, Z. Yang, P. C. J. Donoghue, and M. dos Reis. 2022. A species-level timeline of mammal evolution integrating phylogenomic data. Nature 602:263–267. Nature Publishing Group.

Amemiya, C. T., J. Alföldi, A. P. Lee, S. Fan, H. Philippe, I. MacCallum, I. Braasch, T. Manousaki, I. Schneider, N. Rohner, C. Organ, D. Chalopin, J. J. Smith, M. Robinson, R. A. Dorrington, M. Gerdol, B. Aken, M. A. Biscotti, M. Barucca, D. Baurain, A. M. Berlin, G. L. Blatch, F. Buonocore, T. Burmester, M. S. Campbell, A. Canapa, J. P. Cannon, A. Christoffels, G. De Moro, A. L. Edkins, L. Fan, A. M. Fausto, N. Feiner, M. Forconi, J. Gamieldien, S. Gnerre, A. Gnirke, J. V. Goldstone, W. Haerty, M. E. Hahn, U. Hesse, S. Hoffmann, J. Johnson, S. I. Karchner, S. Kuraku, M. Lara, J. Z. Levin, G. W. Litman, E. Mauceli, T. Miyake, M. G. Mueller, D. R. Nelson, A. Nitsche, E. Olmo, T. Ota, A. Pallavicini, S. Panji, B. Picone, C. P. Ponting, S. J. Prohaska, D. Przybylski, N. R. Saha, V. Ravi, F. J. Ribeiro, T. Sauka-Spengler, G. Scapigliati, S. M. J. Searle, T. Sharpe, O. Simakov, P. F. Stadler, J. J. Stegeman, K. Sumiyama, D. Tabbaa, H. Tafer, J. Turner-Maier, P. van Heusden, S. White, L. Williams, M. Yandell, H. Brinkmann, J.-N. Volff, C. J. Tabin, N. Shubin, M. Schartl, D. B. Jaffe, J. H. Postlethwait, B. Venkatesh, F. Di Palma, E. S. Lander, A. Meyer, and K. Lindblad-Toh. 2013. The African coelacanth genome provides insights into tetrapod evolution. Nature 496:311–316. Nature Publishing Group.

Anderson, J. S., R. R. Reisz, D. Scott, N. B. Fröbisch, and S. S. Sumida. 2008. A stem batrachian from the Early Permian of Texas and the origin of frogs and salamanders. Nature 453:515–518. Nature Publishing Group.

Andreev, P. S., I. J. Sansom, Q. Li, W. Zhao, J. Wang, C.-C. Wang, L. Peng, L. Jia, T. Qiao, and M. Zhu. 2022. Spiny chondrichthyan from the lower Silurian of South China. Nature 609:969–974. Nature Publishing Group.

Archibald, J. D., and D. H. Deutschman. 2001. Quantitative Analysis of the Timing of the Origin and Diversification of Extant Placental Orders. J. Mamm. Evol. 8:107–124.

Arcila, D., L. C. Hughes, B. Meléndez-Vazquez, C. C. Baldwin, W. T. White, K. E. Carpenter, J. T. Williams, M. D. Santos, J. J. Pogonoski, M. Miya, G. Ortí, and R. Betancur-R. 2021. Testing the Utility of Alternative Metrics of Branch Support to Address the Ancient Evolutionary Radiation of Tunas, Stromateoids, and Allies (Teleostei: Pelagiaria). Syst. Biol. 70:1123–1144.

Argyriou, T., S. Giles, and M. Friedman. n.d. A Permian fish reveals widespread distribution of neopterygian-like jaw suspension. eLife 11:e58433.

Argyriou, T., S. Giles, M. Friedman, C. Romano, I. Kogan, and M. R. Sánchez-Villagra. 2018. Internal cranial anatomy of Early Triassic species of †Saurichthys (Actinopterygii:†Saurichthyiformes): implications for the phylogenetic placement of †saurichthyiforms. BMC Evol. Biol. 18:161.

Bagley, J. C., R. L. Mayden, and P. M. Harris. 2018. Phylogeny and divergence times of suckers (Cypriniformes: Catostomidae) inferred from Bayesian total-evidence analyses of molecules, morphology, and fossils. PeerJ 6. PeerJ, Inc.

Bernardi, M., H. Klein, F. M. Petti, and M. D. Ezcurra. 2015. The Origin and Early Radiation of Archosauriforms: Integrating the Skeletal and Footprint Record. PLoS ONE 10:e0128449.

Berv, J. S., and D. J. Field. 2018. Genomic Signature of an Avian Lilliput Effect across the K-Pg Extinction. Syst. Biol. 67:1–13.

Berv, J. S., S. Singhal, D. J. Field, N. Walker-Hale, S. W. McHugh, J. R. Shipley, E. T. Miller, R. T. Kimball, E. L. Braun, A. Dornburg, C. T. Parins-Fukuchi, R. O. Prum, B. M. Winger, M. Friedman, and S. A. Smith. 2024. Genome and life-history evolution link bird diversification to the end-Cretaceous mass extinction. Sci. Adv. 10:eadp0114. American Association for the Advancement of Science.

Betancur-R, R., R. E. Broughton, E. O. Wiley, K. Carpenter, J. A. López, C. Li, N. I. Holcroft, D. Arcila, M. Sanciangco, J. C. C. Ii, F. Zhang, T. Buser, M. A. Campbell, J. A. Ballesteros, A. Roa-Varon, S. Willis, W. C. Borden, T. Rowley, P. C. Reneau, D. J. Hough, G. Lu, T. Grande, G. Arratia, and G. Ortí. 2013. The Tree of Life and a New Classification of Bony Fishes. PLOS Curr. Tree Life, doi: 10.1371/currents.tol.53ba26640df0ccaee75bb165c8c26288. Public Library of Science.

Betancur-R, R., E. O. Wiley, G. Arratia, A. Acero, N. Bailly, M. Miya, G. Lecointre, and G. Ortí. 2017. Phylogenetic classification of bony fishes. BMC Evol. Biol. 17:162.

Bi, X., K. Wang, L. Yang, H. Pan, H. Jiang, Q. Wei, M. Fang, H. Yu, C. Zhu, Y. Cai, Y. He, X. Gan, H. Zeng, D. Yu, Y. Zhu, H. Jiang, Q. Qiu, H. Yang, Y. E. Zhang, W. Wang, M. Zhu, S. He, and G. Zhang. 2021. Tracing the genetic footprints of vertebrate landing in non-teleost ray-finned fishes. Cell 184:1377–1391.e14.

Bian, C., Y. Hu, V. Ravi, I. S. Kuznetsova, X. Shen, X. Mu, Y. Sun, X. You, J. Li, X. Li, Y. Qiu, B.-H. Tay, N. M. Thevasagayam, A. S. Komissarov, V. Trifonov, M. Kabilov, A. Tupikin, J. Luo, Y. Liu, H. Song, C. Liu, X. Wang, D. Gu, Y. Yang, W. Li, G. Polgar, G. Fan, P. Zeng, H. Zhang, Z. Xiong, Z. Tang, C. Peng, Z. Ruan, H. Yu, J. Chen, M. Fan, Y. Huang, M. Wang, X. Zhao, G. Hu, H. Yang, J. Wang, J. Wang, X. Xu, L. Song, G. Xu, P. Xu, J. Xu, S. J. O’Brien, L. Orbán, B. Venkatesh, and Q. Shi. 2016. The Asian arowana (Scleropages formosus) genome provides new insights into the evolution of an early lineage of teleosts. Sci. Rep. 6:1–17. Nature Publishing Group.

Bohn, S., B. R. Kreiser, D. J. Daugherty, and K. A. Bodine. 2017. Natural Hybridization of Lepisosteids: Implications for Managing the Alligator Gar. North Am. J. Fish. Manag. 37:405–413. Taylor & Francis.

Bouckaert, R., J. Heled, D. Kühnert, T. Vaughan, C.-H. Wu, D. Xie, M. A. Suchard, A. Rambaut, and A. J. Drummond. 2014. BEAST 2: A Software Platform for Bayesian Evolutionary Analysis. PLoS Comput. Biol. 10:e1003537.

Bouckaert, R., T. G. Vaughan, J. Barido-Sottani, S. Duchêne, M. Fourment, A. Gavryushkina, J. Heled, G. Jones, D. Kühnert, N. D. Maio, M. Matschiner, F. K. Mendes, N. F. Müller, H. A. Ogilvie, L. du Plessis, A. Popinga, A. Rambaut, D. Rasmussen, I. Siveroni, M. A. Suchard, C.-H. Wu, D. Xie, C. Zhang, T. Stadler, and A. J. Drummond. 2019. BEAST 2.5: An advanced software platform for Bayesian evolutionary analysis. PLOS Comput. Biol. 15:e1006650. Public Library of Science.

Braasch, I., A. R. Gehrke, J. J. Smith, K. Kawasaki, T. Manousaki, J. Pasquier, A. Amores, T. Desvignes, P. Batzel, J. Catchen, A. M. Berlin, M. S. Campbell, D. Barrell, K. J. Martin, J. F. Mulley, V. Ravi, A. P. Lee, T. Nakamura, D. Chalopin, S. Fan, D. Wcisel, C. Cañestro, J. Sydes, F. E. G. Beaudry, Y. Sun, J. Hertel, M. J. Beam, M. Fasold, M. Ishiyama, J. Johnson, S. Kehr, M. Lara, J. H. Letaw, G. W. Litman, R. T. Litman, M. Mikami, T. Ota, N. R. Saha, L. Williams, P. F. Stadler, H. Wang, J. S. Taylor, Q. Fontenot, A. Ferrara, S. M. J. Searle, B. Aken, M. Yandell, I. Schneider, J. A. Yoder, J.-N. Volff, A. Meyer, C. T. Amemiya, B. Venkatesh, P. W. H. Holland, Y. Guiguen, J. Bobe, N. H. Shubin, F. Di Palma, J. Alföldi, K. Lindblad-Toh, and J. H. Postlethwait. 2016. The spotted gar genome illuminates vertebrate evolution and facilitates human-teleost comparisons. Nat. Genet. 48:427–437. Nature Publishing Group.

Brito, P. M., J. Alvarado-Ortega, and F. J. Meunier. 2017. Earliest known lepisosteoid extends the range of anatomically modern gars to the Late Jurassic. Sci. Rep. 7:17830.

Bromham, L., and M. Woolfit. 2004. Explosive Radiations and the Reliability of Molecular Clocks: Island Endemic Radiations as a Test Case. Syst. Biol. 53:758–766.

Brownstein, C. 2022. Unappreciated Cenozoic ecomorphological diversification of stem gars revealed by a large new species. Acta Palaeontol. Pol. 67.

Brownstein, C. D. 2023a. Evidence of large sturgeons in the Paleocene of North America. J. Paleontol. 97:218–222. Cambridge University Press.

Brownstein, C. D. 2023b. Syngnathoid Evolutionary History and the Conundrum of Fossil Misplacement. Integr. Org. Biol. 5:obad011.

Brownstein, C. D., R. C. Harrington, L. R. V. Alencar, D. R. Bellwood, J. H. Choat, L. A. Rocha, P. C. Wainwright, J. Tavera, E. D. Burress, M. M. Muñoz, P. F. Cowman, and T. J. Near. 2025. Phylogenomics establishes an Early Miocene reconstruction of reef vertebrate diversity. Sci. Adv. 11:eadu6149. American Association for the Advancement of Science.

Brownstein, C. D., D. Kim, O. D. Orr, G. M. Hogue, B. H. Tracy, M. W. Pugh, R. Singer, C. Myles-McBurney, J. M. Mollish, J. W. Simmons, S. R. David, G. Watkins-Colwell, E. A. Hoffman, and T. J. Near. 2022. Hidden species diversity in an iconic living fossil vertebrate. Biol. Lett. 18:20220395. Royal Society.

Brownstein, C. D., and T. R. Lyson. 2022. Giant gar from directly above the Cretaceous–Palaeogene boundary suggests healthy freshwater ecosystems existed within thousands of years of the asteroid impact. Biol. Lett. 18:20220118. Royal Society.

Brownstein, C. D., D. J. MacGuigan, D. Kim, O. Orr, L. Yang, S. R. David, B. Kreiser, and T. J. Near. 2024a. The genomic signatures of evolutionary stasis. Evolution 78:821–834.

Brownstein, C. D., and T. J. Near. 2024. Evolutionary origins of the lampriform pelagic radiation. Zool. J. Linn. Soc. 201:422–430.

Brownstein, C. D., and T. J. Near. 2026. Phylogenomics and the origins of sharks. bioRxiv.

Brownstein, C. D., T. R. Simões, M. W. Caldwell, M. S. Y. Lee, D. L. Meyer, and S. G. Scarpetta. 2023a. The affinities of the Late Triassic Cryptovaranoides and the age of crown squamates. R. Soc. Open Sci. 10:230968. Royal Society.

Brownstein, C. D., L. Yang, M. Friedman, and T. J. Near. 2023b. Phylogenomics of the Ancient and Species-Depauperate Gars Tracks 150 Million Years of Continental Fragmentation in the Northern Hemisphere. Syst. Biol. 72:213–227.

Brownstein, C. D., K. L. Zapfe, S. Lott, R. C. Harrington, A. Ghezelayagh, A. Dornburg, and T. J. Near. 2024b. Synergistic innovations enabled the radiation of anglerfishes in the deep open ocean. Curr. Biol. 34:2541–2550.e4. Elsevier.

Brusatte, S. L., M. J. Benton, G. T. Lloyd, M. Ruta, and S. C. Wang. 2010. Macroevolutionary patterns in the evolutionary radiation of archosaurs (Tetrapoda: Diapsida). Earth Environ. Sci. Trans. R. Soc. Edinb. 101:367–382.

Brusatte, S. L., J. K. O’Connor, and E. D. Jarvis. 2015. The Origin and Diversification of Birds. Curr. Biol. 25:R888–R898.

Budd, G. E., and R. P. Mann. 2023. Two Notorious Nodes: A Critical Examination of Relaxed Molecular Clock Age Estimates of the Bilaterian Animals and Placental Mammals. Syst. Biol. syad057.

Burbrink, F. T., F. G. Grazziotin, R. A. Pyron, D. Cundall, S. Donnellan, F. Irish, J. S. Keogh, F. Kraus, R. W. Murphy, B. Noonan, C. J. Raxworthy, S. Ruane, A. R. Lemmon, E. M. Lemmon, and H. Zaher. 2020. Interrogating Genomic-Scale Data for Squamata (Lizards, Snakes, and Amphisbaenians) Shows no Support for Key Traditional Morphological Relationships. Syst. Biol. 69:502–520.

Cao, Y., M. D. Sorenson, Y. Kumazawa, D. P. Mindell, and M. Hasegawa. 2000. Phylogenetic position of turtles among amniotes: evidence from mitochondrial and nuclear genes. Gene 259:139–148.

Carlisle, E., C. M. Janis, D. Pisani, P. C. J. Donoghue, and D. Silvestro. 2023. A timescale for placental mammal diversification based on Bayesian modeling of the fossil record. Curr. Biol. 33:3073–3082.e3.

Casane, D., and P. Laurenti. 2013. Why coelacanths are not ‘living fossils.’ BioEssays 35:332–338.

Cavender-Bares, J., K. H. Kozak, P. V. A. Fine, and S. W. Kembel. 2009. The merging of community ecology and phylogenetic biology. Ecol. Lett. 12:693–715.

Cavin, L., A. Piuz, C. Ferrante, and G. Guinot. 2021. Giant Mesozoic coelacanths (Osteichthyes, Actinistia) reveal high body size disparity decoupled from taxic diversity. Sci. Rep. 11:11812. Nature Publishing Group.

Chafin, T. K., M. R. Douglas, M. R. Bangs, B. T. Martin, S. M. Mussmann, and M. E. Douglas. 2021. Taxonomic Uncertainty and the Anomaly Zone: Phylogenomics Disentangle a Rapid Radiation to Resolve Contentious Species (Gila robusta Complex) in the Colorado River. Genome Biol. Evol. 13:evab200.

Chakrabarty, P., B. C. Faircloth, F. Alda, W. B. Ludt, C. D. Mcmahan, T. J. Near, A. Dornburg, J. S. Albert, J. Arroyave, M. L. J. Stiassny, L. Sorenson, and M. E. Alfaro. 2017. Phylogenomic Systematics of Ostariophysan Fishes: Ultraconserved Elements Support the Surprising Non-Monophyly of Characiformes. Syst. Biol. 66:881–895.

Chen, G., Y. Xie, and G. Zhang. 2025. Phylogenomics and comparative genomic perspective on the avian radiation. Nat. Rev. Biodivers. 1:439–460. Nature Publishing Group.

Chen, M.-Y., D. Liang, and P. Zhang. 2015. Selecting Question-Specific Genes to Reduce Incongruence in Phylogenomics: A Case Study of Jawed Vertebrate Backbone Phylogeny. Syst. Biol. 64:1104–1120.

Chiari, Y., V. Cahais, N. Galtier, and F. Delsuc. 2012. Phylogenomic analyses support the position of turtles as the sister group of birds and crocodiles (Archosauria). BMC Biol. 10:65.

Clarke, J. T., G. T. Lloyd, and M. Friedman. 2016. Little evidence for enhanced phenotypic evolution in early teleosts relative to their living fossil sister group. Proc. Natl. Acad. Sci. 113:11531–11536. Proceedings of the National Academy of Sciences.

Cole, T. L., C. Zhou, M. Fang, H. Pan, D. T. Ksepka, S. R. Fiddaman, C. A. Emerling, D. B. Thomas, X. Bi, Q. Fang, M. R. Ellegaard, S. Feng, A. L. Smith, T. A. Heath, A. J. D. Tennyson, P. G. Borboroglu, J. R. Wood, P. W. Hadden, S. Grosser, C.-A. Bost, Y. Cherel, T. Mattern, T. Hart, M.-H. S. Sinding, L. D. Shepherd, R. A. Phillips, P. Quillfeldt, J. F. Masello, J. L. Bouzat, P. G. Ryan, D. R. Thompson, U. Ellenberg, P. Dann, G. Miller, P. Dee Boersma, R. Zhao, M. T. P. Gilbert, H. Yang, D.-X. Zhang, and G. Zhang. 2022. Genomic insights into the secondary aquatic transition of penguins. Nat. Commun. 13:3912. Nature Publishing Group.

Crawford, N. G., B. C. Faircloth, J. E. McCormack, R. T. Brumfield, K. Winker, and T. C. Glenn. 2012. More than 1000 ultraconserved elements provide evidence that turtles are the sister group of archosaurs. Biol. Lett. 8:783–786.

Crouch, N. M. A., K. Ramanauskas, and B. Igić. 2019. Tip-dating and the origin of Telluraves. Mol. Phylogenet. Evol. 131:55–63.

Cui, X., M. Friedman, T. Qiao, Y. Yu, and M. Zhu. 2022. The rapid evolution of lungfish durophagy. Nat. Commun. 13:2390. Nature Publishing Group.

Darwin, C. 1859. On the Origin of Species by Means of Natural Selection, or the Preservation of Favoured Races in the Struggle for Life. John Murray, London.

Day, J. J., E. M. Steell, T. R. Vigliotta, L. A. Withey, R. Bills, J. P. Friel, M. J. Genner, and M. L. J. Stiassny. 2023. Exceptional levels of species discovery ameliorate inferences of the biogeography and diversification of an Afrotropical catfish family. Mol. Phylogenet. Evol. 182:107754.

Degnan, J. H., and N. A. Rosenberg. 2006. Discordance of Species Trees with Their Most Likely Gene Trees. PLOS Genet. 2:e68. Public Library of Science.

Degnan, J. H., and N. A. Rosenberg. 2009. Gene tree discordance, phylogenetic inference and the multispecies coalescent. Trends Ecol. Evol. 24:332–340.

Delsuc, F., H. Brinkmann, D. Chourrout, and H. Philippe. 2006. Tunicates and not cephalochordates are the closest living relatives of vertebrates. Nature 439:965–968. Nature Publishing Group.

Delsuc, F., H. Philippe, G. Tsagkogeorga, P. Simion, M.-K. Tilak, X. Turon, S. López-Legentil, J. Piette, P. Lemaire, and E. J. P. Douzery. 2018. A phylogenomic framework and timescale for comparative studies of tunicates. BMC Biol. 16:39.

Dornburg, A., M. C. Brandley, M. R. McGowen, and T. J. Near. 2012. Relaxed Clocks and Inferences of Heterogeneous Patterns of Nucleotide Substitution and Divergence Time Estimates across Whales and Dolphins (Mammalia: Cetacea). Mol. Biol. Evol. 29:721–736.

Dornburg, A., and T. J. Near. 2021. The Emerging Phylogenetic Perspective on the Evolution of Actinopterygian Fishes. Annu. Rev. Ecol. Evol. Syst. 52:427–452.

dos Reis, M., P. C. J. Donoghue, and Z. Yang. 2014. Neither phylogenomic nor palaeontological data support a Palaeogene origin of placental mammals. Biol. Lett. 10:20131003. Royal Society.

dos Reis, M., J. Inoue, M. Hasegawa, R. J. Asher, P. C. J. Donoghue, and Z. Yang. 2012. Phylogenomic datasets provide both precision and accuracy in estimating the timescale of placental mammal phylogeny. Proc. Biol. Sci. 279:3491–3500.

Drummond, A. J., S. Y. W. Ho, M. J. Phillips, and A. Rambaut. 2006. Relaxed phylogenetics and dating with confidence. PLoS Biol. 4:e88.

Du, K., M. Stöck, S. Kneitz, C. Klopp, J. M. Woltering, M. C. Adolfi, R. Feron, D. Prokopov, A. Makunin, I. Kichigin, C. Schmidt, P. Fischer, H. Kuhl, S. Wuertz, J. Gessner, W. Kloas, C. Cabau, C. Iampietro, H. Parrinello, C. Tomlinson, L. Journot, J. H. Postlethwait, I. Braasch, V. Trifonov, W. C. Warren, A. Meyer, Y. Guiguen, and M. Schartl. 2020. The sterlet sturgeon genome sequence and the mechanisms of segmental rediploidization. Nat. Ecol. Evol. 4:841–852. Nature Publishing Group.

Duchêne, D. A., A.-A. Chowdhury, J. Yang, M. Iglesias-Carrasco, J. Stiller, S. Feng, S. Bhatt, M. T. P. Gilbert, G. Zhang, J. A. Tobias, and S. Y. W. Ho. 2025. Drivers of avian genomic change revealed by evolutionary rate decomposition. Nature 1–9. Nature Publishing Group.

Dunn, C. W., G. Giribet, G. D. Edgecombe, and A. Hejnol. 2014. Animal Phylogeny and Its Evolutionary Implications*. Annu. Rev. Ecol. Evol. Syst. 45:371–395. Annual Reviews.

Dunn, C. W., A. Hejnol, D. Q. Matus, K. Pang, W. E. Browne, S. A. Smith, E. Seaver, G. W. Rouse, M. Obst, G. D. Edgecombe, M. V. Sørensen, S. H. D. Haddock, A. Schmidt-Rhaesa, A. Okusu, R. M. Kristensen, W. C. Wheeler, M. Q. Martindale, and G. Giribet. 2008. Broad phylogenomic sampling improves resolution of the animal tree of life. Nature 452:745–749. Nature Publishing Group.

Esselstyn, J. A., C. H. Oliveros, M. T. Swanson, and B. C. Faircloth. 2017. Investigating Difficult Nodes in the Placental Mammal Tree with Expanded Taxon Sampling and Thousands of Ultraconserved Elements. Genome Biol. Evol. 9:2308–2321.

Ezcurra, M. D., S. J. Nesbitt, M. Bronzati, F. M. Dalla Vecchia, F. L. Agnolin, R. B. J. Benson, F. Brissón Egli, S. F. Cabreira, S. W. Evers, A. R. Gentil, R. B. Irmis, A. G. Martinelli, F. E. Novas,. 2020. Enigmatic dinosaur precursors bridge the gap to the origin of Pterosauria. Nature 588:445–449. Nature Publishing Group.

Faircloth, B. C. 2016. PHYLUCE is a software package for the analysis of conserved genomic loci. Bioinforma. Oxf. Engl. 32:786–788.

Feng, Y.-J., D. C. Blackburn, D. Liang, D. M. Hillis, D. B. Wake, D. C. Cannatella, and P. Zhang. 2017. Phylogenomics reveals rapid, simultaneous diversification of three major clades of Gondwanan frogs at the Cretaceous–Paleogene boundary. Proc. Natl. Acad. Sci. 114:E5864–E5870. Proceedings of the National Academy of Sciences.

Field, D. J., J. S. Berv, A. Y. Hsiang, R. Lanfear, M. J. Landis, and A. Dornburg. 2019. Timing the extant avian radiation: the rise of modern birds, and the importance of modeling molecular rate variation. PeerJ Preprints.

Foley, N. M., V. C. Mason, A. J. Harris, K. R. Bredemeyer, J. Damas, H. A. Lewin, E. Eizirik, J. Gatesy, E. K. Karlsson, K. Lindblad-Toh, Zoonomia Consortium, M. S. Springer, and W. J. Murphy. 2023. A genomic timescale for placental mammal evolution. Science 380:eabl8189. American Association for the Advancement of Science.

Foley, N. M., M. S. Springer, and E. C. Teeling. 2016. Mammal madness: is the mammal tree of life not yet resolved? Philos. Trans. R. Soc. B Biol. Sci. 371:20150140. Royal Society.

Friedman, M. 2010. Explosive morphological diversification of spiny-finned teleost fishes in the aftermath of the end-Cretaceous extinction. Proc. R. Soc. B Biol. Sci. 277:1675–1683. Royal Society.

Friedman, M. 2022. The Macroevolutionary History of Bony Fishes: A Paleontological View. Annu. Rev. Ecol. Evol. Syst. 53:353–377.

Friedman, M., K. L. Feilich, H. T. Beckett, M. E. Alfaro, B. C. Faircloth, D. Černý, M. Miya, T. J. Near, and R. C. Harrington. 2019. A phylogenomic framework for pelagiarian fishes (Acanthomorpha: Percomorpha) highlights mosaic radiation in the open ocean. Proc. R. Soc. B Biol. Sci. 286:20191502. Royal Society.

Garland, T., Jr, A. F. Bennett, and E. L. Rezende. 2005. Phylogenetic approaches in comparative physiology. J. Exp. Biol. 208:3015–3035.

Gavrilets, S., and J. B. Losos. 2009. Adaptive Radiation: Contrasting Theory with Data. Science 323:732–737. American Association for the Advancement of Science.

Ghezelayagh, A., R. C. Harrington, E. D. Burress, M. A. Campbell, J. C. Buckner, P. Chakrabarty, J. R. Glass, W. T. McCraney, P. J. Unmack, C. E. Thacker, M. E. Alfaro, S. T. Friedman, W. B. Ludt, P. F. Cowman, M. Friedman, S. A. Price, A. Dornburg, B. C. Faircloth, P. C. Wainwright, and T. J. Near. 2022. Prolonged morphological expansion of spiny-rayed fishes following the end-Cretaceous. Nat. Ecol. Evol. 6:1211–1220. Nature Publishing Group.

Giles, S., G.-H. Xu, T. J. Near, and M. Friedman. 2017. Early members of ‘living fossil’ lineage imply later origin of modern ray-finned fishes. Nature 549:265–268. Nature Publishing Group.

Giribet, G., and G. D. Edgecombe. 2019. The Phylogeny and Evolutionary History of Arthropods. Curr. Biol. CB 29:R592–R602.

Glass, J. R., R. C. Harrington, P. F. Cowman, B. C. Faircloth, and T. J. Near. 2023. Widespread sympatry in a species-rich clade of marine fishes (Carangoidei). Proc. R. Soc. B Biol. Sci. 290:20230657. Royal Society.

Gradstein, F. M., J. G. Ogg, M. Schmitz, and G. Ogg. 2021. The Geologic Time Scale 2020. Elsevier Science, Amsterdam, The Netherlands.

Grande, L. 2010. An Empirical Synthetic Pattern Study of Gars (lepisosteiformes) and Closely Related Species, Based Mostly on Skeletal Anatomy. the Resurrection of Holostei. Copeia 2010:iii–871. [American Society of Ichthyologists and Herpetologists (ASIH), Allen Press].

Grande, L., and W. E. Bemis. 1998. A Comprehensive Phylogenetic Study of Amiid Fishes (Amiidae) Based on Comparative Skeletal Anatomy. an Empirical Search for Interconnected Patterns of Natural History. J. Vertebr. Paleontol. 18:1–696. Taylor & Francis.

Grande, L., and W. E. Bemis. 1991. Osteology and Phylogenetic Relationships of Fossil and Recent Paddlefishes (Polyodontidae) with Comments on the Interrelationships of Acipenseriformes. J. Vertebr. Paleontol. 11:1–121. Taylor & Francis.

Grande, L., F. Jin, Y. Yabumoto, and W. E. Bemis. 2002. Protopsephurus liui, a well-preserved primitive paddlefish (Acipenseriformes: Polyodontidae) from the Lower Cretaceous of China. J. Vertebr. Paleontol. 22:209–237. Taylor & Francis.

Hackett, S. J., R. T. Kimball, S. Reddy, R. C. K. Bowie, E. L. Braun, M. J. Braun, J. L. Chojnowski, W. A. Cox, K.-L. Han, J. Harshman, C. J. Huddleston, B. D. Marks, K. J. Miglia, W. S. Moore, F. H. Sheldon, D. W. Steadman, C. C. Witt, and T. Yuri. 2008. A Phylogenomic Study of Birds Reveals Their Evolutionary History. Science 320:1763–1768. American Association for the Advancement of Science.

Hao, S., K. Han, L. Meng, X. Huang, W. Cao, C. Shi, M. Zhang, Y. Wang, Q. Liu, Y. Zhang, H. Sun, I. Seim, X. Xu, X. Liu, and G. Fan. 2020. African Arowana Genome Provides Insights on Ancient Teleost Evolution. iScience 23:101662.

Havelka, M., V. Kašpar, M. Hulák, and M. Flajšhans. 2011. Sturgeon genetics and cytogenetics: a review related to ploidy levels and interspecific hybridization. Folia Zool. 60:93–103. Institute of Vertebrate Biology, Czech Academy of Sciences.

Hilton, E. J., M. A. D. During, L. Grande, and P. E. Ahlberg. 2023. New paddlefishes (Acipenseriformes, Polyodontidae) from the Late Cretaceous Tanis Site of the Hell Creek Formation in North Dakota, USA. J. Paleontol. 97:675–692.

Hilton, E. J., and L. Grande. 2023. Late Cretaceous sturgeons (Acipenseridae) from North America, with two new species from the Tanis site in the Hell Creek Formation of North Dakota. J. Paleontol. 97:189–217. Cambridge University Press.

Hilton, E. J., L. Grande, and W. E. Bemis. 2011. Skeletal Anatomy of the Shortnose Sturgeon, Acipenser brevirostrum Lesueur, 1818, and the Systematics of Sturgeons (Acipenseriformes, Acipenseridae). Fieldiana Life Earth Sci. 2011:1–168. Field Museum of Natural History.

Hime, P. M., A. R. Lemmon, E. C. M. Lemmon, E. Prendini, J. M. Brown, R. C. Thomson, J. D. Kratovil, B. P. Noonan, R. A. Pyron, P. L. V. Peloso, M. L. Kortyna, J. S. Keogh, S. C. Donnellan, R. L. Mueller, C. J. Raxworthy, K. Kunte, S. R. Ron, S. Das, N. Gaitonde, D. M. Green, J. Labisko, J. Che, and D. W. Weisrock. 2020. Phylogenomics Reveals Ancient Gene Tree Discordance in the Amphibian Tree of Life. Syst. Biol. 70:49–66.

Hughes, L. C., G. Ortí, Y. Huang, Y. Sun, C. C. Baldwin, A. W. Thompson, D. Arcila, R. Betancur-R, C. Li, L. Becker, N. Bellora, X. Zhao, X. Li, M. Wang, C. Fang, B. Xie, Z. Zhou, H. Huang, S. Chen, B. Venkatesh, and Q. Shi. 2018. Comprehensive phylogeny of ray-finned fishes (Actinopterygii) based on transcriptomic and genomic data. Proc. Natl. Acad. Sci. U. S. A. 115:6249–6254.

Hughes, L. C., G. Ortí, H. Saad, C. Li, W. T. White, C. C. Baldwin, K. A. Crandall, D. Arcila, and R. Betancur-R. 2021. Exon probe sets and bioinformatics pipelines for all levels of fish phylogenomics. Mol. Ecol. Resour. 21:816–833.

Hurley, I. A., R. L. Mueller, K. A. Dunn, E. J. Schmidt, M. Friedman, R. K. Ho, V. E. Prince, Z. Yang, M. G. Thomas, and M. I. Coates. 2006. A new time-scale for ray-finned fish evolution. Proc. R. Soc. B Biol. Sci. 274:489–498. Royal Society.

Irisarri, I., D. Baurain, H. Brinkmann, F. Delsuc, J.-Y. Sire, A. Kupfer, J. Petersen, M. Jarek, A. Meyer, M. Vences, and H. Philippe. 2017. Phylotranscriptomic consolidation of the jawed vertebrate timetree. Nat. Ecol. Evol. 1:1370–1378. Nature Publishing Group.

Irisarri, I., and A. Meyer. 2016. The Identification of the Closest Living Relative(s) of Tetrapods: Phylogenomic Lessons for Resolving Short Ancient Internodes. Syst. Biol. 65:1057–1075.

Jablonski, D. 2001. Lessons from the past: Evolutionary impacts of mass extinctions. Proc. Natl. Acad. Sci. 98:5393–5398. Proceedings of the National Academy of Sciences.

Jablonski, D. 2005. Mass extinctions and macroevolution. Paleobiology 31:192–210.

Jarvis, E. D., S. Mirarab, A. J. Aberer, B. Li, P. Houde, C. Li, S. Y. W. Ho, B. C. Faircloth, B. Nabholz, J. T. Howard, A. Suh, C. C. Weber, R. R. da Fonseca, J. Li, F. Zhang, H. Li, L. Zhou, N. Narula, L. Liu, G. Ganapathy, B. Boussau, Md. S. Bayzid, V. Zavidovych, S. Subramanian, T. Gabaldón, S. Capella-Gutiérrez, J. Huerta-Cepas, B. Rekepalli, K. Munch, M. Schierup, B. Lindow, W. C. Warren, D. Ray, R. E. Green, M. W. Bruford, X. Zhan, A. Dixon, S. Li, N. Li, Y. Huang, E. P. Derryberry, M. F. Bertelsen, F. H. Sheldon, R. T. Brumfield, C. V. Mello, P. V. Lovell, M. Wirthlin, M. P. C. Schneider, F. Prosdocimi, J. A. Samaniego, A. M. V. Velazquez, A. Alfaro-Núñez, P. F. Campos, B. Petersen, T. Sicheritz-Ponten, A. Pas, T. Bailey, P. Scofield, M. Bunce, D. M. Lambert, Q. Zhou, P. Perelman, A. C. Driskell, B. Shapiro, Z. Xiong, Y. Zeng, S. Liu, Z. Li, B. Liu, K. Wu, J. Xiao, X. Yinqi, Q. Zheng, Y. Zhang, H. Yang, J. Wang, L. Smeds, F. E. Rheindt, M. Braun, J. Fjeldsa, L. Orlando, F. K. Barker, K. A. Jønsson, W. Johnson, K.-P. Koepfli, S. O’Brien, D. Haussler, O. A. Ryder, C. Rahbek, E. Willerslev, G. R. Graves, T. C. Glenn, J. McCormack, D. Burt, H. Ellegren, P. Alström, S. V. Edwards, A. Stamatakis, D. P. Mindell, J. Cracraft, E. L. Braun, T. Warnow, W. Jun, M. T. P. Gilbert, and G. Zhang. 2014. Whole-genome analyses resolve early branches in the tree of life of modern birds. Science 346:1320–1331. American Association for the Advancement of Science.

Jetz, W., G. H. Thomas, J. B. Joy, K. Hartmann, and A. O. Mooers. 2012. The global diversity of birds in space and time. Nature 491:444–448. Nature Publishing Group.

Jones, M. E., C. L. Anderson, C. A. Hipsley, J. Müller, S. E. Evans, and R. R. Schoch. 2013. Integration of molecules and new fossils supports a Triassic origin for Lepidosauria (lizards, snakes, and tuatara). BMC Evol. Biol. 13:208.

Káldy, J., A. Mozsár, G. Fazekas, M. Farkas, D. L. Fazekas, G. L. Fazekas, K. Goda, Z. Gyöngy, B. Kovács, K. Semmens, M. Bercsényi, M. Molnár, and E. Patakiné Várkonyi. 2020. Hybridization of Russian Sturgeon (Acipenser gueldenstaedtii, Brandt and Ratzeberg, 1833) and American Paddlefish (Polyodon spathula, Walbaum 1792) and Evaluation of Their Progeny. Genes 11:753.

Kalyaanamoorthy, S., B. Q. Minh, T. K. F. Wong, A. von Haeseler, and L. S. Jermiin. 2017. ModelFinder: fast model selection for accurate phylogenetic estimates. Nat. Methods 14:587–589. Nature Publishing Group.

Kappas, I., S. Vittas, C. N. Pantzartzi, E. Drosopoulou, and Z. G. Scouras. 2016. A Time-Calibrated Mitogenome Phylogeny of Catfish (Teleostei: Siluriformes). PLOS ONE 11:e0166988. Public Library of Science.

Katoh, K., and D. M. Standley. 2013. MAFFT multiple sequence alignment software version 7: improvements in performance and usability. Mol. Biol. Evol. 30:772–780.

Kear, B. P., V. S. Engelschiøn, Ø. Hammer, A. J. Roberts, and J. H. Hurum. 2023. Earliest Triassic ichthyosaur fossils push back oceanic reptile origins. Curr. Biol. 33:R178–R179. Elsevier.

Kimball, R. T., C. H. Oliveros, N. Wang, N. D. White, F. K. Barker, D. J. Field, D. T. Ksepka, R. T. Chesser, R. G. Moyle, M. J. Braun, R. T. Brumfield, B. C. Faircloth, B. T. Smith, and E. L. Braun. 2019. A Phylogenomic Supertree of Birds. Diversity 11:109. Multidisciplinary Digital Publishing Institute.

Klein, C. G., D. Pisani, D. J. Field, R. Lakin, M. A. Wills, and N. R. Longrich. 2021. Evolution and dispersal of snakes across the Cretaceous-Paleogene mass extinction. Nat. Commun. 12:5335. Nature Publishing Group.

Kligman, B. T., B. M. Gee, A. D. Marsh, S. J. Nesbitt, M. E. Smith, W. G. Parker, and M. R. Stocker. 2023. Triassic stem caecilian supports dissorophoid origin of living amphibians. Nature 614:102–107. Nature Publishing Group.

Ksepka, D. T., J. F. Parham, J. F. Allman, M. J. Benton, M. T. Carrano, K. A. Cranston, P. C. J. Donoghue, J. J. Head, E. J. Hermsen, R. B. Irmis, W. G. Joyce, M. Kohli, K. D. Lamm, D. Leehr, J. L. Patané, P. D. Polly, M. J. Phillips, N. A. Smith, N. D. Smith, M. Van Tuinen, J. L. Ware, and R. C. M. Warnock. 2015. The Fossil Calibration Database—A New Resource for Divergence Dating. Syst. Biol. 64:853–859.

Kück, P., J. Romahn, and K. Meusemann. 2022. Pitfalls of the site-concordance factor (sCF) as measure of phylogenetic branch support. NAR Genomics Bioinforma. 4:lqac064.

Kuhl, H., C. Frankl-Vilches, A. Bakker, G. Mayr, G. Nikolaus, S. T. Boerno, S. Klages, B. Timmermann, and M. Gahr. 2021. An Unbiased Molecular Approach Using 3ʹ-UTRs Resolves the Avian Family-Level Tree of Life. Mol. Biol. Evol. 38:108–127.

Lepage, T., D. Bryant, H. Philippe, and N. Lartillot. 2007. A General Comparison of Relaxed Molecular Clock Models. Mol. Biol. Evol. 24:2669–2680.

Lidgard, S., and A. C. Love. 2018. Rethinking Living Fossils. BioScience 68:760–770.

Linkem, C. W., V. N. Minin, and A. D. Leaché. 2016. Detecting the Anomaly Zone in Species Trees and Evidence for a Misleading Signal in Higher-Level Skink Phylogeny (Squamata: Scincidae). Syst. Biol. 65:465–477.

Liu, L., and S. V. Edwards. 2009. Phylogenetic Analysis in the Anomaly Zone. Syst. Biol. 58:452–460.

Lloyd, G. T., K. E. Davis, D. Pisani, J. E. Tarver, M. Ruta, M. Sakamoto, D. W. E. Hone, R. Jennings, and M. J. Benton. 2008. Dinosaurs and the Cretaceous Terrestrial Revolution. Proc. R. Soc. B Biol. Sci. 275:2483–2490. Royal Society.

Magallón, S., and M. J. Sanderson. 2001. Absolute diversification rates in angiosperm clades. Evol. Int. J. Org. Evol. 55:1762–1780.

Marjanović, D., and M. Laurin. 2007. Fossils, Molecules, Divergence Times, and the Origin of Lissamphibians. Syst. Biol. 56:369–388.

Marlétaz, F., N. Timoshevskaya, V. A. Timoshevskiy, E. Parey, O. Simakov, D. Gavriouchkina, M. Suzuki, K. Kubokawa, S. Brenner, J. J. Smith, and D. S. Rokhsar. 2024. The hagfish genome and the evolution of vertebrates. Nature 627:811–820. Nature Publishing Group.

Melo, B. F., B. L. Sidlauskas, T. J. Near, F. F. Roxo, A. Ghezelayagh, L. E. Ochoa, M. L. J. Stiassny, J. Arroyave, J. Chang, B. C. Faircloth, D. J. MacGuigan, R. C. Harrington, R. C. Benine, M. D. Burns, K. Hoekzema, N. C. Sanches, J. A. Maldonado-Ocampo, R. M. C. Castro, F. Foresti, M. E. Alfaro, and C. Oliveira. 2022. Accelerated Diversification Explains the Exceptional Species Richness of Tropical Characoid Fishes. Syst. Biol. 71:78–92.

Meredith, R. W., J. E. Janečka, J. Gatesy, O. A. Ryder, C. A. Fisher, E. C. Teeling, A. Goodbla, E. Eizirik, T. L. L. Simão, T. Stadler, D. L. Rabosky, R. L. Honeycutt, J. J. Flynn, C. M. Ingram, C. Steiner, T. L. Williams, T. J. Robinson, A. Burk-Herrick, M. Westerman, N. A. Ayoub, M. S. Springer, and W. J. Murphy. 2011. Impacts of the Cretaceous Terrestrial Revolution and KPg extinction on mammal diversification. Science 334:521–524.

Meyer, A., S. Schloissnig, P. Franchini, K. Du, J. M. Woltering, I. Irisarri, W. Y. Wong, S. Nowoshilow, S. Kneitz, A. Kawaguchi, A. Fabrizius, P. Xiong, C. Dechaud, H. P. Spaink, J.-N. Volff, O. Simakov, T. Burmester, E. M. Tanaka, and M. Schartl. 2021. Giant lungfish genome elucidates the conquest of land by vertebrates. Nature 590:284–289. Nature Publishing Group.

Miller, E. C., R. Faucher, P. B. Hart, M. Rincón-Sandoval, A. Santaquiteria, W. T. White, C. C. Baldwin, M. Miya, R. Betancur-R, L. Tornabene, K. Evans, and D. Arcila. 2024. Reduced evolutionary constraint accompanies ongoing radiation in deep-sea anglerfishes. Nat. Ecol. Evol., doi: 10.1038/s41559-024-02586-3.

Minh, B. Q., M. W. Hahn, and R. Lanfear. 2020a. New Methods to Calculate Concordance Factors for Phylogenomic Datasets. Mol. Biol. Evol. 37:2727–2733.

Minh, B. Q., H. A. Schmidt, O. Chernomor, D. Schrempf, M. D. Woodhams, A. von Haeseler, and R. Lanfear. 2020b. IQ-TREE 2: New Models and Efficient Methods for Phylogenetic Inference in the Genomic Era. Mol. Biol. Evol. 37:1530–1534.

Mirarab, S., I. Rivas-González, S. Feng, J. Stiller, Q. Fang, U. Mai, G. Hickey, G. Chen, N. Brajuka, O. Fedrigo, G. Formenti, J. B. W. Wolf, K. Howe, A. Antunes, M. H. Schierup, B. Paten, E. D. Jarvis, G. Zhang, and E. L. Braun. 2024. A region of suppressed recombination misleads neoavian phylogenomics. Proc. Natl. Acad. Sci. 121:e2319506121. Proceedings of the National Academy of Sciences.

Mongiardino Koch, N., J. R. Thompson, A. S. Hiley, M. F. McCowin, A. F. Armstrong, S. E. Coppard, F. Aguilera, O. Bronstein, A. Kroh, R. Mooi, and G. W. Rouse. 2022. Phylogenomic analyses of echinoid diversification prompt a re-evaluation of their fossil record. eLife 11:e72460. eLife Sciences Publications, Ltd.

Mongiardino Koch, N., E. Tilic, A. K. Miller, J. Stiller, and G. W. Rouse. 2023. Confusion will be my epitaph: genome-scale discordance stifles phylogenetic resolution of Holothuroidea. Proc. R. Soc. B Biol. Sci. 290:20230988. Royal Society.

Morris, S. C., and J.-B. Caron. 2014. A primitive fish from the Cambrian of North America. Nature 512:419–422. Nature Publishing Group.

Morris, S. C., and J.-B. Caron. 2012. Pikaia gracilens Walcott, a stem-group chordate from the Middle Cambrian of British Columbia. Biol. Rev. 87:480–512.

Murray, A. M., D. B. Brinkman, D. G. DeMar, and G. P. Wilson. 2020. Paddlefish and sturgeon (Chondrostei: Acipenseriformes: Polyodontidae and Acipenseridae) from lower Paleocene deposits of Montana, U.S.A. J. Vertebr. Paleontol. 40:e1775091. Taylor & Francis.

Murray, A. M., L. E. Nelson, and D. B. Brinkman. 2023. A new sturgeon from the Upper Cretaceous Horseshoe Canyon Formation in central Alberta, Canada. J. Vertebr. Paleontol. 0:e2232846. Taylor & Francis.

Mussini, G., M. P. Smith, J. Vinther, I. A. Rahman, D. J. E. Murdock, D. A. T. Harper, and F. S. Dunn. 2024. A new interpretation of *Pikaia* reveals the origins of the chordate body plan. Curr. Biol. 34:2980–2989.e2.

Near, T. J., A. Dornburg, R. I. Eytan, B. P. Keck, W. L. Smith, K. L. Kuhn, J. A. Moore, S. A. Price, F. T. Burbrink, M. Friedman, and P. C. Wainwright. 2013. Phylogeny and tempo of diversification in the superradiation of spiny-rayed fishes. Proc. Natl. Acad. Sci. 110:12738–12743. Proceedings of the National Academy of Sciences.

Near, T. J., R. I. Eytan, A. Dornburg, K. L. Kuhn, J. A. Moore, M. P. Davis, P. C. Wainwright, M. Friedman, and W. L. Smith. 2012. Resolution of ray-finned fish phylogeny and timing of diversification. Proc. Natl. Acad. Sci. 109:13698–13703.

Near, T. J., and C. E. Thacker. 2024. Phylogenetic classification of living and fossil ray-finned fishes (Actinopterygii). Bull. Peabody Mus. Nat. Hist. 65:3–302.

Nesbitt, S. J. 2011. The Early Evolution of Archosaurs: Relationships and the Origin of Major Clades. Bull. Am. Mus. Nat. Hist. 2011:1–292. American Museum of Natural History.

Nguyen, L.-T., H. A. Schmidt, A. von Haeseler, and B. Q. Minh. 2015. IQ-TREE: A Fast and Effective Stochastic Algorithm for Estimating Maximum-Likelihood Phylogenies. Mol. Biol. Evol. 32:268–274.

O’Leary, M. A., J. I. Bloch, J. J. Flynn, T. J. Gaudin, A. Giallombardo, N. P. Giannini, S. L. Goldberg, B. P. Kraatz, Z.-X. Luo, J. Meng, X. Ni, M. J. Novacek, F. A. Perini, Z. S. Randall, G. W. Rougier, E. J. Sargis, M. T. Silcox, N. B. Simmons, M. Spaulding, P. M. Velazco, M. Weksler, J. R. Wible, and A. L. Cirranello. 2013. The placental mammal ancestor and the post-K-Pg radiation of placentals. Science 339:662–667.

Parey, E., A. Louis, J. Montfort, O. Bouchez, C. Roques, C. Iampietro, J. Lluch, A. Castinel, C. Donnadieu, T. Desvignes, C. Floi Bucao, E. Jouanno, M. Wen, S. Mejri, R. Dirks, H. Jansen, C. Henkel, W.-J. Chen, M. Zahm, C. Cabau, C. Klopp, A. W. Thompson, M. Robinson-Rechavi, I. Braasch, G. Lecointre, J. Bobe, J. H. Postlethwait, C. Berthelot, H. Roest Crollius, and Y. Guiguen. 2023. Genome structures resolve the early diversification of teleost fishes. Science 379:572–575. American Association for the Advancement of Science.

Parham, J. F., P. C. J. Donoghue, C. J. Bell, T. D. Calway, J. J. Head, P. A. Holroyd, J. G. Inoue, R. B. Irmis, W. G. Joyce, D. T. Ksepka, J. S. L. Patané, N. D. Smith, J. E. Tarver, M. van Tuinen, Z. Yang, K. D. Angielczyk, J. M. Greenwood, C. A. Hipsley, L. Jacobs, P. J. Makovicky, J. Müller, K. T. Smith, J. M. Theodor, R. C. M. Warnock, and M. J. Benton. 2012. Best Practices for Justifying Fossil Calibrations. Syst. Biol. 61:346–359.

Phillips, M. J., and C. Fruciano. 2018. The soft explosive model of placental mammal evolution. BMC Evol. Biol. 18:104.

Prum, R. O., J. S. Berv, A. Dornburg, D. J. Field, J. P. Townsend, E. M. Lemmon, and A. R. Lemmon. 2015. A comprehensive phylogeny of birds (Aves) using targeted next-generation DNA sequencing. Nature 526:569–573. Nature Publishing Group.

Pyron, R. A., F. T. Burbrink, and J. J. Wiens. 2013. A phylogeny and revised classification of Squamata, including 4161 species of lizards and snakes. BMC Evol. Biol. 13:93.

Rambaut, A., A. J. Drummond, D. Xie, G. Baele, and M. A. Suchard. 2018. Posterior Summarization in Bayesian Phylogenetics Using Tracer 1.7. Syst. Biol. 67:901–904.

Reddy, S., R. T. Kimball, A. Pandey, P. A. Hosner, M. J. Braun, S. J. Hackett, K.-L. Han, J. Harshman, C. J. Huddleston, S. Kingston, B. D. Marks, K. J. Miglia, W. S. Moore, F. H. Sheldon, C. C. Witt, T. Yuri, and E. L. Braun. 2017. Why Do Phylogenomic Data Sets Yield Conflicting Trees? Data Type Influences the Avian Tree of Life more than Taxon Sampling. Syst. Biol. 66:857–879.

Ribeiro, E., A. M. Davis, R. A. Rivero-Vega, G. Ortí, and R. Betancur-R. 2018. Post-Cretaceous bursts of evolution along the benthic-pelagic axis in marine fishes. Proc. R. Soc. B Biol. Sci. 285:20182010. Royal Society.

Robertson, D. S., W. M. Lewis, P. M. Sheehan, and O. B. Toon. 2013. K-Pg extinction patterns in marine and freshwater environments: The impact winter model. J. Geophys. Res. Biogeosciences 118:1006–1014.

Roxo, F. F., L. E. Ochoa, M. H. Sabaj, N. K. Lujan, R. Covain, G. S. C. Silva, B. F. Melo, J. S. Albert, J. Chang, F. Foresti, M. E. Alfaro, and C. Oliveira. 2019. Phylogenomic reappraisal of the Neotropical catfish family Loricariidae (Teleostei: Siluriformes) using ultraconserved elements. Mol. Phylogenet. Evol. 135:148–165.

Sander, P. M., E. M. Griebeler, N. Klein, J. V. Juarbe, T. Wintrich, L. J. Revell, and L. Schmitz. 2021. Early giant reveals faster evolution of large body size in ichthyosaurs than in cetaceans. Science 374:eabf5787. American Association for the Advancement of Science.

Sanderson, M. J. 2002. Estimating absolute rates of molecular evolution and divergence times: a penalized likelihood approach. Mol. Biol. Evol. 19:101–109.

Sato, H., A. M. Murray, O. Vernygora, and P. J. Currie. 2018. A rare, articulated sturgeon (Chondrostei: Acipenseriformes) from the Upper Cretaceous of Dinosaur Provincial Park, Alberta, Canada. J. Vertebr. Paleontol. 38:(1)-(15). Taylor & Francis.

Schartl, M., J. M. Woltering, I. Irisarri, K. Du, S. Kneitz, M. Pippel, T. Brown, P. Franchini, J. Li, M. Li, M. Adolfi, S. Winkler, J. de Freitas Sousa, Z. Chen, S. Jacinto, E. Z. Kvon, L. R. Correa de Oliveira, E. Monteiro, D. Baia Amaral, T. Burmester, D. Chalopin, A. Suh, E. Myers, O. Simakov, I. Schneider, and A. Meyer. 2024. The genomes of all lungfish inform on genome expansion and tetrapod evolution. Nature 1–8. Nature Publishing Group.

Schultz, D. T., S. H. D. Haddock, J. V. Bredeson, R. E. Green, O. Simakov, and D. S. Rokhsar. 2023. Ancient gene linkages support ctenophores as sister to other animals. Nature 618:110–117. Nature Publishing Group.

Schultze, H.-P., and E. O. Wiley. 1984. The Neopterygian Amia as a Living Fossil. Pp. 153–159 in N. Eldredge and S. M. Stanley, eds. Living Fossils. Springer, New York, NY.

Shan, Y., and R. Gras. 2011. 43 genes support the lungfish-coelacanth grouping related to the closest living relative of tetrapods with the Bayesian method under the coalescence model. BMC Res. Notes 4:49.

Sheehan, P. M., and D. E. Fastovsky. 1992. Major extinctions of land-dwelling vertebrates at the Cretaceous-Tertiary boundary, eastern Montana. Geology 20:556–560.

Simakov, O., J. Bredeson, K. Berkoff, F. Marletaz, T. Mitros, D. T. Schultz, B. L. O’Connell, P. Dear, D. E. Martinez, R. E. Steele, R. E. Green, C. N. David, and D. S. Rokhsar. 2022. Deeply conserved synteny and the evolution of metazoan chromosomes. Sci. Adv. 8:eabi5884. American Association for the Advancement of Science.

Simões, T. R., M. W. Caldwell, M. Tałanda, M. Bernardi, A. Palci, O. Vernygora, F. Bernardini, L. Mancini, and R. L. Nydam. 2018. The origin of squamates revealed by a Middle Triassic lizard from the Italian Alps. Nature 557:706–709. Nature Publishing Group.

Simões, T. R., C. F. Kammerer, M. W. Caldwell, and S. E. Pierce. 2022. Successive climate crises in the deep past drove the early evolution and radiation of reptiles. Sci. Adv. 8:eabq1898. American Association for the Advancement of Science.

Simões, T. R., and R. A. Pyron. 2021. The Squamate Tree of Life. Bull. Mus. Comp. Zool. 163:47–95. Museum of Comparative Zoology, Harvard University.

Singhal, S., T. J. Colston, M. R. Grundler, S. A. Smith, G. C. Costa, G. R. Colli, C. Moritz, R. A. Pyron, and D. L. Rabosky. 2021. Congruence and Conflict in the Higher-Level Phylogenetics of Squamate Reptiles: An Expanded Phylogenomic Perspective. Syst. Biol. 70:542–557.

Smith, J. J., S. Kuraku, C. Holt, T. Sauka-Spengler, N. Jiang, M. S. Campbell, M. D. Yandell, T. Manousaki, A. Meyer, O. E. Bloom, J. R. Morgan, J. D. Buxbaum, R. Sachidanandam, C. Sims, A. S. Garruss, M. Cook, R. Krumlauf, L. M. Wiedemann, S. A. Sower, W. A. Decatur, J. A. Hall, C. T. Amemiya, N. R. Saha, K. M. Buckley, J. P. Rast, S. Das, M. Hirano, N. McCurley, P. Guo, N. Rohner, C. J. Tabin, P. Piccinelli, G. Elgar, M. Ruffier, B. L. Aken, S. M. J. Searle, M. Muffato, M. Pignatelli, J. Herrero, M. Jones, C. T. Brown, Y.-W. Chung-Davidson, K. G. Nanlohy, S. V. Libants, C.-Y. Yeh, D. W. McCauley, J. A. Langeland, Z. Pancer, B. Fritzsch, P. J. de Jong, B. Zhu, L. L. Fulton, B. Theising, P. Flicek, M. E. Bronner, W. C. Warren, S. W. Clifton, R. K. Wilson, and W. Li. 2013. Sequencing of the sea lamprey (Petromyzon marinus) genome provides insights into vertebrate evolution. Nat. Genet. 45:415–421. Nature Publishing Group.

Smith, S. D., M. W. Pennell, C. W. Dunn, and S. V. Edwards. 2020. Phylogenetics is the New Genetics (for Most of Biodiversity). Trends Ecol. Evol. 35:415–425. Elsevier.

Somogyi, S., J. Y. J. Lee, A. Chen, W. Liang, G. S. Bever, and D. J. Field. 2026. Hemiplasy helps explain high rates of apparent morphological convergence in neoavian birds. Curr. Biol. 36:2424–2433.e3. Elsevier.

Song, S., L. Liu, S. V. Edwards, and S. Wu. 2012. Resolving conflict in eutherian mammal phylogeny using phylogenomics and the multispecies coalescent model. Proc. Natl. Acad. Sci. 109:14942–14947. Proceedings of the National Academy of Sciences.

Springer, M. S., W. J. Murphy, E. Eizirik, and S. J. O’Brien. 2003. Placental mammal diversification and the Cretaceous–Tertiary boundary. Proc. Natl. Acad. Sci. 100:1056–1061. Proceedings of the National Academy of Sciences.

Stanley, S. M. 1975. A theory of evolution above the species level. Proc. Natl. Acad. Sci. 72:646–650. Proceedings of the National Academy of Sciences.

Stiller, J., S. Feng, A.-A. Chowdhury, I. Rivas-González, D. A. Duchêne, Q. Fang, Y. Deng, A. Kozlov, A. Stamatakis, S. Claramunt, J. M. T. Nguyen, S. Y. W. Ho, B. C. Faircloth, J. Haag, P. Houde, J. Cracraft, M. Balaban, U. Mai, G. Chen, R. Gao, C. Zhou, Y. Xie, Z. Huang, Z. Cao, Z. Yan, H. A. Ogilvie, L. Nakhleh, B. Lindow, B. Morel, J. Fjeldså, P. A. Hosner, R. R. da Fonseca, B. Petersen, J. A. Tobias, T. Székely, J. D. Kennedy, A. H. Reeve, A. Liker, M. Stervander, A. Antunes, D. T. Tietze, M. Bertelsen, F. Lei, C. Rahbek, G. R. Graves, M. H. Schierup, T. Warnow, E. L. Braun, M. T. P. Gilbert, E. D. Jarvis, S. Mirarab, and G. Zhang. 2024. Complexity of avian evolution revealed by family-level genomes. Nature 1–3. Nature Publishing Group.

Streicher, J. W., and J. J. Wiens. 2017. Phylogenomic analyses of more than 4000 nuclear loci resolve the origin of snakes among lizard families. Biol. Lett. 13:20170393. Royal Society.

Stroud, J. T., and J. B. Losos. 2016. Ecological Opportunity and Adaptive Radiation. Annu. Rev. Ecol. Evol. Syst. 47:507–532.

Su, Z.-H., A. Sasaki, H. Minami, and K. Ozaki. 2024. Arthropod Phylotranscriptomics With a Special Focus on the Basal Phylogeny of the Myriapoda. Genome Biol. Evol. 16:evae189.

Suh, A., L. Smeds, and H. Ellegren. 2015. The Dynamics of Incomplete Lineage Sorting across the Ancient Adaptive Radiation of Neoavian Birds. PLOS Biol. 13:e1002224. Public Library of Science.

Takezaki, N. 2018. Global Rate Variation in Bony Vertebrates. Genome Biol. Evol. 10:1803–1815.

Takezaki, N. 2021. Resolving the Early Divergence Pattern of Teleost Fish Using Genome-Scale Data. Genome Biol. Evol. 13:evab052.

Tarver, J. E., M. dos Reis, S. Mirarab, R. J. Moran, S. Parker, J. E. O’Reilly, B. L. King, M. J. O’Connell, R. J. Asher, T. Warnow, K. J. Peterson, P. C. J. Donoghue, and D. Pisani. 2016. The Interrelationships of Placental Mammals and the Limits of Phylogenetic Inference. Genome Biol. Evol. 8:330–344.

Taylor, A. T., J. M. Long, R. A. Snow, and M. J. Porta. 2020. Hybridization and Population Genetics of Alligator Gar in Lake Texoma. North Am. J. Fish. Manag. 40:544–554.

Thompson, A. W., M. B. Hawkins, E. Parey, D. J. Wcisel, T. Ota, K. Kawasaki, E. Funk, M. Losilla, O. E. Fitch, Q. Pan, R. Feron, A. Louis, J. Montfort, M. Milhes, B. L. Racicot, K. L. Childs, Q. Fontenot, A. Ferrara, S. R. David, A. R. McCune, A. Dornburg, J. A. Yoder, Y. Guiguen, H. Roest Crollius, C. Berthelot, M. P. Harris, and I. Braasch. 2021. The bowfin genome illuminates the developmental evolution of ray-finned fishes. Nat. Genet. 53:1373–1384. Nature Publishing Group.

Title, P. O., S. Singhal, M. C. Grundler, G. C. Costa, R. A. Pyron, T. J. Colston, M. R. Grundler, I. Prates, N. Stepanova, M. E. H. Jones, L. B. Q. Cavalcanti, G. R. Colli, N. Di-Poï, S. C. Donnellan, C. Moritz, D. O. Mesquita, E. R. Pianka, S. A. Smith, L. J. Vitt, and D. L. Rabosky. 2024. The macroevolutionary singularity of snakes. Science 383:918–923. American Association for the Advancement of Science.

Upham, N. S., J. A. Esselstyn, and W. Jetz. 2021. Molecules and fossils tell distinct yet complementary stories of mammal diversification. Curr. Biol. 31:4195–4206.e3.

Valero-Mora, P. M. 2010. ggplot2: Elegant Graphics for Data Analysis. J. Stat. Softw. 35:1–3.

Wang, K., J. Wang, C. Zhu, L. Yang, Y. Ren, J. Ruan, G. Fan, J. Hu, W. Xu, X. Bi, Y. Zhu, Y. Song, H. Chen, T. Ma, R. Zhao, H. Jiang, B. Zhang, C. Feng, Y. Yuan, X. Gan, Y. Li, H. Zeng, Q. Liu, Y. Zhang, F. Shao, S. Hao, H. Zhang, X. Xu, X. Liu, D. Wang, M. Zhu, G. Zhang, W. Zhao, Q. Qiu, S. He, and W. Wang. 2021. African lungfish genome sheds light on the vertebrate water-to-land transition. Cell 184:1362-192.e18.

Wang, Z., J. Pascual-Anaya, A. Zadissa, W. Li, Y. Niimura, Z. Huang, C. Li, S. White, Z. Xiong, D. Fang, B. Wang, Y. Ming, Y. Chen, Y. Zheng, S. Kuraku, M. Pignatelli, J. Herrero, K. Beal, M. Nozawa, Q. Li, J. Wang, H. Zhang, L. Yu, S. Shigenobu, J. Wang, J. Liu, P. Flicek, S. Searle, J. Wang, S. Kuratani, Y. Yin, B. Aken, G. Zhang, and N. Irie. 2013. The draft genomes of soft-shell turtle and green sea turtle yield insights into the development and evolution of the turtle-specific body plan. Nat. Genet. 45:701–706. Nature Publishing Group.

Weaver, L. N., J. R. Kelson, R. M. Holder, N. A. Niemi, and C. Badgley. 2024. On the role of tectonics in stimulating the Cretaceous diversification of mammals. Earth-Sci. Rev. 248:104630.

Wheeler, T. J., and S. R. Eddy. 2013. nhmmer: DNA homology search with profile HMMs. Bioinformatics 29:2487–2489.

Whiteside, D. I., S. A. V. Chambi-Trowell, and M. J. Benton. 2022. A Triassic crown squamate. Sci. Adv. 8:eabq8274. American Association for the Advancement of Science.

Wible, J. R., G. W. Rougier, M. J. Novacek, and R. J. Asher. 2007. Cretaceous eutherians and Laurasian origin for placental mammals near the K/T boundary. Nature 447:1003–1006.

Wu, S., F. E. Rheindt, J. Zhang, J. Wang, L. Zhang, C. Quan, Z. Li, M. Wang, F. Wu, Y. Qu, S. V. Edwards, Z. Zhou, and L. Liu. 2024. Genomes, fossils, and the concurrent rise of modern birds and flowering plants in the Late Cretaceous. Proc. Natl. Acad. Sci. 121:e2319696121. Proceedings of the National Academy of Sciences.

Yu, D., Y. Ren, M. Uesaka, A. J. S. Beavan, M. Muffato, J. Shen, Y. Li, I. Sato, W. Wan, J. W. Clark, J. N. Keating, E. M. Carlisle, R. P. Dearden, S. Giles, E. Randle, R. S. Sansom, R. Feuda, J. F. Fleming, F. Sugahara, C. Cummins, M. Patricio, W. Akanni, S. D’Aniello, C. Bertolucci, N. Irie, C. Alev, G. Sheng, A. de Mendoza, I. Maeso, M. Irimia, B. Fromm, K. J. Peterson, S. Das, M. Hirano, J. P. Rast, M. D. Cooper, J. Paps, D. Pisani, S. Kuratani, F. J. Martin, W. Wang, P. C. J. Donoghue, Y. E. Zhang, and J. Pascual-Anaya. 2024. Hagfish genome elucidates vertebrate whole-genome duplication events and their evolutionary consequences. Nat. Ecol. Evol. 8:519–535. Nature Publishing Group.

Zhang, C., M. Rabiee, E. Sayyari, and S. Mirarab. 2018. ASTRAL-III: polynomial time species tree reconstruction from partially resolved gene trees. BMC Bioinformatics 19:153.

Zhao, T., A. Zwaenepoel, J.-Y. Xue, S.-M. Kao, Z. Li, M. E. Schranz, and Y. Van de Peer. 2021a. Whole-genome microsynteny-based phylogeny of angiosperms. Nat. Commun. 12:3498. Nature Publishing Group.

Zhao, W., X. Zhang, G. Jia, Y. Shen, and M. Zhu. 2021b. The Silurian-Devonian boundary in East Yunnan (South China) and the minimum constraint for the lungfish-tetrapod split. Sci. China Life Sci. 64:1.

Zheng, Y., and J. J. Wiens. 2016. Combining phylogenomic and supermatrix approaches, and a time-calibrated phylogeny for squamate reptiles (lizards and snakes) based on 52 genes and 4162 species. Mol. Phylogenet. Evol. 94:537–547.

Zhu, T., Y. Li, Y. Pang, Y. Han, J. Li, Z. Wang, X. Liu, H. Li, Y. Hua, H. Jiang, H. Teng, J. Quan, Y. Liu, M. Geng, M. Li, F. Hui, J. Liu, Q. Qiu, Q. Li, and Y. Ren. 2021. Chromosome-level genome assembly of Lethenteron reissneri provides insights into lamprey evolution. Mol. Ecol. Resour. 21:448–463.

Zuntini, A. R., T. Carruthers, O. Maurin, P. C. Bailey, K. Leempoel, G. E. Brewer, N. Epitawalage, E. Françoso, B. Gallego-Paramo, C. McGinnie, R. Negrão, S. R. Roy, L. Simpson, E. Toledo Romero, V. M. A. Barber, L. Botigué, J. J. Clarkson, R. S. Cowan, S. Dodsworth, M. G. Johnson, J. T. Kim, L. Pokorny, N. J. Wickett, G. M. Antar, L. DeBolt, K. Gutierrez, K. P. Hendriks, A. Hoewener, A.-Q. Hu, E. M. Joyce, I. A. B. S. Kikuchi, I. Larridon, D. A. Larson, E. J. de Lírio, J.-X. Liu, P. Malakasi, N. A. S. Przelomska, T. Shah, J. Viruel, T. R. Allnutt, G. K. Ameka, R. L. Andrew, M. S. Appelhans, M. Arista, M. J. Ariza, J. Arroyo, W. Arthan, J. B. Bachelier, C. D. Bailey, H. F. Barnes, M. D. Barrett, R. L. Barrett, R. J. Bayer, M. J. Bayly, E. Biffin, N. Biggs, J. L. Birch, D. Bogarín, R. Borosova, A. M. C. Bowles, P. C. Boyce, G. L. C. Bramley, M. Briggs, L. Broadhurst, G. K. Brown, J. J. Bruhl, A. Bruneau, S. Buerki, E. Burns, M. Byrne, S. Cable, A. Calladine, M. W. Callmander, Á. Cano, D. J. Cantrill, W. M. Cardinal-McTeague, M. M. Carlsen, A. J. A. Carruthers, A. de Castro Mateo, M. W. Chase, L. W. Chatrou, M. Cheek, S. Chen, M. J. M. Christenhusz, P.-A. Christin, M. A. Clements, S. C. Coffey, J. G. Conran, X. Cornejo, T. L. P. Couvreur, I. D. Cowie, L. Csiba, I. Darbyshire, G. Davidse, N. M. J. Davies, A. P. Davis, K. van Dijk, S. R. Downie, M. F. Duretto, M. R. Duvall, S. L. Edwards, U. Eggli, R. H. J. Erkens, M. Escudero, M. de la Estrella, F. Fabriani, M. F. Fay, P. de L. Ferreira, S. Z. Ficinski, R. M. Fowler, S. Frisby, L. Fu, T. Fulcher, M. Galbany-Casals, E. M. Gardner, D. A. German, A. Giaretta, M. Gibernau, L. J. Gillespie, C. C. González, D. J. Goyder, S. W. Graham, A. Grall, L. Green, B. F. Gunn, D. G. Gutiérrez, J. Hackel, T. Haevermans, A. Haigh, J. C. Hall, T. Hall, M. J. Harrison, S. A. Hatt, O. Hidalgo, T. R. Hodkinson, G. D. Holmes, H. C. F. Hopkins, C. J. Jackson, S. A. James, R. W. Jobson, G. Kadereit, I. M. Kahandawala, K. Kainulainen, M. Kato, E. A. Kellogg, G. J. King, B. Klejevskaja, B. B. Klitgaard, R. R. Klopper, S. Knapp, M. A. Koch, J. H. Leebens-Mack, F. Lens, C. J. Leon, É. Léveillé-Bourret, G. P. Lewis, D.-Z. Li, L. Li, S. Liede-Schumann, T. Livshultz, D. Lorence, M. Lu, P. Lu-Irving, J. Luber, E. J. Lucas, M. Luján, M. Lum, T. D. Macfarlane, C. Magdalena, V. F. Mansano, L. E. Masters, S. J. Mayo, K. McColl, A. J. McDonnell, A. E. McDougall, T. G. B. McLay, H. McPherson, R. I. Meneses, V. S. F. T. Merckx, F. A. Michelangeli, J. D. Mitchell, A. K. Monro, M. J. Moore, T. L. Mueller, K. Mummenhoff, J. Munzinger, P. Muriel, D. J. Murphy, K. Nargar, L. Nauheimer, F. J. Nge, R. Nyffeler, A. Orejuela, E. M. Ortiz, L. Palazzesi, A. L. Peixoto, S. K. Pell, J. Pellicer, D. S. Penneys, O. A. Perez-Escobar, C. Persson, M. Pignal, Y. Pillon, J. R. Pirani, G. M. Plunkett, R. F. Powell, G. T. Prance, C. Puglisi, M. Qin, R. K. Rabeler, P. E. J. Rees, M. Renner, E. H. Roalson, M. Rodda, Z. S. Rogers, S. Rokni, R. Rutishauser, M. F. de Salas, H. Schaefer, R. J. Schley, A. Schmidt-Lebuhn, A. Shapcott, I. Al-Shehbaz, K. A. Shepherd, M. P. Simmons, A. O. Simões, A. R. G. Simões, M. Siros, E. C. Smidt, J. F. Smith, N. Snow, D. E. Soltis, P. S. Soltis, R. J. Soreng, C. A. Sothers, J. R. Starr, P. F. Stevens, S. C. K. Straub, L. Struwe, J. M. Taylor, I. R. H. Telford, A. H. Thornhill, I. Tooth, A. Trias-Blasi, F. Udovicic, T. M. A. Utteridge, J. C. Del Valle, G. A. Verboom, H. P. Vonow, M. S. Vorontsova, J. M. de Vos, N. Al-Wattar, M. Waycott, C. A. D. Welker, A. J. White, J. J. Wieringa, L. T. Williamson, T. C. Wilson, S. Y. Wong, L. A. Woods, R. Woods, S. Worboys, M. Xanthos, Y. Yang, Y.-X. Zhang, M.-Y. Zhou, S. Zmarzty, F. O. Zuloaga, A. Antonelli, S. Bellot, D. M. Crayn, O. M. Grace, P. J. Kersey, I. J. Leitch, H. Sauquet, S. A. Smith, W. L. Eiserhardt, F. Forest, and W. J. Baker. 2024. Phylogenomics and the rise of the angiosperms. Nature 629:843–850. Nature Publishing Group.

1995. The distribution of the vertebrates in the Late Ordovician and Early Silurian palaeobasins of the Siberian Platform. Bull. Muséum Natl. Hist. Nat. 4ème Sér. – Sect. C – Sci. Terre Paléontol. Géologie Minéralogie 17:39–55.

## References

A.f B. 2004. THE FIRST DISCOVERY OF AN ANGLERFISH (TELEOSTEI, LOPHIIDAE) IN THE EOCENE OF THE NORTHERN CAUCASUS. Available from: https://repository.geologyscience.ru/handle/123456789/39456

Alfaro ME, Faircloth BC, Harrington RC, Sorenson L, Friedman M, Thacker CE, Oliveros CH, Černý D, Near TJ. 2018. Explosive diversification of marine fishes at the Cretaceous–Palaeogene boundary. Nat. Ecol. Evol. 2:688–696.

Álvarez-Carretero S, Tamuri AU, Battini M, Nascimento FF, Carlisle E, Asher RJ, Yang Z, Donoghue PCJ, dos Reis M. 2022. A species-level timeline of mammal evolution integrating phylogenomic data. Nature 602:263–267.

Alves YM, Bergqvist LP, Brito PM. 2019. The dorsal and pectoral fin spines of catfishes (Ostariophysi: Siluriformes) from the Bauru Group (Late Cretaceous), Brazil: A comparative and critical analysis. J. South Am. Earth Sci. 92:32–40.

Amaral CR, Alvarado-Ortega J, Brito PM. 2013. Sapperichthys gen. nov., a new gonorynchid from the Cenomanian of Chiapas, Mexico. Mesoz. Fishes 5:305–323.

and,. 2017. Morphological and histological evidence for the oldest known softshell turtles from Japan. J. Vertebr. Paleontol. 37:e1278606.

Anderson JS, Reisz RR, Scott D, Fröbisch NB, Sumida SS. 2008. A stem batrachian from the Early Permian of Texas and the origin of frogs and salamanders. Nature 453:515–518.

Andrews JV, Schein JP, Friedman M. 2023. An earliest Paleocene squirrelfish (Teleostei: Beryciformes: Holocentroidea) and its bearing on the timescale of holocentroid evolution. J. Syst. Palaeontol. 21:2168571.

Anon. 1998. A NEW PALEOCENE ARMADILLO (MAMMALIA, DASYPODOIDEA) FROM THE ITABORAÍ BASIN, BRAZIL. Publ. Electrónica Asoc. Paleontológica Argent. [Internet] 5. Available from: https://peapaleontologica.org.ar/index.php/peapa/article/view/174

Anon. 2019. Mystacodon selenensis, the earliest known toothed mysticete (Cetacea, Mammalia) from the late Eocene of Peru: anatomy, phylogeny, and feeding adaptations. Geodiversitas 41:401–499.

Anon. Abstract: THE AGE OF DINOSAUR-BEARING STRATA AT PHOEBUS LANDING, CAPE FEAR RIVER, NORTH CAROLINA (Northeastern Section (39th Annual) and Southeastern Section (53rd Annual) Joint Meeting (March 25–27, 2004)). Available from: https://gsa.confex.com/gsa/2004NE/webprogram/Paper69560.html

Arcila D, Ortí G, Vari R, Armbruster JW, Stiassny MLJ, Ko KD, Sabaj MH, Lundberg J, Revell LJ, Betancur-R. R. 2017. Genome-wide interrogation advances resolution of recalcitrant groups in the tree of life. *Nat*. Ecol. Evol. 1:1–10.

Argyriou T, Giles S, Friedman M. A Permian fish reveals widespread distribution of neopterygian-like jaw suspension. eLife 11:e58433.

Argyriou T, Giles S, Friedman M, Romano C, Kogan I, Sánchez-Villagra MR. 2018. Internal cranial anatomy of Early Triassic species of †Saurichthys (Actinopterygii: †Saurichthyiformes): implications for the phylogenetic placement of †saurichthyiforms. BMC Evol. Biol. 18:161.

Arratia G. 2000. Remarkable teleostean fishes from the Late Jurassic of southern Germany and their phylogenetic relationships. Foss. Rec. 3:137–179.

Ascarrunz E, Rage J-C, Legreneur P, Laurin M. 2016. Triadobatrachus massinoti, the earliest known lissamphibian (Vertebrata: Tetrapoda) re-examined by μCT scan, and the evolution of trunk length in batrachians. Contrib. Zool. 85:201–223.

Avilla LS, Mothé D. 2021. Out of Africa: A New Afrotheria Lineage Rises From Extinct South American Mammals. Front. Ecol. Evol. [Internet] 9. Available from: https://www.frontiersin.org/journals/ecology-and-evolution/articles/10.3389/fevo.2021.654302/full

Báez AM. 2013. Anurans from the Early Cretaceous Lagerstätte of Las Hoyas, Spain: New evidence on the Mesozoic diversification of crown-clade Anura. Cretac. Res. 41:90–106.

Bagley JC, Mayden RL, Harris PM. 2018. Phylogeny and divergence times of suckers (Cypriniformes: Catostomidae) inferred from Bayesian total-evidence analyses of molecules, morphology, and fossils. PeerJ [Internet] 6. Available from: https://www.ncbi.nlm.nih.gov/pmc/articles/PMC6035723/

Bannikov AF, Carnevale G. 2010. Bellwoodilabrus landinii n. gen., n. sp., a new genus and species of labrid fish (Teleostei, Perciformes) from the Eocene of Monte Bolca. Geodiversitas 32:201–220.

Beck RMD, Godthelp H, Weisbecker V, Archer M, Hand SJ. 2008. Australia’s Oldest Marsupial Fossils and their Biogeographical Implications. PLoS ONE 3:e1858.

Becker RT, Gradstein FM, Hammer O. 2012. Chapter 22 - The Devonian Period. In: Gradstein Felix M., Ogg JG, Schmitz MD, Ogg GM, editors. The Geologic Time Scale. Boston: Elsevier. p. 559–601. Available from: https://www.sciencedirect.com/science/article/pii/B9780444594259000226

Bellwood D. 1990. A new fossil fish Phyllopharyngodon longipinnis gen. et sp. nov. (family Labridae) from the Eocene, Monte Bolca, Italy. Studi E Ric. Sui Giacimenti Terziari Bolca 6:149–160.

Benito J, Chen A, Wilson LE, Bhullar B-AS, Burnham D, Field DJ. 2022. Forty new specimens of Ichthyornis provide unprecedented insight into the postcranial morphology of crownward stem group birds. PeerJ 10:e13919.

Benito J, Kuo P-C, Widrig KE, Jagt JWM, Field DJ. 2022. Cretaceous ornithurine supports a neognathous crown bird ancestor. Nature 612:100–105.

Benton MJ, Donoghue PC, Asher RJ, Friedman M, Near TJ, Vinther J. 2015. Constraints on the timescale of animal evolutionary history. Palaeontol Electron 18:1–106.

Benton MJ, Donoghue PCJ. 2007. Paleontological Evidence to Date the Tree of Life. Mol. Biol. Evol. 24:26–53.

Bergqvist LP, Pereira PVLG, Machado AS, Castro MCD, Melki LB, Lopes RT. 2019. Osteoderm microstructure of Riostegotherium yanei, the oldest Xenarthra. An. Acad. Bras. Ciênc. 91:e20181290.

Bernardi M, Klein H, Petti FM, Ezcurra MD. 2015. The Origin and Early Radiation of Archosauriforms: Integrating the Skeletal and Footprint Record. PLoS ONE 10:e0128449.

Berta A. 1991. New Enaliarctos (Pinnipedimorpha) from the Oligocene and Miocene of Oregon and the Role of “Enaliarctids” in Pinniped Phylogeny. Available from: http://repository.si.edu/xmlui/handle/10088/19145

Berv JS, Field DJ. 2018. Genomic Signature of an Avian Lilliput Effect across the K-Pg Extinction. Syst. Biol. 67:1–13.

Berv JS, Singhal S, Field DJ, Walker-Hale N, McHugh SW, Shipley JR, Miller ET, Kimball RT, Braun EL, Dornburg A, et al. 2024. Genome and life-history evolution link bird diversification to the end-Cretaceous mass extinction. Sci. Adv. 10:eadp0114.

Betancur-R. R, Arcila D, Vari RP, Hughes LC, Oliveira C, Sabaj MH, Ortí G. 2019. Phylogenomic incongruence, hypothesis testing, and taxonomic sampling: The monophyly of characiform fishes*. Evolution 73:329–345.

Betancur-R R, Broughton RE, Wiley EO, Carpenter K, López JA, Li C, Holcroft NI, Arcila D, Sanciangco M, Ii JCC, et al. 2013. The Tree of Life and a New Classification of Bony Fishes. PLOS Curr. Tree Life [Internet]. Available from: http://currents.plos.org/treeoflife/article/the-tree-of-life-and-a-new-classification-of-bony-fishes/

Bever GS, Lyson TR, Field DJ, Bhullar B-AS. 2015. Evolutionary origin of the turtle skull. Nature 525:239–242.

Bi X, Wang K, Yang L, Pan H, Jiang Haifeng, Wei Q, Fang M, Yu H, Zhu C, Cai Y, et al. 2021a. Tracing the genetic footprints of vertebrate landing in non-teleost ray-finned fishes. Cell 184:1377–1391.e14.

Bi X, Wang K, Yang L, Pan H, Jiang Haifeng, Wei Q, Fang M, Yu H, Zhu C, Cai Y, et al. 2021b. Tracing the genetic footprints of vertebrate landing in non-teleost ray-finned fishes. Cell 184:1377–1391.e14.

Bian C, Hu Y, Ravi V, Kuznetsova IS, Shen X, Mu X, Sun Y, You X, Li J, Li X, et al. 2016. The Asian arowana (Scleropages formosus) genome provides new insights into the evolution of an early lineage of teleosts. Sci. Rep. 6:1–17.

Bittencourt JS, Simões TR, Caldwell MW, Langer MC. 2020. Discovery of the oldest South American fossil lizard illustrates the cosmopolitanism of early South American squamates. *Commun*. Biol. 3:1–11.

Boessenecker RW, Beatty BL, Geisler JH. 2023. New specimens and species of the Oligocene toothed baleen whale Coronodon from South Carolina and the origin of Neoceti. PeerJ 11:e14795.

Bona P, Ezcurra MD, Barrios F, Fernandez Blanco MV. 2018. A new Palaeocene crocodylian from southern Argentina sheds light on the early history of caimanines. Proc. R. Soc. B Biol. Sci. 285:20180843.

Borsuk-Bialynicka M, Moody SM. 1984. Priscagaminae, a new subfamily of the Agamidae (Sauria) from the Late Cretaceous of the Gobi Desert. Acta Palaeontol. Pol. [Internet] 29. Available from: https://agro.icm.edu.pl/agro/element/bwmeta1.element.agro-a23634c5-3ba7-4810-86b8-46315cd4a85a

Bowman J, Enard D, Lynch VJ. 2023. Phylogenomics reveals an almost perfect polytomy among the almost ungulates (Paenungulata). bioRxiv:2023.12.07.570590.

Braasch I, Gehrke AR, Smith JJ, Kawasaki K, Manousaki T, Pasquier J, Amores A, Desvignes T, Batzel P, Catchen J, et al. 2016. The spotted gar genome illuminates vertebrate evolution and facilitates human-teleost comparisons. Nat. Genet. 48:427–437.

Brazeau MD, Giles S, Dearden RP, Jerve A, Ariunchimeg Y, Zorig E, Sansom R, Guillerme T, Castiello M. 2020. Endochondral bone in an Early Devonian ‘placoderm’ from Mongolia. *Nat*. Ecol. Evol. 4:1477–1484.

Brinkmann H, Venkatesh B, Brenner S, Meyer A. 2004. Nuclear protein-coding genes support lungfish and not the coelacanth as the closest living relatives of land vertebrates. Proc. Natl. Acad. Sci. 101:4900–4905.

Brito PM, Alvarado-Ortega J, Meunier FJ. 2017. Earliest known lepisosteoid extends the range of anatomically modern gars to the Late Jurassic. Sci. Rep. 7:17830.

Brito PM, Figueiredo FJ, Leal MEC. 2020. A revision of Laeliichthys ancestralis Santos, 1985 (Teleostei: Osteoglossomorpha) from the Lower Cretaceous of Brazil: Phylogenetic relationships and biogeographical implications. PLOS ONE 15:e0241009.

Brochu CA. 1997. Morphology, Fossils, Divergence Timing, and the Phylogenetic Relationships of Gavialis. Syst. Biol. 46:479–522.

Brownstein CD. 2018. The distinctive theropod assemblage of the Ellisdale site of New Jersey and its implications for North American dinosaur ecology and evolution during the Cretaceous. J. Paleontol. 92:1115–1129.

Brownstein CD. 2019. First Record of a Small Juvenile Giant Crocodyliform and its Ontogenetic and Biogeographic Implications. Bull. Peabody Mus. Nat. Hist. 60:81–90.

Brownstein Chase Doran. 2023. Palaeospondylus and the early evolution of gnathostomes. Nature 620:E20–E22.

Brownstein CD. 2023. Syngnathoid Evolutionary History and the Conundrum of Fossil Misplacement. Integr. Org. Biol. 5:obad011.

Brownstein CD, Dornburg A, Near TJ. 2025. Cenozoic evolutionary history obscures the Mesozoic origins of acanthopterygian fishes. Evolution:qpaf040.

Brownstein CD, Lyson TR. 2022. Giant gar from directly above the Cretaceous–Palaeogene boundary suggests healthy freshwater ecosystems existed within thousands of years of the asteroid impact. Biol. Lett. 18:20220118.

Brownstein CD, MacGuigan DJ, Kim D, Orr O, Yang L, David SR, Kreiser B, Near TJ. 2024. The genomic signatures of evolutionary stasis. Evolution 78:821–834.

Brownstein CD, Meyer DL, Fabbri M, Bhullar B-AS, Gauthier JA. 2022. Evolutionary origins of the prolonged extant squamate radiation. Nat. Commun. 13:7087.

Brownstein CD, Near TJ. 2024. A giant bowfin from a Paleocene hothouse ecosystem in North America. Zool. J. Linn. Soc. 202:zlae042.

Brownstein CD, Near TJ, Dearden RP. 2024. The Palaeozoic assembly of the holocephalan body plan far preceded post-Cretaceous radiations into the ocean depths. Proc. R. Soc. B Biol. Sci. 291:20241824.

Brownstein Chase D., Simões TR, Caldwell MW, Lee MSY, Meyer DL, Scarpetta SG. 2023. The affinities of the Late Triassic *Cryptovaranoides* and the age of crown squamates. R. Soc. Open Sci. 10:230968.

Brownstein Chase Doran, Yang L, Friedman M, Near TJ. 2023. Phylogenomics of the Ancient and Species-Depauperate Gars Tracks 150 Million Years of Continental Fragmentation in the Northern Hemisphere. Syst. Biol. 72:213–227.

Brownstein CD, Zapfe KL, Lott S, Harrington RC, Ghezelayagh A, Dornburg A, Near TJ. 2024. Synergistic innovations enabled the radiation of anglerfishes in the deep open ocean. Curr. Biol. 34:2541–2550.e4.

Budd GE, Mann RP. 2023. Two Notorious Nodes: A Critical Examination of Relaxed Molecular Clock Age Estimates of the Bilaterian Animals and Placental Mammals. Syst. Biol.:syad057.

Burbrink FT, Grazziotin FG, Pyron RA, Cundall D, Donnellan S, Irish F, Keogh JS, Kraus F, Murphy RW, Noonan B, et al. 2020. Interrogating Genomic-Scale Data for Squamata (Lizards, Snakes, and Amphisbaenians) Shows no Support for Key Traditional Morphological Relationships. Syst. Biol. 69:502–520.

Caldwell MW, Nydam RL, Palci A, Apesteguía S. 2015. The oldest known snakes from the Middle Jurassic-Lower Cretaceous provide insights on snake evolution. Nat. Commun. 6:5996.

Capobianco A, Friedman M. 2024. Fossils indicate marine dispersal in osteoglossid fishes, a classic example of continental vicariance. Proc. R. Soc. B Biol. Sci. 291:20241293.

Capobianco A, Zouhri S, Friedman M. 2025. A long-snouted marine bonytongue (Teleostei: Osteoglossidae) from the early Eocene of Morocco and the phylogenetic affinities of marine osteoglossids. Zool. J. Linn. Soc. 203:zlae015.

Carnevale G, Pietsch TW. 2009. An Eocene Frogfish from Monte Bolca, Italy: The Earliest Known Skeletal Record for the Family. Palaeontology 52:745–752.

Carnevale G, Pietsch TW. 2010. Eocene handfishes from Monte Bolca, with description of a new genus and species, and a phylogeny of the family Brachionichthyidae (Teleostei: Lophiiformes). Zool. J. Linn. Soc. 160:621–647.

Carnevale G, Pietsch TW. 2011. Batfishes from the Eocene of Monte Bolca. Geol. Mag. 148:461–472.

Casanovas-Vilar I, Garcia-Porta J, Fortuny J, Sanisidro Ó, Prieto J, Querejeta M, Llácer S, Robles JM, Bernardini F, Alba DM. Oldest skeleton of a fossil flying squirrel casts new light on the phylogeny of the group. eLife 7:e39270.

Cavin L, Suteethorn V, Buffetaut E, Tong H. 2007. A new Thai Mesozoic lungfish (Sarcopterygii, Dipnoi) with an insight into post-Palaeozoic dipnoan evolution. Zool. J. Linn. Soc. 149:141–177.

Celerino de Carvalho J, Santucci RM. 2024. A new fossil Squamata from the Quiricó Formation (Lower Cretaceous), Sanfranciscana Basin, Minas Gerais, Brazil. Cretac. Res. 154:105717.

Chakrabarty P, Faircloth BC, Alda F, Ludt WB, Mcmahan CD, Near TJ, Dornburg A, Albert JS, Arroyave J, Stiassny MLJ, et al. 2017. Phylogenomic Systematics of Ostariophysan Fishes: Ultraconserved Elements Support the Surprising Non-Monophyly of Characiformes. Syst. Biol. 66:881–895.

Chen M-Y, Liang D, Zhang P. 2015. Selecting Question-Specific Genes to Reduce Incongruence in Phylogenomics: A Case Study of Jawed Vertebrate Backbone Phylogeny. Syst. Biol. 64:1104–1120.

Chester SGB, Bloch JI, Boyer DM, Clemens WA. 2015. Oldest known euarchontan tarsals and affinities of Paleocene Purgatorius to Primates. Proc. Natl. Acad. Sci. 112:1487–1492.

Chester SGB, Williamson TE, Bloch JI, Silcox MT, Sargis EJ. 2017. Oldest skeleton of a plesiadapiform provides additional evidence for an exclusively arboreal radiation of stem primates in the Palaeocene. R. Soc. Open Sci. 4:170329.

Chiari Y, Cahais V, Galtier N, Delsuc F. 2012. Phylogenomic analyses support the position of turtles as the sister group of birds and crocodiles (Archosauria). BMC Biol. 10:65.

Choo B, Zhu M, Qu Q, Yu X, Jia L, Zhao W. 2017. A new osteichthyan from the late Silurian of Yunnan, China. PLOS ONE 12:e0170929.

Claramunt S, Cracraft J. 2015. A new time tree reveals Earth history’s imprint on the evolution of modern birds. Sci. Adv. 1:e1501005.

Clarke JA, Norell MA, Dashzeveg D. 2005. New Avian Remains from the Eocene of Mongolia and the Phylogenetic Position of the Eogruidae (Aves, Gruoidea). Am. Mus. Novit. 2005:1–17.

Cloutier R, Clement AM, Lee MSY, Noël R, Béchard I, Roy V, Long JA. 2020. Elpistostege and the origin of the vertebrate hand. Nature 579:549–554.

Coates MI, Finarelli JA, Sansom IJ, Andreev PS, Criswell KE, Tietjen K, Rivers ML, La Riviere PJ. 2018. An early chondrichthyan and the evolutionary assembly of a shark body plan. Proc. R. Soc. B Biol. Sci. 285:20172418.

Cole TL, Zhou C, Fang M, Pan H, Ksepka DT, Fiddaman SR, Emerling CA, Thomas DB, Bi X, Fang Q, et al. 2022. Genomic insights into the secondary aquatic transition of penguins. Nat. Commun. 13:3912.

Conrad JL. 2008. Phylogeny And Systematics Of Squamata (Reptilia) Based On Morphology. Bull. Am. Mus. Nat. Hist. 2008:1–182.

Cossette AP, Brochu CA. 2020. A systematic review of the giant alligatoroid Deinosuchus from the Campanian of North America and its implications for the relationships at the root of Crocodylia. J. Vertebr. Paleontol. 40:e1767638.

Crane A, Benito J, Chen A, Musser G, Torres CR, Clarke JA, Lautenschlager S, Ksepka DT, Field DJ. 2024. Taphonomic damage obfuscates interpretation of the retroarticular region of the Asteriornis mandible. Geobios [Internet]. Available from: https://www.sciencedirect.com/science/article/pii/S0016699524000536

Crawford NG, Faircloth BC, McCormack JE, Brumfield RT, Winker K, Glenn TC. 2012. More than 1000 ultraconserved elements provide evidence that turtles are the sister group of archosaurs. Biol. Lett. 8:783–786.

Croghan JA, Palci A, Onary S, Lee MSY, Caldwell MW. 2024. Morphology and systematics of a new fossil snake from the early Rupelian (Oligocene) White River Formation, Wyoming. Zool. J. Linn. Soc.:zlae073.

Cui X, Friedman M, Qiao T, Yu Y, Zhu M. 2022. The rapid evolution of lungfish durophagy. Nat. Commun. 13:2390.

Cui X, Friedman M, Yu Y, Zhu Y, Zhu M. 2023. Bony-fish-like scales in a Silurian maxillate placoderm. Nat. Commun. 14:7622.

Cui X, Qiao T, Zhu M. 2019. Scale morphology and squamation pattern of Guiyu oneiros provide new insights into early osteichthyan body plan. Sci. Rep. 9:4411.

Darlim G, Lee MSY, Walter J, Rabi M. 2022. The impact of molecular data on the phylogenetic position of the putative oldest crown crocodilian and the age of the clade. Biol. Lett. 18:20210603.

Davesne D, Carnevale G, Friedman M. 2017. Bajaichthys elegans from the Eocene of Bolca (Italy) and the overlooked morphological diversity of Zeiformes (Teleostei, Acanthomorpha). Palaeontology 60:255–268.

Davesne D, Friedman M, Schmitt AD, Fernandez V, Carnevale G, Ahlberg PE, Sanchez S, Benson RBJ. 2021. Fossilized cell structures identify an ancient origin for the teleost whole-genome duplication. Proc. Natl. Acad. Sci. 118:e2101780118.

Dawson MR, Beard KC. 1996. New late Paleocene rodents (Mammalia) from big multi quarry, Washakie Basin, Wyoming. Palaeovertebrata 25:301–321.

Dawson MR, Janis CM, Gunnell GF, Uhen MD. 2008. Evolution of tertiary mammals of North America.

Denton R. 2022. ALBEMARLE SOUND NC - A MODERN ANALOG FOR THE ELLISDALE FOSSIL SITE (LATE CRETACEOUS, CAMPANIAN, NJ). In: GSA. Available from: https://gsa.confex.com/gsa/2022NC/webprogram/Paper373518.html

Dingus L, Loope D, Dashzeveg D, Swisher C, Minjin C, Novacek M, Norell M. 2008. The Geology of Ukhaa Tolgod (Djadokhta Formation, Upper Cretaceous, Nemegt Basin, Mongolia). Dep. Earth Atmospheric Sci. Fac. Publ. [Internet]. Available from: https://digitalcommons.unl.edu/geosciencefacpub/217

Diogo R. 2009. GRANDE, T., F. POYATO-ARIZA & R. DIOGO. (2009). Gonorynchiform interrelationships: historical overview, analysis, and revised systematics of the group. In: Grande, T., F. Poyato-Ariza & R. Diogo (eds.), Gonorynchiformes and ostariophysan relationships – a comprehensive review, Science Publishers and Taylor & Francis (Oxford, UK): 221-231.

Dornburg A, Friedman M, Near TJ. 2015. Phylogenetic analysis of molecular and morphological data highlights uncertainty in the relationships of fossil and living species of Elopomorpha (Actinopterygii: Teleostei). Mol. Phylogenet. Evol. 89:205–218.

Dornburg A, Near TJ. 2021. The Emerging Phylogenetic Perspective on the Evolution of Actinopterygian Fishes. Annu. Rev. Ecol. Evol. Syst. 52:427–452.

Dornburg A, Townsend JP, Brooks W, Spriggs E, Eytan RI, Moore JA, Wainwright PC, Lemmon A, Lemmon EM, Near TJ. 2017. New insights on the sister lineage of percomorph fishes with an anchored hybrid enrichment dataset. Mol. Phylogenet. Evol. 110:27–38.

Du K, Stöck M, Kneitz S, Klopp C, Woltering JM, Adolfi MC, Feron R, Prokopov D, Makunin A, Kichigin I, et al. 2020. The sterlet sturgeon genome sequence and the mechanisms of segmental rediploidization. *Nat*. Ecol. Evol. 4:841–852.

Esselstyn JA, Oliveros CH, Swanson MT, Faircloth BC. 2017. Investigating Difficult Nodes in the Placental Mammal Tree with Expanded Taxon Sampling and Thousands of Ultraconserved Elements. Genome Biol. Evol. 9:2308–2321.

Estes R, de Queiroz K, Gauthier J. 1988. Phylogenetic relationships within Squamata. In: Phylogenetic Relationships of the lizard families. Stanford, California: Stanford University Press. p. 119–281. Available from: https://books.google.com/books?hl=en&lr=&id=h5fIP1X7YvoC&oi=fnd&pg=PA119&ots=fW_Kh2cy1A&sig=Ukv57Cm2U52V1_tXWHr5fGB15b4

Evans SE, Borsuk-Bialynicka M. 1998. A stem-group frog from the Early Triassic of Poland. Acta Palaeontol. Pol. 43:573–580.

Evans SE, Raia P, Barbera C. 2006. The Lower Cretaceous lizard genus *Chometokadmon* from Italy. Cretac. Res. 27:673–683.

Ezcurra MD, Nesbitt SJ, Bronzati M, Dalla Vecchia FM, Agnolin FL, Benson RBJ, Brissón Egli F, Cabreira SF, Evers SW, Gentil AR, et al. 2020. Enigmatic dinosaur precursors bridge the gap to the origin of Pterosauria. Nature 588:445–449.

Ezcurra MD, Scheyer TM, Butler RJ. 2014. The Origin and Early Evolution of Sauria: Reassessing the Permian Saurian Fossil Record and the Timing of the Crocodile-Lizard Divergence. PLOS ONE 9:e89165.

Field DJ, Benito J, Chen A, Jagt JWM, Ksepka DT. 2020. Late Cretaceous neornithine from Europe illuminates the origins of crown birds. Nature 579:397–401.

Field DJ, Bercovici A, Berv JS, Dunn R, Fastovsky DE, Lyson TR, Vajda V, Gauthier JA. 2018. Early Evolution of Modern Birds Structured by Global Forest Collapse at the End-Cretaceous Mass Extinction. Curr. Biol. 28:1825–1831.e2.

de Figueiredo FJ, Gallo V, Leal MEC. 2012. Phylogenetic relationships of the elopomorph fish †*Paraelops cearensis* Silva Santos revisited: Evidence from new specimens. Cretac. Res. 37:148–154.

Filleul A, Maisey JG. 2004. Redescription of Santanichthys diasii (Otophysi, Characiformes) from the Albian of the Santana Formation and comments on its implications for otophysan relationships. American Museum novitates ; no. 3455. Available from: http://hdl.handle.net/2246/2765

Foley NM, Mason VC, Harris AJ, Bredemeyer KR, Damas J, Lewin HA, Eizirik E, Gatesy J, Karlsson EK, Lindblad-Toh K, et al. 2023. A genomic timescale for placental mammal evolution. Science 380:eabl8189.

Ford DP, Evans SE, Choiniere JN, Fernandez V, Benson RBJ. 2021. A reassessment of the enigmatic diapsid *Paliguana whitei* and the early history of Lepidosauromorpha. Proc. R. Soc. B Biol. Sci. 288:20211084.

Fordyce RE, Marx FG. 2018. Gigantism Precedes Filter Feeding in Baleen Whale Evolution. Curr. Biol. 28:1670–1676.e2.

Fox RC, Youzwyshyn GP. 1994. New primitive carnivorans (Mammalia) from the Paleocene of western Canada, and their bearing on relationships of the order. J. Vertebr. Paleontol. 14:382–404.

Freisem LS, Müller J, Sues H-D, Sobral G. 2024. A new sphenodontian (Diapsida: Lepidosauria) from the Upper Triassic (Norian) of Germany and its implications for the mode of sphenodontian evolution. BMC Ecol. Evol. 24:35.

Frey L, Coates M, Ginter M, Hairapetian V, Rücklin M, Jerjen I, Klug C. 2019. The early elasmobranch Phoebodus: phylogenetic relationships, ecomorphology and a new time-scale for shark evolution. Proc. R. Soc. B Biol. Sci. 286:20191336.

Frey L, Coates MI, Tietjen K, Rücklin M, Klug C. 2020. A symmoriiform from the Late Devonian of Morocco demonstrates a derived jaw function in ancient chondrichthyans. *Commun*. Biol. 3:681.

Friedman M. 2007. The interrelationships of Devonian lungfishes (Sarcopterygii: Dipnoi) as inferred from neurocranial evidence and new data from the genus Soederberghia Lehman, 1959. Zool. J. Linn. Soc. 151:115–171.

Friedman M. 2022. The Macroevolutionary History of Bony Fishes: A Paleontological View. Annu. Rev. Ecol. Evol. Syst. 53:353–377.

Friedman M, Carnevale G. 2018. The Bolca Lagerstätten: shallow marine life in the Eocene. J. Geol. Soc. 175:569–579.

Friedman M, Feilich KL, Beckett HT, Alfaro ME, Faircloth BC, Černý D, Miya M, Near TJ, Harrington RC. 2019. A phylogenomic framework for pelagiarian fishes (Acanthomorpha: Percomorpha) highlights mosaic radiation in the open ocean. Proc. R. Soc. B Biol. Sci. 286:20191502.

de la Fuente MS, Iturralde-Vinent M. 2001. A New Pleurodiran Turtle from the Jagua Formation (Oxfordian) of Western Cuba. J. Paleontol. 75:860–869.

Gao K, Norell MA. 2000. Taxonomic composition and systematics of Late Cretaceous lizard assemblages from Ukhaa Tolgod and adjacent localities, Mongolian Gobi Desert. Bull. Am. Mus. Nat. Hist. 2000:1–118.

Gauthier JA, Kearney M, Maisano JA, Rieppel O, Behlke ADB. 2012. Assembling the Squamate Tree of Life: Perspectives from the Phenotype and the Fossil Record. Bull. Peabody Mus. Nat. Hist. 53:3–308.

Gheerbrant E. 2009. Paleocene emergence of elephant relatives and the rapid radiation of African ungulates. Proc. Natl. Acad. Sci. 106:10717–10721.

Gheerbrant E, Schmitt A, Kocsis L. 2018. Early African Fossils Elucidate the Origin of Embrithopod Mammals. Curr. Biol. 28:2167–2173.e2.

Ghezelayagh A, Harrington RC, Burress ED, Campbell MA, Buckner JC, Chakrabarty P, Glass JR, McCraney WT, Unmack PJ, Thacker CE, et al. 2022. Prolonged morphological expansion of spiny-rayed fishes following the end-Cretaceous. *Nat*. Ecol. Evol. 6:1211–1220.

Gil-Delgado A, Delclòs X, Sellés A, Galobart À, Oms O. 2023. The Early Cretaceous coastal lake Konservat-Lagerstätte of La Pedrera de Meià (Southern Pyrenees). Geol. Acta 21:1–XIII.

Giles S, Darras L, Clément G, Blieck A, Friedman M. 2015. An exceptionally preserved Late Devonian actinopterygian provides a new model for primitive cranial anatomy in ray-finned fishes. Proc. R. Soc. B Biol. Sci. 282:20151485.

Giles S, Friedman M, Brazeau MD. 2015. Osteichthyan-like cranial conditions in an Early Devonian stem gnathostome. Nature 520:82–85.

Giles S, Xu G-H, Near TJ, Friedman M. 2017. Early members of ‘living fossil’ lineage imply later origin of modern ray-finned fishes. Nature 549:265–268.

Goloboff PA, Catalano SA. 2016. TNT version 1.5, including a full implementation of phylogenetic morphometrics. Cladistics 32:221–238.

Gómez RO. 2016. A new pipid frog from the Upper Cretaceous of Patagonia and early evolution of crown-group Pipidae. Cretac. Res. 62:52–64.

Gottmann-Quesada A, Sander P. 2009. A redescription of the early archosauromorph Protorosaurus speneri MEYER, 1832, and its phylogenetic relationships. Palaeontogr. A 287:123–220.

Gradstein FM, Ogg JG, Schmitz M, Ogg G. 2021. The Geologic Time Scale 2020. Amsterdam, The Netherlands: Elsevier Science

Grande L. 2010. An Empirical Synthetic Pattern Study of Gars (lepisosteiformes) and Closely Related Species, Based Mostly on Skeletal Anatomy. the Resurrection of Holostei. Copeia 2010:iii–871.

Grande L, Jin F, Yabumoto Y, Bemis WE. 2002. Protopsephurus liui, a well-preserved primitive paddlefish (Acipenseriformes: Polyodontidae) from the Lower Cretaceous of China. J. Vertebr. Paleontol. 22:209–237.

Grande T, Poyato-Ariza FJ. 1999. Phylogenetic relationships of fossil and Recent gonorynchiform fishes (Teleostei: Ostariophysi). Zool. J. Linn. Soc. 125:197–238.

Grande TC, Borden WC, Wilson MVH, Scarpitta L. 2018. Phylogenetic Relationships among Fishes in the Order Zeiformes Based on Molecular and Morphological Data. Copeia 106:20–48.

Griffiths EF, Ford DP, Benson RBJ, Evans SE. 2021. New information on the Jurassic lepidosauromorph Marmoretta oxoniensis.Ruta M, editor. Pap. Palaeontol. 7:2255–2278.

Hackett SJ, Kimball RT, Reddy S, Bowie RCK, Braun EL, Braun MJ, Chojnowski JL, Cox WA, Han K-L, Harshman J, et al. 2008. A Phylogenomic Study of Birds Reveals Their Evolutionary History. Science 320:1763–1768.

Han G, Mao F, Bi S, Wang Y, Meng J. 2017. A Jurassic gliding euharamiyidan mammal with an ear of five auditory bones. Nature 551:451–456.

Hand SJ, Maugoust J, Beck RMD, Orliac MJ. 2023. A 50-million-year-old, three-dimensionally preserved bat skull supports an early origin for modern echolocation. Curr. Biol. 33:4624–4640.e21.

Hao S, Han K, Meng L, Huang X, Cao W, Shi C, Zhang M, Wang Y, Liu Q, Zhang Y, et al. 2020. African Arowana Genome Provides Insights on Ancient Teleost Evolution. iScience 23:101662.

Hedges SB. 2012. Amniote phylogeny and the position of turtles. BMC Biol. 10:64.

Henrici AC. 1998. A New Pipoid Anuran from the Late Jurassic Morrison Formation at Dinosaur National Monument, Utah. J. Vertebr. Paleontol. 18:321–332.

Hernández-Guerrero C, Cantalice KM, González-Rodríguez KA, Bravo-Cuevas VM. 2021. The first record of a pterothrissin (Albuliformes, Albulidae) from the Muhi Quarry, mid-Cretaceous (Albian-Cenomanian) of Hidalgo, central Mexico. J. South Am. Earth Sci. 107:103032.

Hilton EJ, Grande L, Bemis WE. 2011. Skeletal Anatomy of the Shortnose Sturgeon, Acipenser brevirostrum Lesueur, 1818, and the Systematics of Sturgeons (Acipenseriformes, Acipenseridae). Fieldiana Life Earth Sci. 2011:1–168.

Hime PM, Lemmon AR, Lemmon ECM, Prendini E, Brown JM, Thomson RC, Kratovil JD, Noonan BP, Pyron RA, Peloso PLV, et al. 2020. Phylogenomics Reveals Ancient Gene Tree Discordance in the Amphibian Tree of Life. Syst. Biol. 70:49–66.

Hirayama R, Isaji S, Hibino T. 2013. Kappachelys okurai gen. et sp. nov., a New Stem Soft-Shelled Turtle from the Early Cretaceous of Japan. In: Brinkman DB, Holroyd PA, Gardner JD, editors. Morphology and Evolution of Turtles. Dordrecht: Springer Netherlands. p. 179–185. Available from: 10.1007/978-94-007-4309-0_12

Hoffmann S, Beck RMD, Wible JR, Rougier GW, Krause DW. 2020. Phylogenetic placement of Adalatherium hui (Mammalia, Gondwanatheria) from the Late Cretaceous of Madagascar: implications for allotherian relationships. J. Vertebr. Paleontol. 40:213–234.

Houde P, Dickson M, Camarena D. 2023. Basal Anseriformes from the Early Paleogene of North America and Europe. Diversity 15:233.

Houssaye A, Rage J-C, Torcida Fernández-Baldor F, Huerta P, Bardet N, Pereda Suberbiola X. 2013. A new varanoid squamate from the Early Cretaceous (Barremian–Aptian) of Burgos, Spain. Cretac. Res. 41:127–135.

Hughes LC, Ortí G, Huang Y, Sun Y, Baldwin CC, Thompson AW, Arcila D, Betancur-R R, Li C, Becker L, et al. 2018. Comprehensive phylogeny of ray-finned fishes (Actinopterygii) based on transcriptomic and genomic data. Proc. Natl. Acad. Sci. U. S. A. 115:6249–6254.

Hughes LC, Ortí G, Saad H, Li C, White WT, Baldwin CC, Crandall KA, Arcila D, Betancur-R R. 2021. Exon probe sets and bioinformatics pipelines for all levels of fish phylogenomics. Mol. Ecol. Resour. 21:816–833.

Hurley IA, Mueller RL, Dunn KA, Schmidt EJ, Friedman M, Ho RK, Prince VE, Yang Z, Thomas MG, Coates MI. 2006. A new time-scale for ray-finned fish evolution. Proc. R. Soc. B Biol. Sci. 274:489–498.

Huttenlocker AK, Grossnickle DM, Kirkland JI, Schultz JA, Luo Z-X. 2018. Late-surviving stem mammal links the lowermost Cretaceous of North America and Gondwana. Nature 558:108–112.

Irisarri I, Baurain D, Brinkmann H, Delsuc F, Sire J-Y, Kupfer A, Petersen J, Jarek M, Meyer A, Vences M, et al. 2017. Phylotranscriptomic consolidation of the jawed vertebrate timetree. *Nat*. Ecol. Evol. 1:1370–1378.

Irisarri I, Meyer A. 2016. The Identification of the Closest Living Relative(s) of Tetrapods: Phylogenomic Lessons for Resolving Short Ancient Internodes. Syst. Biol. 65:1057–1075.

Iwabe N, Hara Y, Kumazawa Y, Shibamoto K, Saito Y, Miyata T, Katoh K. 2005. Sister Group Relationship of Turtles to the Bird-Crocodilian Clade Revealed by Nuclear DNA–Coded Proteins. Mol. Biol. Evol. 22:810–813.

Jäger KRK, Luo Z-X, Martin T. 2020. Postcranial Skeleton of Henkelotherium guimarotae (Cladotheria, Mammalia) and Locomotor Adaptation. J. Mamm. Evol. 27:349–372.

Jarvis ED, Mirarab S, Aberer AJ, Li B, Houde P, Li C, Ho SYW, Faircloth BC, Nabholz B, Howard JT, et al. 2014. Whole-genome analyses resolve early branches in the tree of life of modern birds. Science 346:1320–1331.

Jones ME, Anderson CL, Hipsley CA, Müller J, Evans SE, Schoch RR. 2013. Integration of molecules and new fossils supports a Triassic origin for Lepidosauria (lizards, snakes, and tuatara). BMC Evol. Biol. 13:208.

Jones MEH, Benson RBJ, Skutschas P, Hill L, Panciroli E, Schmitt AD, Walsh SA, Evans SE. 2022. Middle Jurassic fossils document an early stage in salamander evolution. Proc. Natl. Acad. Sci. 119:e2114100119.

Jones MF, Beard KC, Simmons NB. 2024. Phylogeny and systematics of early Paleogene bats. J. Mamm. Evol. 31:18.

Joyce WG, Parham JF, Lyson TR, Warnock RCM, Donoghue PCJ. 2013. A Divergence Dating Analysis of Turtles Using Fossil Calibrations: An Example of Best Practices. J. Paleontol. 87:612–634.

Kemp A, Cavin L, Guinot G. 2017. Evolutionary history of lungfishes with a new phylogeny of post-Devonian genera. Palaeogeogr. Palaeoclimatol. Palaeoecol. 471:209–219.

Keqin G, Lianhai H. 1995. Iguanians From the Upper Cretaceous Djadochta Formation, Gobi Desert, China. J. Vertebr. Paleontol. 15:57–78.

Kimball RT, Oliveros CH, Wang N, White ND, Barker FK, Field DJ, Ksepka DT, Chesser RT, Moyle RG, Braun MJ, et al. 2019. A Phylogenomic Supertree of Birds. Diversity 11:109.

King B, Qiao T, Lee MSY, Zhu M, Long JA. 2017. Bayesian Morphological Clock Methods Resurrect Placoderm Monophyly and Reveal Rapid Early Evolution in Jawed Vertebrates. Syst. Biol. 66:499–516.

Kligman BT, Gee BM, Marsh AD, Nesbitt SJ, Smith ME, Parker WG, Stocker MR. 2023. Triassic stem caecilian supports dissorophoid origin of living amphibians. Nature 614:102–107.

Klug C, Coates M, Frey L, Greif M, Jobbins M, Pohle A, Lagnaoui A, Haouz WB, Ginter M. 2023. Broad snouted cladoselachian with sensory specialization at the base of modern chondrichthyans. Swiss J. Palaeontol. 142:2.

Korth WW. 1994. The Tertiary record of rodents in North America. Springer Science & Business Media Available from: https://books.google.com/books?hl=en&lr=&id=F4yxJ3M06TgC&oi=fnd&pg=PA3&ots=Xb6AGlvLbM&sig=zjbRD4YVyZvV_C9JhP5doS9NtZI

Kowallis BJ, Christiansen EH, Deino AL. 1991. Age of the Brushy Basin Member of the Morrison Formation, Colorado Plateau, western USA. Cretac. Res. 12:483–493.

Kramarz A, MacPhee R. 2022. Did some extinct South American native ungulates arise from an afrothere ancestor? A critical appraisal of Avilla and Mothé’s (2021) Sudamericungulata – Panameridiungulata hypothesis. J. Mamm. Evol. 30:1–11.

Krause DW, Hoffmann S, Hu Y, Wible JR, Rougier GW, Kirk EC, Groenke JR, Rogers RR, Rossie JB, Schultz JA, et al. 2020. Skeleton of a Cretaceous mammal from Madagascar reflects long-term insularity. Nature 581:421–427.

Krebs B. 1991. Das Skelett von Henkelotherium guimarotae gen. et sp. nov.(Eupantotheria, Mammalia) aus dem Oberen Jura von Portugal. Selbstverlag Fachbereich Geowissenschaften, FU Berlin Available from: https://e-docs.geo-leo.de/entities/publication/a0f63f76-85c0-4364-a06b-090b3a046f69

Kroth M, Trabucho-Alexandre JP, Pimenta MP, Vis G-J, Boever ED. 2024. Facies characterisation and stratigraphy of the upper Maastrichtian to lower Danian Maastricht Formation, South Limburg, the Netherlands. Neth. J. Geosci. 103:e13.

Ksepka DT, Bertelli S, Giannini NP. 2006. The phylogeny of the living and fossil Sphenisciformes (penguins). Cladistics 22:412–441.

Ksepka DT, Field DJ, Heath TA, Pett W, Thomas DB, Giovanardi S, Tennyson AJD. 2023. Largest-known fossil penguin provides insight into the early evolution of sphenisciform body size and flipper anatomy. J. Paleontol. 97:434–453.

Ksepka DT, Grande L, Mayr G. 2019. Oldest Finch-Beaked Birds Reveal Parallel Ecological Radiations in the Earliest Evolution of Passerines. Curr. Biol. 29:657–663.e1.

Ksepka DT, Stidham TA, Williamson TE. 2017. Early Paleocene landbird supports rapid phylogenetic and morphological diversification of crown birds after the K–Pg mass extinction. Proc. Natl. Acad. Sci. 114:8047–8052.

Kuhl H, Frankl-Vilches C, Bakker A, Mayr G, Nikolaus G, Boerno ST, Klages S, Timmermann B, Gahr M. 2021. An Unbiased Molecular Approach Using 3ʹ-UTRs Resolves the Avian Family-Level Tree of Life. Mol. Biol. Evol. 38:108–127.

Lambert O, Martínez-Cáceres M, Bianucci G, Celma CD, Salas-Gismondi R, Steurbaut E, Urbina M, Muizon C de. 2017. Earliest Mysticete from the Late Eocene of Peru Sheds New Light on the Origin of Baleen Whales. Curr. Biol. 27:1535–1541.e2.

Lee MSY. 1993. The Origin of the Turtle Body Plan: Bridging a Famous Morphological Gap. Science 261:1716–1720.

Lee MSY, Yates AM. 2018. Tip-dating and homoplasy: reconciling the shallow molecular divergences of modern gharials with their long fossil record. Proc. R. Soc. B Biol. Sci. 285:20181071.

Liu G, Pan Q, Dai Y, Wang X, Li M, Zhu P, Zhou X. 2024. Phylogenomics of Afrotherian mammals and improved resolution of extant Paenungulata. Mol. Phylogenet. Evol. 195:108047.

Liu J. 2021. Redescription of ‘Amyzon’ brevipinne and remarks on North American Eocene catostomids (Cypriniformes: Catostomidae). J. Syst. Palaeontol. 19:677–689.

Longrich NR, Bhullar B-AS, Gauthier JA. 2012. Mass extinction of lizards and snakes at the Cretaceous–Paleogene boundary. Proc. Natl. Acad. Sci. 109:21396–21401.

Longrich NR, Vinther J, Pyron RA, Pisani D, Gauthier JA. 2015. Biogeography of worm lizards (Amphisbaenia) driven by end-Cretaceous mass extinction. Proc. R. Soc. B Biol. Sci. [Internet]. Available from: https://royalsocietypublishing.org/doi/10.1098/rspb.2014.3034

López-Arbarello A. 2012. Phylogenetic Interrelationships of Ginglymodian Fishes (Actinopterygii: Neopterygii). PLOS ONE 7:e39370.

López-Arbarello A, Sferco E. 2018. Neopterygian phylogeny: the merger assay. R. Soc. Open Sci. 5:172337.

López-Conde OA, Sterli J, Alvarado-Ortega J, Chavarría-Arellano ML. 2017. A new platychelyid turtle (Pan-Pleurodira) from the Late Jurassic (Kimmeridgian) of Oaxaca, Mexico. Pap. Palaeontol. 3:161–174.

L Recinos M, Cantalice, Kleyton M., Caballero-Viñas, Carmen, and Alvarado-Ortega J. 2023. A new Mesozoic teleost of the subfamily Albulinae (Albuliformes: Albulidae) highlights the proto-Gulf of Mexico in the early diversification of extant bonefishes. J. Syst. Palaeontol. 21:2223797.

Lu J, Giles S, Friedman M, den Blaauwen JL, Zhu M. 2016. The Oldest Actinopterygian Highlights the Cryptic Early History of the Hyperdiverse Ray-Finned Fishes. Curr. Biol. 26:1602–1608.

Lu J, Giles S, Friedman M, Zhu M. 2017. A new stem sarcopterygian illuminates patterns of character evolution in early bony fishes. Nat. Commun. 8:1932.

Lu J, Zhu M, Ahlberg PE, Qiao T, Zhu Y, Zhao W, Jia L. 2016. A Devonian predatory fish provides insights into the early evolution of modern sarcopterygians. Sci. Adv. 2:e1600154.

Luo Z-X, Gatesy SM, Jenkins FA, Amaral WW, Shubin NH. 2015. Mandibular and dental characteristics of Late Triassic mammaliaform Haramiyavia and their ramifications for basal mammal evolution. Proc. Natl. Acad. Sci. U. S. A. 112:E7101–E7109.

Luo Z-X, Martin T. 2023. Mandibular and dental characteristics of the Late Jurassic mammal Henkelotherium guimarotae (Paurodontidae, Dryolestida). PalZ 97:569–619.

Luo Z-X, Meng Q-J, Ji Q, Liu D, Zhang Y-G, Neander AI. 2015. Evolutionary development in basal mammaliaforms as revealed by a docodontan. Science 347:760–764.

Maidment SCR, Balikova D, Muxworthy AR. 2017. Magnetostratigraphy of the Upper Jurassic Morrison Formation at Dinosaur National Monument, Utah, and Prospects for Using Magnetostratigraphy as a Correlative Tool in the Morrison Formation. In: Zeigler KE, Parker WG, editors. Terrestrial Depositional Systems. Elsevier. p. 279–302. Available from: https://www.sciencedirect.com/science/article/pii/B9780128032435000078

Mallik R, Wcisel DJ, Near TJ, Yoder JA, Dornburg A. 2025. Investigating the Impact of Whole-Genome Duplication on Transposable Element Evolution in Teleost Fishes. Genome Biol. Evol. 17:evae272.

Marivaux L, Vianey-Liaud M, Jaeger J-J. 2004. High-level phylogeny of early Tertiary rodents: dental evidence. Zool. J. Linn. Soc. 142:105–134.

Martínez RN, Simões TR, Sobral G, Apesteguía S. 2021. A Triassic stem lepidosaur illuminates the origin of lizard-like reptiles. Nature 597:235–238.

Marugán-Lobón J, Martín-Abad H, Buscalioni ÁD. 2023. The Las Hoyas Lagerstätte: a palaeontological view of an Early Cretaceous wetland. J. Geol. Soc. 180:jgs2022-079.

Mateus O, Puértolas-Pascual E, Callapez PM. 2019. A new eusuchian crocodylomorph from the Cenomanian (Late Cretaceous) of Portugal reveals novel implications on the origin of Crocodylia. Zool. J. Linn. Soc. 186:501–528.

Mayr G. 2015. The middle Eocene European “ratite” Palaeotis (Aves, Palaeognathae) restudied once more. Paläontol. Z. 89:503–514.

Mayr G, De Pietri VL, Paul Scofield R. 2017. A new fossil from the mid-Paleocene of New Zealand reveals an unexpected diversity of world’s oldest penguins. Sci. Nat. 104:9.

Mayr G, Pietri VLD, Love L, Mannering AA, Scofield RP. 2017. A well-preserved new mid-paleocene penguin (Aves, Sphenisciformes) from the Waipara Greensand in New Zealand. J. Vertebr. Paleontol. [Internet]. Available from: https://www.tandfonline.com/doi/full/10.1080/02724634.2017.1398169

Mayr G, Zelenkov N. 2021. Extinct crane-like birds (Eogruidae and Ergilornithidae) from the Cenozoic of Central Asia are indeed ostrich precursors. Ornithology 138:ukab048.

McInerney PL, Blokland JC, Worthy TH. 2024. Skull morphology of the enigmatic Genyornis newtoni Stirling and Zeitz, 1896 (Aves, Dromornithidae), with implications for functional morphology, ecology, and evolution in the context of Galloanserae. Hist. Biol. 36:1093–1165.

Melo BF, Sidlauskas BL, Near TJ, Roxo FF, Ghezelayagh A, Ochoa LE, Stiassny MLJ, Arroyave J, Chang J, Faircloth BC, et al. 2022. Accelerated Diversification Explains the Exceptional Species Richness of Tropical Characoid Fishes. Syst. Biol. 71:78–92.

Melstrom KM, Irmis RB. 2019. Repeated Evolution of Herbivorous Crocodyliforms during the Age of Dinosaurs. Curr. Biol. 29:2389–2395.e3.

Meredith RW, Janečka JE, Gatesy J, Ryder OA, Fisher CA, Teeling EC, Goodbla A, Eizirik E, Simão TLL, Stadler T, et al. 2011. Impacts of the Cretaceous Terrestrial Revolution and KPg extinction on mammal diversification. Science 334:521–524.

Mertz DF, Renne PR. 2005. A numerical age for the Messel fossil deposit (UNESCO World Heritage Site derived from^4^0Ar/^3^9Ar dating on a basaltic rock fragment. Cour.-FORSCHUNGSINSTITUT SENCKENBERG 255:67.

Meyer A, Schloissnig S, Franchini P, Du K, Woltering JM, Irisarri I, Wong WY, Nowoshilow S, Kneitz S, Kawaguchi A, et al. 2021. Giant lungfish genome elucidates the conquest of land by vertebrates. Nature 590:284–289.

Meyer D, Brownstein CD, Jenkins KM, Gauthier JA. 2023. A Morrison stem gekkotan reveals gecko evolution and Jurassic biogeography. Proc. R. Soc. B [Internet]. Available from: https://royalsocietypublishing.org/doi/10.1098/rspb.2023.2284

Milàn J, Lucas S, Lockley M, Spielmann J, Schwimmer D. 2010. BITE MARKS OF THE GIANT CROCODYLIAN DEINOSUCHUS ON LATE CRETACEOUS (CAMPANIAN) BONES. 51.

Miller EC, Martinez CM, Friedman ST, Wainwright PC, Price SA, Tornabene L. 2022. Alternating regimes of shallow and deep-sea diversification explain a species-richness paradox in marine fishes. Proc. Natl. Acad. Sci. 119:e2123544119.

Miller KG, Sugarman PJ, Browning JV, Kominz MA, Olsson RK, Feigenson MD, Hernandez JC. 2004. Upper Cretaceous sequences and sea-level history, New Jersey Coastal Plain. Geol. Soc. Am. Bull. 116:368–393.

Mirarab S, Rivas-González I, Feng S, Stiller J, Fang Q, Mai U, Hickey G, Chen G, Brajuka N, Fedrigo O, et al. 2024. A region of suppressed recombination misleads neoavian phylogenomics. Proc. Natl. Acad. Sci. 121:e2319506121.

Mohler BF, McDonald AT, Wolfe DG. 2021. First remains of the enormous alligatoroid Deinosuchus from the Upper Cretaceous Menefee Formation, New Mexico. PeerJ 9:e11302.

Monsch KA, Bannikov AF. 2011. New taxonomic synopses and revision of the scombroid fishes (Scombroidei, Perciformes), including billfishes, from the Cenozoic of territories of the former USSR. Earth Environ. Sci. Trans. R. Soc. Edinb. 102:253–300.

Musilova Z, Cortesi F, Matschiner M, Davies WIL, Patel JS, Stieb SM, de Busserolles F, Malmstrøm M, Tørresen OK, Brown CJ, et al. 2019. Vision using multiple distinct rod opsins in deep-sea fishes. Science 364:588–592.

Musser G, Clarke JA. 2024. A new Paleogene fossil and a new dataset for waterfowl (Aves: Anseriformes) clarify phylogeny, ecological evolution, and avian evolution at the K-Pg Boundary. PLOS ONE 19:e0278737.

Near TJ, Dornburg A, Friedman M. 2014. Phylogenetic relationships and timing of diversification in gonorynchiform fishes inferred using nuclear gene DNA sequences (Teleostei: Ostariophysi). Mol. Phylogenet. Evol. 80:297–307.

Near TJ, Eytan RI, Dornburg A, Kuhn KL, Moore JA, Davis MP, Wainwright PC, Friedman M, Smith WL. 2012. Resolution of ray-finned fish phylogeny and timing of diversification. Proc. Natl. Acad. Sci. 109:13698–13703.

Near TJ, Thacker CE. 2024. Phylogenetic classification of living and fossil ray-finned fishes (Actinopterygii). Bull. Peabody Mus. Nat. Hist. 65:3–302.

Nesbitt SJ, Butler RJ, Ezcurra MD, Barrett PM, Stocker MR, Angielczyk KD, Smith RMH, Sidor CA, Niedźwiedzki G, Sennikov AG, et al. 2017. The earliest bird-line archosaurs and the assembly of the dinosaur body plan. Nature 544:484–487.

Nesbitt SJ, Clarke JA. 2016. The Anatomy and Taxonomy of the Exquisitely Preserved Green River Formation (Early Eocene) Lithornithids (Aves) and the Relationships of Lithornithidae. Bull. Am. Mus. Nat. Hist. 2016:1–91.

Nesbitt SJ, Patellos E, Kammerer CF, Ranivoharimanana L, Wyss AR, Flynn JJ. 2023. The earliest-diverging avemetatarsalian: a new osteoderm-bearing taxon from the Triassic (?Earliest Late Triassic) of Madagascar and the composition of avemetatarsalian assemblages prior to the radiation of dinosaurs. Zool. J. Linn. Soc. 199:327–353.

Ni X, Gebo DL, Dagosto M, Meng J, Tafforeau P, Flynn JJ, Beard KC. 2013. The oldest known primate skeleton and early haplorhine evolution. Nature 498:60–64.

Ogg JG, Ogg GM, Gradstein FM. 2016. 14 - Paleogene. In: Ogg JG, Ogg GM, Gradstein FM, editors. A Concise Geologic Time Scale. Elsevier. p. 187–201. Available from: https://www.sciencedirect.com/science/article/pii/B9780444594679000145

O’Leary MA, Bloch JI, Flynn JJ, Gaudin TJ, Giallombardo A, Giannini NP, Goldberg SL, Kraatz BP, Luo Z-X, Meng J, et al. 2013. The placental mammal ancestor and the post-K-Pg radiation of placentals. Science 339:662–667.

Olsen PE. 1984. The skull and pectoral girdle of the parasemionotid fish Watsonulus eugnathoides from the Early Triassic Sakamena Group of Madagascar, with comments on the relationships of the holostean fishes. J. Vertebr. Paleontol. 4:481–499.

Onary S, Hsiou AS, Lee MSY, Palci A. 2021. Redescription, taxonomy and phylogenetic relationships of Boavus Marsh, 1871 (Serpentes: Booidea) from the early–middle Eocene of the USA. J. Syst. Palaeontol. 19:1601–1622.

Palci A, Onary S, Lee MSY, Smith KT, Wings O, Rabi M, Georgalis GL. 2024. A new booid snake from the Eocene (Lutetian) Konservat-Lagerstätte of Geiseltal, Germany, and a new phylogenetic analysis of Booidea. Zool. J. Linn. Soc. 202:zlad179.

Parey E, Louis A, Montfort J, Bouchez O, Roques C, Iampietro C, Lluch J, Castinel A, Donnadieu C, Desvignes T, et al. 2023. Genome structures resolve the early diversification of teleost fishes. Science 379:572–575.

Paterson RS, Rybczynski N, Kohno N, Maddin HC. 2020. A Total Evidence Phylogenetic Analysis of Pinniped Phylogeny and the Possibility of Parallel Evolution Within a Monophyletic Framework. Front. Ecol. Evol. [Internet] 7. Available from: https://www.frontiersin.org/journals/ecology-and-evolution/articles/10.3389/fevo.2019.00457/full

Phillips MJ, Fruciano C. 2018. The soft explosive model of placental mammal evolution. BMC Evol. Biol. 18:104.

Poust A, Boessenecker R. 2018. Expanding the geographic and geochronologic range of early pinnipeds: new specimens of Enaliarctos from Northern California and Oregon. Acta Palaeontol. Pol. [Internet] 63. Available from: http://www.app.pan.pl/article/item/app003992017.html

Poyato-Ariza FJ. 1996a. The phylogenetic relationships of Rubiesichthys gregalis and Gordichthys conquensis (Ostariophysi, Chanidae), from the Early Cretaceous of Spain. Mesoz. Fishes—Systematics Paleoecol. Verl. Dr Fredrich Pfeil Munich Ger.:329–348.

Poyato-Ariza FJ. 1996b. A revision of Rubiesichthys gregalis WENZ 1984 (Ostariophysi, Gonorynchiformes), from the Early Cretaceous of Spain. Syst Paleoecol 1984:329–348.

Prokofiev AM. 2002. A remarkable new genus of Carangidae fron1 the Upper Paleocene of Turkmenistan (Osteichthyes: Perciformes). Available from: https://www.zin.ru/Journals/zsr/content/2002/zr_2002_11_1_Prokofiev_2.pdf

Prothero D, Bitboul C, Moore G, Niem A, Section AA of PGP, Section GS of AC. 2001. Magnetic stratigraphy of the Pacific Coast Cenozoic: a symposium volume based on Proceedings of the 1997 Pacific Section AAPG-SEPM Meeting, Bakersfield, California, and the 2001 Pacific Section AAPG-SEPM-Cordilleran Section GSA Meeting. Fullerton, CA: Pacific Section SEPM

Prum RO, Berv JS, Dornburg A, Field DJ, Townsend JP, Lemmon EM, Lemmon AR. 2015. A comprehensive phylogeny of birds (Aves) using targeted next-generation DNA sequencing. Nature 526:569–573.

Pyron RA, Burbrink FT, Wiens JJ. 2013. A phylogeny and revised classification of Squamata, including 4161 species of lizards and snakes. BMC Evol. Biol. 13:93.

Qiao T, King B, Long JA, Ahlberg PE, Zhu M. 2016. Early Gnathostome Phylogeny Revisited: Multiple Method Consensus. PLoS ONE 11:e0163157.

Qu Q, Zhu M, Wang W. 2013. Scales and Dermal Skeletal Histology of an Early Bony Fish Psarolepis romeri and Their Bearing on the Evolution of Rhombic Scales and Hard Tissues. PLOS ONE 8:e61485.

Reddy S, Kimball RT, Pandey A, Hosner PA, Braun MJ, Hackett SJ, Han K-L, Harshman J, Huddleston CJ, Kingston S, et al. 2017. Why Do Phylogenomic Data Sets Yield Conflicting Trees? Data Type Influences the Avian Tree of Life more than Taxon Sampling. Syst. Biol. 66:857–879.

dos Reis M, Donoghue PCJ, Yang Z. 2014. Neither phylogenomic nor palaeontological data support a Palaeogene origin of placental mammals. Biol. Lett. 10:20131003.

dos Reis M, Inoue J, Hasegawa M, Asher RJ, Donoghue PCJ, Yang Z. 2012. Phylogenomic datasets provide both precision and accuracy in estimating the timescale of placental mammal phylogeny. Proc. Biol. Sci. 279:3491–3500.

Renesto S, Bernardi M. 2014. Redescription and phylogenetic relationships of Megachirella wachtleri Renesto et Posenato, 2003 (Reptilia, Diapsida). Paläontol. Z. 88:197–210.

Ribeiro AC, Poyato-Ariza FJ, Bockmann FA, Carvalho MR de. 2018. Phylogenetic relationships of Chanidae (Teleostei: Gonorynchiformes) as impacted by Dastilbe moraesi, from the Sanfranciscana basin, Early Cretaceous of Brazil. Neotropical Ichthyol. 16:e180059.

Rietbergen TB, van den Hoek Ostende LW, Aase A, Jones MF, Medeiros ED, Simmons NB. 2023. The oldest known bat skeletons and their implications for Eocene chiropteran diversification. PLOS ONE 18:e0283505.

Rio JP, Mannion PD. 2021. Phylogenetic analysis of a new morphological dataset elucidates the evolutionary history of Crocodylia and resolves the long-standing gharial problem. PeerJ 9:e12094.

Russell DE. 1987. The Paleogene of Asia: mammals and stratigraphy. Mém. Muséum Natl. Hist. Nat. Sci. Terre 52:1–488.

Santaquiteria A, Siqueira AC, Duarte-Ribeiro E, Carnevale G, White WT, Pogonoski JJ, Baldwin CC, Ortí G, Arcila D, Ricardo B-R. 2021. Phylogenomics and Historical Biogeography of Seahorses, Dragonets, Goatfishes, and Allies (Teleostei: Syngnatharia): Assessing Factors Driving Uncertainty in Biogeographic Inferences. Syst. Biol. 70:1145–1162.

Santini F, Tyler JC. 2003. A phylogeny of the families of fossil and extant tetraodontiform fishes (Acanthomorpha, Tetraodontiformes), Upper Cretaceous to Recent. Zool. J. Linn. Soc. 139:565–617.

Scarpetta SG. 2024. A Palaeogene stem crotaphytid (Aciprion formosum) and the phylogenetic affinities of early fossil pleurodontan iguanians. R. Soc. Open Sci. [Internet]. Available from: https://royalsocietypublishing.org/doi/10.1098/rsos.221139

Schartl M, Woltering JM, Irisarri I, Du K, Kneitz S, Pippel M, Brown T, Franchini P, Li J, Li M, et al. 2024. The genomes of all lungfish inform on genome expansion and tetrapod evolution. Nature:1–8.

Schoch RR, Sues H-D. 2018. A new lepidosauromorph reptile from the Middle Triassic (Ladinian) of Germany and its phylogenetic relationships. J. Vertebr. Paleontol. 38:e1444619.

Schoch RR, Werneburg R, Voigt S. 2020. A Triassic stem-salamander from Kyrgyzstan and the origin of salamanders. Proc. Natl. Acad. Sci. 117:11584–11588.

Schwimmer DR. 2002. King of the crocodylians: the paleobiology of Deinosuchus. Indiana University Press Available from: https://books.google.com/books?hl=en&lr=&id=0OsPJnC4CCwC&oi=fnd&pg=PR5&dq=King+of+the+Crocodylians:+The+Paleobiology+of+Deinosuchus+(Life+of+the+Past)&ots=8MNNMJwHvk&sig=pvFH3mGhJyykN1opYJWPTcqYfqY

Seiffert ER. 2007. A new estimate of afrotherian phylogeny based on simultaneous analysis of genomic, morphological, and fossil evidence. BMC Evol. Biol. 7:224.

Seiffert ER, Heritage S, de Vries D, Sallam HM, Vitek NS, Aoron E, Princehouse P. 2025. Oldest record of a crown anomaluroid rodent from sub-Saharan Africa: a new genus and species from the early Oligocene Topernawi Formation of northern Kenya. Hist. Biol. 37:1568–1578.

Shan Y, Gras R. 2011. 43 genes support the lungfish-coelacanth grouping related to the closest living relative of tetrapods with the Bayesian method under the coalescence model. BMC Res. Notes 4:49.

Simmons NB, Geisler JH. 1998. Phylogenetic Relationships of Icaronycteris, Archaeonycteris, Hassianycteris, and Palaeochiropteryx to Extant Bat Lineages, with Comments on the Evolution of Echolocation and Foraging Strategies in Microchiroptera. American Museum of Natural History

Simões TR, Caldwell MW, Pierce SE. 2020. Sphenodontian phylogeny and the impact of model choice in Bayesian morphological clock estimates of divergence times and evolutionary rates. BMC Biol. 18:191.

Simões TR, Caldwell MW, Tałanda M, Bernardi M, Palci A, Vernygora O, Bernardini F, Mancini L, Nydam RL. 2018. The origin of squamates revealed by a Middle Triassic lizard from the Italian Alps. Nature 557:706–709.

Simões TR, Kammerer CF, Caldwell MW, Pierce SE. 2022. Successive climate crises in the deep past drove the early evolution and radiation of reptiles. Sci. Adv. 8:eabq1898.

Singhal S, Colston TJ, Grundler MR, Smith SA, Costa GC, Colli GR, Moritz C, Pyron RA, Rabosky DL. 2021. Congruence and Conflict in the Higher-Level Phylogenetics of Squamate Reptiles: An Expanded Phylogenomic Perspective. Syst. Biol. 70:542–557.

Siu-Ting K, Torres-Sánchez M, San Mauro D, Wilcockson D, Wilkinson M, Pisani D, O’Connell MJ, Creevey CJ. 2019. Inadvertent Paralog Inclusion Drives Artifactual Topologies and Timetree Estimates in Phylogenomics. Mol. Biol. Evol. 36:1344–1356.

Slack KE, Jones CM, Ando T, Harrison GL (Abby), Fordyce RE, Arnason U, Penny D. 2006. Early Penguin Fossils, Plus Mitochondrial Genomes, Calibrate Avian Evolution. Mol. Biol. Evol. 23:1144–1155.

Slater GJ. 2015. Iterative adaptive radiations of fossil canids show no evidence for diversity-dependent trait evolution. Proc. Natl. Acad. Sci. 112:4897–4902.

Smith ME, Carroll AR, Singer BS. 2008. Synoptic reconstruction of a major ancient lake system: Eocene Green River Formation, western United States. GSA Bull. 120:54–84.

Sobral G, Simões TR, Schoch RR. 2020. A tiny new Middle Triassic stem-lepidosauromorph from Germany: implications for the early evolution of lepidosauromorphs and the Vellberg fauna. Sci. Rep. 10:2273.

Spiekman SNF, Fraser NC, Scheyer TM. 2021. A new phylogenetic hypothesis of Tanystropheidae (Diapsida, Archosauromorpha) and other “protorosaurs”, and its implications for the early evolution of stem archosaurs. PeerJ 9:e11143.

Springer MS, Murphy WJ, Eizirik E, O’Brien SJ. 2003. Placental mammal diversification and the Cretaceous–Tertiary boundary. Proc. Natl. Acad. Sci. 100:1056–1061.

Stiller J, Feng S, Chowdhury A-A, Rivas-González I, Duchêne DA, Fang Q, Deng Y, Kozlov A, Stamatakis A, Claramunt S, et al. 2024. Complexity of avian evolution revealed by family-level genomes. Nature:1–3.

Straube N, Li C, Mertzen M, Yuan H, Moritz T. 2018. A phylogenomic approach to reconstruct interrelationships of main clupeocephalan lineages with a critical discussion of morphological apomorphies. BMC Evol. Biol. 18:158.

Streicher JW, Wiens JJ. 2017. Phylogenomic analyses of more than 4000 nuclear loci resolve the origin of snakes among lizard families. Biol. Lett. 13:20170393.

Takezaki N. 2018. Global Rate Variation in Bony Vertebrates. Genome Biol. Evol. 10:1803–1815.

Takezaki N. 2021. Resolving the Early Divergence Pattern of Teleost Fish Using Genome-Scale Data. Genome Biol. Evol. 13:evab052.

Takezaki N, Nishihara H. 2017. Support for Lungfish as the Closest Relative of Tetrapods by Using Slowly Evolving Ray-Finned Fish as the Outgroup. Genome Biol. Evol. 9:93–101.

Tałanda M, Fernandez V, Panciroli E, Evans SE, Benson RJ. 2022. Synchrotron tomography of a stem lizard elucidates early squamate anatomy. Nature 611:99–104.

Tambussi CP, Degrange FJ, De Mendoza RS, Sferco E, Santillana S. 2019. A stem anseriform from the early Palaeocene of Antarctica provides new key evidence in the early evolution of waterfowl. Zool. J. Linn. Soc. 186:673–700.

Tarver JE, dos Reis M, Mirarab S, Moran RJ, Parker S, O’Reilly JE, King BL, O’Connell MJ, Asher RJ, Warnow T, et al. 2016. The Interrelationships of Placental Mammals and the Limits of Phylogenetic Inference. Genome Biol. Evol. 8:330–344.

Thomas DB, Tennyson AJD, Marx FG, Ksepka DT. 2023. Pliocene fossils support a New Zealand origin for the smallest extant penguins. J. Paleontol. 97:711–721.

Thomas DB, Tennyson AJD, Scofield RP, Heath TA, Pett W, Ksepka DT. 2020. Ancient crested penguin constrains timing of recruitment into seabird hotspot. Proc. R. Soc. B Biol. Sci. 287:20201497.

Thompson AW, Hawkins MB, Parey E, Wcisel DJ, Ota T, Kawasaki K, Funk E, Losilla M, Fitch OE, Pan Q, et al. 2021. The bowfin genome illuminates the developmental evolution of ray-finned fishes. Nat. Genet. 53:1373–1384.

Title PO, Singhal S, Grundler MC, Costa GC, Pyron RA, Colston TJ, Grundler MR, Prates I, Stepanova N, Jones MEH, et al. 2024. The macroevolutionary singularity of snakes. Science 383:918–923.

Tomiya S. 2011. A New Basal Caniform (Mammalia: Carnivora) from the Middle Eocene of North America and Remarks on the Phylogeny of Early Carnivorans. PLOS ONE 6:e24146.

Tomiya S, Tseng ZJ. 2016. Whence the beardogs? Reappraisal of the Middle to Late Eocene ‘Miacis’ from Texas, USA, and the origin of Amphicyonidae (Mammalia, Carnivora). R. Soc. Open Sci. 3:160518.

Torres CR, Norell MA, Clarke JA. 2021. Bird neurocranial and body mass evolution across the end-Cretaceous mass extinction: The avian brain shape left other dinosaurs behind. Sci. Adv. 7:eabg7099.

Townsend TM, Larson A, Louis E, Macey JR. 2004. Molecular Phylogenetics of Squamata: The Position of Snakes, Amphisbaenians, and Dibamids, and the Root of the Squamate Tree. Syst. Biol. 53:735–757.

Trujillo K, Kowallis B. 2015. Recalibrated legacy 40Ar/39Ar ages for the Upper Jurassic Morrison Formation, Western Interior, U.S.A. Geol. Intermt. West 2:1–8.

Tyler JC, Santini F. 2005. A phylogeny of the fossil and extant zeiform-like fishes, Upper Cretaceous to Recent, with comments on the putative zeomorph clade (Acanthomorpha). Zool. Scr. 34:157–175.

Upham NS, Esselstyn JA, Jetz W. 2021. Molecules and fossils tell distinct yet complementary stories of mammal diversification. Curr. Biol. 31:4195–4206.e3.

Velazco PM, Buczek AJ, Hoffman E, Hoffman DK, O’Leary MA, Novacek MJ. 2022. Combined data analysis of fossil and living mammals: a Paleogene sister taxon of Placentalia and the antiquity of Marsupialia. Cladistics 38:359–373.

Vianey-Liaud M, Marivaux L. 2021. The beginning of the adaptive radiation of Theridomorpha (Rodentia) in Western Europe: morphological and phylogenetic analyses of early and middle Eocene taxa; implications for systematics. Palaeovertebrata 44:2–e2.

Vidal N, Hedges SB. 2005. The phylogeny of squamate reptiles (lizards, snakes, and amphisbaenians) inferred from nine nuclear protein-coding genes. C. R. Biol. 328:1000–1008.

Vlachos E. 2018. A Review of the Fossil Record of North American Turtles of the Clade Pan-Testudinoidea. Bull. Peabody Mus. Nat. Hist. 59:3–94.

Wang H, Wang Y. 2023. Middle ear innovation in Early Cretaceous eutherian mammals. Nat. Commun. 14:6831.

Wang K, Wang J, Zhu C, Yang L, Ren Y, Ruan J, Fan G, Hu J, Xu W, Bi X, et al. 2021. African lungfish genome sheds light on the vertebrate water-to-land transition. Cell 184:1362–1376.e18.

Wang X, Tedford R. 1996. Canidae. In: p. 433–452.

Wang Z, Pascual-Anaya J, Zadissa A, Li W, Niimura Y, Huang Z, Li C, White S, Xiong Z, Fang D, et al. 2013. The draft genomes of soft-shell turtle and green sea turtle yield insights into the development and evolution of the turtle-specific body plan. Nat. Genet. 45:701–706.

Wescott WA, Diggens JN. 1998. Depositional history and stratigraphical evolution of the Sakamena group (Middle Karoo Supergroup) in the southern Morondava Basin, Madagascar. J. Afr. Earth Sci. 27:461–479.

Wesley-Hunt GD, Flynn JJ. 2005. Phylogeny of the carnivora: Basal relationships among the carnivoramorphans, and assessment of the position of ‘miacoidea’ relative to carnivora. J. Syst. Palaeontol. 3:1–28.

Wesley-Hunt GD, Werdelin L. 2005. Basicranial morphology and phylogenetic position of the upper Eocene carnivoramorphan Quercygale. Acta Palaeontol. Pol. 50:837.

Wible JR, Rougier GW, Novacek MJ, Asher RJ. 2007. Cretaceous eutherians and Laurasian origin for placental mammals near the K/T boundary. Nature 447:1003–1006.

Wilson Mantilla GP, Chester SGB, Clemens WA, Moore JR, Sprain CJ, Hovatter BT, Mitchell WS, Mans WW, Mundil R, Renne PR. 2021. Earliest Palaeocene purgatoriids and the initial radiation of stem primates. R. Soc. Open Sci. 8:210050.

Worthy TH, Degrange FJ, Handley WD, Lee MSY. 2017. The evolution of giant flightless birds and novel phylogenetic relationships for extinct fowl (Aves, Galloanseres). R. Soc. Open Sci. 4:170975.

Xu G-H. 2019. Osteology and phylogeny of Robustichthys luopingensis, the largest holostean fish in the Middle Triassic. PeerJ 7:e7184.

Xu G-H, Zhao L-J, Coates MI. 2014. The oldest ionoscopiform from China sheds new light on the early evolution of halecomorph fishes. Biol. Lett. 10:20140204.

Zaher H, Smith KT. 2020. Pythons in the Eocene of Europe reveal a much older divergence of the group in sympatry with boas. Biol. Lett. 16:20200735.

Zhang C, Rabiee M, Sayyari E, Mirarab S. 2018. ASTRAL-III: polynomial time species tree reconstruction from partially resolved gene trees. BMC Bioinformatics 19:153.

Zhao W, Zhang X, Jia G, Shen Y, Zhu M. 2021. The Silurian-Devonian boundary in East Yunnan (South China) and the minimum constraint for the lungfish-tetrapod split. Sci. China Life Sci. 64:1.

Zheng Y, Wiens JJ. 2016. Combining phylogenomic and supermatrix approaches, and a time-calibrated phylogeny for squamate reptiles (lizards and snakes) based on 52 genes and 4162 species. Mol. Phylogenet. Evol. 94:537–547.

Zhu M, Yu X, Ahlberg PE. 2001. A primitive sarcopterygian fish with an eyestalk. Nature 410:81–84.

Zhu M, Yu X, Ahlberg PE, Choo B, Lu J, Qiao T, Qu Q, Zhao W, Jia L, Blom H, et al. 2013. A Silurian placoderm with osteichthyan-like marginal jaw bones. Nature 502:188–193.

Zhu M, Zhao W, Jia L, Lu J, Qiao T, Qu Q. 2009. The oldest articulated osteichthyan reveals mosaic gnathostome characters. Nature 458:469–474.

Zhu Y, Giles S, Young GC, Hu Y, Bazzi M, Ahlberg PE, Zhu M, Lu J. 2021. Endocast and Bony Labyrinth of a Devonian “Placoderm” Challenges Stem Gnathostome Phylogeny. Curr. Biol. 31:1112–1118.e4.

